# Targeted protein modification with an antibody-based system

**DOI:** 10.1101/2023.07.28.551006

**Authors:** Oded Rimon, Juraj Konc, Montader Ali, Vaidehi Roy Chowdhury, Inga Černauskienė, Pietro Sormanni, Gonçalo J. L. Bernardes, Michele Vendruscolo

## Abstract

The chemical modification of proteins is one of the major mechanisms used to regulate the properties and functions of these macromolecules in the cell. It is therefore of great interest to develop tools to exploit this type of modifications for applications in molecular biology, medicine and biotechnology. Here we present a method of using antibodies to perform post-translational covalent modifications of endogenous proteins in complex environments by exploiting proximity-driven chemistry. The method is based on the ability of antibodies to hold a weakly reactive group close to its intended site of reaction by binding the target protein on a nearby epitope. We characterise this approach by modifying the green fluorescent protein in increasingly complex environments, and illustrate its applicability by targeting the disease-associated protein beta-2 microglobulin.

---

Chemically modifying a specific protein in a complex environment, such as that within a cell, is highly challenging but holds great promise. As well as being versatile labels for monitoring the behaviour of proteins in their natural milieu, post-translational modifications (PTMs) can effectively influence protein structure and function [1]. This phenomenon has long been exploited pharmacologically: covalent inhibitors such as aspirin bind, or attach a covalent payload, to the active sites of enzymes. Following this paradigm, rationally designed targeted covalent inhibitors (TCIs) adapt non-covalent ligands to react with nucleophilic residues adjacent to their binding site [2]. However, proteins that lack ligand binding sites are largely inaccessible to both traditional drugs and TCIs [3]. Notably, the very reasons that make these proteins difficult to target with traditional small-molecule drugs make them near-ideal candidates for modulation *via* post-translational modification [4].

It is therefore important to develop technologies for targeted post-translational modification (TPTM) – the covalent modification of endogenous proteins in complex environments at a specific surface-exposed site. To be therapeutically relevant, one would like these technologies to fulfil several requirements: (1) To engage endogenous targets, which are genetically and chemically unmodified; (2) To bind chosen sites, without relying on proximity to naturally occurring binding sites on the target; (3) To be traceless, i.e., to break up, with the non-covalent binding moiety dissociating and only leaving behind a covalent payload [5] (traceless reactions uncouple the reactivity and selectivity of the reaction from the identity of the covalent payload, uncovering a vast chemical derivatisation landscape and leaving no junk atoms behind that may cause haptenisation and other unpredictable behaviours [2]); (4) To be selective for their targets, in order to minimize adverse effects due to non-specific modification of other macromolecules and small-molecule nucleophiles; and (5) To be site selective, to reduce the immunogenicity of the modified product and produce consistent and uniform effects [6].

Despite many advances in establishing methods for the *in vivo* PTM of proteins, to our knowledge no technique or compound exists at this time that fulfils all the requirements listed above. Here, we present an approach for developing such tools, and demonstrate its potential and its limitations by targeting the green fluorescent protein (GFP) in environments of increasing complexity. We then apply the lessons learned from this model system to modify the disease-associated protein beta-2 microglobulin (β2m) in complex mixtures.

In essence, the targeted post-translational modifiers that we introduce in this work are hybrid molecules consisting of a high-affinity antibody and a synthetic linker featuring a weakly reactive group susceptible to induction through the proximity effect.

## Results

### PEABS: an antibody-based design for targeted post-translational modification

A key feature of TPTM is selectivity. In complex biological environments, no chemical functionality can be expected to be unique to one macromolecule, and reactive chemical groups non-specifically modify almost inevitably off-target proteins at least to some extent. Therefore, weak reactivity is required for any ligand-directed chemistry, warhead for TCI or reactive group for TPTM. On the other hand, to accomplish its task, the reactive moiety of any covalent drug must act during the period of time that its non-covalent moiety is bound, i.e. within its residence time [2].

A way to bridge these two seemingly contradictory requirements is through the exploitation of the proximity effect. Non-covalent binding of a covalent drug to a macromolecule reduces its translational and rotational entropy. As a result, the activation energy for its covalent reaction with a nearby functional group loses a portion of its entropic penalty and is therefore decreased. This effect can account for more than a three orders of magnitude acceleration in reaction rates [7].

To achieve the long residence times required for TPTM, one can increase the interaction surface area. This is often challenging with small-molecule drugs, but less so with biological macromolecules – and particularly antibodies [8]. Antibodies bind to unmodified endogenous proteins, fulfilling the first requirement of TPTM. With methods ranging from immunisation to *in silico* rational design, antibodies can be developed to target any unique feature on the surface of a target protein, with high affinity and selectivity [9]. This fulfils the second requirement for TPTM, allowing one to create a TPTM agent for a desired target residue, rather than choosing the target residue based on its proximity to a known, well-structured binding pocket. Thanks to the versatility of antibodies, the choice of a target epitope can be made with additional considerations in mind. The first of these would be the choice of a naturally occurring nucleophile on the target protein to be used for substitution with the payload. While cysteine is a common choice in TCI design due to its high nucleophilicity and low abundance, targeting it would limit the range to surface-exposed, free-form cysteine residues. Primary amines, on the other hand, are common on the surfaces of proteins. As well as the N-terminal amine (on non-acetylated proteins), lysine residues constitute 6% of the human proteome, and they are quite nucleophilic despite being protonated at physiological pH [10], with some 9000 lysine residues estimated as available for nucleophilic attack in human proteomes [11].

With these observations in mind, we opted for a design in which an antibody is produced recombinantly with an engineered cysteine residue at a carefully selected position (see below). A synthetic linker is then conjugated to the engineered cysteine *in vitro* to form the active hybrid species. Upon the binding of this species to its target antigen, its activated ester moiety is positioned precisely to transfer the payload on to one amine group of the target, forming an amide link between them and releasing the spent antibody, which can then be cleared without loss of the PTM (Figure 1a).

**Figure 1.**
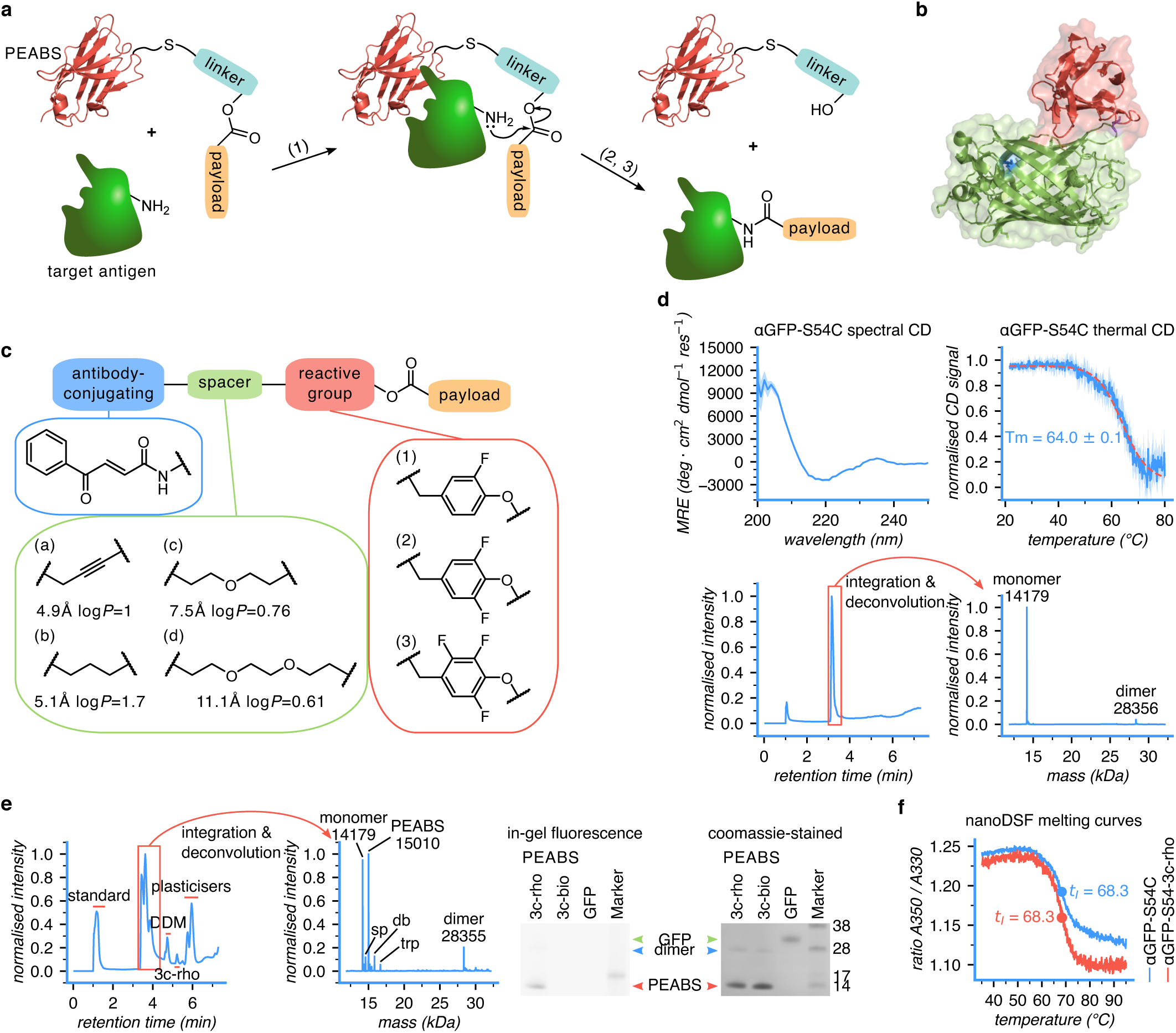
Design of a protein-editing antibody-based system (PEABS). **(a)** General scheme of a PEABS reaction. A linker featuring an amine-reactive group is bioconjugated to an antibody prior to the reaction; the non-covalent binding of antibody and antigen brings the reactive group in proximity to a primary amine group of the target (1); a nucleophilic attack transfers the payload onto the target, with the linker leaving (2); the dissociation of the spent PEABS leaves no trace on the target except the (iso)peptide-bound payload (3). **(b)** Structure of the antibody–antigen pair used in this work, which consists of GFP (green, with lysine 41 in blue) and anti-GFP nanobody (red with residue 54, chosen for linker conjugation, in purple) [PDB: 3OGO][12]. **(c)** Schematic structure of a loaded PEABS linker. Linkers are named ***#l*** according to their reactive group number (#) and spacer letter (*l*), e.g. **2b**. A benzoylacrylic-based Michael acceptor (blue) is responsible for conjugation with the engineered cysteine on the antibody. Spacers (green) connect it to a reactive group; values under each spacer indicate its fully extended length and partition coefficient as predicted by ChemDraw. Reactive groups (red) are responsible for carrying the payload and transferring it tracelessly to the target; different fluorophenols were employed for this purpose. **(d)** Characterisation of αGFP-S54C, the cysteine mutant used as basis for PEABS systems, using circular dichroism (CD), heat-induced denaturation monitored by CD, and intact protein LC-MS. **(e)** Confirmation of bioconjugation of **3c-rho** to αGFP-S54C using LC-MS and SDS-PAGE; 40% of the nanobody was converted, with very low residual free linker. αGFP-S54C *exp.* 14179 *obs.* 14179; PEABS *exp.* 15010 *obs.* 15010; sp (spent PEABS) *exp.* 14586 *obs.* 14586; dbl (PEABS + **3c-rho**) *exp.* 15841 *obs.* 15842; trp (PEABS + 2 × **3c-rho**) *exp.* 16673 *obs.* 16676. **(f)** nanoDSF profiles confirm minimal destabilisation of the nanobody structure following bioconjugation.

Referring to its ability to post-translationally modify proteins, as well as extending them by forming new peptide bonds at the N-terminus, we termed this design the Protein-Editing Antibody-Based System, or *PEABS*.

### Design and characterisation of GFP-modifying PEABS

For the non-covalent binding moiety of our designs, we chose single-domain antibodies (sdAbs), which are heavy-chain–only, functional antibody fragments that weigh 12–15 kDa, lack the constant region (*fragment crystallizable region*, or Fc) that facilitates the immune response, and feature stability, solubility and expressibility in bacteria and yeast [13, 14]. Despite the small size of sdAbs, they can be highly specific and have low-nanomolar and even picomolar affinity for their targets. We begin with the ∼1 nM binding, llama-derived αGFP nanobody (also referred to as GFP-enhancer), discovered and crystallised independently by Kirchhofer *et al.* [15] and Kubala *et al.* [12]. Serine 54 was then chosen as the site of mutation to cysteine and subsequent bioconjugation to the synthetic linker, based on the minimal impact of serine to cysteine mutations, its low conservation score in multiple sequence alignment of sdAbs, and its position in the bound structure (Figure 1b).

On the linker side, a three-part modular design was envisaged. On one end, we included a cysteine-selective, strong electrophile, to enable its *in vitro* bioconjugation to the antibody. On the other end, we placed the payload to be carried on a weakly reactive group. This payload – on proximity with a primary amine – is meant to undergo nucleophilic substitution, transferring the payload to the target. Between the electrophile and the payload, a spacer domain was inserted to determine the reach, flexibility and interaction profile of the linker, thereby affecting the site-selectivity of the reaction and its rate by varying the level of entropic restriction (Figure 1c).

Cysteine-selective, fast-reacting benzoylacrylate derivatives proved highly suitable for the first purpose [16]. Spacers ranging 4.9–11.1 Å in length, corresponding to full linker lengths of about 12–18 Å (Figure S11) and varying in hydrophobicity (with predicted partition coefficients in the 0.6–1.7 range) were explored. For the amine-reactive chemistry, the range of options can be divided into four groups: substitution at carbon, substitution at sulphur, direct addition on carbonyl carbon, and conjugate addition, all varying in chemoselectivity to primary amines, stability in aqueous environments, uninduced and proximity-induced reaction rates, and the reversibility and tracelessness of the conjugation reaction [17]. Here, we chose substitution at carbon and used phenols with varying levels of fluorination, covering a range of reaction rates while providing protection from hydrolysis [18].

Finally, as the only part of our compound to stay behind on the target, the payload is in essence the post-translational modification itself. To enable various techniques of monitoring the reaction and characterising its products, the payloads we chose were biotin, a vitamin and commonly used affinity tag (*bio*), and rhodamine B, a water-soluble and photostable fluorescent dye (*rho*).

Medium-length, trifluorophenol-based linkers loaded with rhodamine B (**3c-rho**) or biotin (**3c-bio**) were synthesised, and a gene for the αGFP nanobody was cloned into a bacterial expression vector with and without introducing an S54C mutation. αGFP and αGFP-S54C were expressed and purified, and prior to conjugation of any linkers, the fold and thermal stability of the mutant were assessed using circular dichroism (CD) (Figure 1d). These spectra and melting temperatures were consistent with correctly folded, active sdAbs [19]. The mass and purity of the cysteine mutant were also confirmed by SDS-PAGE and LC-MS (Figure 1d). Binding assays using surface plasmon resonance (SPR) confirmed binding of the cysteine mutant to GFP, with a 25-fold reduced affinity compared to the wild type, but still well within the high-picomolar/low-nanomolar range (Table S1).

To form the PEABS molecule αGFP-S54-3c-rho, αGFP-S54C was reduced, desalted and mixed with 10 equivalents of **3c-rho**. After a 50 min incubation at 25 °C, the formed PEABS molecule was purified by desalting into the reaction buffer (40 mM potassium phosphate and 40 mM NaCl, pH 7.2). Intact protein LC-MS was used to validate the formation and purity of PEABS (Figure 1e). Peaks in the chromatogram were first classified into broad categories (e.g. nanobody derivatives, surfactants, free linker species) by manual inspection. The peaks corresponding to nanobody derivatives were then marginalised over retention time to produce a spectrum, which was deconvoluted using the UniDec software [20] to afford the final mass/intensity spectrum. SDS-PAGE analysis comparing in-gel fluorescence of rhodamine B (prior to staining) and Coomassie-stained total protein further confirmed the conjugation of the antibody and the small-molecule linker (Figure 1e).

Finally, we compared the stabilities of the fully formed PEABS molecules to those of the antibody they are based on. Nano differential scanning fluorimetry (nanoDSF) was used to establish its unfolding inflection temperature. This method gave comparable results for the unconjugated αGFP-S54C and αGFP-S54-3c-rho, suggesting that the stability is unaffected by linker conjugation (Fig. 1f). Furthermore, conjugation of a PEABS linker had no measurable effect on the affinity of the nanobody to its target (Table S1).

### PEABS post-translationally modifies its target at a single position

Several methods to monitor the reaction between the PEABS and its target were explored. Many, including following the fluorescence polarisation of rhodamine B on its transfer and using in-gel fluorescence SDS-PAGE, presented difficulties in detecting any changes, mostly due to high background signals from minute residual amounts of rhodamine B in solution (released from slow hydrolysis of PEABS at the reactive group or present in the linker preparation). Serving as the gold standard for the analysis of protein post-translational modifications [21], mass spectrometry–based methods were thus sought.

Intact-protein LC-MS may not be strictly quantitative when comparing different species, as they may be ionised to different degrees and fly differently. However, when comparing very similar species, such as unmodified nanobody *vs* PEABS or GFP *vs* rhodamine-conjugated GFP (GFP-rho), this technique proved to be highly informative. To obtain the most quantitative results, ion series corresponding to the expected masses of species were identified, and extracted ion chromatograms (XICs) for each such series were isolated from the chromatogram. Subsequently, the time segment containing the peaks corresponding to all species in a set (e.g. all GFP species) was identified, and the integrated intensity over that segment was used as a surrogate for its quantity (Figure 2a). As expected, the integrals of very dissimilar species, such as nanobody and GFP, do not accurately capture their relative concentrations in solution. Therefore, each species was normalised to the sum of its own group of species (Figure 2a). This proved useful for monitoring reaction progress in a quantitative way (see below).

**Figure 2.**
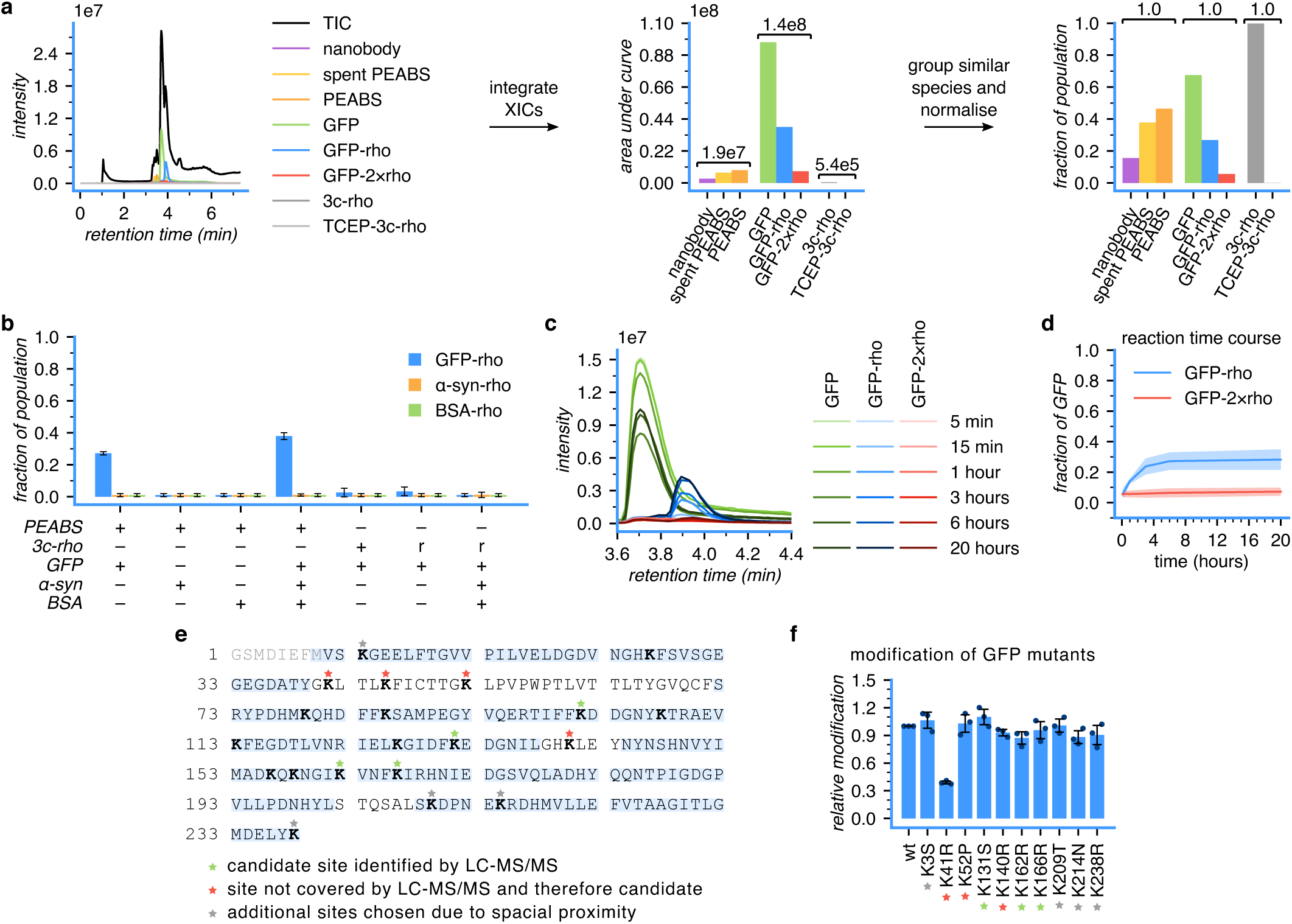
αGFP PEABS post-translationally modifies GFP in a target-selective and site-selective manner. αGFP-S54-3c-rho was used in all cases shown. **(a)** Representative example of an intact-protein LC-MS chromatogram and its extracted ion chromatograms corresponding to the experimentally relevant species in solution. Analysis of the abundance of different species in solution based on summation of signals allows semi-quantitative comparison of similar species (e.g. GFP *vs* GFP-rho), but not of dissimilar ones (e.g. GFP *vs* nanobody). **(b)** For each experimental condition in N=3 independent experiments, semi-quantitative analysis as in panel (a) was used to derive the relative populations of similar species out of the total population of that species group. In solution, PEABS transfers the rhodamine B payload to GFP, but not to α-synuclein or to BSA, and finds its target in a mixture of the three. Free linkers, whether in their benzoylacrylic form (*+*) or bound to TCEP (*r*) only modify GFP to a very small extent, and not at all when other proteins are present. Values under 1% are within the uncertainty of measurements and are therefore represented as 1 ± 1%. **(c)** A representative example of intact-protein LC-MS monitoring of the rhodamine payload transfer reaction at 18 °C. **(d)** Kinetics of GFP modification as derived from N=3 independent experiments; see panel (c), Figure S3 and Table S2. Shaded area represents 95% confidence intervals. Single-site modification is significantly preferred to multiple-site modification, suggesting selectivity for a specific site. **TIC** total ion count; **XIC** extracted ion count. **(e)** GFP modified by αGFP-S54-3c-bio was digested with chymotrypsin and analysed by LC-MS/MS. Regions covered by the analysis are highlighted blue. Four sites were identified as potential candidates for the modification site (green stars), and four more sites were in regions not covered by the analysis. Four additional sites were chosen for further inspection due to their proximity to residue S54C of the nanobody in the bound structure. **(f)** The twelve candidate residues identified in the analysis were systematically mutated out, and subjected to modification by αGFP-S54-3c-rho. Data at the 3 hour time point reveal a clear preference to modification at residue 41. GFP-K101R and GFP-K45R could not be produced. Means represent N=3 independent experiments and error bars standard deviations; full data in Figure S8.

With this tool in hand, we set out to examine the reaction between PEABS and its target. αGFP-S54-3c-rho was prepared as above, and its concentration estimated by its fraction of total nanobody species and the known starting nanobody concentration. Then, GFP, α-synuclein, BSA or all three were added at an approximate 1:1 molar ratio. After 5 hours at 25 °C, samples were analysed by LC-MS. GFP was modified to 27 ± 1% when alone in solution, and its level of modification increased to 38 ± 3% in the presence of α-synuclein or BSA, probably due to crowding effects; the latter two proteins were unmodified by PEABS, even if they were the only proteins in solution (Figure 2b).

To further validate that the antibody determines target-selectivity, free **3c-rho** linker or the small-molecule conjugate **TCEP-3c-rho** were incubated with GFP, or with the three-protein mixture. GFP alone was very minimally modified by free linkers (4.5 ± 2.5%), and in a mixture was not modified. This is in contrast to the difference observed with PEABS, indicating that GFP does not have a unique property causing its modification, and it is truly brought about by the proximity effect conferred by the antibody.

To better understand the payload transfer reaction, we monitored its kinetics. In a similar setup as before, incubated at 18 °C, samples were taken at six time-points ranging 5–1200 min after GFP addition and subjected to intact protein LC-MS (Figure 2c). A pattern of emergence of rhodamine-conjugated GFP was observed, and could be quantified by integrating extracted ion chromatograms and calculating ratios, as described above (Figure 2d). A rapidly increasing population of GFP-rho, with double-modified GFP-2×rho only appearing in very small amounts and at a much slower rate, indicated selectivity towards a specific amine of the GFP target.

We employed a two-step process to identify the preferred site of modification by PEABS. First, GFP modified using PEABS was isolated by SDS-PAGE and subjected to chymotrypsin digestion followed by LC-MS/MS analysis (Cambridge Centre for Proteomics). Such analysis is sensitive to low levels of modified peptides, and may flag sites of low-abundance and non-specific modification. On the other hand, it does not cover the entire length of the protein, and may therefore miss sites of modification even if they are abundant. Therefore, we used it for preliminary identification of candidates for further investigation. Sites that were covered by the analysis but did not show the modification were ruled out; sites that appeared modified in either GFP-rho or GFP-bio but not both were deemed low-likelihood candidates; sites that appeared in both were deemed high-likelihood candidates; and sites that were not covered had to be investigated further to be decided. We found four low-likelihood sites, identified in GFP-bio but not GFP-rho, and no high-likelihood candidates (Figure 2e). These four sites, along with four sites that were outside the covered regions, and four additional sites that were suspected for structural reasons, were chosen for further investigation (Figure 2e). In the second step, the suspected lysine residues were mutated out one by one, and modification by αGFP-S54-3c-rho was monitored. Most mutations had very minor or no effect on modification by PEABS, except for K41R, which reduced the modification rate as well as the final modification level to a fraction of their normal values, indicating that K41 is the preferred site of modification by αGFP-S54-3c-rho (Figure 2f). We note that the LC-MS peak for the modified GFP-K41-rho is shifted compared to unmodified GFP, whereas the non-specific modification does not shift the peak (Figure S8). This helped us verify that the majority of modification in wild type GFP is indeed at K41 (Figure 2c).

### PEABS is selective towards its target

After observing an apparent specificity of PEABS to its target GFP in a simple mixture of three proteins (Figure 2b), we measured its target-selectivity in a complex mixture of cellular components. For this purpose, a lysate of BL21(DE3) bacteria was prepared and its concentration measured using the Bradford assay. 1.25 µg of GFP alone, 125 µg of lysate, or a mixture of the two were combined with αGFP-S54-3c-bio at a 1:1 molar equivalence to GFP, or an equivalent volume of buffer. The six mixtures were incubated at 25 °C for 18 h. Samples of each reaction were then run on two identical SDS-PAGE gels in parallel, in the same gel box. One gel was developed using standard Coomassie stain, while the other was blotted and developed using streptavidin–Alexa Fluor™ 647, to reveal biotinylation patterns.

Remarkably, GFP was selectively modified even when it constituted only 1% of the solution by weight, to about the same extent as when it is on its own (Figure 3a). In fact, GFP alone accounted for nearly 80% of the biotinylation in solution, with only a fraction distributed among the other 99% (Figure 3b). Assuming a uniform distribution of modifiable residues, we can approximate the selectivity ratio for on-target GFP activity as 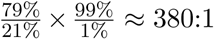. It is likely that even this low level of non-specific reactivity can be attributed to some non-specific binding of the antibody rather than non-specific reactivity of the PEABS reactive group. These findings indicate high target-selectivity, an important feature of TPTM agents.

**Figure 3.**
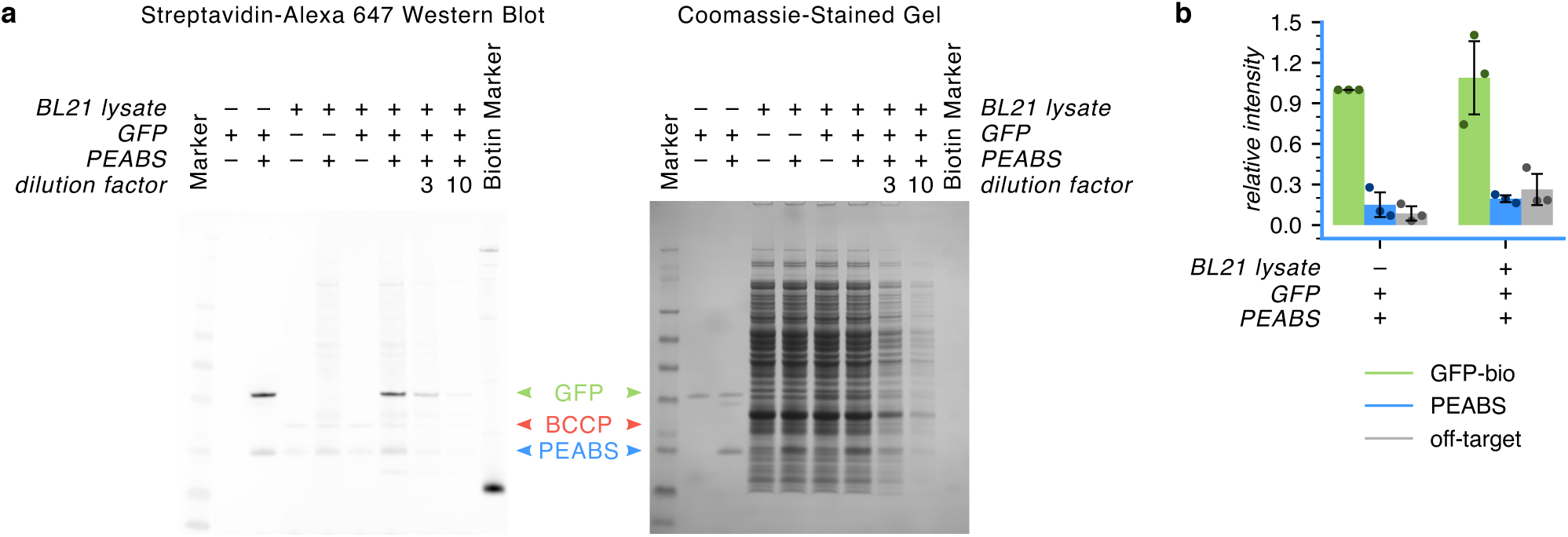
PEABS finds and modifies its target in a complex milieu of cellular components. 1.25 µg (45 pmol) of GFP was incubated with 45 pmol of αGFP-S54-3c-bio (or buffer), with or without 125 µg of soluble protein lysate from *E. coli*. **(a)** Western blot using biotinylated protein-detecting Streptavidin–Alexa 647. No signal is detected from unmodified GFP (lane 2), but after overnight incubation with PEABS it shows a strong signal, indicating its successful biotinylation (lane 3). Lysate not containing GFP presents two biotinylated bands at ∼14 kDa and ∼20 kDa (likely biotin carboxyl carrier protein [22]; lane 4) and when incubated overnight with PEABS, only a faint non-specific signal is added (lane 5). In a lysate containing GFP, only the lysate signals appear (lane 6), but with PEABS, GFP becomes highly biotinylated, whereas other proteins in solution are hardly affected (lanes 7–9). **(b)** Quantification of the intensities of the bands corresponding to biotinylated GFP and PEABS, and the total signal from all off-target bands in the western blot (lanes 3 and 7) confirms that GFP is modified to a similar extent with and without off-target proteins in solution and that, despite constituting only 1% of the protein mass in solution, it accounts for 79 ± 11% of the signal. Bars represent N=3 independent experiments and are internally normalised to the intensity of the treated GFP-only band, with error bars representing standard deviations (Figure S5).

### Characterisation of the targeted post-translational modification reaction

During experimentation with **3c-rho** and **3c-bio**, we noticed differences in the level of modification that they caused at the tested time points. To verify that these differences arose from slight differences in reactivity, we measured the TPTM kinetics on pure GFP at a 1:1 molar ratio. We found that rhodamine B is transferred at a slightly higher rate than biotin (Figure 4a). This minor but noticeable effect from the payload may be an important consideration in the design of future implementations.

**Figure 4.**
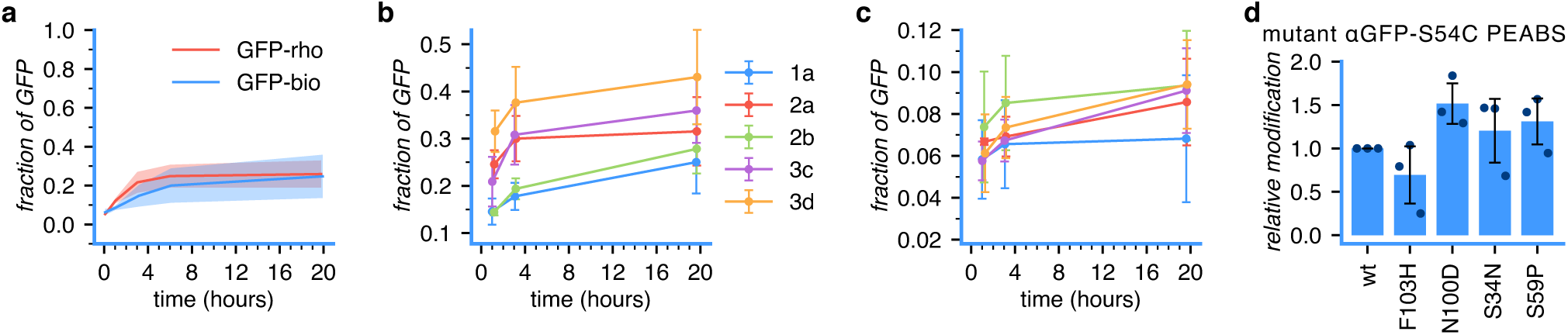
Parameters affecting the specific and non-specific activities of PEABS. **(a)** Comparison of payload transfer reaction time courses between αGFP-S54-3c-rho and αGFP-S54-3c-bio, differing only by the payload they carry. While minor differences can be detected, the overall reaction rate is similar. **(b)** Time-course comparison of αGFP-S54C PEABS based on different linkers, all carrying rhodamine B as their payload. The intrinsic reactivity of the fluorophenol and the hydrophilicity of the linker both contribute to the overall reactivity of the system. **(c)** Time-course of second rhodamine B transfer. Note the different y scale. Here, the intrinsic reactivity of the fluorophenol is the stronger determinant of site non-specific reaction. **(d)** Relative modification by mutants of GFP-S54-3c-rho at the 3-hour time point. Mutations were designed to alter the affinity of the nanobody to GFP without negatively affecting its stability. Several mutants show faster modification of GFP and reach overall higher modification levels. Shaded areas represent 95% confidence intervals, error bars represent standard deviations, all errors from N=3 independent experiments (Figures S7, S9).

Based on the principles of the PEABS design, and our results with trifluorophenol-based agents, we thought that reactivity and selectivity could also be fine-tuned by varying the spacers and electrophiles of the PEABS linkers. To investigate this possibility, we synthesised four more linkers carrying rhodamine B: a longer, more hydrophilic trifluorophenol-based linker (**3d-rho**) and three shorter, less hydrophilic and presumably less reactive monofluorophenol- and difluorophenol-based linkers (**1a-rho**, **2a-rho** and **2b-rho**). The reaction rates of the linkers are clearly affected by the hydrophilicity of the spacer, with **1a-rho** and **2b-rho** displaying identical reaction rates despite the more reactive difluorophenol in **2b-rho**, and **3d-rho** reacting faster than **3c-rho** despite the identical reactive group (Figure 4a–b). No non-specific reactivity towards α-synuclein or BSA was observed with any of the linkers. However, double modification was observed to different extents in all. We used this is an indirect measure of site non-specificity. Here, the level of non-specificity seems to follow mostly the intrinsic reactivity of the electrophile (Figure 4d). Combining these results, we expect combinations of longer, more hydrophilic spacers, with reactive groups of lower intrinsic reactivity, to be ideal for future implementations of the PEABS approach.

Next, we wanted to examine the role of affinity in the rate of the TPTM reaction. For this purpose, we designed four mutants of the αGFP-S54C nanobody that should decrease its affinity towards GFP without negatively affecting its stability. It was interesting to find that PEABS systems based on several of these mutants had significantly improved reactivity compared to the original (Figure 4d). In the mutants tested, this effect seemed to correlate with increased off-rates (Table S1). We postulate that there is an ideal window of affinity for a PEABS nanobody, where the competition between active PEABS and inactive nanobodies (e.g. unconjugated nanobodies or hydrolysed PEABS) will be impeded if their affinity is too high, while insufficient affinity would not allow the reaction to occur. This would be an important consideration in future PEABS designs.

### General applicability of the PEABS approach

Having characterised the key features of the PEABS system and their effects on its TPTM reaction, we set out to test its applicability to different targets.

Beta-2 microglobulin (β2m) is a major histocompatibility complex (MHC) I component involved in several renal diseases [23]. Most notably, it is known to accumulate and form amyloids in dialysis patients, a condition known as dialysis-related amyloidosis [24]. It may also play a role in the progression of multiple myeloma [25]. Nanobodies against this 10.9 kDa protein had been generated, and several had been shown to successfully inhibit its aggregation [26]. Based on our results with αGFP mutants, we chose to base β2m PEABS on the mid-nanomolar–affinity camel-derived nb24 [26], whose bound structure to β2m had been solved [27] (Figure 5a).

**Figure 5.**
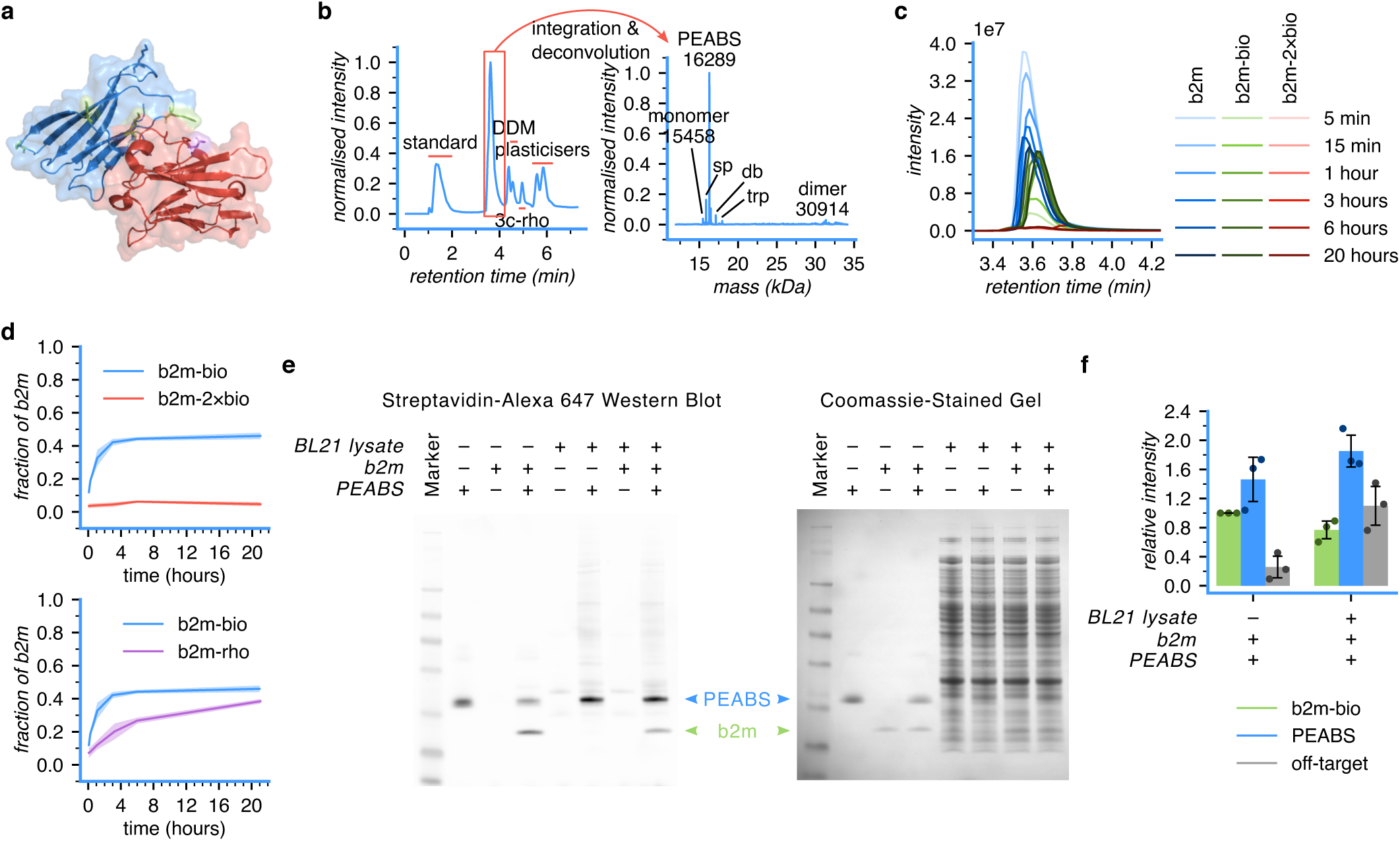
Application of PEABS to β2m. **(a)** Structure of the antibody–antigen pair consisting of β2m (blue, with residues K41, K48, K75 (front) and K58 (back) in green) and nanobody nb24 (red with residue 113, chosen for linker conjugation, in purple) [PDB: 4KDT][27]. **(b)** Confirmation of bioconjugation of **3c-rho** to nb24-D113C using LC-MS; >80% of the nanobody was converted, with low residual free linker. nb24-D113C *exp.* 14179 *obs.* 14179; PEABS *exp.* 16289 *obs.* 16289; sp (spent PEABS) *exp.* 15865 *obs.* 15865; dbl (PEABS + **3c-rho**) *exp.* 17121 *obs.* 17121; trp (PEABS + 2×**3c-rho**) *exp.* 17953 *obs.* 17953. **(c)** A representative example of intact-protein LC-MS monitoring of the biotin payload transfer reaction at 25 °C. **(d)** Kinetics of β2m modification with biotin using nb24-D113-3c-bio (blue) or rhodamine B using nb24-D113-3c-rho (purple) as derived from N=3 independent experiments each; see Figure S4 and Tables S4, S5. Here, transfer of biotin occurs more rapidly than transfer of rhodamine B. **(e)** Western blot for detection of off-target modification by nb24-D113-3c-bio, using Streptavidin–Alexa 647. **(f)** Quantification of the intensities of the bands corresponding to biotinylated β2m and PEABS, and the total signal from all off-target bands in the western blot (lanes 4 and 8) shows lower specificity of the nb24-based PEABS compared to αGFP-based PEABS; however, the on-target signal still accounts for 42 ± 4% of the non-PEABS signal, giving an estimated selectivity ratio of 70:1. Bars represent N=3 independent experiments and are internally normalised to the intensity of the treated β2m-only band (Figure S6). All means represent N=3 independent experiments, shaded areas represent 95% confidence intervals and error bars standard deviations.

Based on the bound structure, we chose residue D113 to be mutated to cysteine and used to bind PEABS linkers. Mutating this residue did not compromise stability or binding to β2m (Table S1 and Figure S10a). The same linkers used for GFP PEABS were repurposed, and formation of the PEABS molecule followed the same procedure (Figure 5b). The mass spectrometry-based technique that we developed for GFP PEABS proved effective for β2m PEABS as well (Figure 5c). Monitoring the reaction of nb24-D113-3c-bio with recombinant β2m at 25 °C revealed highly effective modification, with little to no double modification, indicating a highly site-selective TPTM reaction (Figure 5d). Isolation of β2m species by SDS-PAGE followed by chymotrypsin digestion-LC-MS/MS (Cambridge Centre for Proteomics) did not confidently identify the site of modification, but narrowed the options down to four, three of which are supported by structural analysis (Figure S10c and 5a). Intriguingly, the two payloads tested had the inverse effect on the reaction rate compared to GFP PEABS (Figure 5d, compare 4a). Finally, target-selectivity in a complex mixture was measured, giving an approximate selectivity ratio of 70:1 (Figure 5e–f). This value is lower than that of GFP PEABS, most likely due to lower specificity of the nanobody employed, but is still indicative of a highly effective TPTM agent formed against this disease-causing target.

## Discussion and Conclusions

With every step in the flow of information from DNA, to RNA, to protein, new errors and aberrant behaviours can be introduced. DNA errors can be addressed with genome editing and, increasingly, RNA errors can be addressed with RNA editing [28]. Currently, however, there is no convenient tool to systematically address *in vivo* problems at the protein level, such as accumulation, misfolding, aggregation and phase separation.

In this work, we have considered the set of requirements that a system for targeted post-translational modification (TPTM) of proteins would need to fulfil to become a viable tool for addressing such problems in the complex environment of a living cell. These requirements include having an endogenous target, binding pocket independence, tracelessness, target selectivity and site selectivity.

Towards meeting these conditions, we have reported PEABS, an antibody-based system for chemically modifying proteins, and illustrated its use by targeting two distinct proteins, GFP and β2m. Targeting endogenous proteins and being independent of ligand binding pockets is built into the design by using antibodies for their non-covalent affinity and specificity. Tracelessness is also built into the design, by employing substitution-at-carbon electrophiles – here, fluorophenols – for the reactive group. Target selectivity was shown first in pure protein solutions and simple mixtures, where LC-MS analysis could determine explicitly that GFP PEABS only modifies GFP and not α-synuclein or BSA (Figure 2c). Then, target selectivity was further monitored in a lysate, where the majority of a biotin payload was transferred to the desired target, despite it accounting for only 1% of the total protein mass (Figures 3 and 5e–f). Both systems featured insignificant double-modification, suggesting selectivity to a specific site on the target. In GFP, we showed that this site is lysine 41.

Comprising two independent components – an antibody and a synthetic linker – the PEABS design strategy allows to choose the site of modification first, and only then construct a TPTM agent to modify it by developing antibodies that bind an adjacent epitope. Additionally, the two-component design decouples the choice of binding site from the choice of chemistry – as we have shown, an existing linker can be fitted onto different antibodies, targeting different targets. Similarly, the same antibody can deploy different payloads, and even react with different adjacent residues, by attaching a different linker to its engineered cysteine. We explored the effects that the payload and linker play in specific and non-specific reactivity of PEABS in Figure 4.

We note that the target selectivity of PEABS can only be as good as that of the antibody it is based on. Of note, higher affinity was also found to not always be better, suggesting that further improvements can be made by minor modifications to the nanobody on which the system is based. Site selectivity was achieved in both GFP and β2m PEABS. However, absolute site specificity is challenging, as high payload transfer activity requires longer, more hydrophilic linkers (Figure 4b), but these also have more nucleophiles of the target within their reach. While relatively resistant to hydrolysis, PEABS made with fluorophenyl esters do hydrolyse in solution over time, especially at physiological pH at or above room temperature (Figure S2). Improving stability would be a necessary step towards using PEABS systems *in vivo* for therapeutic purposes.

In perspective, we anticipate this method to be applicable in a wide range of settings in basic research, diagnostics and therapeutics, from labelling specific proteins in the complex biological milieu to post-translationally modify disease-associated proteins to promote their degradation or protect them from aggregation. Due to the important role played by high-affinity, high-specificity antibodies against selected epitopes, novel approaches to rational antibody design will undoubtedly be crucial for future implementations of PEABS to accomplish the goal of drugging the undruggable.

## Methods

### Chemical linker synthesis

Linkers **1a**, **2a**, **2b**, **3c** and **3d** were synthesised as detailed in the Supplementary Methods. Biotin (**3c-bio**) or rhodamine B (**1a-rho**, **2a-rho**, **2b-rho**, **3c-rho** and **3d-rho**) were loaded by esterification and the products purified. A few milligrams of each loaded linker were dissolved in anhydrous dimethyl sulphoxide (DMSO) at a 20 mM, divided into 6 µL (120 nmol) aliquots, lyophilised and stored at 4–10 °C in the dark. Just before use, one aliquot was taken out, dissolved back into 6 µL DMSO, and diluted into 6 µL of 4% dodecyl maltoside (DDM) in water, for a final 10 mM in 50 : 48 : 2 DMSO : H_2_O : DDM.

### PEABS preparation

A 25 nmol (e.g. 400 µL at 63 µM) stock of purified cysteine-mutant nanobody (αGFP-S54C or nb24-D113C) in 40 mM 2-ethanesulphonic acid (MES) + 40 mM NaCl, pH 6.1 was thawed from −80 °C to room temperature quickly, incubated on ice for 45 min, centrifuged to remove aggregates (20 min / 4 °C / 20000 × g) and monomerised by reduction using 8 molar equivalents tris(2-carboxyethyl)phosphine (TCEP) (2 hours at 25 °C, no shaking). A stock of loaded linker (e.g. **3c-rho**) was prepared at 10 mM in 50 : 48 : 2 DMSO : H_2_O : DDM. The monomerised antibody was then desalted using PD MiniTrap desalting columns with Sephadex G-25 resin (Cytiva) to remove excess TCEP (which competes with the cysteine for the reaction with the benzoylacrylic moiety) and its concentration was determined using a NanoDrop spectrophotometer. The nanobody was diluted to 20 µM, mixed with a 1 : 50-diluted linker (final 200 µM, 10 eq) and supplemented with 60 µM TCEP, then incubated for 50 min at 25 °C without shaking. These conditions were used to achieve maximal formation of PEABS while minimising over-modified nanobody species (Figure S1, Supplementary Note 2). Finally, the formed PEABS molecule was purified from the unreacted excess linker, surfactant and DMSO by desalting into TPTM reaction buffer (40 mM potassium phosphate + 40 mM NaCl, pH 7.2) and centrifuging to remove suspended material (15 min / 4 °C / 20000 × g).

### GFP selectivity assay

PEABS αGFP-S54-3c-rho was prepared as above. Small-molecule linker controls were prepared by diluting a 10 mM stock of **3c-rho** into the TPTM reaction buffer in the absence (*+* condition) or presence of 2 molar equivalents of TCEP (*r* condition). Protein stocks of GFP, α-synuclein and BSA in the TPTM reaction buffer were prepared at 30 µM, and used to prepare 1.8 µM solutions of each or an equimolar mixture of all three (1.8 µM each). 90 pmol (e.g. 9 µL at 10 µM) of either PEABS, free **3c-rho** or TCEP-3c-rho were added to a 50 µL sample of each protein or mixture thereof, representing a 1:1 molar ratio between PEABS and the target. The reaction was allowed to proceed at 25 °C for 5 hours (or at 18 °C for 3 h and then at 25 °C for 3 h) and analysed by LC-MS.

### Time-course experiments

PEABS molecules αGFP-S54-3c-rho and αGFP-S54-3c-bio, or nb24-D113-3c-rho and nb24-D113-3c-bio, were prepared as described. A solution of 1.8 µM target (GFP or β2m) in the TPTM reaction buffer was prepared, and to a 100 µL sample of it, 180 pmol (e.g. 18 µL at 10 µM) PEABS was added for a final 1 : 1 (PEABS : target) molar ratio. The reaction was allowed to proceed at 18 °C or 25 °C, while samples were analysed at 5, 15, 60, 180, 360 and 1200 minutes by LC-MS. Reaction starting times were staggered by 30 minute increments to allow parallel analysis. For rhodamine-loaded PEABS comparisons, PEABS molecules were prepared as above, except the final desalting step which was performed using 7K MWCO, 0.7 mL polyacrylamide spin desalting columns (Pierce) in small scale (100 µL + 20 µL stacker). Reactions were started at 10 minute intervals to allow parallel analysis, allowed to proceed at 25 °C, and measured by LC-MS at the 1, 3 and 20 h time points.

### Selectivity in a complex environment

PEABS αGFP-S54-3c-bio or nb24-D113-3c-bio was prepared as described. BL21(DE3) clarified lysate was prepared as described in the Supplementary Methods. 50 µg/mL target (1.8 µM GFP or 4.6 µM β2m, lysate (5 mg/mL) or a mixture of both (1:100 target:lysate) were supplemented with an equimolar amount of PEABS, or the same volume of buffer (*untreated controls*). Reactions were allowed to proceed at 25 °C overnight. Samples of each condition were loaded onto two identical SDS-PAGE gels, which were run in parallel; one was Coomassie-stained and the other analysed by western blot to detect biotinylated proteins as described in the Supplementary Methods.

### Other materials and methods

For further discussion of the considerations that went into the above procedures, please refer to Supplementary Note 2. Gene and protein sequences and methods for site-directed mutagenesis, protein expression and purification, organic synthesis of compounds, mass spectrometry, Western blot, and protein and small molecule quantification and characterisation are available in the Supplementary Methods.

## Data Availability

All substantial data to support the results in this paper are contained within it and its supplementary materials. The mass spectrometry proteomics data supporting Figures 2e and S10c have been deposited to the ProteomeXchange Consortium via the PRIDE partner repository with the dataset identifier PXD049693. Raw intact protein mass spectrometry data are available on request from the corresponding authors.

## Code Availability

Code for all analyses from raw files is available on GitHub at https://github.com/vendruscolo-lab/PEABS-LC-MS. It is dependent on the MassLynx API, which is freely available from Waters at https://support.waters.com/KB_Inst/Mass_Spectrometry/WKB1775_What_is_the_MassLynx_Raw_Data_Reader_Interface_Library.

## Acknowledgements

The authors thank Dr Cristina Visentin and Prof. Stefano Ricagno (University of Milan) for the β2m protein, Dr Rebecca C. Gregory for producing α-synuclein and proof-reading the manuscript, and Dr Thomas Löhr, Dr Phil Lindstedt, Ross Taylor, Dr Roxine Staats and Magdalena Nowinska for helpful discussions. O.R. was supported by a scholarship from the Reuben Foundation. P.S. is a Royal Society University Research Fellow (URF\R1\201461). This work was supported by the UKRI (10059436, 10061100) and the Centre for Misfolding Diseases.

## Author Contributions

The project was conceived by O.R. and M.V.. Chemical linkers and synthetic pathways were designed by J.K. and O.R., and performed by O.R. and I.G.. Except where mentioned otherwise, proteins were produced by O.R., M.A. (nb24 and its variants) or V.R. (GFP-wt). Experiments were designed by O.R. and performed by O.R., M.A. (β2m PEABS and BLI) and V.R. (SPR). The synthetic side of the work was supervised by G.J.L.B.. β2m work and nanobody design were supervised by P.S.. Funding was secured by M.V., who supervised all other aspects of the project. The manuscript was written by O.R. and M.V. and approved by all authors.

## Notes

### Competing Interest Statement

The authors have declared no competing interest.

### Summary of Updates

New results, including implementation of the PEABS approach for the biologically-relevant target beta-2 microglobulin. Additionally, methods were clarified and expanded upon, and the main text was restructured to improve readability.

