## Supplementary Information for "Targeted protein modification with an antibody-based system"

Centre for Misfolding Diseases, Yusuf Hamied Department of Chemistry,
University of Cambridge, Cambridge, CB2 1EW, United Kingdom

[†] Currently at: The Francis Crick Institute, London, UK

[‡] Currently at: St. Anna Children's Cancer Research Institute (CCRI) and CeMM Research Center for Molecular Medicine of the Austrian Academy of Sciences, Vienna, Austria

✉ Corresponding authors:

| | |
|---|---|
| Prof. Michele Vendruscolo | Dr Oded Rimon |
| | |
| Tel. +44 (0)1223 763873 | |

**Instrument Reports** 56

**References** 151

### Extended Data

| Ligand | Analyte | $k_a$ ($M^{-1}s^{-1}$) | $k_d$ ($s^{-1}$) | $k_D$ (M) |
|---|---|---|---|---|
| *αGFP* | GFP | $1.093 \times 10^6$ | $4.756 \times 10^{-5}$ | $4.350 \times 10^{-11}$ |
| *αGFP-S54C* | GFP | $1.690 \times 10^5$ | $1.883 \times 10^{-4}$ | $1.115 \times 10^{-9}$ |
| *αGFP-S54-2b* | GFP | $2.222 \times 10^5$ | $1.087 \times 10^{-4}$ | $4.894 \times 10^{-10}$ |
| *αGFP-S54C-F103H* | GFP | $7.248 \times 10^4$ | $4.868 \times 10^{-4}$ | $6.716 \times 10^{-9}$ |
| *αGFP-S54C-S34N* | GFP | $3.827 \times 10^5$ | $6.635 \times 10^{-3}$ | $1.733 \times 10^{-8}$ |
| αGFP-S54C | *GFP-K41R* | $1.503 \times 10^5$ | $1.740 \times 10^{-4}$ | $1.158 \times 10^{-9}$ |
| αGFP-S54C | *GFP-K131S* | $1.445 \times 10^5$ | $1.680 \times 10^{-4}$ | $1.162 \times 10^{-9}$ |
| αGFP-S54C | *GFP-K140R* | $1.619 \times 10^5$ | $1.693 \times 10^{-4}$ | $1.046 \times 10^{-9}$ |
| αGFP-S54C | *GFP-K162R* | $1.354 \times 10^5$ | $1.709 \times 10^{-4}$ | $1.263 \times 10^{-9}$ |
| αGFP-S54C | *GFP-K166R* | $1.579 \times 10^5$ | $1.753 \times 10^{-4}$ | $1.110 \times 10^{-9}$ |
| β2m | *nb24* | $1.708 \times 10^5$ | $1.795 \times 10^{-2}$ | $1.051 \times 10^{-7}$ |
| β2m | *nb24-D113C* | $1.819 \times 10^5$ | $1.795 \times 10^{-2}$ | $9.869 \times 10^{-8}$ |

**Table S1 | Kinetic parameters of binding of αGFP and its variants to GFP.** Surface plasmon resonance (SPR) analysis, fitted by a 1:1 kinetic binding model reveals that the serine to a cysteine mutation at position 54 reduces the affinity by a factor of about 9. Conjugation of a PEABS linker at the cysteine does not have a measurable additional effect on the affinity. Full SPR reports are found in pages 56 (αGFP), 62 (αGFP-S54C), 67 (αGFP-S54-2b), 72 (αGFP-S54C-F103H), 77 (αGFP-S54C-S34N), 82 (GFP-K41R), 87 (GFP-K131S), 92 (GFP-K140R), 97 (GFP-K162R), 102 (GFP-K166R). BLI data appear in Figure S10b.

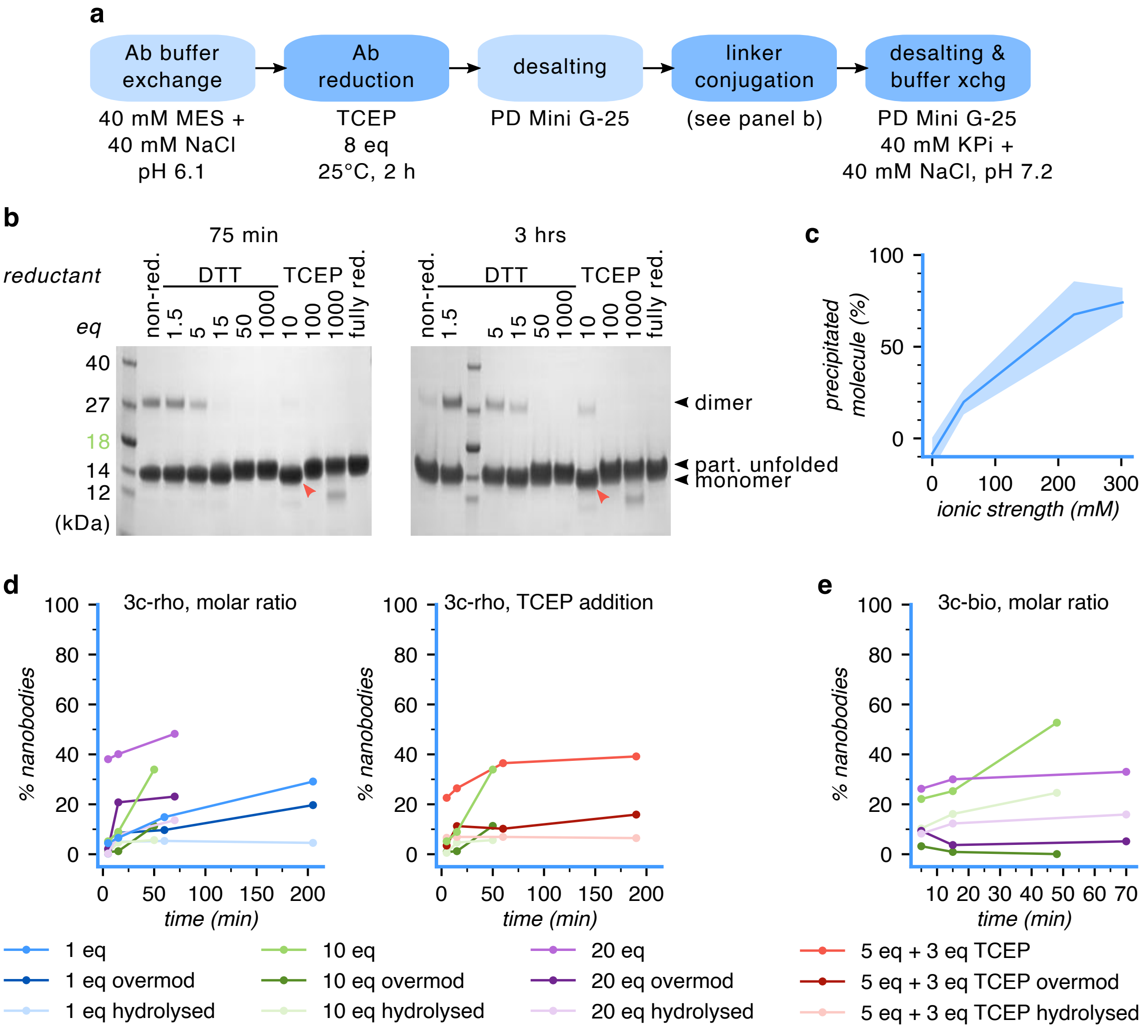

**Figure S1 | Parameters in PEABS experimental design.** **(a)** General procedure for the bioconjugation of an antibody and a synthetic linker to form PEABS molecules. **(b)** Optimisation of reduction conditions of DesAb-HET-A112C for the elimination of dimers while retaining the intramolecular disulphide bond. Non-reduced control and under-reduced samples show presence of dimers, whereas fully-reduced control and over-reduced samples unfold due to loss of disulphide, evident by the higher apparent protein size. 10 eq TCEP removes most of the dimers while retaining the folded form of the protein (red arrowhead). **(c)** Buffer ionic strength effect on compound 2b-rho precipitation in buffered aqueous solutions. **(d)** Time course measurements of the conjugation of αGFP-S54C to **3c-rho** at 1 : 1, 1 : 10 and 1 : 20 molar ratios, or a 1 : 3 : 5 ratio of nanobody : TCEP : linker. Reintroduction of sub-stoichiometric levels of TCEP may prove useful for suppressing over-modification (right-hand panel). **(e)** Time course measurements of the conjugation of αGFP-S54C to **3c-bio** at 1 : 10 and 1 : 20 molar ratios shows similar trends.

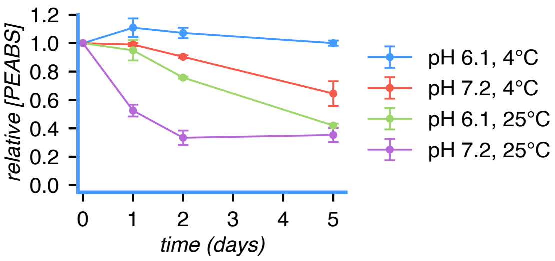

**Figure S2 | Stability of αGFP-S54-3c-rho.** PEABS constructs were prepared according to the general procedure and either desalted into the linker-conjugation buffer (pH 6.1) or buffer exchanged into the TPTM reaction buffer (pH 7.2). Then, each was divided into two portions, with one kept at 25 °C and the other in the refrigerator (4–6 °C). At several time points over the course of five days, the fraction of PEABS in solution, relative to total nanobody species, was measured by LC-MS and compared to the starting fraction after desalting. Error bars represent standard deviations over N=2 trials.

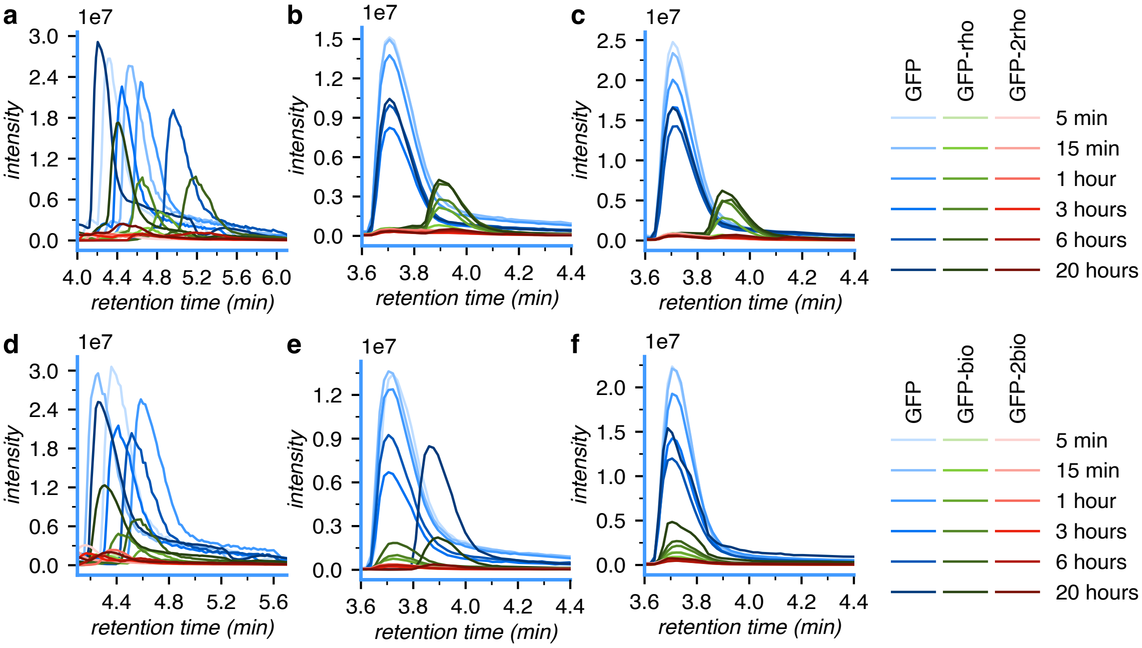

**Figure S3 | GFP PEABS 3c-rho and 3c-bio TPTM reaction kinetics.** **(a)** Extracted ion chromatograms of GFP species in a time-course experiment of TPTM reaction by αGFP-S54-3c-rho. The area under the curve in the shown region is used to arrive at the numbers in Tables S2, S3. **(b)** As in (a), from second independent experiment. **(c)** As in (a), from third independent experiment. **(d)** Extracted ion chromatograms of GFP species in a time-course experiment of TPTM reaction by αGFP-S54-3c-bio. **(e)** As in (d), from second independent experiment. **(f)** As in (d), from third independent experiment.

| *Repeat* | Time / min | Area under XIC curve | | | Fraction | |
|---|---|---|---|---|---|---|
| | | GFP | GFP-rho | GFP-2×rho | GFP-rho | GFP-2×rho |
| *1* | 5 | 7409077 | 436579 | 578135 | 0.051827 | 0.068631 |
| | 15 | 7777425 | 571442 | 581340 | 0.063990 | 0.065098 |
| | 61 | 6791886 | 1086520 | 572438 | 0.128569 | 0.067737 |
| | 177 | 5890933 | 2132684 | 673085 | 0.245229 | 0.077396 |
| | 361 | 6011579 | 2720353 | 1005434 | 0.279373 | 0.103255 |
| | 1174 | 7907271 | 4003605 | 1547130 | 0.297489 | 0.114960 |
| *2* | 5 | 2804594 | 163784 | 239754 | 0.051053 | 0.074733 |
| | 15 | 2786424 | 215579 | 246050 | 0.066372 | 0.075753 |
| | 61 | 2486069 | 367317 | 233117 | 0.119008 | 0.075528 |
| | 188 | 1406268 | 417924 | 148423 | 0.211863 | 0.075242 |
| | 359 | 1689374 | 610572 | 194301 | 0.244792 | 0.077900 |
| | 1188 | 1697486 | 639569 | 224706 | 0.249660 | 0.087716 |
| *3* | 5 | 3696606 | 243289 | 298701 | 0.057399 | 0.070472 |
| | 14 | 3585675 | 258747 | 285838 | 0.062647 | 0.069206 |
| | 60 | 3087006 | 444708 | 257112 | 0.117374 | 0.067861 |
| | 178 | 2603006 | 679532 | 239255 | 0.192951 | 0.067936 |
| | 362 | 2273538 | 717894 | 241005 | 0.222091 | 0.074558 |
| | 1195 | 2700101 | 913475 | 347605 | 0.230607 | 0.087753 |
| *Mean* | 5 | – | – | – | 0.053 ± 0.003 | 0.071 ± 0.003 |
| | 15 | – | – | – | 0.064 ± 0.002 | 0.070 ± 0.004 |
| | 60 | – | – | – | 0.122 ± 0.005 | 0.070 ± 0.004 |
| | 181 | – | – | – | 0.217 ± 0.022 | 0.074 ± 0.004 |
| | 360 | – | – | – | 0.249 ± 0.024 | 0.085 ± 0.013 |
| | 1186 | – | – | – | 0.259 ± 0.028 | 0.097 ± 0.013 |

**Table S2 | GFP PEABS 3c-rho TPTM kinetics data**

| *Repeat* | Time / min | Area under XIC curve | | | Fraction | |
|---|---|---|---|---|---|---|
| | | GFP | GFP-bio | GFP-2×bio | GFP-bio | GFP-2×bio |
| *1* | 5 | 8193952 | 492116 | 276705 | 0.054907 | 0.030873 |
| | 14 | 8084148 | 545100 | 278458 | 0.061194 | 0.031260 |
| | 59 | 7525904 | 761278 | 285724 | 0.088800 | 0.033329 |
| | 189 | 5911625 | 1358086 | 306361 | 0.179260 | 0.040438 |
| | 372 | 5800194 | 2061179 | 397301 | 0.249577 | 0.048107 |
| | 1194 | 7342661 | 3626413 | 774775 | 0.308793 | 0.065973 |
| *2* | 5 | 2612749 | 174809 | 109369 | 0.060343 | 0.037754 |
| | 15 | 2626029 | 190911 | 112801 | 0.065163 | 0.038502 |
| | 60 | 2377203 | 212471 | 103902 | 0.078881 | 0.038574 |
| | 200 | 1230673 | 192526 | 59693 | 0.129832 | 0.040255 |
| | 363 | 1653664 | 351145 | 85875 | 0.167957 | 0.041075 |
| | 1193 | 1425613 | 386208 | 86338 | 0.203465 | 0.045485 |
| *3* | 5 | 3182689 | 262106 | 153983 | 0.072832 | 0.042788 |
| | 14 | 3146701 | 263822 | 149630 | 0.074104 | 0.042029 |
| | 59 | 2900510 | 289386 | 135704 | 0.087018 | 0.040806 |
| | 181 | 2164031 | 351398 | 106481 | 0.134024 | 0.040612 |
| | 361 | 1879393 | 443739 | 105922 | 0.182680 | 0.043606 |
| | 1200 | 2641549 | 844339 | 185604 | 0.229972 | 0.050553 |
| *Mean* | 5 | – | – | – | 0.063 ± 0.008 | 0.037 ± 0.005 |
| | 15 | – | – | – | 0.067 ± 0.005 | 0.037 ± 0.004 |
| | 59 | – | – | – | 0.085 ± 0.004 | 0.038 ± 0.003 |
| | 190 | – | – | – | 0.148 ± 0.022 | 0.040 ± 0.000 |
| | 365 | – | – | – | 0.200 ± 0.036 | 0.044 ± 0.003 |
| | 1195 | – | – | – | 0.247 ± 0.045 | 0.054 ± 0.009 |

**Table S3 | GFP PEABS 3c-bio TPTM kinetics data**

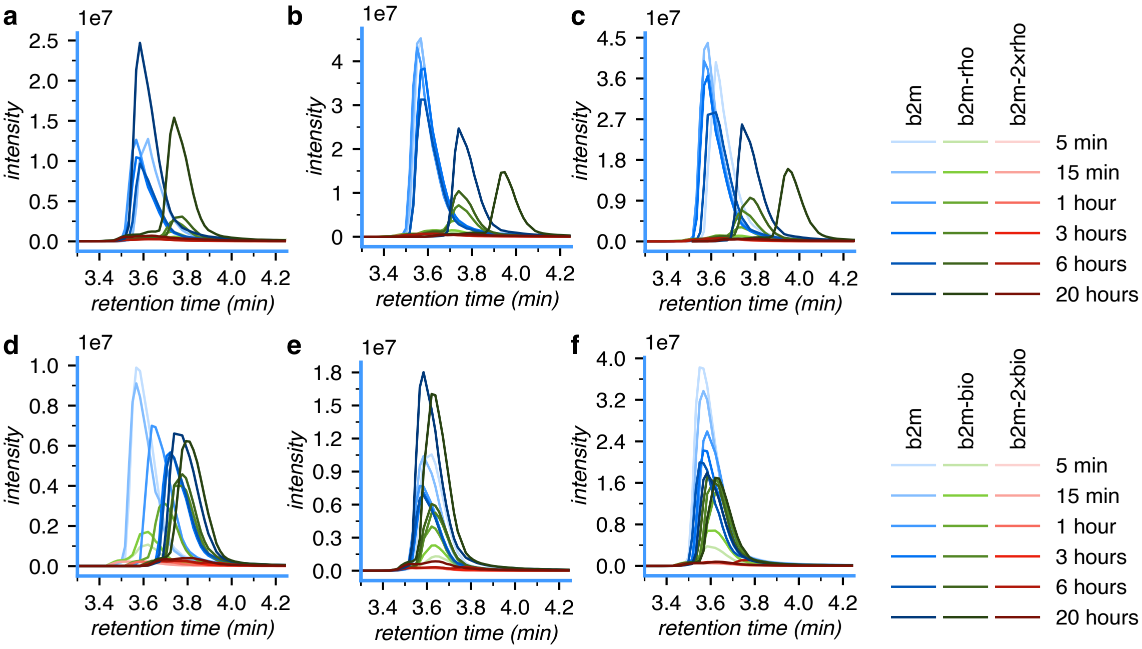

**Figure S4 | β2m PEABS 3c-rho and 3c-bio TPTM reaction kinetics. (a)** Extracted ion chromatograms of β2m species in a time-course experiment of TPTM reaction by nb24-D113-3c-rho. The area under the curve in the shown region is used to arrive at the numbers in Tables S4, S5. **(b)** As in (a), from second independent experiment. **(c)** As in (a), from third independent experiment. **(d)** Extracted ion chromatograms of GFP species in a time-course experiment of TPTM reaction by nb24-D113-3c-bio. The area under the curve in the shown region is used to arrive at the numbers in Table SS5. **(e)** As in (d), from second independent experiment. **(f)** As in (d), from third independent experiment.

| *Repeat* | Time / min | Area under XIC curve | | | Fraction | |
|---|---|---|---|---|---|---|
| | | β2m | β2m-rho | β2m-2×rho | β2m-rho | β2m-2×rho |
| *1* | 5 | 1445045 | 150804 | 91827 | 0.089356 | 0.054410 |
| | 15 | 1618163 | 175134 | 96002 | 0.092698 | 0.050814 |
| | 65 | 1494737 | 241245 | 89441 | 0.132159 | 0.048998 |
| | 223 | 1286270 | 410735 | 97070 | 0.228940 | 0.054106 |
| | 364 | 1159384 | 477503 | 95804 | 0.275585 | 0.055292 |
| | 1396 | 3041238 | 2071383 | 235439 | 0.387315 | 0.044023 |
| *2* | 5 | 5071651 | 391030 | 230152 | 0.068688 | 0.040428 |
| | 13 | 5322618 | 441631 | 225708 | 0.073729 | 0.037681 |
| | 60 | 5030704 | 696689 | 225463 | 0.117035 | 0.037875 |
| | 177 | 4547620 | 1105995 | 225969 | 0.188108 | 0.038433 |
| | 356 | 3810849 | 1477146 | 215106 | 0.268421 | 0.039088 |
| | 1196 | 3030144 | 1939555 | 183582 | 0.376373 | 0.035624 |
| *3* | 5 | 4538128 | 303686 | 197653 | 0.060262 | 0.039221 |
| | 15 | 5021024 | 378708 | 204066 | 0.067581 | 0.036416 |
| | 63 | 4810821 | 592880 | 198878 | 0.105823 | 0.035498 |
| | 180 | 4301518 | 1049453 | 210908 | 0.188687 | 0.037920 |
| | 361 | 3914993 | 1436387 | 206121 | 0.258459 | 0.037089 |
| | 1202 | 3175390 | 2153607 | 202716 | 0.389320 | 0.036646 |
| *Mean* | 5 | – | – | – | 0.073 ± 0.012 | 0.045 ± 0.007 |
| | 15 | – | – | – | 0.078 ± 0.011 | 0.042 ± 0.007 |
| | 62 | – | – | – | 0.118 ± 0.011 | 0.041 ± 0.006 |
| | 193 | – | – | – | 0.202 ± 0.019 | 0.043 ± 0.008 |
| | 360 | – | – | – | 0.267 ± 0.007 | 0.044 ± 0.008 |
| | 1264 | – | – | – | 0.384 ± 0.006 | 0.039 ± 0.004 |

**Table S4 | β2m PEABS 3c-rho TPTM kinetics data**

| *Repeat* | Time / min | Area under XIC curve | | | Fraction | |
|---|---|---|---|---|---|---|
| | | β2m | β2m-rho | β2m-2×rho | β2m-rho | β2m-2×rho |
| *1* | 5 | 1269601 | 189706 | 62611 | 0.124650 | 0.041140 |
| | 15 | 1159051 | 270585 | 62237 | 0.181373 | 0.041717 |
| | 60 | 953156 | 439904 | 62487 | 0.302226 | 0.042931 |
| | 182 | 777006 | 594840 | 70453 | 0.412425 | 0.048848 |
| | 356 | 768573 | 664823 | 89302 | 0.436609 | 0.058648 |
| | 1202 | 1030000 | 936602 | 110816 | 0.450849 | 0.053343 |
| *2* | 5 | 1448105 | 217787 | 68154 | 0.125595 | 0.039304 |
| | 15 | 1360543 | 337859 | 70472 | 0.191002 | 0.039840 |
| | 60 | 989107 | 521824 | 69226 | 0.330235 | 0.043810 |
| | 180 | 872514 | 691828 | 78630 | 0.421083 | 0.047859 |
| | 355 | 912546 | 815093 | 117660 | 0.441713 | 0.063762 |
| | 1376 | 2360401 | 2184922 | 221366 | 0.458373 | 0.046440 |
| *3* | 5 | 5011896 | 633873 | 160465 | 0.109171 | 0.027637 |
| | 16 | 4339945 | 1052935 | 166461 | 0.189399 | 0.029943 |
| | 81 | 3468777 | 1963449 | 183289 | 0.349647 | 0.032640 |
| | 177 | 2847450 | 2309091 | 196406 | 0.431368 | 0.036691 |
| | 361 | 2647693 | 2426693 | 347660 | 0.447560 | 0.064120 |
| | 1201 | 2351248 | 2261082 | 197440 | 0.470102 | 0.041050 |
| *Mean* | 5 | – | – | – | 0.120 ± 0.008 | 0.036 ± 0.006 |
| | 15 | – | – | – | 0.187 ± 0.004 | 0.037 ± 0.005 |
| | 67 | – | – | – | 0.327 ± 0.019 | 0.040 ± 0.005 |
| | 179 | – | – | – | 0.422 ± 0.008 | 0.044 ± 0.006 |
| | 357 | – | – | – | 0.442 ± 0.004 | 0.062 ± 0.002 |
| | 1260 | – | – | – | 0.460 ± 0.008 | 0.047 ± 0.005 |

**Table S5 | β2m PEABS 3c-rho TPTM kinetics data**

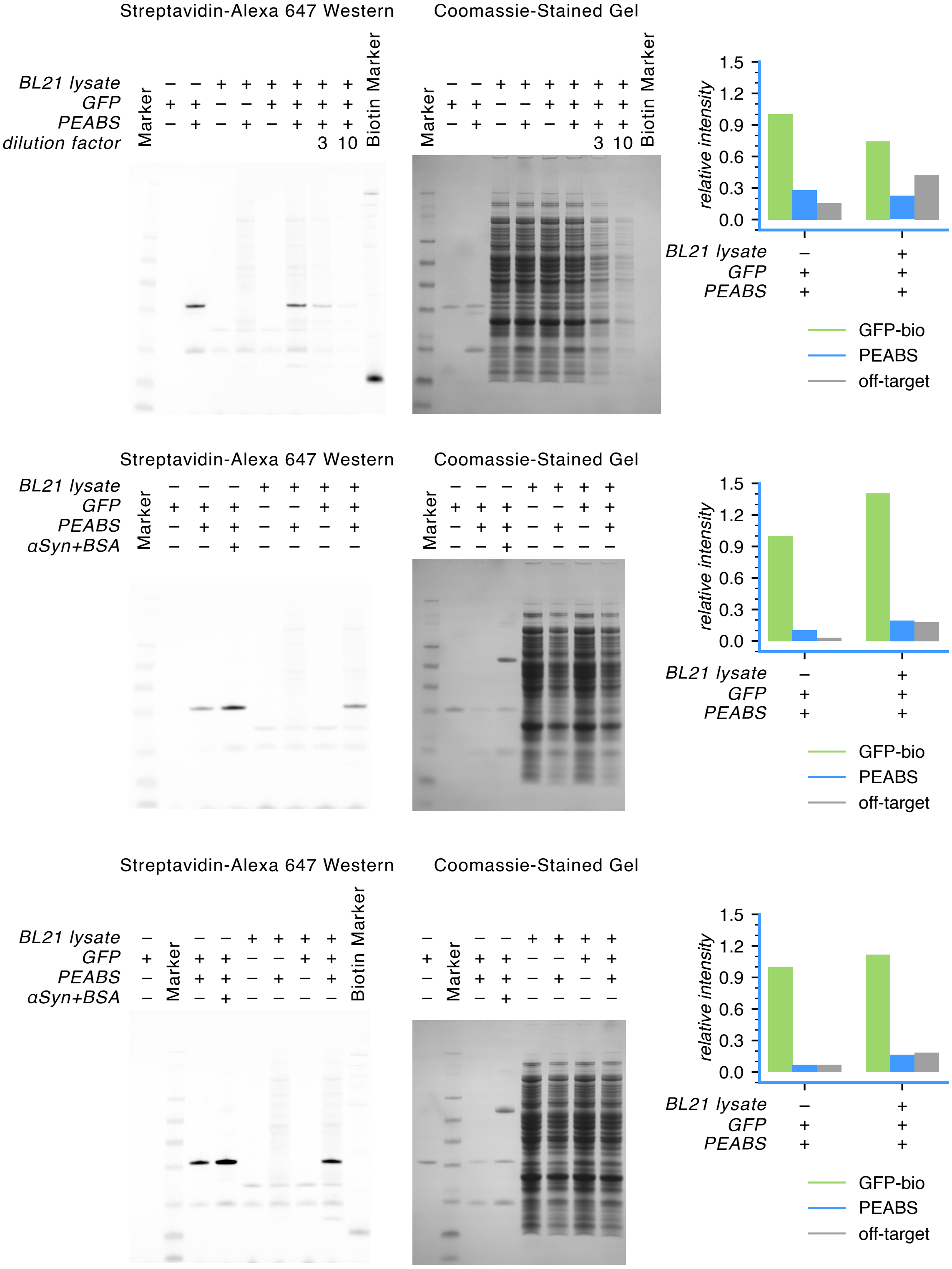

**Figure S5 | GFP PEABS modifies its target in a complex milieu of cellular components.** Three repeats of the biotin-detecting western blot experiment, with their respective quantification plots. Full report on pages 107, 115, 123.

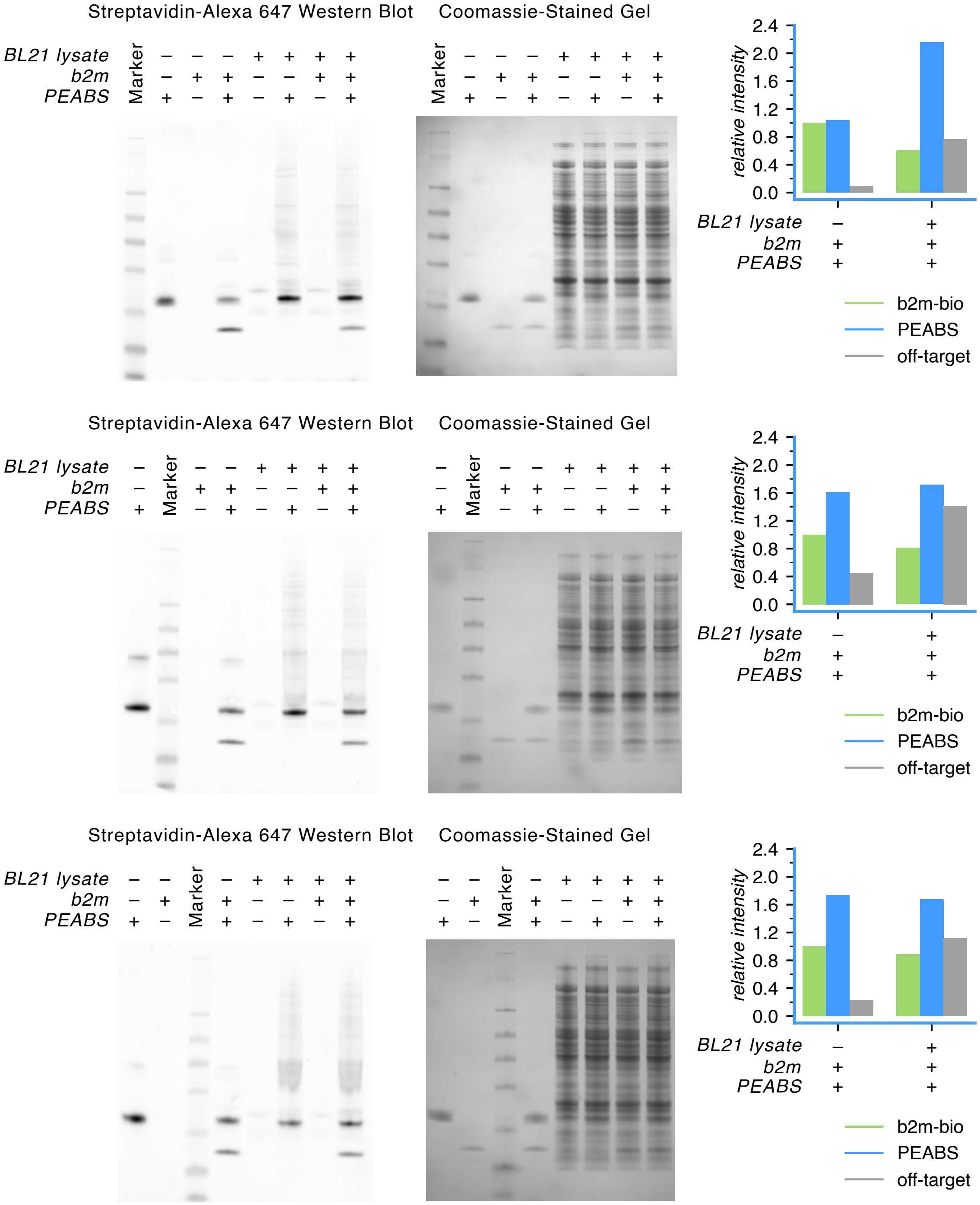

**Figure S6 | β2m PEABS modifies its target in a complex milieu of cellular components.** Three repeats of the biotin-detecting western blot experiment, with their respective quantification plots. Full report on pages 130, 137, 144.

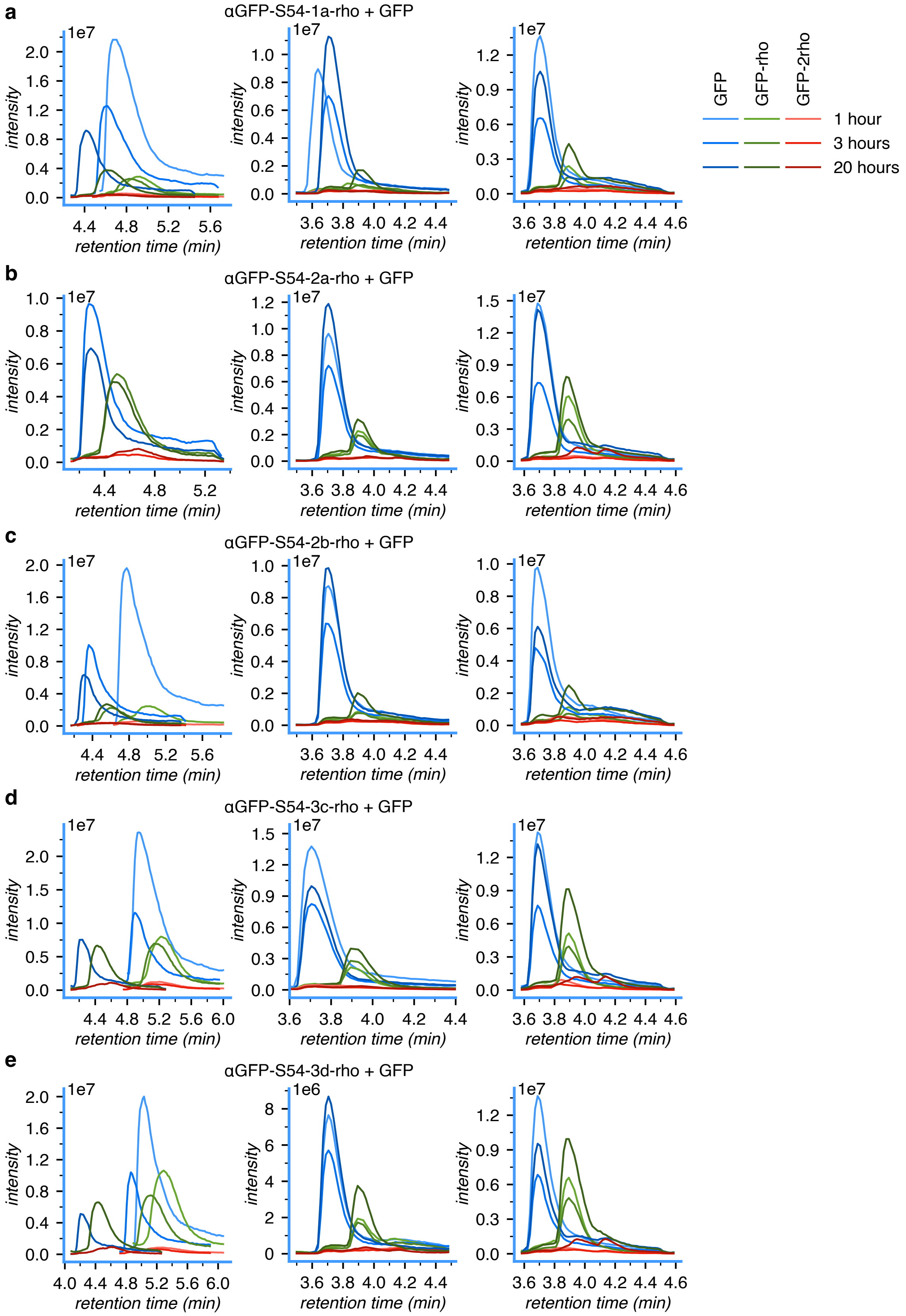

**Figure S7 | GFP PEABS 1a-rho, 2a-rho, 2b-rho, 3c-rho and 3d-rho TPTM reaction kinetics.** Extracted ion chromatograms of GFP species in three independent time-course experiments of each TPTM reaction by αGFP-S54-1a-rho (a), 2a-rho (b), 2b-rho (c), 3c-rho (d) and 3d-rho (e).

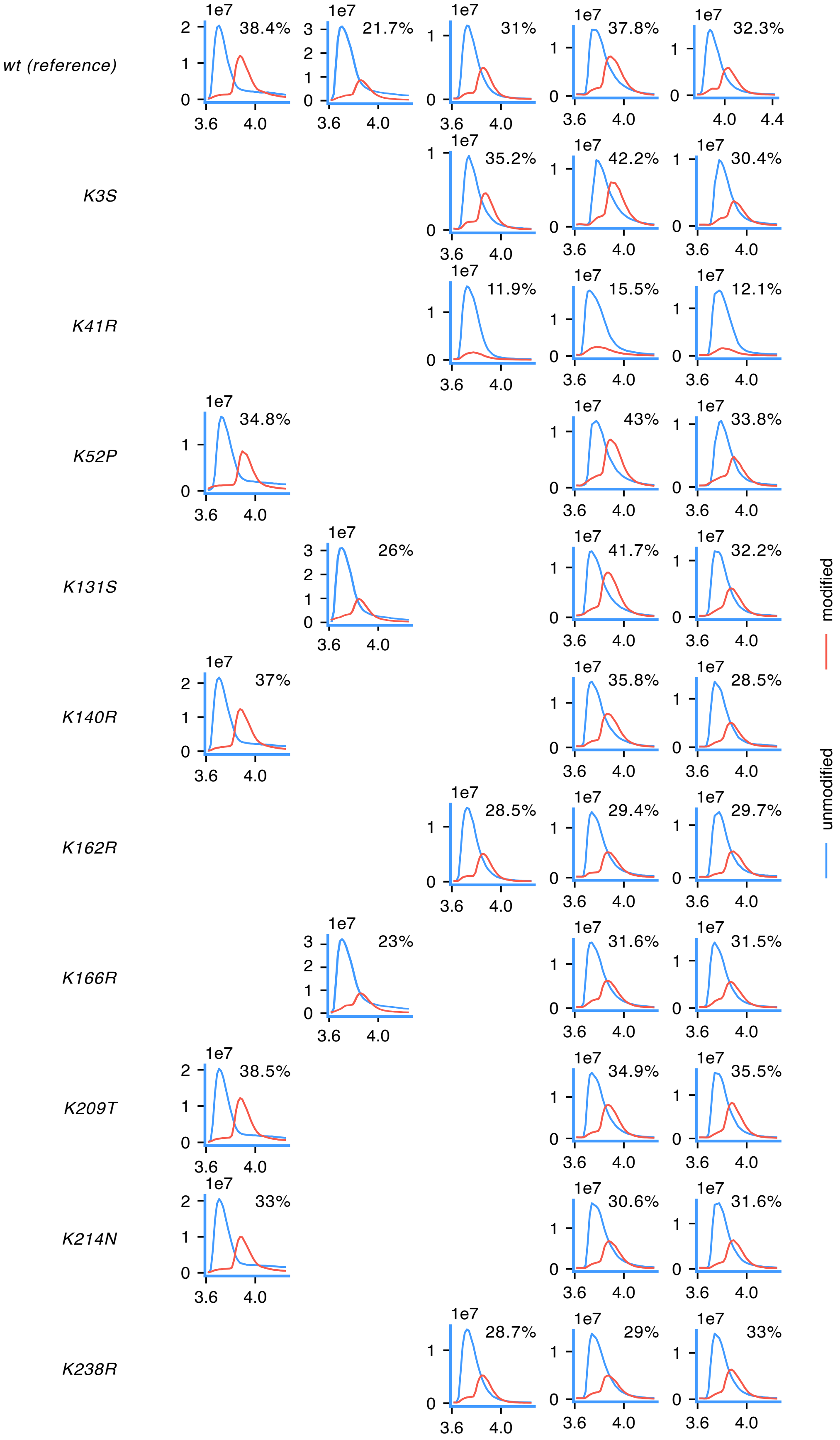

**Figure S8 | Systematic removal of lysine residues from GFP.** 3 hour data.

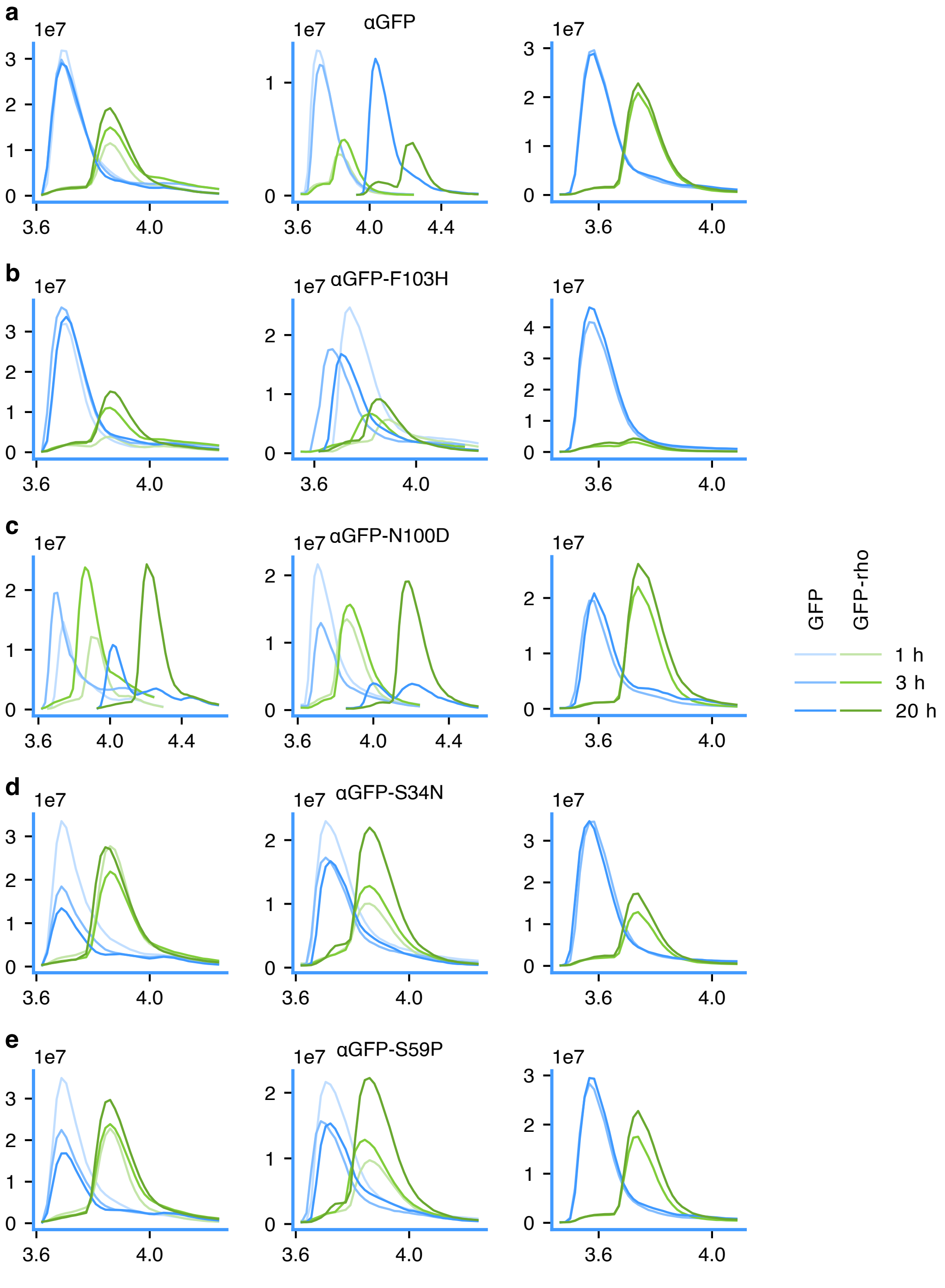

**Figure S9 | αGFP nanobodies with altered affinity and their TPTM reaction kinetics.** Mutations were designed to affect the affinity of the nanobody to GFP without reducing its stability, and PEABS variants were formed from all at the same time and tested against the same batch of GFP. Modification was measured at 1 hour, 3 hours and 20 hours into incubation at 20 °C.

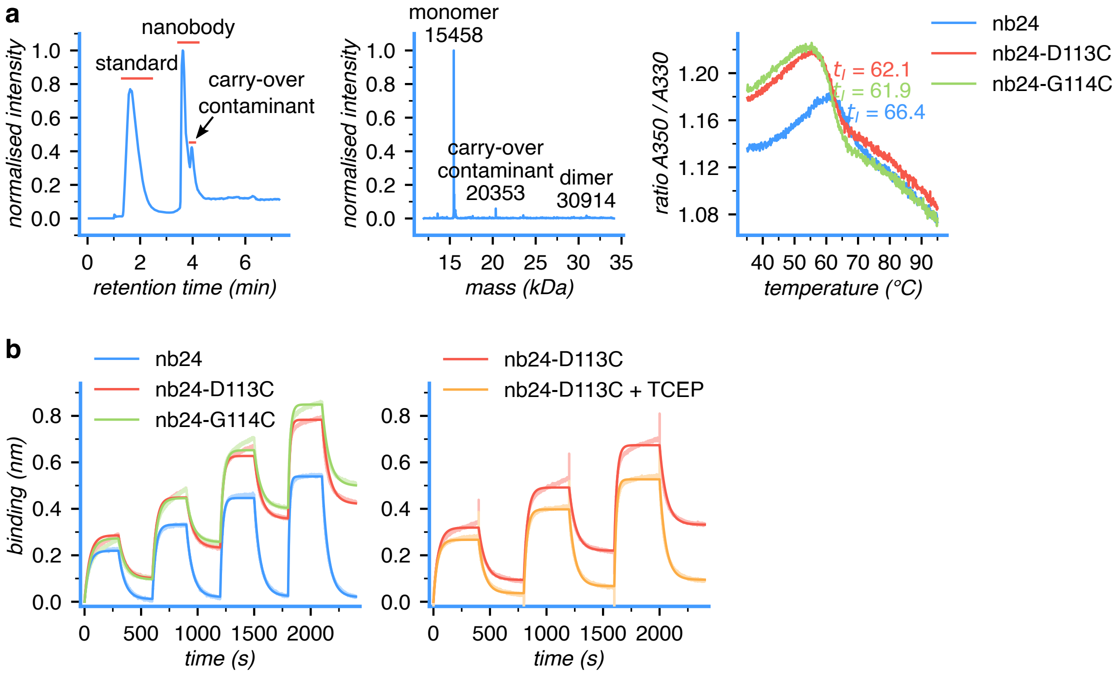

**c**

```
 0 MIQRTPKIQV YSRHPAENGK SNFLNCYVSG FHPSDIEVDL
     *        *         *          *
40 LKNGERIEKV EHSDLSFSKD WSFYLLYYTE FTPTEKDEYA
80 CRVNHVTLSQ PKIVKWDRDM
```

* (green) candidate site identified by LC-MS/MS

* (red) site not covered by LC-MS/MS and therefore candidate

**Figure S10 | Characterisation of nb24-based PEABS. (a)** Characterisation of nb24-D113C, the cysteine mutant used as basis for PEABS systems, using intact protein LC-MS and nanoDSF melting profiles. **(b)** BLI binding profiles of nb24 and its cysteine mutants to β2m (biotinylated and immobilised to streptavidin biosensors) show comparable dissociation constants among the variants. Adding 8 eq TCEP improves the fit and brings the fitted $k_D$ of nb24-D113C to ~100 nM, essentially identical to that of wild-type nb24 (~105 nM). **(c)** β2m modified by nb24-D113-3c-bio was digested with chymotrypsin and analysed by LC-MS/MS. Regions covered by the analysis are highlighted blue. One site, K75, was identified as a potential candidate for the modification site (green star), and three more sites – K41, K48 and K58, were in regions not covered by the analysis, and are therefore potential sites as well.

### Supplementary Notes

#### Linker design

To estimate the lengths of the various PEABS linkers, starting structures were drawn in ChemDraw and imported into GaussView as MOL files. After initial structure clean up and automatic addition of hydrogen atoms, geometry was optimised using density functional theory (DFT) calculations with the B3LYP hybrid functional [1, 2] and the 6-31G basis set [3] as implemented in Gaussian 03 [4]. Converged structures were exported as PDB files and loaded into PyMOL for visualisation.

Small-scale *in vitro* experiments with fluorophenols led to two observations: (1) that these leaving groups would increase the reactivity of their esters towards amines with increasing fluorination; and (2) that the fluorine atoms in the *ortho* position relative to the hydroxyl protect them from hydrolysis. To gain better understanding of this, we performed DFT simulations of the aminolysis of 2,6-difluorophenyl acetate and 2,3,5,6-tetrafluorophenylacetate. First, geometry optimisations of starting materials, intermediates and products in the gas phase were carried out using DFT calculations with the B3LYP hybrid functional (Becke [1], 3 parameter Lee-Yang-Parr [2]) and the 6-31++G basis set [3] as implemented in Gaussian 03 [4]. The tetrahedral intermediate was used as a starting structure for scanning the coordinate of the alcohol hydrogen along the axis of its movement towards the nitrogen (i.e. towards the starting material) or towards the phenol oxygen (i.e. towards the product). These scans were conducted at loose convergence criteria and using the 6-31G basis set without diffuse functions. The transition states were identified as the highest energy coordination along the scan, and optimised as described above with the 6-31++G basis set using the Berny algorithm for transition states [5]. All transition states were then verified by the intrinsic reaction coordinate (IRC) method [6].

Several reaction mechanisms were explored, with the only one that consistently converged and predicted the correct compounds forming being that shown in Figure S11b. The predicted activation energies ($\Delta G^{\ddagger}{}_{1}$), 31.89 kcal/mol (difluorophenol) and 29.98 kcal/mol (tetrafluorophenol), correspond to a predicted 25-fold higher reaction rate in the latter and mirror our first experimental observation. The high energy tetrahedral transition state TS1 helps capture the potential effect of steric hindrance by the *ortho* position fluorine atoms and explain our second observation. While the conditions of these simulations were not physiological, they helped point towards more fluorinated phenols, and particularly 2,3,6-trifluorphenol and 2,3,5,6-tetrafluorophenol, as stable but more reactive groups for the next generation of cleavable linkers.

#### The PEABS experimental protocol

In this note, we discuss the considerations and observations that have informed the different parameters of the PEABS formation protocol.

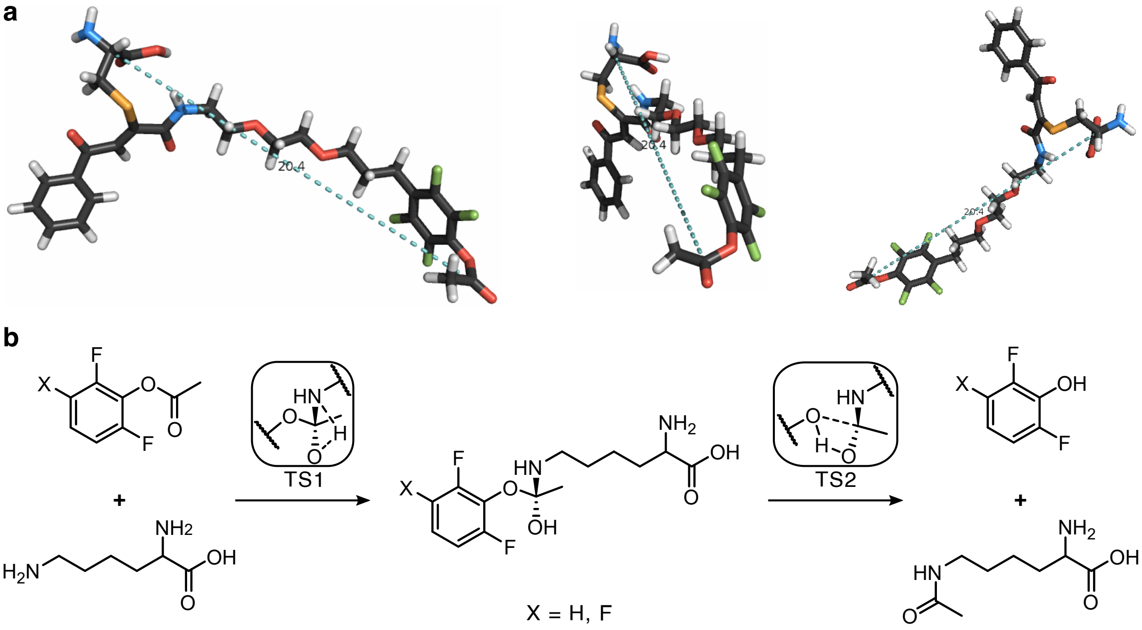

**Figure S11 | *In silico* analysis of PEABS linkers. (a)** A representative optimised structure of a loaded linker (**3d-acetate**) bound to cysteine at the antibody-binding domain, calculated using density functional theory (DFT). The same structure is viewed from several angles. The distance between the Cα atom of the cysteine and the reactive carbonyl carbon is predicted to be 20.4 Å. **(b)** Mechanism of fluorophenyl ester aminolysis according to simulations. Step 1 is the rate determining step, and is expected to be significantly influenced by steric hindrance from the *ortho* position fluorine atoms. **TS** transition state (only parts differing from the intermediate are shown).

#### Nanobody reduction

Due to the small size and dynamic nature of nanobodies, as well as the strategic placement of engineered cysteine residues for PEABS linker conjugation, cysteine-mutant nanobodies were found to be highly reactive, forming nanobody dimers and oxidised cysteine species *in vitro*. These species cannot be conjugated to the benzoylacrylic functional group, and must therefore be reduced prior to conjugation. However, the structurally necessary internal disulphide bond within these proteins must be retained during reduction.

Using a designed nanobody (DesAb) with an exposed cysteine on its CDR3 loop (A112C), we determined that 5–10 molar equivalents of TCEP successfully diminished the dimer population without significantly affecting the folding state of the monomeric nanobody (Figure S1b). In later experiments, an 8-fold molar ratio of TCEP and quiescent conditions during a 2-hour incubation were found to ideally balance the integrity of the cysteine mutant nanobodies and the reactivity of their cysteine residue towards the PEABS linkers.

Depending on the specific requirements in applications of the PEABS method, the populations of doubly and triply conjugated nanobodies can be lessened by using lower concentrations of TCEP at this step, at the price of a potentially lower yield of PEABS.

##### Preventing aggregation

In preliminary experiments using the synthesised PEABS linkers, we found that under standard aqueous Michael addition conditions, the linker conjugation reaction was inconsistent in its rate. Often, the linker would precipitate out of solution before binding the nanobody. Systematic examination revealed that parameters contributing to linker precipitation included alkaline pH, high reaction temperature, low DMSO content, salt content (Figure S1c) and most notably shaking of the reaction vessel. On the other hand, higher DMSO concentrations caused misfolding of the nanobody, leading to its aggregation and off-target reactions with the linker.

To solve this, we tested an array of mass spec–compatible detergents, and found that addition of n-dodecyl-β-maltoside at a final concentration of 0.04 % allowed the linker to remain soluble for long periods of time under quiescent conditions and react with the nanobody's free cysteine at molar ratios as low as 1 : 1. Reducing the ionic strength of the solution to 80 mM and lowering the pH to 6.1 (see below) had no observable negative effects on the reaction and contributed further to linker solubility.

Once the reaction is completed, desalting allows the full removal of the detergent, excess linker and DMSO and readjustment of pH and ionic strength to physiological conditions, without affecting the stability of the PEABS construct. Remaining free linker after desalting precipitates readily and is removed by centrifugation as described in the main text.

##### Linker stoichiometry

Cysteine-reactive electrophiles, and particularly "soft" ones such as the benzoylacrylic functional group, are often used in considerable stoichiometric excess to facilitate full conversion of cysteine residues [7]. However, nanobodies are small and dynamic and feature many sites of potential reactivity, including the two cysteine residues involved in the intramolecular disulphide bond. By varying the stoichiometric ratio of linker to nanobody, as well as reaction temperature, duration, pH and reduction potential, we found the conditions described in the main text – namely, 1 : 10 nanobody : linker ratio, 25 °C without shaking, 50 min at pH 6.1 with 0–3 equivalents of TCEP – led to good conversion of the nanobody to PEABS with minimal double- and triple-modified nanobody (Figure S1d–e).

To maintain the balance between on-target conjugation, off-target reactivity, lab-side practicality and procedure scalability, we would recommend using our experimental conditions as good starting points for case-by-case optimisation, rather than immutable protocols, in applications of the PEABS method to other protein targets.

### Supplementary Methods

#### Sequences

##### Gene: αGFP-S54C as cloned into pET-29b(+)

The amino acid sequence for anti-GFP nanobody was derived from the crystal structure PDB:3OGO. Reverse translation was performed using a Markov chain-based algorithm developed in-house, generating bacterial-like codon sequences by preferentially selecting codon combinations (pairs, triplets) that co-occur naturally in the *E. coli* genome, then ranking them based on features that are known to inhibit or promote soluble expression in bacteria.

`catATGCAAGTCCAGTTAGTTGAATCCGGCGGCGCGTTAGTGCAGCCTGGCGGTTCACTGCGCCTGTCTTGC`
`GCTGCGTCCGGGTTCCCTGTCAACCGTTATTCCATGCGCTGGTACCGCCAGGCACCAGGTAAAGAACGCGAG`
`TGGGTAGCGGGTATGAGT`***`TGC`***`GCGGGCGATCGCTCCAGCTATGAGGATTCTGTCAAAGGCCGTTTCACCATT`
`TCACGTGACGACGCGCGCAACACCGTTTATCTGCAAATGAACAGCCTGAAACCTGAAGATACCGCCGTCTAC`
`TACTGCAACGTGAATGTTGGTTTTGAATACTGGGGCCAGGGCACTCAGGTCACCGTCAGTAGCCTCGAG`

Cloning sites: NdeI/XhoI

##### Primers: Increase nanobody pI and extend His-tag

**fwd** `GTCAGTAGCCGCAGtggcggtcatcaccaccaccaccaccac`

**rev** `CTGCGGCTACTGACggtgacctgagtgccctg`

##### Gene: αGFP-S54C, final construct

`ATGCAAGTCCAGTTAGTTGAATCCGGCGGCGCGTTAGTGCAGCCTGGCGGTTCACTGCGCCTGTCTTGCGCT`
`GCGTCCGGGTTCCCTGTCAACCGTTATTCCATGCGCTGGTACCGCCAGGCACCAGGTAAAGAACGCGAGTGG`
`GTAGCGGGTATGAGT`***`TGC`***`GCGGGCGATCGCTCCAGCTATGAGGATTCTGTCAAAGGCCGTTTCACCATTTCA`
`CGTGACGACGCGCGCAACACCGTTTATCTGCAAATGAACAGCCTGAAACCTGAAGATACCGCCGTCTACTAC`
`TGCAACGTGAATGTTGGTTTTGAATACTGGGGCCAGGGCACTCAGGTCACCGTCAGTAGCCGCAGTGGCGGT`
`CATCACCACCACCACCACCACTGA`

##### Protein: αGFP-S54C

`MQVQLVESGGALVQPGGSLRLSCAASGFPVNRYSMRWYRQAPGKEREWVAGMS`***`C`***`AGDRSSYEDSVKGRFTIS`
`RDDARNTVYLQMNSLKPEDTAVYYCNVNVGFEYWGQGTQVTVSSRSGGHHHHHHH`

##### Primers: αGFP-S54C F103H

**fwd** `gttggtcatgaatactgGGGCCAGGGCACTC`

**rev** `cagtattcatgaccaacATTCACGTTGCAGTAGTAGACG`

**Primers: αGFP-S54C N100D**

**fwd** caacgtggatgttgGTTTTGAATACTGGGGCCAG

**rev** caacatccacgttgCAGTAGTAGACGGCGGTATC

**Primers: αGFP-S54C S34N**

**fwd** ccgttataacatgcgCTGGTACCGCCAGGC

**rev** cgcatgttataacggTTGACAGGGAACCCGG

**Primers: αGFP-S54C S59P**

**fwd** gatcgcccgagcTATGAGGATTCTGTCAAAGGCCG

**rev** gctcgggcgatcGCCCGCGCAACTCATAC

**Gene: GFP in pHAT3 (modified)**

pHAT3 vector carrying the gene for GFP was generously gifted by Dr Marko Hyvönen and Dr Matthew Watson (Department of Biochemistry, University of Cambridge), and minor adjustments were made at the C-terminus.

ATGAACACCATTCATCACCATCACCATCACAACACTAGTGGATCTGGTGGTGGTGGCGGCCGCCTGGTTCCG
AGGGGATCCATGGATATCGAATTCATGGTGAGCAAGGGCGAGGAGCTGTTCACCGGGGTGGTGCCCATCCTG
GTCGAGCTGGACGGCGACGTAAACGGCCACAAGTTCAGCGTGTCCGGCGAGGGCGAGGGCGATGCCACCTAC
GGCAAGCTGACCCTGAAGTTCATCTGCACCACCGGCAAGCTGCCCGTGCCCTGGCCCACCCTCGTGACCACC
CTGACCTACGGCGTGCAGTGCTTCAGCCGCTACCCCGACCACATGAAGCAGCACGACTTCTTCAAGTCCGCC
ATGCCCGAAGGCTACGTCCAGGAGCGCACCATCTTCTTCAAGGACGACGGCAACTACAAGACCCGCGCCGAG
GTGAAGTTCGAGGGCGACACCCTGGTGAACCGCATCGAGCTGAAGGGCATCGACTTCAAGGAGGACGGCAAC
ATCCTGGGGCACAAGCTGGAGTACAACTACAACAGCCACAACGTCTATATCATGGCCGACAAGCAGAAGAAC
GGCATCAAGGTGAACTTCAAGATCCGCCACAACATCGAGGACGGCAGCGTGCAGCTCGCCGACCACTACCAG
CAGAACACCCCCATCGGCGACGGCCCCGTGCTGCTGCCCGACAACCACTACCTGAGCACCCAGTCCGCCCTG
AGCAAAGACCCCAACGAGAAGCGCGATCACATGGTCCTGCTGGAGTTCGTGACCGCCGCCGGGATCACTCTC
GGCATGGACGAGCTGTACAAATGA

**Protein: GFP (uncleaved)**

MNTIHHHHHHNTSGSGGGGGRLVPRGSMDIEFMVSKGEELFTGVVPILVELDGDVNGHKFSVSGEGEGDATY
GKLTLKFICTTGKLPVPWPTLVTTLTYGVQCFSRYPDHMKQHDFFKSAMPEGYVQERTIFFKDDGNYKTRAE
VKFEGDTLVNRIELKGIDFKEDGNILGHKLEYNYNSHNVYIMADKQKNGIKVNFKIRHNIEDGSVQLADHYQ
QNTPIGDGPVLLPDNHYLSTQSALSKDPNEKRDHMVLLEFVTAAGITLGMDELYK

**Protein: GFP (thrombin-cleaved)**

GSMDIEFMVSKGEELFTGVVPILVELDGDVNGHKFSVSGEGEGDATYGKLTLKFICTTGKLPVPWPTLVTTL
TYGVQCFSRYPDHMKQHDFFKSAMPEGYVQERTIFFKDDGNYKTRAEVKFEGDTLVNRIELKGIDFKEDGNI
LGHKLEYNYNSHNVYIMADKQKNGIKVNFKIRHNIEDGSVQLADHYQQNTPIGDGPVLLPDNHYLSTQSALS
KDPNEKRDHMVLLEFVTAAGITLGMDELYK

**Primers: GFP K3S**

**fwd** gagcagcggcgAGGAGCTGTTCACCGGG

**rev** cgccgctgctcACCATGAATTCGATATCCATGGATCC

**Primers: GFP K41R**

**fwd** cggccgcctgACCCTGAAGTTCATCTGCAC

**rev** caggcggccgTAGGTGGCATCGCCCTC

**Primers: GFP K52P**

**fwd** cggcccgctgCCCGTGCCCTGGC

**rev** cagcgggccgGTGGTGCAGATGAACTTCAGG

**Primers: GFP K131S**

**fwd** cttcagcgaggacGGCAACATCCTGGGGC

**rev** gtcctcgctgaagTCGATGCCCTTCAGCTC

**Primers: GFP K140R**

**fwd** gcaccgcctggAGTACAACTACAACAGCCACAAC

**rev** ccaggcggtgcCCCAGGATGTTGCCGTC

**Primers: GFP K162R**

**fwd** catccgcgtgaacTTCAAGATCCGCCACAACATC

**rev** gttcacgcggatgCCGTTCTTCTGCTTGTCGG

**Primers: GFP K166R**

**fwd** cttccgcatccgCCACAACATCGAGGACG

**rev** cggatgcggaagTTCACCTTGATGCCGTTC

**Primers: GFP K209T**

**fwd** gagcaccgaccCCAACGAGAAGCGCG

**rev** ggtcggtgctcAGGGCGGACTGGG

**Primers: GFP K214N**

**fwd** cgagaaccgcgATCACATGGTCCTGCTG

**rev** cgcggttctcgTTGGGGTCTTTGCTCAG

**Primers: GFP K238R**

**fwd** ctgtaccgctgataagGATGCATAAGCTTGAGTATTCTATAGTGTC

**rev** cttatcagcggtacagCTCGTCCATGCCGAGAG

**Gene: nb24**

The amino acid sequence for the anti-β2m nanobody nb24 was derived from the crystal structure PDB:4KDT. Reverse translation was performed using the same method described for αGFP above.

ccATGgaaCAAGTGCAGTTACAGGAATCTGGTGGTGGTTCCGTGCAGGCGGGCGGCAGTTTACGCCTGTCGT
GTGCTGCTTCAGGCTATACCGATTCTCGCTACTGCATGGCATGGTTTCGACAAGCGCCAGGCAAGGAACGTG
AATGGGTAGCACGTATTAACAGCGGTCGCGATATTACTTACTATGCGGACAGTGTTAAGGGGCGCTTTACCT
TCAGTCAGGATAACGCAAAGAACACCGTTTATCTGCAGATGGATTCGCTGGAACCAGAAGATACCGCCACTT
ATTACTGCGCGACAGATATTCCTTTACGCTGCCGTGATATCGTGGCAAAAGGCGGTGATGGCTTTCGCTATT
GGGGTCAGGGAACTCAAGTTACTGTCAGTTCACTCGAG

Cloning: using NcoI/XhoI into pET-28b(+)

**Protein: nb24**

MEQVQLQESGGGSVQAGGSLRLSCAASGYTDSRYCMAWFRQAPGKEREWVARINSGRDITYYADSVKGRFTF
SQDNAKNTVYLQMDSLEPEDTATYYCATDIPLRCRDIVAKGGDGFRYWGQGTQVTVSSLEHHHHHH

**Primers: nb24 D113C**

**fwd** cggttgcggctttcGCTATTGGGGTCAGGGAACTCAAG

**rev** gaaagccgcaaccgCCTTTTGCCACGATATCACGGCAG

**Primers: nb24 G114C**

**fwd** ggtgattgctttcgCTATTGGGGTCAGGGAAC

**rev** cgaaagcaatcaccGCCTTTTGCCACGATATC

#### Protein properties

| protein | amino acids | unprocessed molecular weight (Da)[a] | final molecular weight (Da)[b] | attenuation coefficient $\varepsilon$ ($M^{-1}\cdot cm^{-1}$)[a] | isoelectric point[a] |
|---|---|---|---|---|---|
| αGFP | 127 | 14164.6 | 14162.6 | 27000 ± 60 | 8.6[c] |
| αGFP-S54C | 127 | 14180.7 | 14178.7 | 27000 ± 60 | 8.5[c] |
| αGFP-S54C-F103H | 127 | 14170.7 | 14168.7 | 27000 ± 60 | 8.5 |
| αGFP-S54C-N100D | 127 | 14181.7 | 14179.7 | 27000 ± 60 | 7.8 |
| αGFP-S54C-S34N | 127 | 14207.7 | 14205.7 | 27000 ± 60 | 8.5 |
| αGFP-S54C-S59P | 127 | 14190.8 | 14188.7 | 27000 ± 60 | 8.5 |
| nb24 | 138 | 15474.1 | 15470.0 | 28550 ± 130 | 6.3 |
| nb24-D113C | 138 | 15462.1 | 15458.1 | 28550 ± 130 | 6.5 |
| nb24-G114C | 138 | 15520.1 | 15516.1 | 28550 ± 130 | 6.3 |
| His-GFP | 271 | 30357.2 | 30337.2 | 21950 ± 60 | 6.0 |
| GFP | 246 | 27721.3 | 27701.3 | 21950 ± 60 | 5.4 |
| GFP-K#S | 246 | 27680.2 | 27660.2 | 21950 ± 60 | 5.3 |
| GFP-K#R | 246 | 27749.4 | 27729.3 | 21950 ± 60 | 5.4 |
| GFP-K52P | 246 | 27690.3 | 27670.2 | 21950 ± 60 | 5.3 |
| GFP-K209T | 246 | 27694.3 | 27674.2 | 21950 ± 60 | 5.3 |
| GFP-K214N | 246 | 27707.3 | 27687.2 | 21950 ± 60 | 5.3 |
| β2m | 100 | 11862.4 | 10860.3 | 20000 ± 60 | 6.0 |

[a] Values predicted by Expasy ProtParam. [b] Predicted mass following disulphide bond or active site formation. [c] We found it is beneficial to keep the pI of the nanobody outside the range experienced in PEABS experiments (6.1–7.4), and therefore introduced the modified C-terminus shown above.

#### Molecular biology

##### Chemically competent bacterial cells

This is a modified version of Inoue's method [8]. XL10 gold (Agilent) bacteria were plated on LB+Agar plates with selective antibiotics, and single colonies were used to inoculate 10 mL LB cultures and incubated at 37 °C overnight. 240 mL LB media containing 10 mM $MgSO_4$ and 10 mM $MgCl_2$ were inoculated with a 1:100 diluted O/N culture and incubated at 37 °C until $0.3 < OD_{600} < 0.5$. The culture was incubated on ice for 30 min, then cells were pelleted (15 min / 4 °C / 4000 × g), re-suspended in 240 mL ice-cold 100 mM $MgCl_2$, incubated on ice 10 min and pelleted again under same conditions. Pellet was resuspended in 120 mL ice-cold "TB Buffer" (250 mM KCl + 10 mM HEPES stock + 15 mM $CaCl_2$ + 55 mM $MnCl_2$), incubated at 4 °C for 10 min and pelleted. Pellet was then resuspended again in 5 mL ice-cold "TB Buffer", incubated at 4 °C for 10 min and pelleted one last time. Finally, pellet was re-suspended in 5 mL ice-cold "TB Buffer" + 7 % DMSO (v/v) and incubated on ice. The cells were aliquoted into 100 µL portions, shock-frozen in liquid $N_2$ and stored at -80 °C.

##### Choice of site mutations

Site mutations for affinity reduction were chosen using a two-step process. First, the unbound structure of the αGFP nanobody was uploaded to the CamSol Combination tool [9], and sites on the GFP-binding surface where mutations were found to stabilise – or at least not dramatically destabilise – the free nanobody, were noted. Then, the GFP-bound structure was uploaded to the same platform, and the sites identified in the previous scan were explicitly listed as custom mutation sites. Mutations that had a predicted destabilising effect on the complex, but not on the nanobody itself, were selected as candidates for affinity reduction.

Substitute residues for GFP lysines were selected similarly. The structure of GFP was uploaded to CamSol Combination [9] and the lysine residues to be removed were listed as custom mutation sites. For each site, the substitute that was predicted to cause the least destabilisation to the structure was selected. Where no suitable substitutes were identified using this tool, arginine (K → R) was used by default.

##### Site-directed mutagenesis

Template plasmids were mixed with forward and reverse primers, dNTP mix, HF buffer, DMSO and Phusion polymerase enzyme (NEB) according to the manufacturer's protocol. PCR reactions were performed on a Bio-Rad C1000 Touch Thermal Cycler using the sequence: 98 °C for 2 minutes; 20 × (98 °C for 10 sec; annealing temperature (varies) for 30 sec; 72 °C for 165 sec); then another 5 × (98 °C for 10 sec; annealing temperature - 10 °C for 30 sec; 72 °C for 165 sec); and finally 72 °C for 10 min. The annealing temperature for primers designed using the Naismith method [10] was the primer–primer overlap's melting temperature plus 3 to 4 °C. Products were digested using DpnI (NEB), purified using a GeneJET PCR Purification kit (Thermo Scientific), quantified on a NanoDrop and transformed at 1 µL (typically, 100–150 ng) into chemically competent XL10 bacteria (made as described). Bacteria were selected on antibiotics-containing LB+agar plates. Several colonies were picked up, grown overnight, and used to harvest plasmids using a GeneJET Plasmid Miniprep kit (Thermo Scientific).

#### Protein expression and purification

Gene integrity and correctness was verified by sequencing at the DNA Sequencing Facility, Department of Biochemistry, University of Cambridge. Plasmids were then transformed into chemically competent *E. coli* BL21-gold(DE3) (Agilent) or SHuffle T7 (NEB) by adding 10 ng of plasmid DNA into a vial of competent bacteria and applying heat shock of 42 °C for 45 seconds. Transformed cells were diluted 10-fold into pre-warmed SOC medium, recovered for 45 minutes at 37 °C under 300 rpm shaking, then selected by resistance to antibiotics (ampicillin or kanamycin) on LB-agar plates. Colonies from overnight growth were picked into liquid LB, grown overnight, supplemented with glycerol (final 10 %), flash frozen in liquid $N_2$ and kept at -80 °C. For expression tests, solubility tests and protein purifications, fresh antibiotics selection plates were streaked with bacteria from the -80 °C stocks, and colonies from overnight-grown plates were picked and used to inoculate starter cultures, which in turn – upon overnight growth – were used to inoculate expression cultures by 100-fold dilution.

##### General procedure: expression and affinity purification of soluble proteins

1 L of LB media was inoculated with an O/N starter culture of cells. Cells were grown to $OD_{600}$ of 0.7–0.9 at 37 °C; then expression was induced by adding IPTG to a final concentration of 0.5 mM. Cells were cultured overnight at 18 °C, then harvested from the media by centrifugation (25 min / 4 °C / 6200 × g) and resuspended in 50 mL buffer. The resuspended bacteria were then pelleted again under similar conditions, resuspended in 50 mL buffer + 10 mM imidazole and supplemented with a tablet of cOmplete EDTA-free protease inhibitor cocktail (Roche). After 30 min incubation on ice, the cells were sonicated using a Sonics Vibra-Cell for 5 minutes of active time at 40 % amplitude, in 15 sec on 45 sec off pulses. The clarified lysate was centrifuged (15 min / 4 °C / 39000 × g) to remove cell debris and insoluble or aggregated proteins. Proteins were purified from the 0.45 µm–filtered supernatant of the lysate using a gravitational column loaded with 4 mL Ni-NTA Agarose resin (Thermo) and equilibrated against buffer + 10 mM imidazole. The flow through was re-introduced into the column for maximal binding; this was repeated twice more. Non-specifically bound proteins were then washed with 50 mL buffer + 55 mM imidazole, and the tagged protein was eluted with 6 × 8 mL fractions of buffer + 300 mM imidazole and immediately supplemented with 0.2 mM ethylenediaminetetraacetic acid (EDTA) to prevent aggregation on nickel. Fractions were assessed by SDS-PAGE, and the more pure product-containing fractions were pooled.

**αGFP nanobody and its variants** pET-29b(+) vector carrying the gene for αGFP or one of its variants was transformed into SHuffle T7 bacteria as described, and selected for kanamycin resistance. Expression and affinity purification followed the general procedure, with 50 mM potassium phosphate + 500 mM NaCl, pH 7.0 as the buffer. Unwanted proteins were precipitated using ammonium sulphate at 30 % saturation (170 mg/mL) on a tube roller at 4 °C for at least 1 hour, and pelleted by centrifugation (15 min / 4 °C / 10000 × g). The supernatant was decanted into a fresh tube, supplemented with an additional 300 mg/mL ammonium sulphate (final saturation 75 %), incubated on a tube roller at 4 °C for at least 1 hour, and pelleted by centrifugation. The supernatant was discarded, and the pellet resuspended in 3 mL buffer, mixed thoroughly at 4 °C, and centrifuged to remove suspended material (5 min / 4 °C / 5000–10000 × g). The clarified supernatant was reduced using 200 µM TCEP for 2 h at rt, then loaded onto a HiLoad 16/600 Superdex 75 prep-grade size exclusion chromatography column (GE Healthcare) on an Äkta Prime system (GE Healthcare). The buffer used for resuspension, equilibration and elution was 40 mM MES + 40 mM NaCl, pH 6.1. Peak fractions were collected, pooled, flash frozen in liquid $N_2$ and stored at -80 °C.

**GFP** GFP in pHAT3 was transformed into BL21(DE3) cells (New England Biolabs) as described, and selected for ampicillin resistance. Expression and affinity purification followed the general procedure, with PBS as the buffer and 40 mM imidazole in the washing step. Imidazole was removed by dialysis to PBS at 4 °C overnight. The His-tag was cleaved using thrombin protease (GE Healthcare) according to manufacturer instructions. The cleaved protein was purified by reverse affinity chromatography followed by size-exclusion chromatography, eluting with PBS. Peak fractions were collected, pooled, flash frozen in liquid $N_2$ and stored at -80 °C.

**GFP variants** Mutated GFP in pHAT3 was transformed into BL21(DE3) and selected for ampicillin resistance. Expression and affinity purification followed the general procedure, with PBS as the buffer and 40 mM imidazole in the washing step. Unwanted proteins were precipitated using ammonium sulphate at 30 % saturation (170 mg/mL) on a tube roller at 4 °C for at least 1 hour and pelleted by centrifugation (15 min / 4 °C / 5000 × g). The supernatant was decanted into a fresh tube, supplemented with an additional 300 mg/mL ammonium sulphate (final saturation 75 %), incubated on a tube roller at 4 °C overnight, and pelleted by centrifugation (15 min / 4 °C / 12000 × g). The supernatant was discarded, and the pellet resuspended in 2 mL buffer, mixed thoroughly on ice, and centrifuged to remove suspended material (10 min / 4 °C / 20000 × g). Protein in the clarified supernatant was quantified on NanoDrop and approx. 2 mg of protein was transferred to a fresh tube, supplemented with 20 U bovine thrombin (Antibodies.com) and incubated on a tube roller in the dark at room temperature overnight. The digested GFP variant was then loaded onto a HiLoad 16/600 Superdex 75 prep-grade size exclusion chromatography column (GE Healthcare) on an Äkta Prime system (GE Healthcare). The buffer used for equilibration and elution was PBS pH 7.3. Peak fractions were collected, pooled, brought to 30 µM, flash frozen in liquid $N_2$ and stored at -80 °C.

**nb24 and its variants** pET-28b(+) vector carrying the gene for nb24 or one of its variants was transformed into SHuffle T7 bacteria as described, and selected for kanamycin resistance. Expression followed the general procedure with PBS as the buffer.

Affinity purification was conducted using an AmMag™ SA Plus Semi-automated System (Genscript), with PBS as buffer and PBS-washed His Mag Sepharose Excel magnetic beads (Cytiva) for capturing the his-tagged nanobodies. The nanobody was then loaded onto a Superdex 75 10/300 exclusion chromatography column (GE Healthcare) on an Äkta Pure system (GE Healthcare). The buffer used for equilibration and elution was 40 mM MES + 40 mM NaCl, pH 6.1. Peak fractions were collected, pooled, flash frozen in liquid $N_2$ and stored at -80 °C.

**β2m** Recombinant human beta-2 microglobulin was a gift from Dr Cristina Visentin and Prof. Stefano Ricagno (University of Milan), produced as previously described [11, 12].

#### Organic synthesis

##### General methods

Where not mentioned otherwise, general procedures are for 1 mmol of compound.

**Sonogashira coupling (1a-3, 2a-3, 3a-3 and 3b-3)** Example for 2 mmol: Iodinated fluorophenol $X_1$ (2 mmol) was weighed in a round-bottom flask. Alkyne $X_2$ (3 mmol, 1.5 eq), 31 mg copper(I) iodide (0.16 mmol, 8 %mol) and 141 mg bis(triphenylphosphine) palladium(II) dichloride (0.2 mmol, 10 %mol) were added, dissolved in dry DMF (5 mL) and stirred under $N_2$ for 5 min. 558 µL triethylamine (405 mg, 4 mmol, 2 eq) was added and the mixture was stirred for $X_3$ (time) at RT. Purifications varied, see individual compounds.

**Boc deprotection** Boc-protected amine $X_1$ was weighed in a round-bottom flask and dissolved in anhydrous DCM at 0 °C for a final 0.1 M (10 mL). Trifluoroacetic acid (994 µL, 13 mmol, 13 eq) was added slowly and stirred on ice for 15 min. Reaction was then brought to room temperature and stirred for another $X_2$ (time). Reaction was monitored by TLC, and once completed, the solvent and residual acid were evaporated *in vacuo* as an azeotrope using toluene in 3–5 rounds.

**HATU amide coupling (1a, 2a, 2b, 3c, 3d)** 1.1 mmol acid $X_1$ (1.1 eq) and 475 mg HATU (1.25 mmol, 1.25 eq) were weighed into a dry round-bottom flask and dissolved in dry DMF (1.5 mL) at 0 °C. After 10 min, 436 µL *N*,*N*-diisopropylethylamine (2.5 mmol, 2.5 eq) were added dropwise and the solution was stirred for $X_2$ (temperature and time). In a separate vial, amine $X_3$ (1.0 mmol) was dissolved in dry DMF (1.5 mL), then added dropwise to the stirred reaction mixture. The reaction was brought to room temperature and stirred for $X_4$ (time). The reaction mixture was then diluted 20 times in EtOAc, acidified by washing twice with 3M HCl and then washed twice with $H_2O$ and twice with brine. The organic layer was dried over $MgSO_4$, filtered and concentrated *in vacuo*, then purified by flash chromatography on silica gel (mobile phase $X_5$).

**EDC esterification (1a-rho, 2a-rho, 2b-rho, 3c-rho, 3c-bio, 3d-rho)** Example for 100 µmol: acid $X_1$ (2.5 eq), 23 mg *N*-(3-Dimethylaminopropyl)-*N* -ethylcarbodiimide

| *symbol* | *name* | *formula* | *mono-isotopic mass* | *molecular weight* |
|---|---|---|---|---|
| **1a-2** | 2-fluoro-4-iodophenol | $C_6H_4FIO$ | 237.93 | 238.00 |
| **1a-3** | 4-(N-Boc-3-aminoprop-1-yn-1-yl)-2-fluorophenol | $C_{14}H_{16}FNO_3$ | 265.11 | 265.28 |
| **1a-4** | 3-(3-fluoro-4-hydroxyphenyl)prop-2-yn-1-aminium trifluoroacetate | $C_9H_9FNO^+ \cdot C_2F_3O_2^-$ | 166.07 | 166.17 + 113.02 = 279.19 |
| **1a** | Linker 1a | $C_{19}H_{14}FNO_3$ | 323.10 | 323.32 |
| **1a-rho** | Linker 1a Rhodamin B ester | $C_{47}H_{43}FN_3O_5^+ \cdot Cl^-$ | 748.32 | 748.87 + 35.45 = 784.32 |
| **2a-3** | 4-(N-Boc-3-aminoprop-1-yn-1-yl)-2,6-difluorophenol | $C_{14}H_{15}F_2NO_3$ | 283.10 | 283.27 |
| **2a-4** | 3-(3,5-difluoro-4-hydroxyphenyl)prop-2-yn-1-aminium trifluoroacetate | $C_9H_8F_2NO^+ \cdot C_2F_3O_2^-$ | 184.06 | 184.17 + 113.02 = 297.19 |
| **2a** | Linker 2a | $C_{19}H_{13}F_2NO_3$ | 341.09 | 341.31 |
| **2a-rho** | Linker 2a Rhodamin B ester | $C_{47}H_{42}F_2N_3O_5^+ \cdot Cl^-$ | 766.31 | 766.87 + 35.45 = 802.32 |
| **2b-3** | 4-(N-Boc-3-aminopropan-1-yl)-2,6-difluorophenol | $C_{14}H_{19}F_2NO_3$ | 287.13 | 287.31 |
| **2b** | Linker 2b | $C_{19}H_{17}F_2NO_3$ | 345.12 | 345.35 |
| **2b-rho** | Linker 2b Rhodamin B ester | $C_{47}H_{46}F_2N_3O_5^+ \cdot Cl^-$ | 770.34 | 770.90 + 35.45 = 806.35 |
| **c-2** | N-Boc-2-(prop-2-yn-1-yloxy)ethan-1-amine | $C_{10}H_{17}NO_3$ | 199.12 | 199.25 |
| **3a-2** | 2,3,6-trifluoro-4-iodophenol | $C_6H_2F_3IO$ | 273.91 | 273.98 |
| **3c-3** | 4-(3-(2-(Boc-amino)ethoxy)prop-1-yn-1-yl)-2,3,6-trifluorophenol | $C_{16}H_{18}F_3NO_4$ | 345.12 | 345.32 |
| **3c-4** | 4-(3-(2-(Boc-amino)ethoxy)propyl)-2,3,6-trifluorophenol | $C_{16}H_{22}F_3NO_4$ | 349.15 | 349.35 |
| **3c-5** | 2-(3-(2,3,5-trifluoro-4-hydroxyphenyl)propoxy)ethan-1-aminium trifluoroacetate | $C_{11}H_{15}F_3NO_2^+ \cdot C_2F_3O_2^-$ | 250.10 | 250.24 + 113.02 = 363.26 |
| **3c** | Linker 3c | $C_{21}H_{20}F_3NO_4$ | 407.13 | 407.39 |
| **3c-rho** | Linker 3c Rhodamin B ester | $C_{49}H_{49}F_3N_3O_6^+ \cdot Cl^-$ | 832.35 | 832.94 + 35.45 = 868.39 |
| **3c-bio** | Linker 3c biotin ester | $C_{31}H_{34}F_3N_3O_6S$ | 633.21 | 633.68 |
| **d-2** | N-Boc-2-(2-(prop-2-yn-1-yloxy)ethoxy)ethan-1-amine | $C_{12}H_{21}NO_4$ | 243.15 | 243.30 |
| **3d-3** | 4-(3-(2-(2-(Boc-amino)ethoxy)ethoxy)prop-1-yn-1-yl)-2,3,6-trifluorophenol | $C_{18}H_{22}F_3NO_5$ | 389.15 | 389.37 |
| **3d-4** | 4-(2-(3-(2-(Boc-amino)ethoxy)propoxy)ethyl)-2,3,6-trifluorophenol | $C_{18}H_{26}F_3NO_5$ | 393.18 | 393.40 |
| **3d-5** | (3-(3-(2,3,5-trifluoro-4-hydroxyphenyl)propoxy)propoxy)methanaminium trifluoroacetate | $C_{13}H_{19}F_3NO_3^+ \cdot C_2F_3O_2^-$ | 294.13 | 294.29 + 113.02 = 407.31 |
| **3d** | Linker 3d | $C_{23}H_{24}F_3NO_5$ | 451.16 | 451.44 |
| **3d-rho** | Linker 3d Rhodamin B ester | $C_{51}H_{53}F_3N_3O_7^+ \cdot Cl^-$ | 876.38 | 876.99 + 35.45 = 912.44 |

**Table S6 | Summary of molecules synthesised in this work.**

hydrochloride (EDC·HCl) (120 µmol, 1.2 eq) and 3.1 mg 4-dimethylaminopyridine (25 µmol, 0.25 eq) were weighed in an oven-dried round-bottom flask and dissolved in dry DCM (1.5 mL) at 0 °C. In a separate vial, $X_2$ was dissolved in a mixture of dry DMF (90 µL) and dry DCM (1.5 mL), then added dropwise over the course of 30 min to the stirred reaction mixture. The reaction was brought to room temperature and stirred for 4 h. The reaction mixture was then diluted 10-fold in DCM, washed once with $H_2O$ then separated;

aqueous phase was extracted with DCM, the two organic phases were joined, washed twice with $H_2O$ and once with brine, dried over $MgSO_4$, filtered and dried *in vacuo*. The residue was purified by flash chromatography on silica gel (mobile phase $X_3$)

**Alkyne reduction (2b-3, 3c-4, 3d-4)** Alkyne $X_1$ was weighed in an oven-dried 100 mL round-bottom flask. $X_2$ (mass) palladium on carbon (1:10 w/w) was added and the flask was flushed with $N_2$. The mixture was suspended in anhydrous methanol (20 mL) and stirred at room temperature for 10 min. 5–10 L of hydrogen gas were bubbled into the solution while stirring and venting with a needle, and the flask was then stirred under $H_2$ atmosphere for $X_3$ (time). The reaction was quenched by bubbling nitrogen gas as above. Once safe to open, the reaction mixture was filtered through 2 cm of celite, and washed several times with methanol. The solvent was then removed *in vacuo* to dryness, and products were used without further purification.

##### Linker 1a

Reaction scheme: Fig. S12

**2-fluoro-4-iodophenol (1a-2): 238.0 g/mol** 1.28 g sodium hydroxide (32 mmol, 2 eq) was dissolved in $H_2O$ (40 mL, 800 mM) and cooled to 0 °C. To a stirred solution of 2.54 g iodine (10 mmol, 0.625 eq) and 1.66 g potassium iodide (10 mmol, 0.625 eq) in $H_2O$ (40 mL) at 0 °C was added dropwise 1.79g **1a** (16 mmol) and stirred for 10 min, after which the cold sodium hydroxide solution was added slowly and stirred for 45 min at 0 °C. The reaction was quenched with a large excess of ammonium chloride (10 g) and 3.16 g sodium thiosulphate (20 mmol, 1.25 eq) was added to reduce any remaining iodine. The reaction mixture was acidified with HCl, extracted with DCM (3 × 25 mL) then washed with $H_2O$ (3 × 100 mL) and with brine (100 mL). The organic layer was dried over $MgSO_4$, filtered and concentrated *in vacuo*, then purified by flash chromatography on silica gel (100 % DCM).

Yields 992 mg of **1a-2** (4.17 mmol, 26 %).

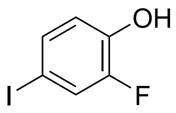

**1a-2**
$C_6H_4FIO$

$^1$H-NMR ($CDCl_3$, δ/ppm): 7.40 (1H, dd, H5), 7.34 (1H, dt, H3), 6.77 (1H, t, H6), 5.13 (1H, s, OH); $^{19}$F-NMR ($CDCl_3$, δ/ppm): -138.0.

**4-(*N*-Boc-3-aminoprop-1-yn-1-yl)-2-fluorophenol (1a-3): 265.28 g/mol Sonogashira coupling** of 3 mmol with $X_1$ = 714 mg **1a-2**; $X_2$ = 698 mg *N*-Boc-propargylamine; $X_3$ = 2 h 30 min. The solution was basified with 50 mL 10 % sodium hydroxide solution and stirred for 10 min. The solution was then diluted with 450 mL $H_2O$ and impurities were extracted using DCM. The aqueous layer was acidified to pH 1–2 using HCl and the resulting protonated product was extracted with EtOAc (2 portions). This organic layer

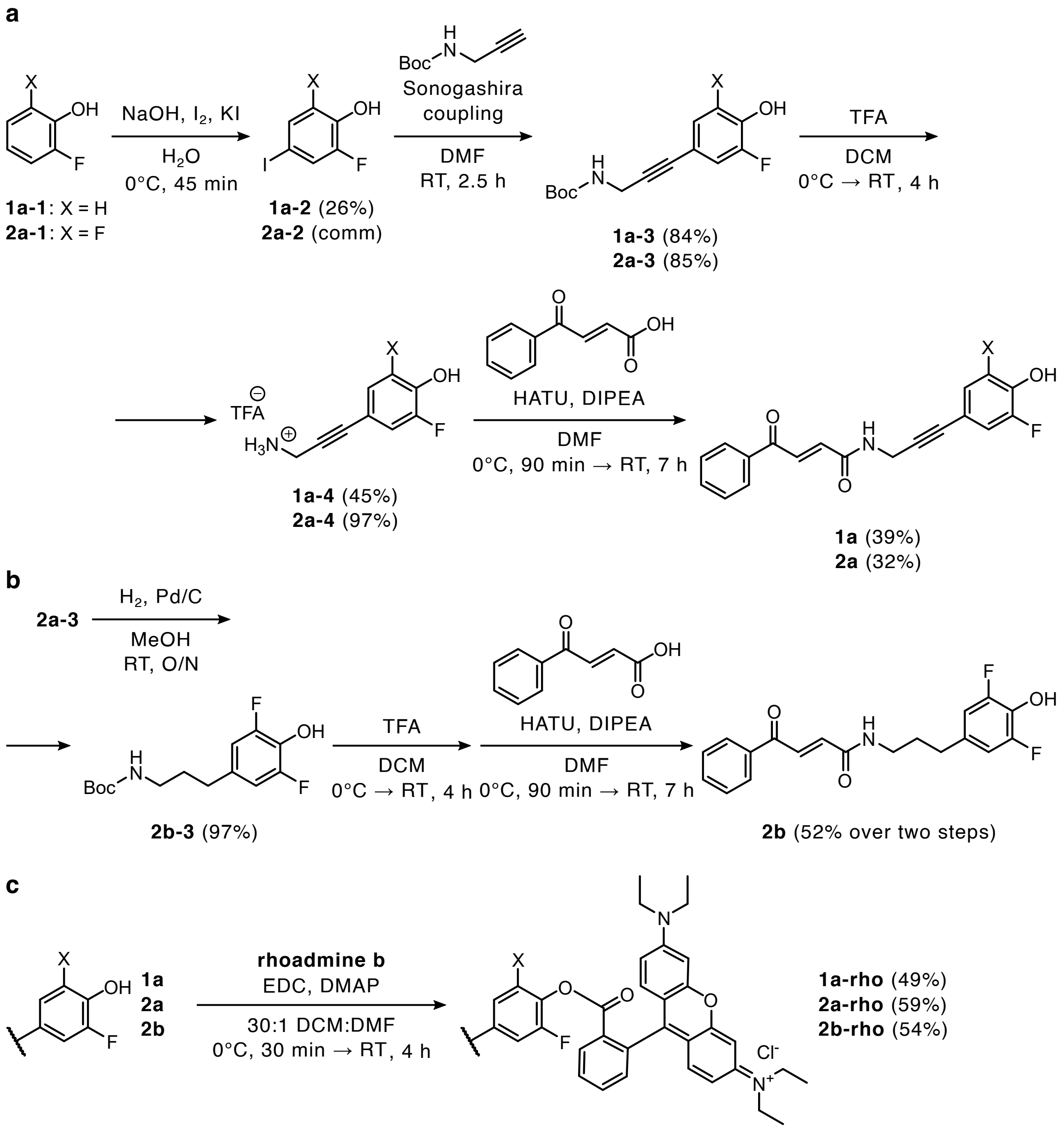

**Figure S12 | Synthetic routes to groups 1 and 2 linkers.** **(a)** Synthetic route to **1a** and **2a**. **(b)** Synthetic route to **2b**. **(c)** Loading rhodamine as a payload. **comm** commercially available; **TFA** trifluoroacetic acid; **DCM** dichloromethane; **DMF** dimethylformamide; **DIPEA** diisopropylethylamine; **EDC** ethyl(dimethylaminopropyl)carbodiimide; **DMAP** dimethylaminopyridine.

was then washed with $H_2O$ (2 portions) and brine (2 portions), dried over $MgSO_4$, filtered and concentrated *in vacuo*. The residue was dry-loaded on silica gel and purified by flash chromatography (4:1 PE:EtOAc).

Yields 668 mg of **1a-3** (2.52 mmol, 84 %) as viscous liquid.

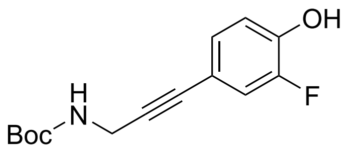

**1a-3**
$C_{14}H_{16}FNO_3$

$^{1}$H-NMR ($CDCl_3$, δ/ppm): 7.10 (1H, dd, H5), 7.06 (1H, dt, H3), 6.91 (1H, t, H6), 5.96 (1H, s, OH), 4.80 (1H, s, NH), 4.11 (2H, s, $CH_2$), 1.47 (9H, s, Boc); $^{19}$F-NMR ($CDCl_3$, δ/ppm): -139.7.

**3-(3-fluoro-4-hydroxyphenyl)prop-2-yn-1-aminium trifluoroacetate (1a-4): 279.19 g/mol** **Boc deprotection** of 1.6 mmol with $X_1$ = 425 mg **1a-3**; $X_2$ = 4 h. Loaded on silica gel and purified by flash chromatography (85:15 DCM:MeOH).

Yields 256 mg at 78 %wt purity of **1a-4** (199 mg pure product + 57 mg excess TFA) (0.71 mmol, 45 %), used without further purification.

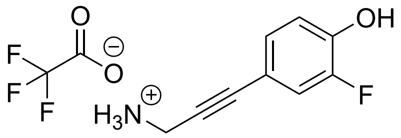

**1a-4**
$C_9H_9FNO^+ \cdot C_2F_3O_2^-$

$^{1}$H-NMR (MeOD, δ/ppm): 7.74 (1H, dd, H5), 7.02 (1H, dt, H3), 6.89 (1H, t, H6), 3.99 (2H, s, $CH_2$); $^{19}$F-NMR (MeOD, δ/ppm): -138.5, -77.0 (TFA).

**Linker 1a: 323.32 g/mol** **HATU amide coupling** of 0.52 mmol with $X_1$ = 119 mg trans-3-benzoylacrylic acid (0.66 mmol, 1.25 eq); $X_2$ = 0 °C for 90 min; $X_3$ = 167 mg **1a-4** (214 mg at 78 %wt); $X_4$ = 7 h; $X_5$ = 30:1 DCM:MeOH.

Yields 65 mg of **1a** (0.2 mmol, 39 %).

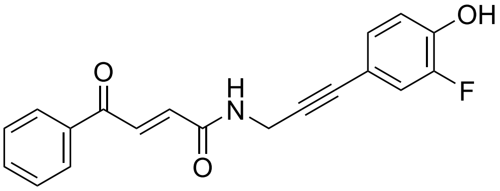

**1a**
$C_{19}H_{14}FNO_3$

$^{1}$H-NMR (MeOD, δ/ppm): 8.04 (2H, d), 7.92 (1H, d, $J$ = 15.3), 7.67 (1H, t), 7.56 (2H, t), 7.12 (1H, dd), 7.07 (1H, dt), 7.03 (1H, d, $J$ = 15.3), 6.85 (1H, t), 4.30 (2H, s); $^{19}$F-NMR (MeOD, δ/ppm): -138.9. Full spectra and assignments appear in Fig. S13.

**1a-rho: 784.33 g/mol** **EDC esterification** of 85 µmol with $X_1$ = 100 mg Rhodamine B; $X_2$ = 27 mg **1a**; $X_3$ = 92:8 DCM:MeOH.

Yields 33 mg of **1a-rho** (42 µmol, 49 %).

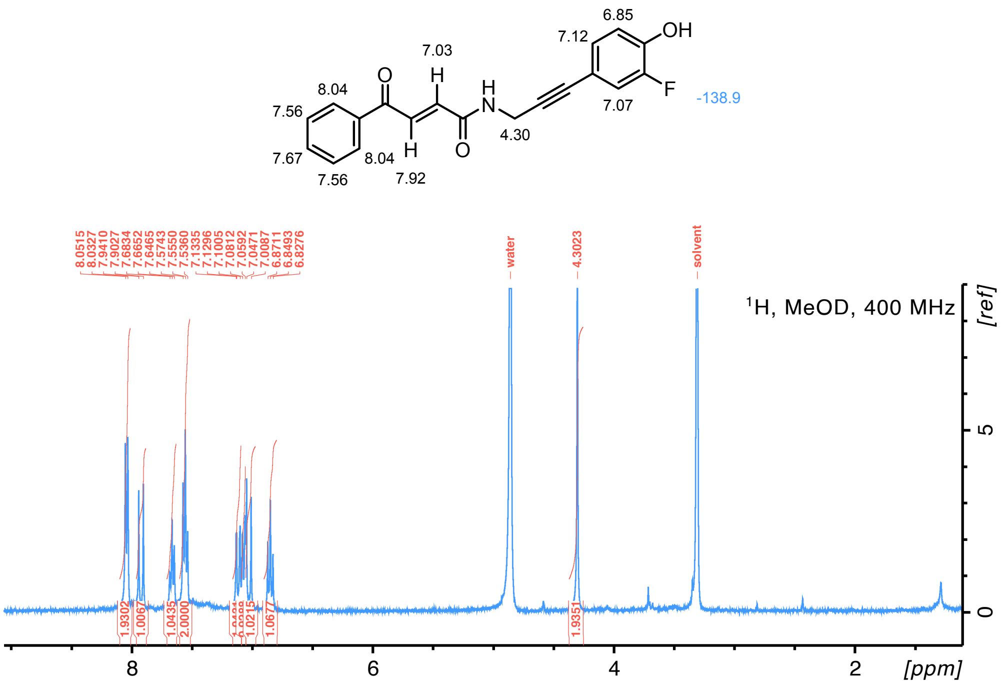

**Figure S13 | Linker 1a NMR spectra**

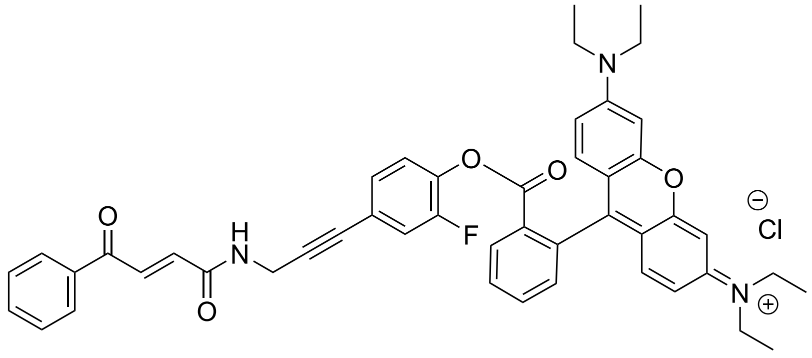

**1a-rho**
$C_{47}H_{43}FN_3O_5^+ \cdot Cl^-$

$^1$H-NMR ($CDCl_3$, δ/ppm): see Fig S14; $^{19}$F-NMR ($CDCl_3$, δ/ppm): -128.5. LC-MS$^+$: 748.3 (*exp* 748.3).

#### Linker 2a

Reaction scheme: Fig. S12

**4-(*N*-Boc-3-aminoprop-1-yn-1-yl)-2,6-difluorophenol (2a-3): 283.27 g/mol Sonogashira coupling** with $X_1$ = 512 mg **2a-2**; $X_2$ = 466 mg *N*-Boc-propargylamine; $X_3$ = 3 h. Solvent was removed *in vacuo* as an azeotrope using toluene (7 rounds of dilution and evaporation), and the residue was dry-loaded on silica gel and purified by flash chromatography (3:1 PE:EtOAc).

Yields 483 mg of **2a-3** (1.7 mmol, 85 %).

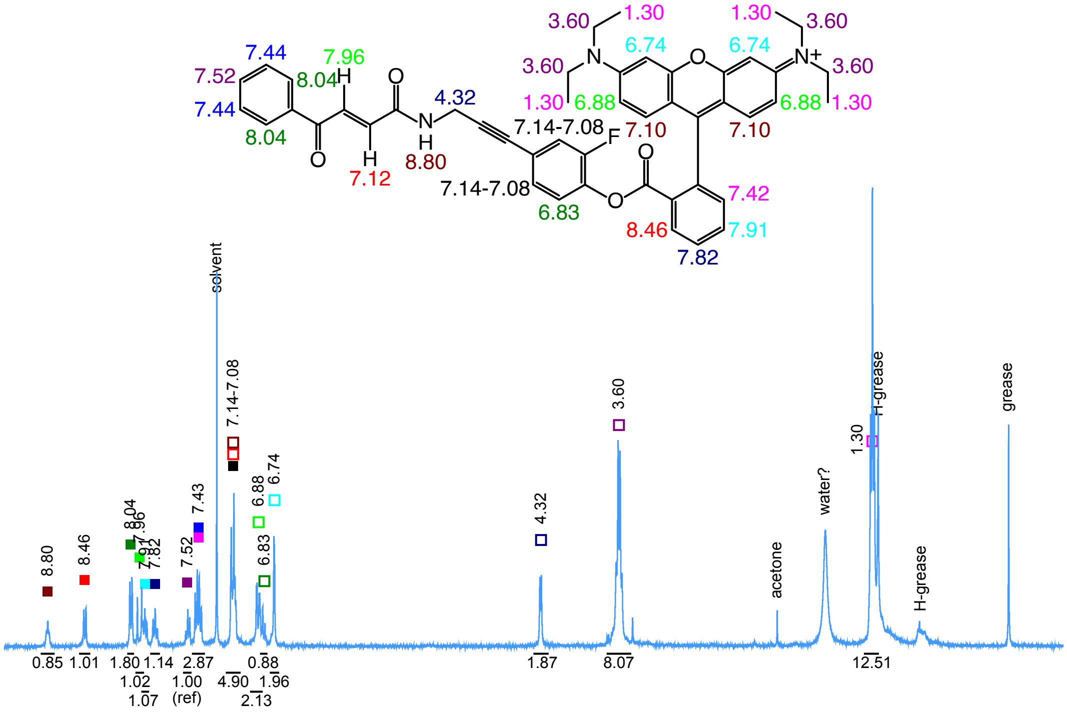

**Figure S14 | 1a-rho NMR spectra**

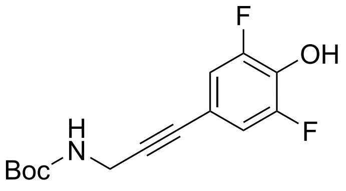

**2a-3**
$C_{14}H_{15}F_2NO_3$

[1]H-NMR ($CDCl_3$, δ/ppm): 6.90 (2H, d, H3 and H5), 6.25 (1H, s-br, OH), 4.82 (1H, s-br, NH), 4.10 (2H, s, $CH_2$), 1.47 (9H, s, Boc); [19]F-NMR ($CDCl_3$, δ/ppm): -134.9.

**3-(3,5-difluoro-4-hydroxyphenyl)prop-2-yn-1-aminium trifluoroacetate (2a-4): 297.18 g/mol** **Boc deprotection** with $X_1$ = 283 mg **2a-3**; $X_2$ = 2 h 30 min. Loaded on silica gel and purified by flash chromatography (85:15 DCM:MeOH).

Yields 346mg of **2a-4** at 83 %wt purity (288 mg pure product + 58 mg excess TFA) (0.97 mmol, 97 %) as a white crystalline solid, used without further purification.

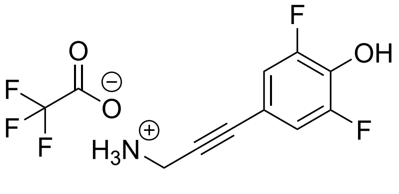

**2a-4**
$C_9H_8F_2NO^+ \cdot C_2F_3O_2^-$

[1]H-NMR (MeOD, δ/ppm): 7.05 (2H, dq, H3 and H5), 3.99 (2H, s, $CH_2$); [19]F-NMR (MeOD, δ/ppm): -135.0, -77.0 (TFA).

**Linker 2a: 341.31 g/mol HATU amide coupling** of 0.4 mmol with $X_1$ = 70 mg trans-3-benzoylacrylic acid (0.4 mmol, 1.0 eq); $X_2$ = 0 °C for 1 hour; $X_3$ = 119 mg **2a-4**; $X_4$ = 2 h; $X_5$ = 64:36 PE:EtOAc.

Yields 44 mg of **2a** (129 µmol, 32 %).

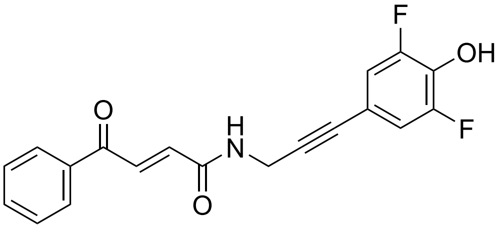

**2a**
$C_{19}H_{13}F_2NO_3$

$^1$H-NMR (MeOD, δ/ppm): 8.04 (2H, d), 7.92 (1H, d, $J$ = 15.3), 7.66 (1H, t), 7.56 (2H, t), 7.02 (1H, d, $J$ = 15.3), 6.98 (2H, s), 4.30 (2H, s); $^{19}$F-NMR (MeOD, δ/ppm): -135.5. Full spectra and assignments appear in Fig. S15.

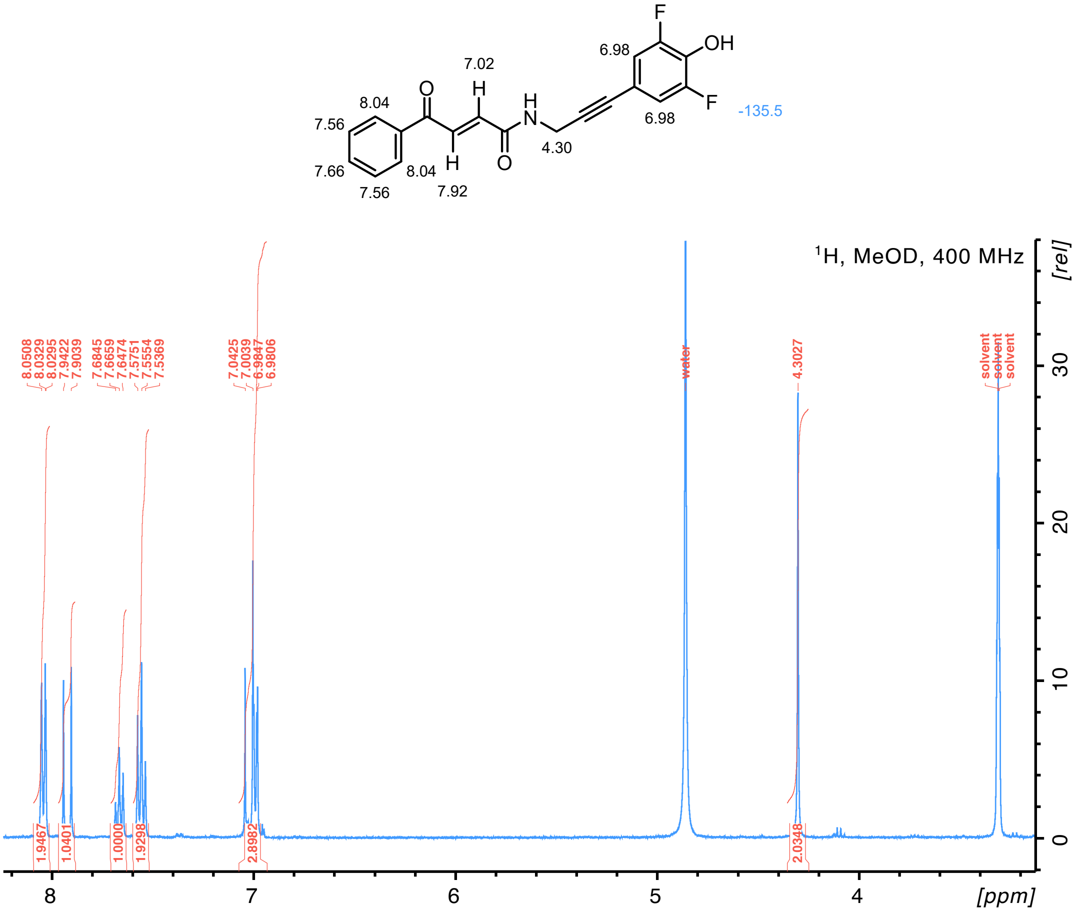

**Figure S15 | Linker 2a NMR spectra**

**2a-rho: 802.32 g/mol EDC esterification** of 70 µmol with $X_1$ = 81 mg Rhodamine B; $X_2$ = 25 mg **2a**; $X_3$ = 9:1 DCM:MeOH.

Yields 33 mg of **2a-rho** (41 µmol, 59 %).

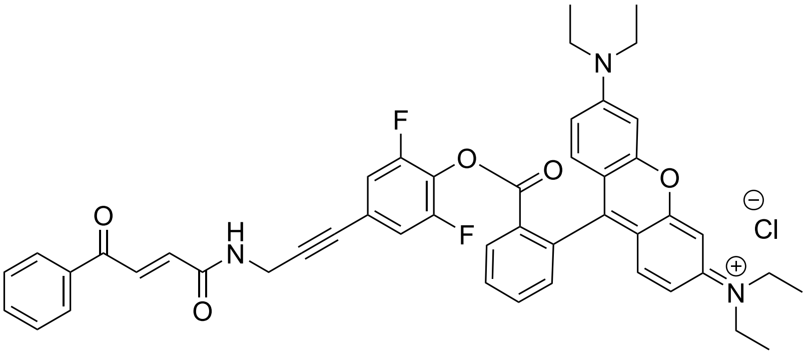

**2a-rho**

$C_{47}H_{42}F_2N_3O_5^+ \cdot Cl^-$

$^1$H-NMR ($CDCl_3$, δ/ppm): see Fig S16; $^{19}$F-NMR ($CDCl_3$, δ/ppm): -126.4. LC-MS$^+$: 766.3 (*exp* 766.3).

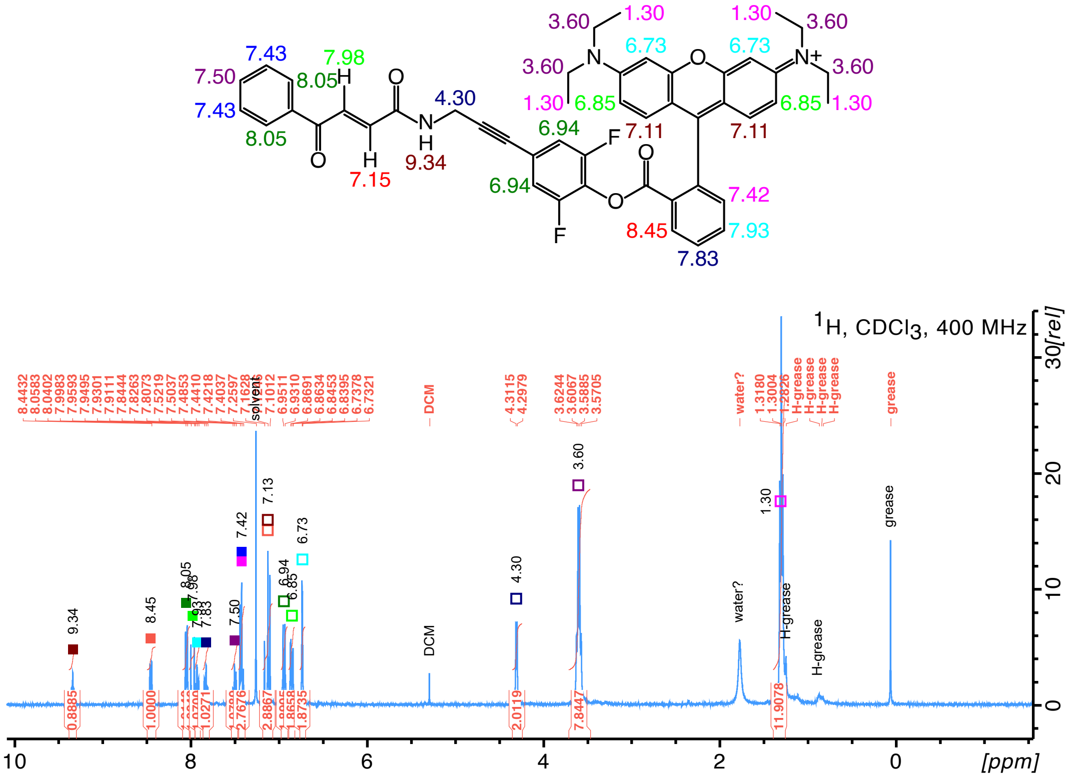

**Figure S16 | 2a-rho NMR spectra**

#### Linker 2b

Reaction scheme: Fig. S12

**4-(*N*-Boc-3-aminopropan-1-yl)-2,6-difluorophenol (2b-3): 287.31 g/mol Alkyne reduction** of 1.63 mmol with $X_1$ = 462 mg **2a-3**; $X_2$ = 46 mg; $X_3$ = 7 h.

Yields 454 mg of **2b-3** (1.58 mmol, 97 %).

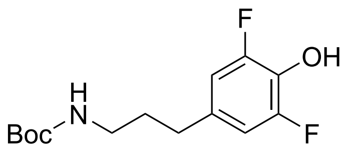

**2b-3**
$C_{14}H_{19}F_2NO_3$

$^1$H-NMR ($CDCl_3$, δ/ppm): 6.70 (2H, d, H3 and H5), 5.65 (1H, s-br, OH), 4.55 (1H, s-br, NH), 3.13 (2H, s, -HN-C$H_2$-), 2.54 (2H, t, Ph-C$H_2$-), 1.75 (2H, quintet, -$CH_2$-C$H_2$-$CH_2$-), 1.44 (9H, s, Boc); $^{19}$F-NMR ($CDCl_3$, δ/ppm): -135.5.

**Linker 2b: 345.35 g/mol** **Boc deprotection** of 1.78 mmol with $X_1$ = 512 mg **2b-3**; $X_2$ = 2 h. Yields 89 %.

Followed by **HATU amide coupling** of 1.2 mmol, $X_1$ = 238 mg trans-3-benzoylacrylic acid (1.35 mmol, 1.125 eq); $X_2$ = 0 °C for 90 min; $X_3$ = 362 mg **2a-4**; $X_4$ = 7 h; $X_5$ = 64:36 PE:EtOAc.

Yields 215 mg of **2b** (0.62 mmol, 52 % over two steps).

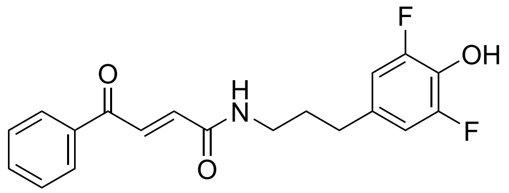

**2b**
$C_{19}H_{17}F_2NO_3$

$^1$H-NMR (MeOD, δ/ppm): 8.03 (2H, d, Ph-*o*), 7.87 (1H, d, $J$ = 15.3, alkene Ph side), 7.66 (1H, t, Ph-*p*), 7.55 (2H, t, Ph-*m*), 7.01 (1H, d, $J$ = 15.3, alkene NH side), 6.78 (2H, d, H3 and H5), 3.32 (2H, s, -HN-C$H_2$-), 2.59 (2H, t, Ph-C$H_2$-), 1.85 (2H, quintet, -$CH_2$-C$H_2$-$CH_2$-), 1.44 (9H, s, Boc); $^{19}$F-NMR (MeOD, δ/ppm): -136.2.

**2b-rho: 806.35 g/mol** **EDC esterification** with $X_1$ = 120 mg Rhodamine B; $X_2$ = 35 mg **2b**; $X_3$ = 92:8 DCM:MeOH.

Yields 44 mg of **2b-rho** (54 µmol, 54 %).

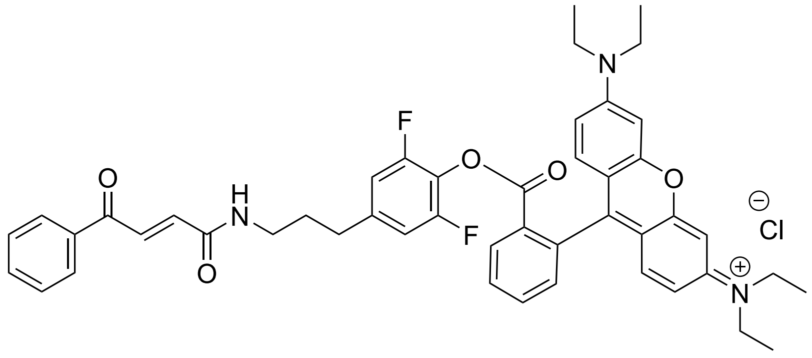

**2b-rho**
$C_{47}H_{46}F_2N_3O_5^+ \cdot Cl^-$

$^1$H-NMR ($CDCl_3$, δ/ppm): see Fig S17; $^{19}$F-NMR ($CDCl_3$, δ/ppm): -127.5. LC-MS$^+$: 770.3 (*exp* 770.3).

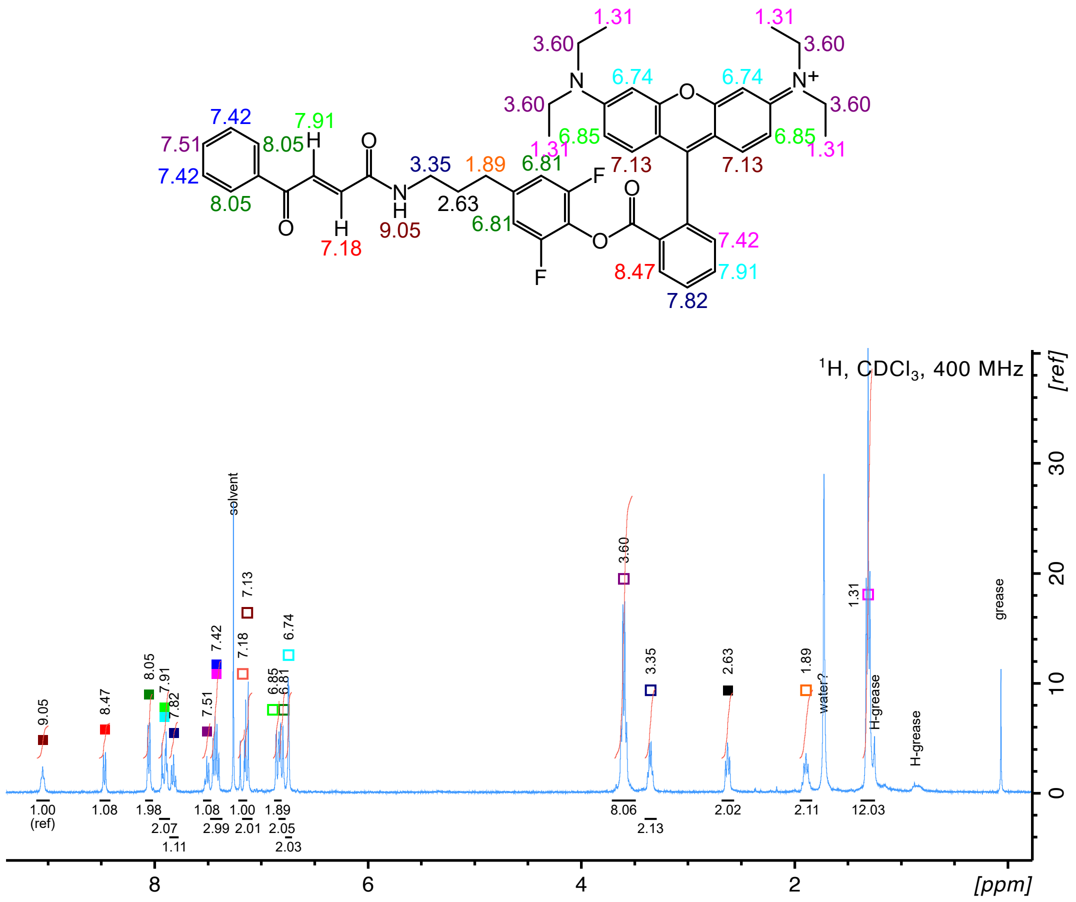

**Figure S17 | 2b-rho NMR spectra**

#### Group 3 linkers

The spacers used in linkers **3c** and **3d** were not commercially available with a terminal alkyne, but published syntheses were available and produced good yields (Fig. S18a). Additionally, the iodination of the fluorophenols had to be changed from the one used for 2-fluorophenol; a modified reaction optimised for 2,3,6-trifluorophenol led to a high yield at very high purity (Fig. S18b, first step) of this previously-unpublished compound.

#### Linker 3c

Reaction scheme: Fig. S18

***N*-Boc-2-(prop-2-yn-1-yloxy)ethan-1-amine (c-2): 199.25 g/mol** Williamson ether synthesis, based on refs [13, 14]. 3.23 g *N*-Boc-ethanolamine (20 mmol) was weighed in a round-bottom flask with 2.98 g of 80 %wt propargyl bromine in toluene stabilised with MgO (2.38 g net weight, 20 mmol, 1 eq), and then dissolved in tetrahydrofuran (20 mL). 476 mg tetra-(*n*-butyl)ammonium iodide (TBAI) (1.2 mmol, 0.06 eq), 498 mg potassium iodide (3 mmol, 0.15 eq) and 1.12 g potassium hydroxide (20 mmol, 1 eq) were added and the reaction mixture was stirred at room temperature O/N. On completion, the mixture was filtered and the filtrate was concentrated *in vacuo*, diluted with $H_2O$ (20 mL) and ex-

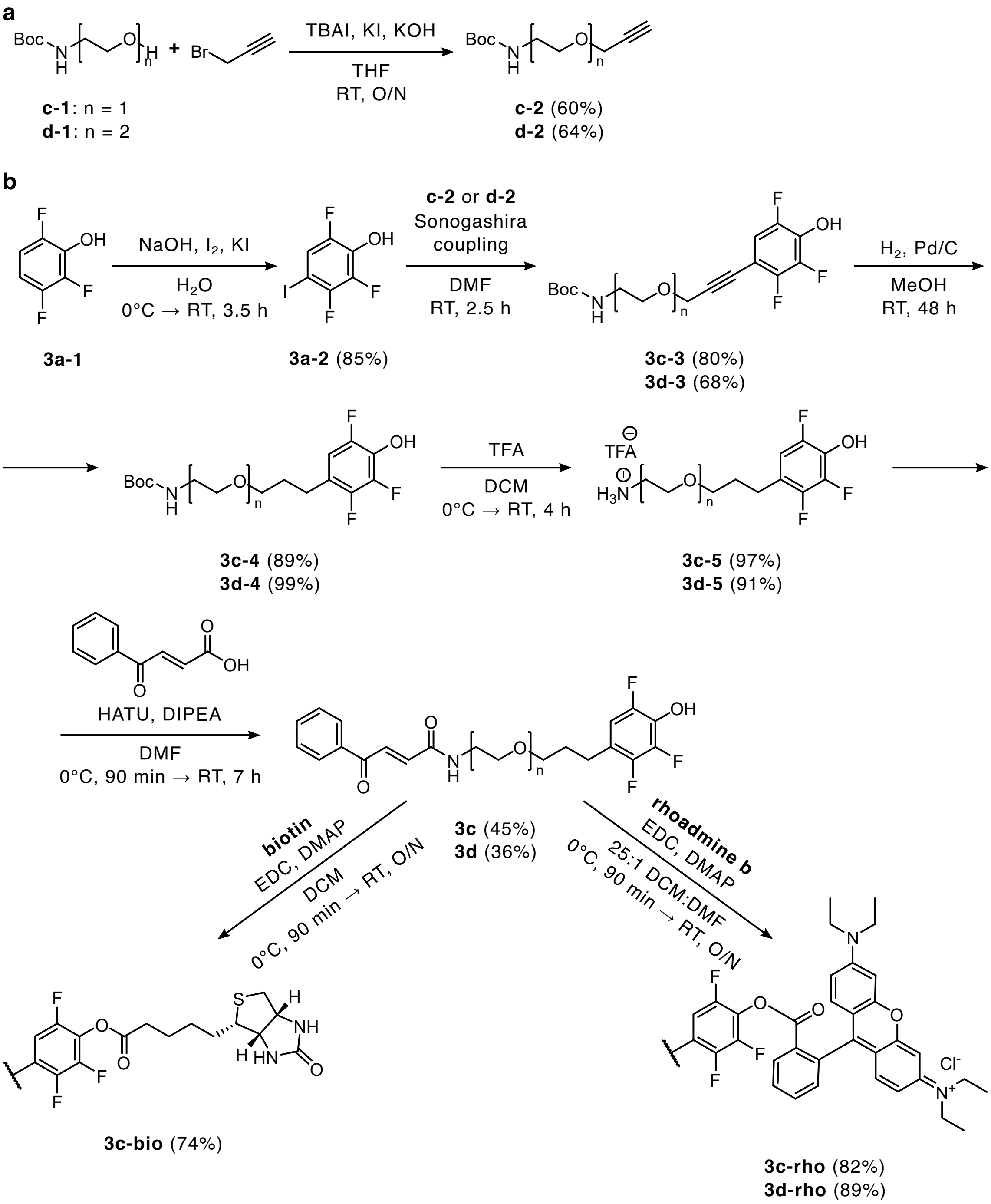

**Figure S18 | Synthetic routes to group 3 linkers.** **(a)** Route to spacer precursors **c-2** and **d-2**. **(b)** Route to linkers **3c** and **3d**, and on to **3c-bio**, **3c-rho** and **3d-rho**. **TBAI** tetrabutylammonium iodide; **THF** tetrahydrofuran.

tracted twice with EtOAc (50 mL). The organic phase was concentrated *in vacuo*, loaded on silica gel and purified by flash chromatography (5:1 PE:EtOAc).

Yields 2.39 g of **c-2** (12 mmol, 60 %) as a colourless oil.

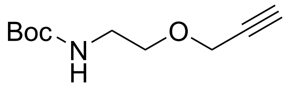

**c-2**
$C_{10}H_{17}NO_3$

$^1$H-NMR ($CDCl_3$, δ/ppm): 4.89 (1H, s-br, NH), 4.15 (2H, s, -C$H_2$-C ), 3.58 (2H, t, -O-C$H_2$-$CH_2$-), 3.34 (2H, q, -O-$CH_2$-C$H_2$-), 2.44 (1H, s, CH), 1.44 (9H, s, Boc).

**2,3,6-trifluoro-4-iodophenol (3a-2): 273.98 g/mol** 540 mg of sodium hydroxide (13.5 mmol, 2 eq) was dissolved in $H_2O$ (17 mL for final 800 mM) and cooled to 4 °C. To a stirred solution of 1.71 g iodine (6.75 mmol, 1 eq) and 1.12 g potassium iodide (6.75 mmol, 1 eq) in $H_2O$ (17 mL) at 0 °C was added dropwise 1 g 2,3,6-trifluorophenol (6.75 mmol) and stirred for 10 min, after which the cold sodium hydroxide solution was added slowly and the flask was brought to room temperature and stirred for 3.5 hours. The reaction was quenched with saturated ammonium chloride and 1.34 g (8.45 mmol, 1.25 eq) sodium thiosulphate was added to reduce any remaining iodine. The reaction mixture was then carefully acidified with HCl, extracted with DCM (3 × 40 mL) then washed with $H_2O$ (2 × 100 mL) and 100 mL brine. The organic layer was dried over $MgSO_4$, filtered and concentrated *in vacuo.*

Yields 1.56 g of **3a-2** (5.73 mmol, 85 %) at very high purity with no need for further purification.

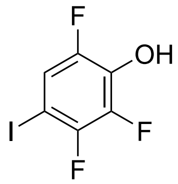

**3a-2**
$C_6H_2F_3IO$

$^1$H-NMR ($CDCl_3$, δ/ppm): 7.28 (1H, ddd, H5), 5.54 (1H, s-br, OH); $^{13}$C-NMR ($CDCl_3$, δ/ppm): 149.3, 146.9, 139.1, 134.8, 119.2, 67.6; $^{19}$F-NMR ($CDCl_3$, δ/ppm): -119.8 (1F, dd, $J = 10.6, 21.8$, F3), -138.9 (1F, dd, $J = 6.8, 10.6$, F6), -153.3 (1F, dd, $J = 6.8, 21.8$, F2). The spectra were analysed and compared to the starting material spectra to ascertain the correct position of the iodine on the ring – see Fig. S19.

**4-(3-(2-(Boc-amino)ethoxy)prop-1-yn-1-yl)-2,3,6-trifluorophenol (3c-3): 345.32 g/mol** **Sonogashira coupling** with $X_1$ = 548 mg **3a-2**; $X_2$ = 598 mg **c-2**; $X_3$ = 3 h. The reaction mixture was diluted into 100 mL saturated ammonium chloride (1:20) and immediately extracted with EtOAc (3 × 30 mL). The joined organic fractions were washed with water (2 × 100 mL) and brine (100 mL), dried over $MgSO_4$, filtered and concentrated *in vacuo.* The residue was purified by flash chromatography on silica gel (74:25:1 PE:EtOAc:AcOH).

Yields 610 mg of **3c-3** at 90 % purity (549 mg pure product, 1.59 mmol, 80 %) as a golden-brown viscous liquid, used without further purification.

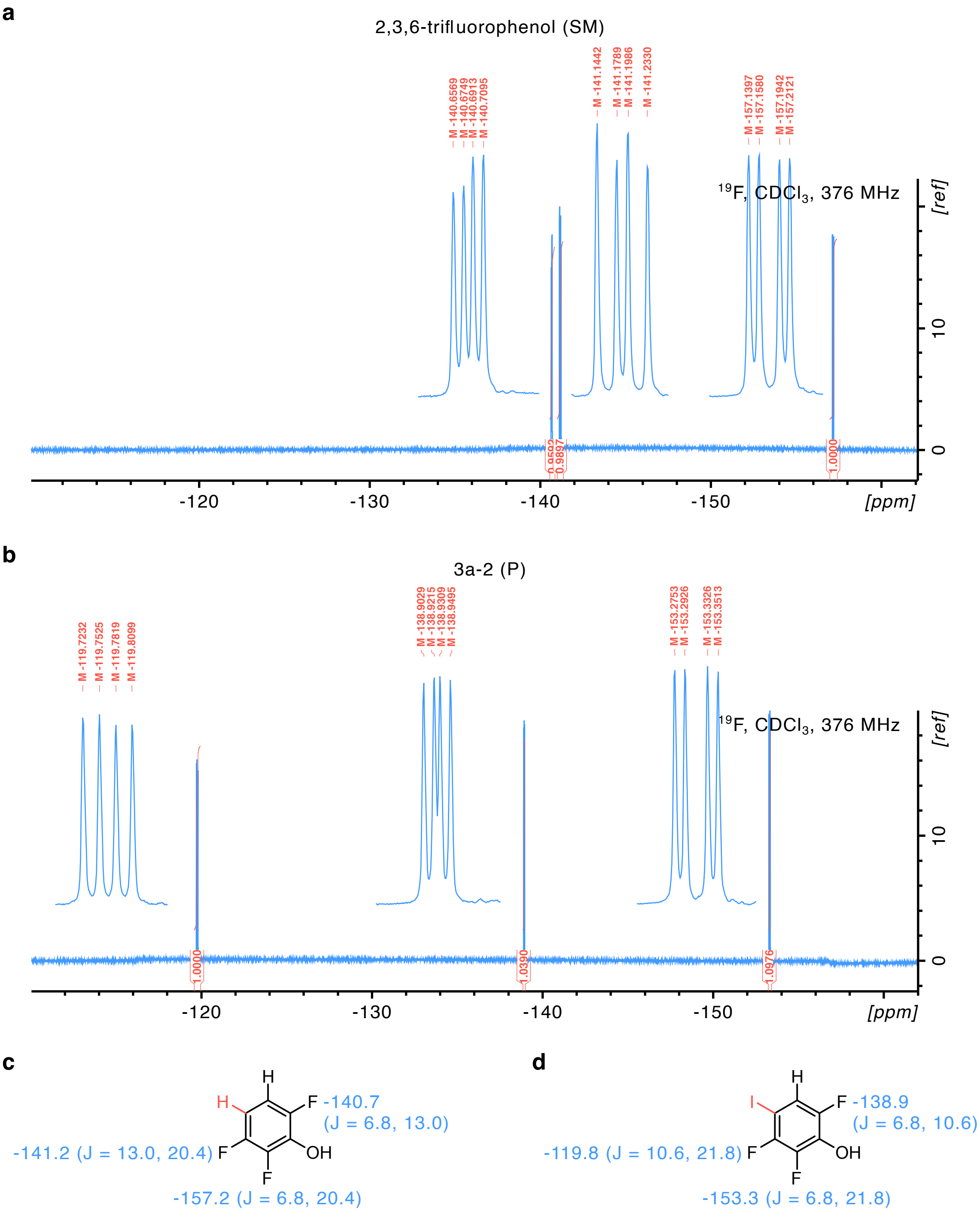

**Figure S19 | Confirmation of the desired structure of the product 3a-2.** $^{19}$F-NMR spectra of 2,3,6-trifluorophenol (panels a,c) and product **3a-2** (panels b,d) identify fluorine 3 as the one experiencing the most significant shift on addition of the fluorine atom (+21.4 ppm), consistent with the predicted +19.9 ppm shift from an orthogonal iodine on a fluoroaryl compound [15]. The J-coupling pattern supports the identification of the new -119.8 ppm peak as F3, located at -141.2 ppm in the SM (panels c,d). These observations place the iodine atom at position 4, confirming the correct structure of compound **3a-2** (2,3,6-trifluoro-4-iodophenol).

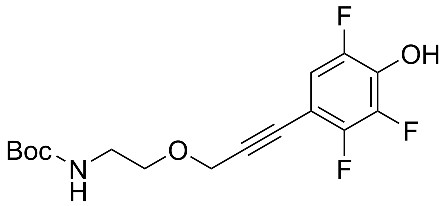

**3c-3**
$C_{16}H_{18}F_3NO_4$

[1]H-NMR ($CDCl_3$, δ/ppm): 6.91 (1H, ddd, H5), 4.99 (1H, s-br, NH), 4.38 (2H, s, -O-C$H_2$-C ), 3.65 (2H, t, -O-C$H_2$-$CH_2$-), 3.37 (2H, q, -O-$CH_2$-C$H_2$-), 1.44 (9H, s, Boc); [19]F-NMR ($CDCl_3$, δ/ppm): -138.6 (1F, dd, F6), -140.4 (1F, dd, F3), -156.9 (1F, dd, F2).

**4-(3-(2-(Boc-amino)ethoxy)propyl)-2,3,6-trifluorophenol (3c-4): 349.35 g/mol**
**Alkyne reduction** of 1.2 mmol with $X_1$ = 462 mg **3c-3** at 90 %wt (416 mg pure compound); $X_2$ = 47 mg; $X_3$ = 42 h.

Yields 375 mg of **3c-4** (1.07 mmol, 89 %).

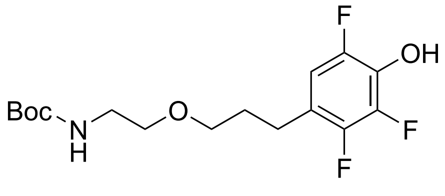

**3c-4**
$C_{16}H_{22}F_3NO_4$

[1]H-NMR ($CDCl_3$, δ/ppm): 6.68 (1H, ddd, H5), 4.91 (1H, s-br, NH), 3.47 (2H, t, -O-C$H_2$-$CH_2$-NH-), 3.43 (2H, t, -O-C$H_2$-$CH_2$-$CH_2$-), 3.31 (2H, dt, -O-$CH_2$-C$H_2$-NH-), 2.64 (2H, t, Ar-C$H_2$-), 1.83 (2H, quintet, Ar-$CH_2$-C$H_2$-), 1.45 (9H, s, Boc); [13]C-NMR ($CDCl_3$, δ/ppm): 156.4, 148.9, 147.3, 144.9, 132.6, 120.2, 110.5, 79.8, 69.97, 69.93, 40.6, 29.8, 28.5, 25.0; [19]F-NMR ($CDCl_3$, δ/ppm): -141.6 (1F, dd, F3), -147.5 (1F, dd, F6), -158.0 (1F, dd, F2).

**2-(3-(2,3,5-trifluoro-4-hydroxyphenyl)propoxy)ethan-1-aminium trifluoroacetate (3c-5): 363.26** **Boc deprotection** with $X_1$ = 349 mg **3c-4**; $X_2$ = 5 h.

Yields 352 mg of **3c-5** (0.97 mmol, 97 %) as a flaky white solid.

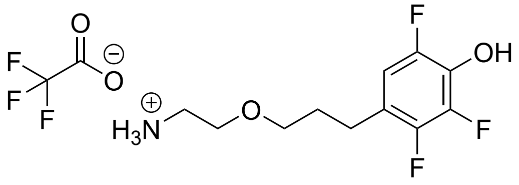

**3c-5**
$C_{11}H_{15}F_3NO_2^+ \cdot C_2F_3O_2^-$

[1]H-NMR (MeOD, δ/ppm): 6.79 (1H, ddd, H5), 3.64 (2H, t, -O-C$H_2$-$CH_2$-NH-), 3.52 (2H, t, -O-C$H_2$-$CH_2$-$CH_2$-), 3.12 (2H, dt, -O-$CH_2$-C$H_2$-NH-), 2.69 (2H, t, Ar-C$H_2$-), 1.88 (2H, quintet, Ar-$CH_2$-C$H_2$-); [19]F-NMR (MeOD, δ/ppm): -77.9 (3F, s, TFA), -142.0 (1F, dd, F3), -150.7 (1F, dd, F6), -160.4 (1F, dd, F2).

**Linker 3c: 407.39 g/mol** **HATU amide coupling** of 0.9 mmol with $X_1$ = 176 mg trans-3-benzoylacrylic acid (1.0 mmol, 1.1 eq); $X_2$ = RT for 1.5 h; $X_3$ = 327 mg **3c-5**; $X_4$ = O/N; $X_5$ = 100 % EtOAc ($R_f$ = 0.74).

Yields 183 mg of **3c** (0.45 mmol, 45 %).

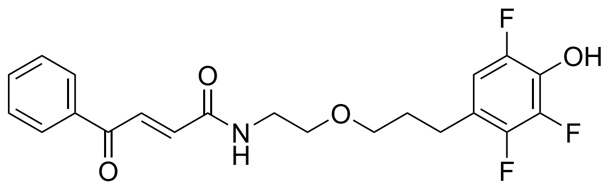

**3c**
$C_{21}H_{20}F_3NO_4$

$^{1}$H-NMR (DMSO-d$_6$, δ/ppm): 8.00 (2H, d, Ph-*o*), 7.76 (1H, d, *J* = 15.3, alkene Ph side), 7.70 (1H, t, Ph-*p*), 7.57 (2H, t, Ph-*m*), 7.04 (1H, d, *J* = 15.3, alkene NH side), 6.96 (1H, ddd, H5), 4.04 (1H, s-br, NH), 3.46 (2H, t, -O-C$H_2$-CH$_2$-NH-), 3.40 (2H, t, -O-C$H_2$-CH$_2$-CH$_2$-), 3.37 (2H, t, -O-CH$_2$-C$H_2$-NH-), 2.60 (2H, t, Ar-C$H_2$-), 1.76 (2H, quintet, Ar-CH$_2$-C$H_2$-); $^{13}$C-NMR (DMSO-d$_6$, δ/ppm): 189.9, 163.5, 146.8, 144.4, 142.8, 136.7, 136.4, 133.7, 132.9, 132.0, 129.0, 128.6, 119.0, 110.8, 69.1, 68.5, 39.0, 29.4, 24.1; $^{19}$F-NMR (DMSO-d$_6$, δ/ppm): -137.5 (1F, dd, F3), -147.4 (1F, dd, F6), -156.2 (1F, dd, F2). Full spectra and assignments appear in Fig. S20.

**3c-rho: 868.39 g/mol** Modified **EDC esterification**: 96 mg rhodamine B (200 µmol, 2 eq), 46 mg *N*-(3-Dimethylaminopropyl)-*N* -ethylcarbodiimide hydrochloride (EDC·HCl) (240 µmol, 2.4 eq) and 4.9 mg 4-dimethylaminopyridine (40 µmol, 0.4 eq) were weighed in an oven-dried round-bottom flask and dissolved in dry DCM (1.5 mL) at 0 °C. In a separate vial, 41 mg **3c** (100 µmol) was dissolved in a mixture of dry DMF (100 µL) and dry DCM (1 mL), then added dropwise over the course of 45 min to the stirred reaction mixture. The reaction was brought to room temperature and stirred overnight. The reaction mixture was then diluted 20-fold in DCM, washed once with $H_2O$ then separated; aqueous phase was extracted with DCM, the two organic phases were joined, washed twice with 100 mL $H_2O$ and once with 100 mL brine, dried over $MgSO_4$, filtered and dried *in vacuo*. The residue was purified by flash chromatography on silica gel.

Yields 71 mg of **3c-rho** (82 µmol, 82 %).

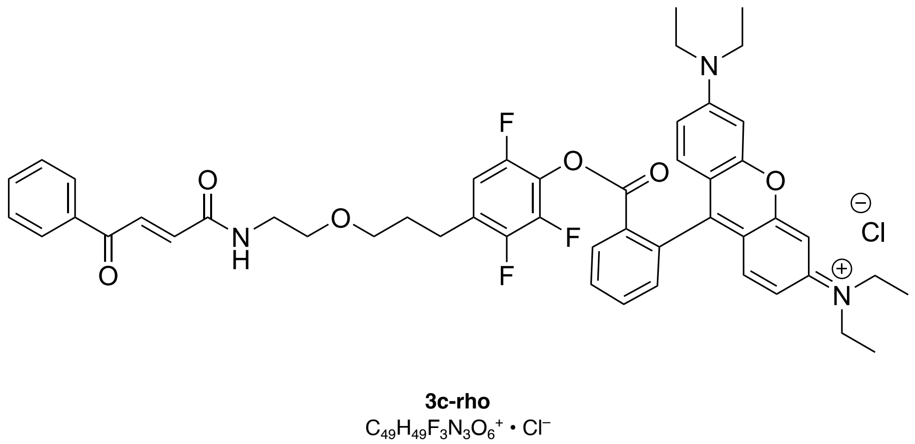

**3c-rho**
$C_{49}H_{49}F_3N_3O_6^+ \cdot Cl^-$

$^{1}$H-NMR (CDCl$_3$, δ/ppm): see Fig S21; $^{13}$C-NMR (CDCl$_3$, δ/ppm): 245.0, 190.4, 164.7, 157.9, 157.1, 155.7, 137.1, 136.1, 134.6, 133.6, 133.0, 132.2, 131.1, 131.0, 130.9, 129.1, 128.9, 127.9, 114.5, 113.6, 96.6, 77.4, 69.7, 69.1, 46.3, 39.7, 29.8, 29.5, 25.6, 12.8; $^{19}$F-NMR (CDCl$_3$, δ/ppm): -132.8 (1F, d, F6), -145.8 (1F, dd, F3), -149.0 (1F, d, F2).

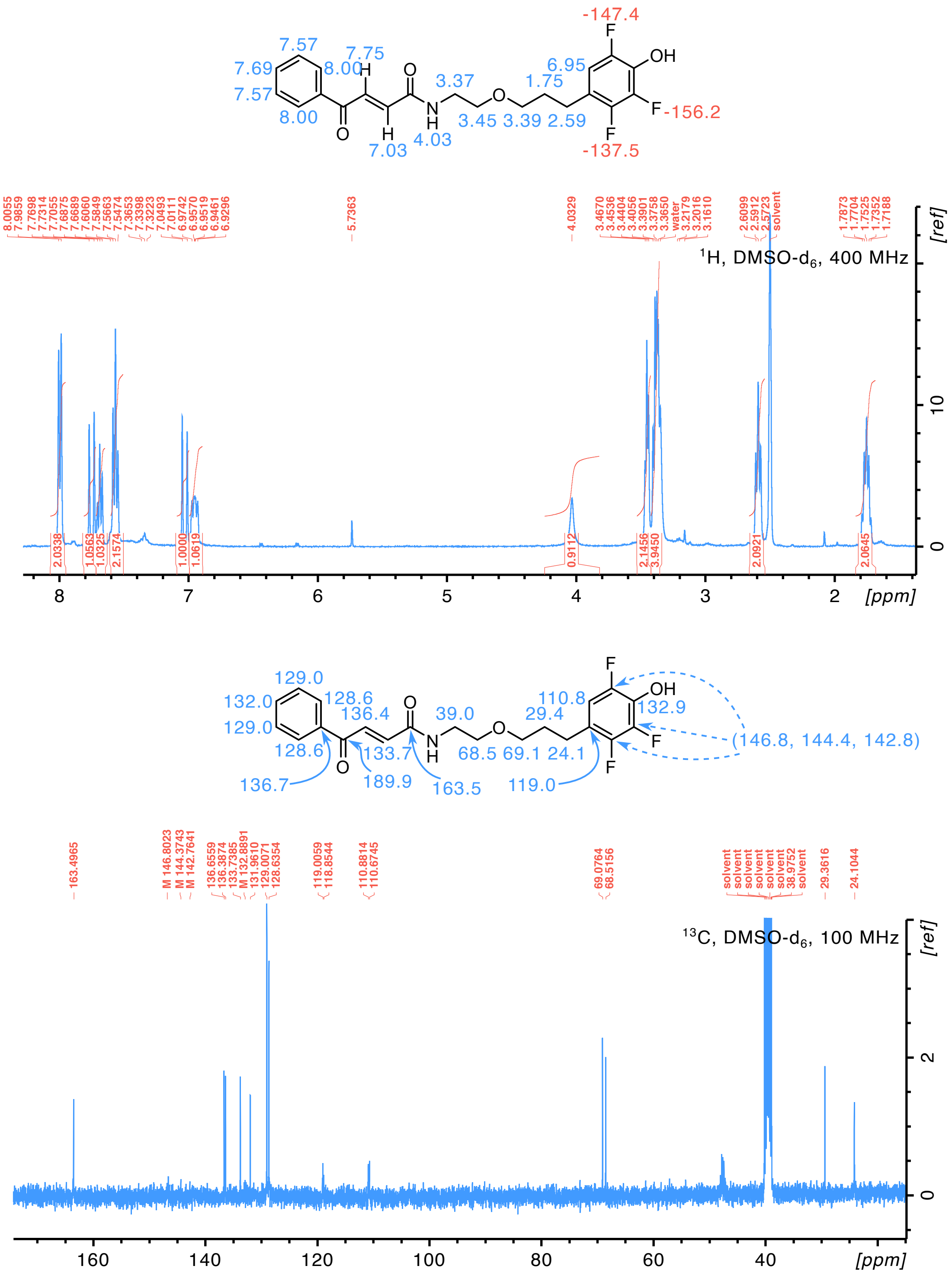

**Figure S20 | Linker 3c NMR spectra**

**3c-bio: 633.68 g/mol** Modified **EDC esterification**: 49 mg biotin (200 µmol, 2 eq), 46 mg *N*-(3-Dimethylaminopropyl)-*N* -ethylcarbodiimide hydrochloride (EDC·HCl) (240 µmol, 2.4 eq) and 4.9 mg 4-dimethylaminopyridine (40 µmol, 0.4 eq) were weighed in an oven-dried round-bottom flask and dissolved in dry DCM (1.5 mL) at 0 °C. In a separate vial, 41 mg **3c** (100 µmol) was dissolved in dry DCM (1 mL), then added dropwise over

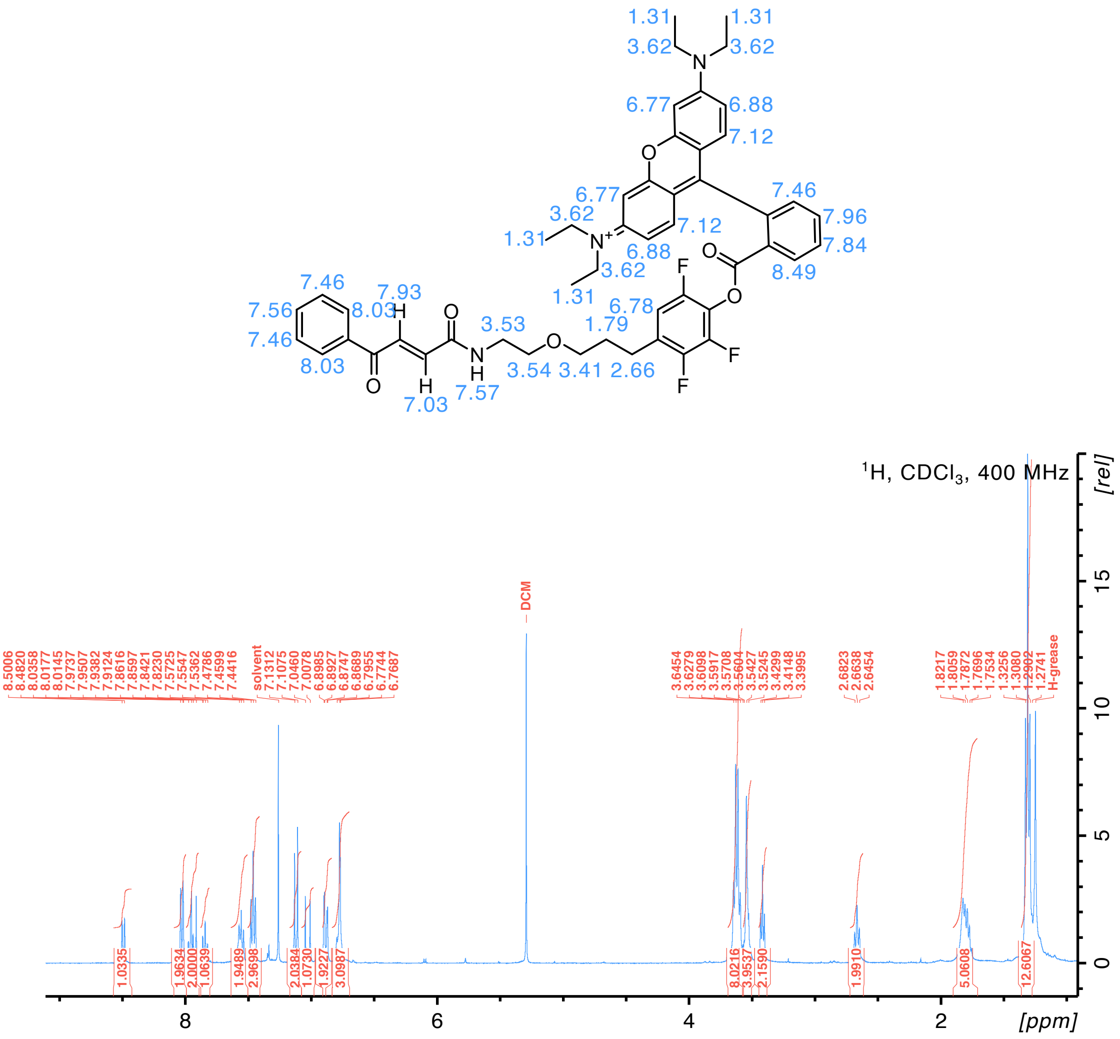

**Figure S21 | 3c-rho NMR spectra**

the course of 45 min to the stirred reaction mixture. The reaction was brought to room temperature and stirred overnight. The reaction mixture was then decanted into a 15 mL centrifuge tube using DCM and a precipitate was pelleted by centrifugation (3 min / 4000 × g) and the supernatant was purified by flash chromatography on silica gel (gradient 95:5 → 92:8 DCM:MeOH).

Yields 47 mg of **3c-bio** (74 µmol, 74 %).

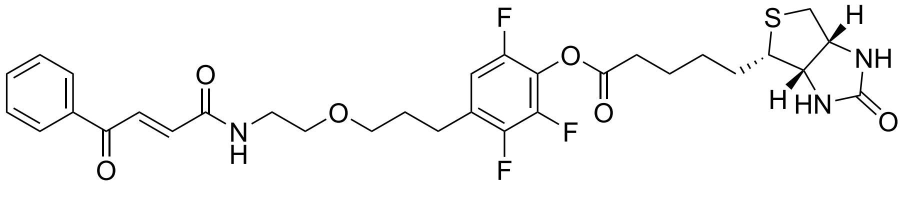

**3c-bio**
$C_{31}H_{34}F_3N_3O_6S$

$^{1}$H-NMR ($CDCl_3$, δ/ppm): see Fig S22; $^{13}$C-NMR ($CDCl_3$, δ/ppm): 190.0, 170.2, 164.3, 163.4, 137.0, 135.4, 133.9, 133.3, 129.0, 129.0, 111.3, 111.1, 77.4, 70.0, 69.3, 62.0, 60.2, 55.4, 40.7, 39.9, 33.2, 29.6, 28.4, 28.3, 25.6, 24.8; $^{19}$F-NMR ($CDCl_3$, δ/ppm): -132.8

(1F, d, F6), -146.2 (1F, dd, F3), -148.8 (1F, d, F2).

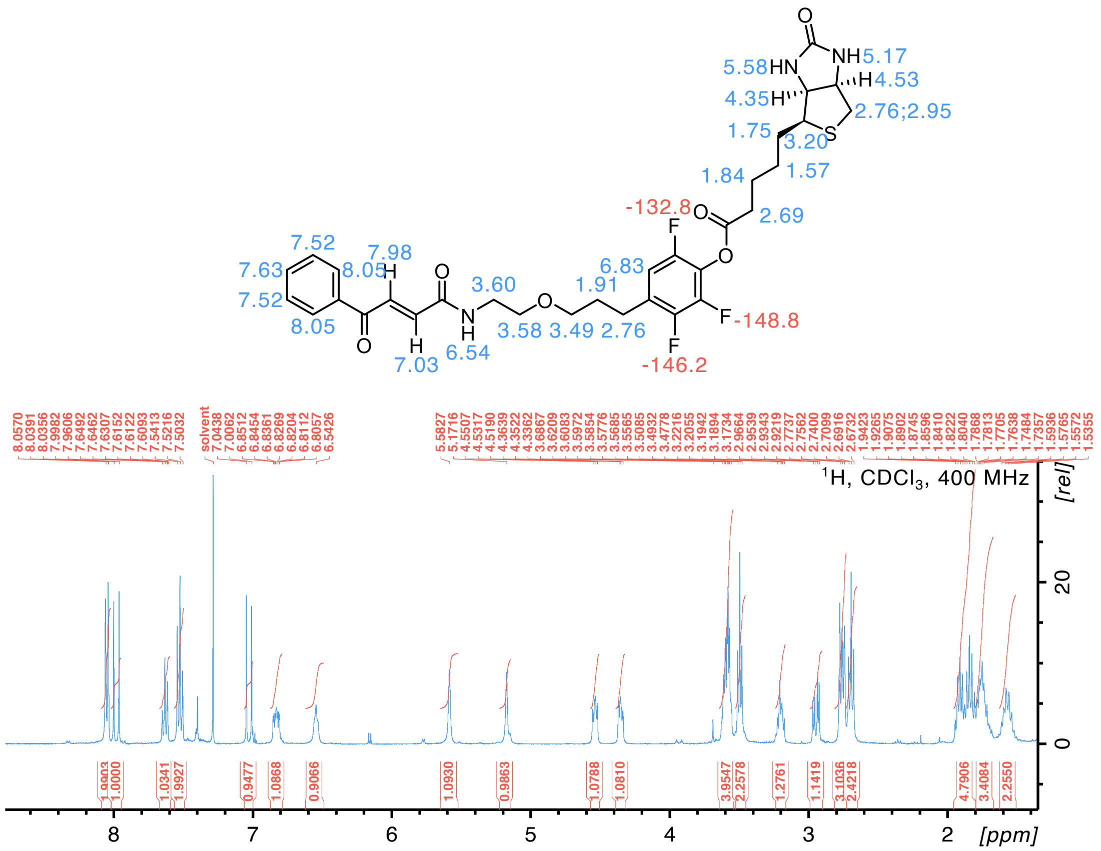

**Figure S22 | 3c-bio NMR spectra**

##### Linker 3d

Reaction scheme: Fig. S18

***N*-Boc-2-(2-(prop-2-yn-1-yloxy)ethoxy)ethan-1-amine (d-2): 243.30 g/mol** Williamson ether synthesis, based on refs [13, 14]. 4.11 g 2-[2-(Boc-amino)ethoxy]ethanol (20 mmol) was weighed in a round-bottom flask with 2.98 g of 80 %wt propargyl bromine in toluene stabilised with MgO (2.38 g net weight, 20 mmol, 1 eq), and then dissolved in tetrahydrofuran (20 mL). 476 mg tetra-(*n*-butyl)ammonium iodide (TBAI) (1.2 mmol, 0.06 eq), 498 mg potassium iodide (3 mmol, 0.15 eq) and 1.12 g potassium hydroxide (20 mmol, 1 eq) were added and the reaction mixture was stirred at room temperature overnight. On completion, the mixture was filtered and the filtrate was concentrated *in vacuo*, diluted with $H_2O$ (20 mL) and extracted twice with EtOAc (50 mL). The organic phase was concentrated *in vacuo*, loaded on silica gel and purified by flash chromatography (5:1 PE:EtOAc).

Yields 3.1 g of **d-2** (12.7 mmol, 64 %) as a colourless oil.

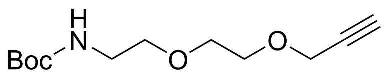

**d-2**
$C_{12}H_{21}NO_4$

$^1$H-NMR ($CDCl_3$, δ/ppm): 4.94 (1H, s-br, NH), 4.20 (2H, s, -C$H_2$C ), 3.69 (2H, t, -O-C$H_2$-$CH_2$-O-), 3.64 (2H, t, -O-$CH_2$-C$H_2$-NH-), 3.54 (2H, t, -O-C$H_2$-$CH_2$-NH-), 3.31 (2H, t, -O-$CH_2$-C$H_2$-O-), 2.44 (1H, s,  CH), 1.44 (9H, s, Boc).

**4-(3-(2-(2-(Boc-amino)ethoxy)ethoxy)prop-1-yn-1-yl)-2,3,6-trifluorophenol (3d-3): 389.37 Sonogashira coupling** with $X_1$ = 548 mg **3a-2**; $X_2$ = 730 mg **d-2**; $X_3$ = 3 h. The reaction mixture was diluted into 100 mL saturated ammonium chloride (1:20) and immediately extracted with EtOAc (3 × 30 mL). The joined organic fractions were washed with water (2 × 100 mL) and brine (100 mL), dried over $MgSO_4$, filtered and concentrated *in vacuo*. The residue was purified by flash chromatography on silica gel (39:1 → 29:1 DCM:MeOH).

Yields 702 mg at 75 %wt purity of **3d-3** (529 mg pure product, 1.36 mmol, 68 %), used without further purification.

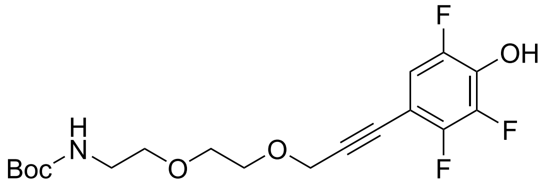

**3d-3**
$C_{18}H_{22}F_3NO_5$

$^1$H-NMR ($CDCl_3$, δ/ppm): 6.89 (1H, ddd, H5), 5.05 (1H, s-br, NH), 4.42 (2H, s, -O-C$H_2$-C ), 3.75 (2H, t, -O-$CH_2$-C$H_2$-O-), 3.67 (2H, t, -O-C$H_2$-$CH_2$-O-), 3.56 (2H, t, -O-C$H_2$-$CH_2$-NH-), 3.33 (2H, t, -O-$CH_2$-C$H_2$-NH-), 1.43 (9H, s, Boc); $^{19}$F-NMR ($CDCl_3$, δ/ppm): -138.8 (1F, dd, F6), -140.3 (1F, dd, F3), -156.9 (1F, dd, F2).

**4-(2-(3-(2-(Boc-amino)ethoxy)propoxy)ethyl)-2,3,6-trifluorophenol (3d-4): 393.40 g/mol Alkyne reduction** of 1.2 mmol with $X_1$ = 622 mg **3d-3** at 75 %wt (467 mg pure compound); $X_2$ = 50 mg; $X_3$ = 42 h. Full conversion by NMR.

Yields 470 mg of **3d-4** (1.19 mmol, 99 %).

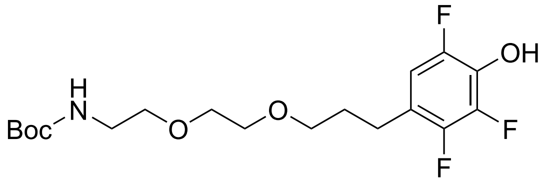

**3d-4**
$C_{18}H_{26}F_3NO_5$

**(3-(3-(2,3,5-trifluoro-4-hydroxyphenyl)propoxy)propoxy)methanaminium trifluoroacetate (3d-5): 407.31 g/mol Boc deprotection** with $X_1$ = 393 mg **3d-4**; $X_2$ = 5 h. Following toluene co-evaporation, the residue was re-dissolved in DCM and re-dried, several times.

Yields 403 mg of **3d-5** at 92 %wt (370 mg pure product, 0.91 mmol, 91 %) as a colourless oil, used without further purification.

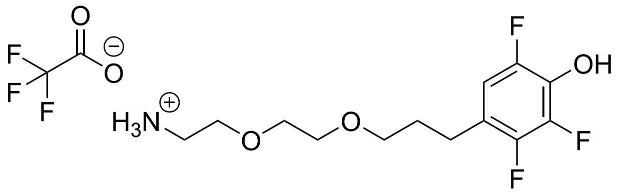

**3d-5**
$C_{13}H_{19}F_3NO_3^+ \cdot C_2F_3O_2^-$

$^1$H-NMR (MeOD, δ/ppm): 6.79 (1H, ddd, H5), 3.70 (2H, t, -O-CH$_2$-C$H_2$-O-), 3.67 (2H, t, -O-C$H_2$-CH$_2$-NH-), 3.62 (2H, t, -O-C$H_2$-CH$_2$-O-), 3.50 (2H, t, -O-CH$_2$-C$H_2$-NH-), 3.13 (2H, t, -O-C$H_2$-CH$_2$-CH$_2$-), 2.66 (2H, t, Ar-C$H_2$-), 1.85 (2H, quintet, Ar-CH$_2$-C$H_2$-); $^{19}$F-NMR (MeOD, δ/ppm): -77.8 (3F, s, TFA), -142.0 (1F, dd, F3), -150.8 (1F, dd, F6), -160.4 (1F, dd, F2).

**Linker 3d: 451.44 g/mol** **HATU amide coupling** of 0.9 mmol with $X_1$ = 176 mg trans-3-benzoylacrylic acid (1.0 mmol, 1.1 eq); $X_2$ = RT for >1 hour; $X_3$ = 400 mg **3d-5** at 91 %wt (0.9 mmol); $X_4$ = O/N; $X_5$ = 100 % EtOAc.

Yields 161 mg of **3d** (0.36 mmol, 36 %).

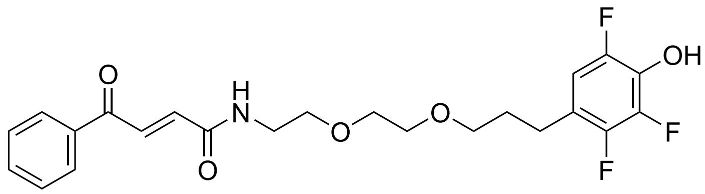

**3d**
$C_{23}H_{24}F_3NO_5$

$^1$H-NMR (MeOD, δ/ppm): 8.00 (2H, d, Ph-*o*), 7.86 (1H, d, $J$ = 15.3, alkene Ph side), 7.65 (1H, t, Ph-*p*), 7.53 (2H, t, Ph-*m*), 7.03 (1H, d, $J$ = 15.3, alkene NH side), 6.76 (1H, ddd, H5), 3.637 (2H, t, -O-CH$_2$-C$H_2$-O-), 3.625 (2H, t, -O-C$H_2$-CH$_2$-NH-), 3.60 (2H, t, -O-C$H_2$-CH$_2$-O-), 3.52 (2H, t, -O-CH$_2$-C$H_2$-NH-), 3.47 (2H, t, -O-C$H_2$-CH$_2$-CH$_2$-), 2.63 (2H, t, Ar-C$H_2$-), 1.81 (2H, quintet, Ar-CH$_2$-C$H_2$-); $^{13}$C-NMR (MeOD, δ/ppm): 191.5, 166.7, 150.7, 148.4, 146.2, 138.2, 136.5, 134.9, 134.6, 134.0, 130.0, 129.8, 120.4, 111.6, 71.3, 71.2, 71.1, 70.3, 40.8, 31.0, 25.7; $^{19}$F-NMR (DMSO-d$_6$, δ/ppm): -137.6 (1F, dd, F3), -147.6 (1F, dd, F6), -156.3 (1F, dd, F2). $^{19}$F-NMR (MeOD, δ/ppm): -141.0 (1F, dd, F3), -149.7 (1F, dd, F6), -159.4 (1F, dd, F2). Full spectra and assignments appear in Fig. S23.

**3d-rho: 912.44 g/mol** Modified **EDC esterification**: See **3c-rho**.

Yields 81 mg of **3d-rho** (89 µmol, 89 %).

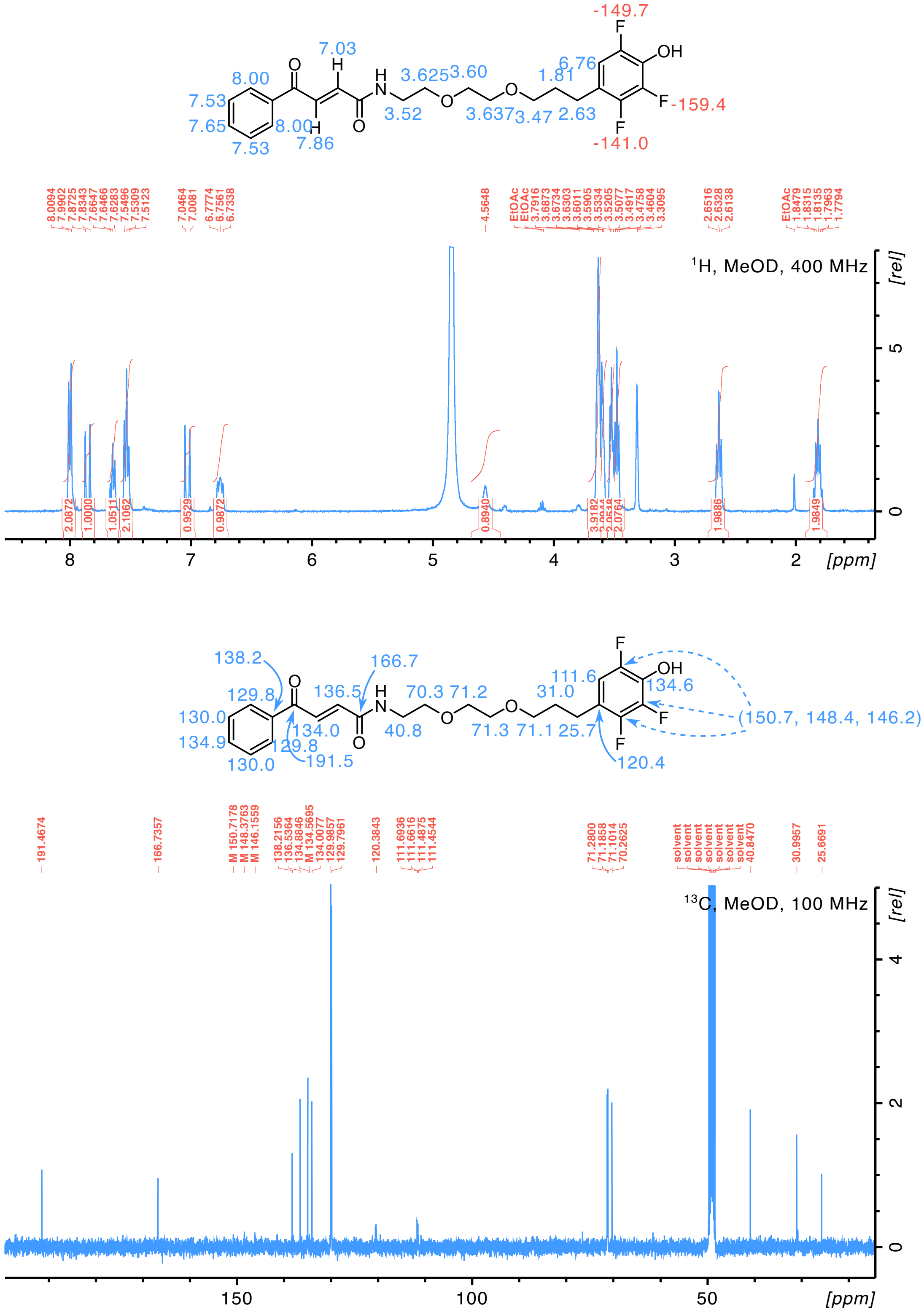

**Figure S23 | Linker 3d NMR spectra**

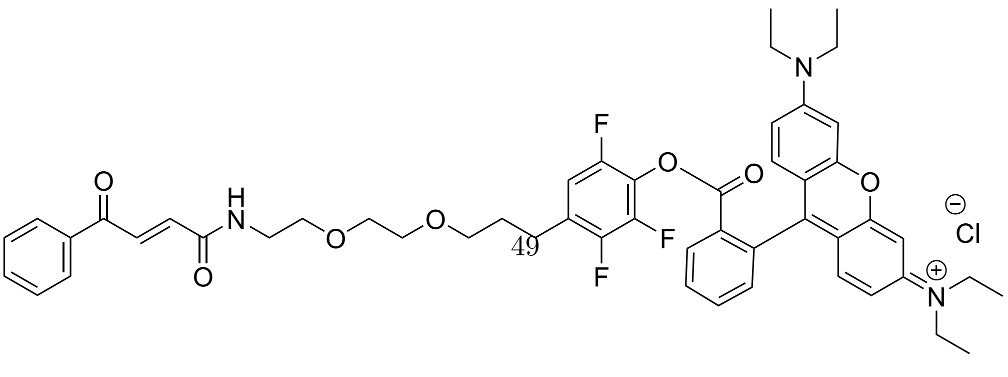

3d-rho

$^{1}$H-NMR ($CDCl_3$, δ/ppm): see Fig S24; $^{13}$C-NMR ($CDCl_3$, δ/ppm): 190.3, 162.0, 157.9, 157.1, 155.7, 137.1, 136.0, 134.6, 133.6, 133.0, 132.2, 131.1, 131.0, 130.9, 129.1, 128.9, 127.8, 114.5, 113.5, 96.6, 70.4, 70.2, 69.8, 69.4, 46.3, 39.6, 29.8, 29.4, 25.4, 12.8; $^{19}$F-NMR ($CDCl_3$, δ/ppm): -132.8 (1F, d, F6), -145.8 (1F, dd, F3), -148.9 (1F, d, F2).

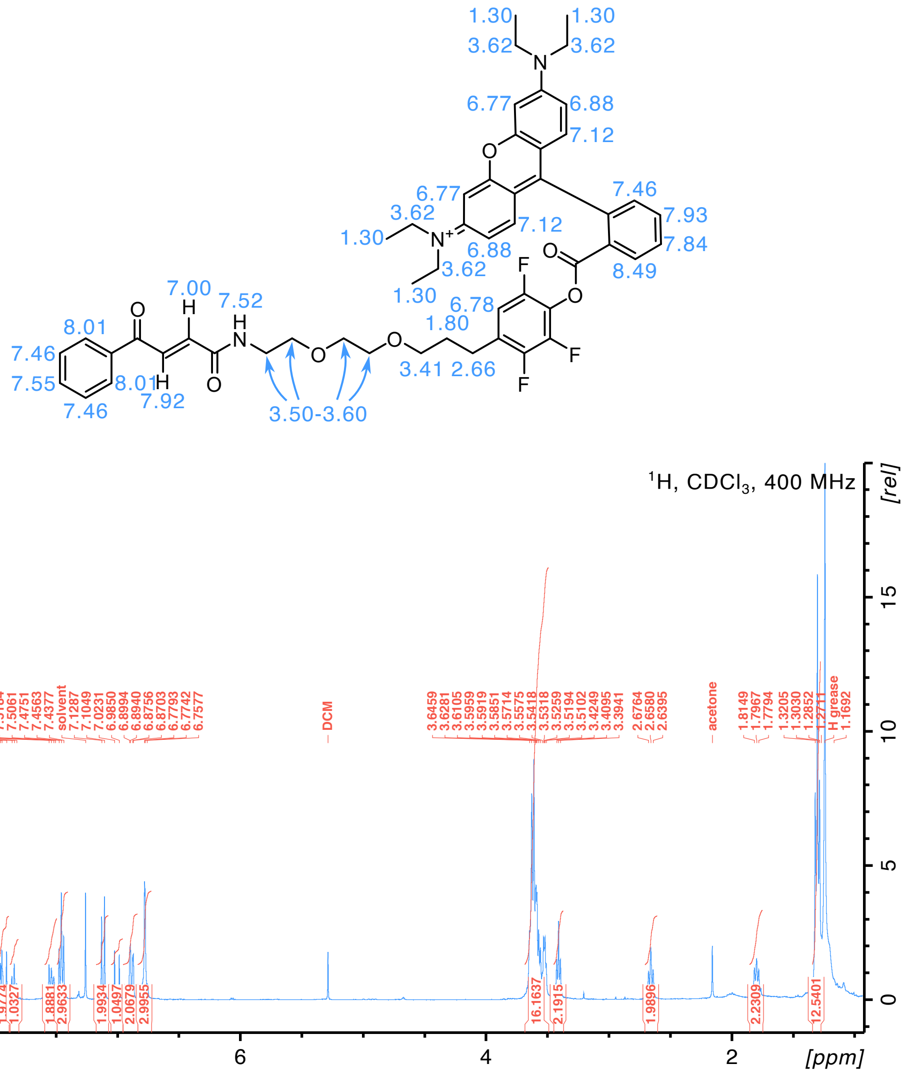

**Figure S24 | 3d-rho NMR spectra**

#### Analytical techniques

##### SDS-PAGE and in-gel fluorescence

Precast Novex NuPAGE Bis-Tris 4–12% or 12–20% gels (Thermo) were used for all SDS-PAGE analyses, running at 200 V for 30–40 min with NuPAGE MES SDS buffer (Thermo). SeeBlue Plus2 (Thermo) pre-stained protein marker was used as standard. Gels were imaged using a NuGenius gel imager (Syngene) either before staining using the UV fluorescence mode, or after staining with InstantBlue (Abcam) using a white light converter.

##### Western blot for detection of biotinylated proteins

An unstained SDS-PAGE gel was transferred to a nitrocellulose membrane by dry-blotting using an iBlot™ 2 device (ThermoFisher), applying 20 V for 1 min, 23 V for 4 min and finally 25 V for 2 min (GFP experiments) or 20 V for 1 min, 23 V for 3.5 min and 25 V for 1 min (β2m experiments). The membrane was blocked using 5 % BSA in TBS (50 mM Tris + 150 mM NaCl, pH 7.4), in a 50 mL tube, on a tube roller at 6 °C overnight. The membrane was washed twice with TBST (TBS + 0.1 % Tween-20 (w/v)) and twice with TBS, before being incubated with 5 % BSA + 0.5  g/mL Streptavidin–Alexa Fluor™ 647 conjugate (Invitrogen) (1:4000 dilution) in TBS for 60 min. Then, the membrane was washed twice with TBST and twice with TBS, and imaged using a ChemiDoc™ MP Imager (Bio-Rad). Band and lane fluorescence intensity were quantified using Image Lab (Bio-Rad), and the total intensities of untreated control lanes were subtracted from each respective treated sample intensity to calculate the signal of non-specific biotinylation by PEABS.

##### UV/vis spectrophotometry

Samples were transferred to a UV/vis-transparent cuvette and spectra were recorded between 220–340 nm on a Cary 400 spectrophotometer. Samples of low volume (e.g. plasmid DNA; or proteins prepared for PEABS generation by reduction and precipitation of aggregates) were measured by depositing a 2 µL sample on a NanoDrop 2000 spectrophotometer (Thermo). In both techniques, the buffer was measured and subtracted as a blank.

For determining protein concentration, absorbance at 280 nm was used according to Beer-Lambert law ($c = \frac{A}{\varepsilon l}$, where $l = 1$ cm for the cuvette used and $\varepsilon$ is the predicted attenuation coefficient of the protein).

##### Far-UV CD spectroscopy

**Spectral CD** Far-UV CD spectra were recorded between 198–250 nm using a Chirascan system (AppliedPhotopysics). Proteins at 5–10 µM were measured in a quartz cell with a 0.1 cm path length at 25 °C. All spectra were measured as θ (in mdeg) then corrected

by subtracting the buffer spectrum. Samples were then transferred to a UV/vis cuvette and their exact concentration was measured as described. Ten spectra per sample were averaged and normalised to Mean Residue Ellipticity (MRE; in $\deg \cdot \mathrm{cm}^2 \cdot \mathrm{dmol}^{-1}$) using the sample concentration (in M) and the number of residues in the protein:
$\mathrm{MRE} = \frac{\theta}{\frac{c}{10} \times 0.1\ \mathrm{cm} \times \#_{\mathrm{residues}}}$

**Thermal denaturation CD** Samples were prepared as above and covered in a thin layer of mineral oil to prevent evaporation. Circular dichroism was measured at 207 nm (1 nm bandwidth) and five measurements taken over ten seconds were averaged. The signal was recorded every 0.2 °C from 25 °C to 95 °C (cell temperature) at a rate of 1 °C $\mathrm{min}^{-1}$ and corrected using the sample temperature, measured by a probe. A two-state unfolding model was used to fit the data for determination of melting temperature.

**Nano differential scanning fluorimetry (nanoDSF)** nanoDSF was measured using a Tycho™ instrument (NanoTemper) following manufacturer instructions.

##### Intact protein LC-MS

Samples were prepared by transferring 20 µL into 1.5 mL microcentrifuge tubes and centrifuging for 7 min / 4 °C / 20000 × g. 12 µL of the supernatant liquids were then transferred to 300 µL screw thread vials (Supelco) with pre-slit screw thread caps (Fisher Scientific), and injected to an Acquity UPLC system using an Acquity UPLC Protein BEH C4 column (300 Å pore diameter, 1.7 µm, 2.1 mm × 50 mm) held at 40 °C. Solvent A consisted of 0.1 % formic acid in water, and solvent B of 5 % water + 0.1 % formic acid in acetonitrile. At a flow rate of 0.2 mL/min, a gradient of 7.29 min was utilised for most samples (hold at 5 % B for 0.93 min, increase to 100 % B over 4.28 min, hold for another 1.04 min then decrease back to 5 % B over 1.04 min). For better separation of some samples, a longer gradient of 19.9 min was used (hold at 5 % B for 1.0 min, jump to 24 % B in 0.1 min and hold for 1.9 min, then slowly up to 55 % B over 14 min, jump to 100 % B in 0.1 min and hold for another 1.7 min then decrease back to 5 % B over 0.1 min and hold for 1.1 min). Data were recorded on a Waters Xevo G2-S qTOF mass spectrometer using an electrospray ionisation (ESI) source at 120 °C with a capillary voltage of 3.0 kV and a cone voltage of 40 V. Nitrogen was used as the desolvation gas at a flow rate of 600 L/h. LockSpray was infused throughout the measurement for internal calibration.

**Basic analysis** The recorded 3-dimensional spectra (retention time × m/z × intensity) were searched for protein peaks by a combination of manual inspection and a custom-written python library. Each peak was then 'summed' (marginalised over retention times) to receive a 2d spectrum of the entire peak (m/z × intensity). These spectra were deconvoluted using UniDec [16] with the following parameters:

- For nanobodies (~15 kDa): m/z range 600–2000; charge range 6–55; mass range 12000–32000 or 12000–16000; sampling resolution 1 Da.

- For GFP (~27 kDa): m/z range 600–1800; charge range 12–55; mass range 24000–32000; sampling resolution 1 Da.

Resulting spectra (intact mass × intensity), internally normalised to the range 0–1, were used to identify the abundant species in solution.

**Semi-quantitative analysis** After identification of the existing species in each sample, the peaks representing ions corresponding to each species were identified using a custom-written python library, using a combination of predictions and observed signals. In brief, the first prediction for a given mass $M$ consisted of 20 ion peaks, with its "anchor ion" closest to 1000 Da (e.g. for $M = 14179$, the anchor ion would be $z = 14$, $m/z = \frac{14179+14}{14} = 1013.79$), and ranging from two z-values below (e.g. $z = 12$, $m/z = \frac{14179+12}{12} = 1182.59$) to 18 z-values above the anchor (e.g. $z = 32$, $m/z = \frac{14179+32}{32} = 444.10$). In parallel, the theoretical span of masses for the species arising from isotopic variability was predicted using an estimation of the number of carbon atoms and a 1.11 % natural abundance of carbon-13. Based on this, each peak was given the predicted width of this span divided by the corresponding z-value. Using these predictions and an iterative process incorporated the recorded data for the sample, these predictions were refined to catch as many ion peaks as possible without including m/z values arising from other species in solution or from measurement noise.

Once predictions were refined, the corresponding m/z ranges were sliced out from the 3-dimensional spectrum and marginalised over the m/z axis to form a 2-dimensional extracted ion chromatogram (XIC) representing the species. The process was repeated for each relevant species. To further increase signal-to-noise ratios, the retention time (rt) range corresponding to a group of species (e.g. all nanobody derivatives) was identified by manual inspection of the XICs.

Quantification of species followed directly by summing their corresponding XICs in the identified rt range. As discussed, the absolute values derived from this quantification cannot be used directly, but the relative values within a group of species was found to capture their relative abundances well, and was consistent across independent experiments conduced over a range of months or years and a range of concentrations and injection volumes.

##### Surface plasmon resonance (SPR)

SPR analysis was carried out on a Biacore™ T200 instrument (GE Healthcare). Nanobodies were immobilised on a Series S Sensor Chip CM5 (Cytiva): the chip surface was primed with the running buffer (PBS pH 7.4 + 0.01% Tween-20) and activated with 50 mM N-hydroxysuccinimide (NHS) and 200 mM 1-ethyl-3-(3-dimethylaminopropyl) carbodiimide hydrochloride (EDC) at a flow rate of 10 µL/min for 420 seconds. The ligand was diluted to 1 µM in 10 mM sodium acetate, pH 5.5 and loaded onto the sample flow cell surface at a flow rate of 5 µL/min in short injections totalling between 540–840 seconds. The sensor chip surface was deactivated with 1 M ethanolamine·HCl, pH 8.5 at a flow rate of 10 µL/min for 420 seconds. Surface activity was tested by performing four injections of 30 seconds each of 10 mM glycine, pH 2.0 at a flow rate of 30 µL/min.

Recombinant GFP analyte was injected in 7 concentrations ranging 0.03–333 nM in duplicate, at a flow rate of 30 µL/min for 240 seconds, followed by a dissociation time of 1200 seconds. After each cycle, the sensor surface was regenerated by performing two injections of 30 seconds each of 10 mM glycine, pH 2.0 at a flow rate of 30 µL/min, followed by a stabilisation period of 60 seconds. The binding sensorgrams were fitted according to a 1 : 1 binding model and the data were processed using Biacore T200 Evaluation software.

##### Biolayer interferometry (BLI)

BLI measurements were conducted using an Octet-BLI K2 system (ForteBio) with streptavidin (SA) sensors. To prevent non-specific interactions with the sensors, all experiments were carried out in PBS supplemented with 0.05% Tween-20 (Sigma). A black 96-well plate (Greiner 655209) was used, with 200 µl per well, and the sensors were pre-hydrated in the assay buffer for at least 15 minutes before use. The assay plate was maintained at 30°C / 1000 rpm. The loading wells were loaded with 50 nM of biotinylated β2-microglobulin, as previously described [12].

Each experimental run consisted of a baseline step, a loading step, a second baseline step, followed by multiple association and brief dissociation steps. The association phase was observed in wells containing six different concentrations of nb24 nanobody variants ranging 25–400 nM. In all experiments, a reference sensor, treated in the same way as the assay sensors but placed in buffer wells during association steps, was employed. Its signal was subtracted from the signal of each assay sensor before data analysis. The binding data were then globally fitted with a 1:1 partial dissociation binding model, using Rmax, on-rate, and off-rate as global parameters, and $\mathtt{Y}_{t\rightarrow\text{inf}}$ as a local parameter.

##### Small molecule nuclear magnetic resonance (NMR) spectroscopy

Samples for NMR were dissolved in deuterated solvents as indicated, at concentrations ranging from 10 mg/mL (short $^{1}$H spectra) to 50 mg/mL (overnight $^{13}$C or other complex spectra). Once fully dissolved, samples were transferred to Wilmad 528-PP 7 inch × 5 mm NMR tubes and loaded onto a 400 MHz Avance III HD Smart Probe spectrometer (Bruker). Data were analysed using Bruker TopSpin.

##### Small molecule LC-MS

Samples were prepared by dissolving in water or water:acetonitrile mixtures of up to 90 % acetonitrile. Once fully dissolved, the sample was filtered using a 4 mm PVDF syringe filter with a 0.22 µm pore size (Millipore) into a 300 µL screw thread vial (Supelco) and covered with a pre-slit screw thread cap (Fisher Scientific). Samples were injected on an

Acquity UPLC system equiped with an Acquity UPLC BEH C18 column (130 Å pore diameter, 1.7 µm, 2.1 mm × 50 mm). Solvent A consisted of 0.1 % formic acid in water, and solvent B of 0.1 % formic acid in acetonitrile. At a flow rate of 0.37 mL/min, a gradient of 8.00 min was utilised (hold at 10 % B for 0.38 min, increase to 100 % B over 6.8 min, hold for another 0.38 min then decrease back to 10 % B over 0.44 min). Data were recorded on a Waters Xevo G2-S qTOF mass spectrometer using an electrospray ionisation (ESI) source at 120 °C with a capillary voltage of 3.0 kV and a cone voltage of 40 V. Nitrogen was used as the desolvation gas at a flow rate of 600 L/h. LockSpray was infused throughout the measurement for internal calibration.

#### Instrument Reports

#### File Properties

**Evaluation File**
Name: N/A

**User Information**
Performed By: Administrator
Current User: Administrator

**Created With Software**
Name: Biacore T200 Evaluation Software
Version: 1.0

**Notebook**

**Result**

**Result File**
Name: 20230225_MCK_GFP-analyte_aGFP-RSGGH-ligand_FC4-3.blr
Path: C:\Bia Users\Vaidehi vr358
Size: 6 625 280 bytes

**Run Information**
Type: Kinetics/Affinity
Cycles: 21
Start: 3/25/2023 7:43:02 PM
End: 3/26/2023 7:44:59 AM

**Instrument**
Instrument Type: BiacoreT200
Instrument Id: 1646191
IFC: TYPE105

**User Information**
Run Performed By: Administrator

**Created With Software**
Name: Biacore T200 Control Software
Version: 1.0

**Chip Information**
Chip Id: Vaidehi 230321 aGFP wt
Chip Lot No: 10335900
Chip Name: CM5
First Dock Date: 3/21/2023 1:14:13 PM
Last Modification Date: 3/21/2023 1:14:13 PM
Last Use Date: 3/25/2023 11:26:54 AM

**Immobilization in Fc=1**
Immobilization Date:
Immobilization Result:
Ligand:
Final Response [RU]:

**Immobilization in Fc=2**
Immobilization Date:
Immobilization Result:
Ligand:
Final Response [RU]:

**Immobilization in Fc=3**
Immobilization Date:
Immobilization Result:
Ligand:
Final Response [RU]:

**Immobilization in Fc=4**
Immobilization Date:

#### File Properties (continued)

Immobilization Result:
Ligand:
Final Response [RU]:

**Notebook**

**Kinetics: 'GFP 2', fit: '1. 1:1 Binding'**

Curve: Fc=4-3 Ligand: N/A Sample: GFP Temp: 25°C

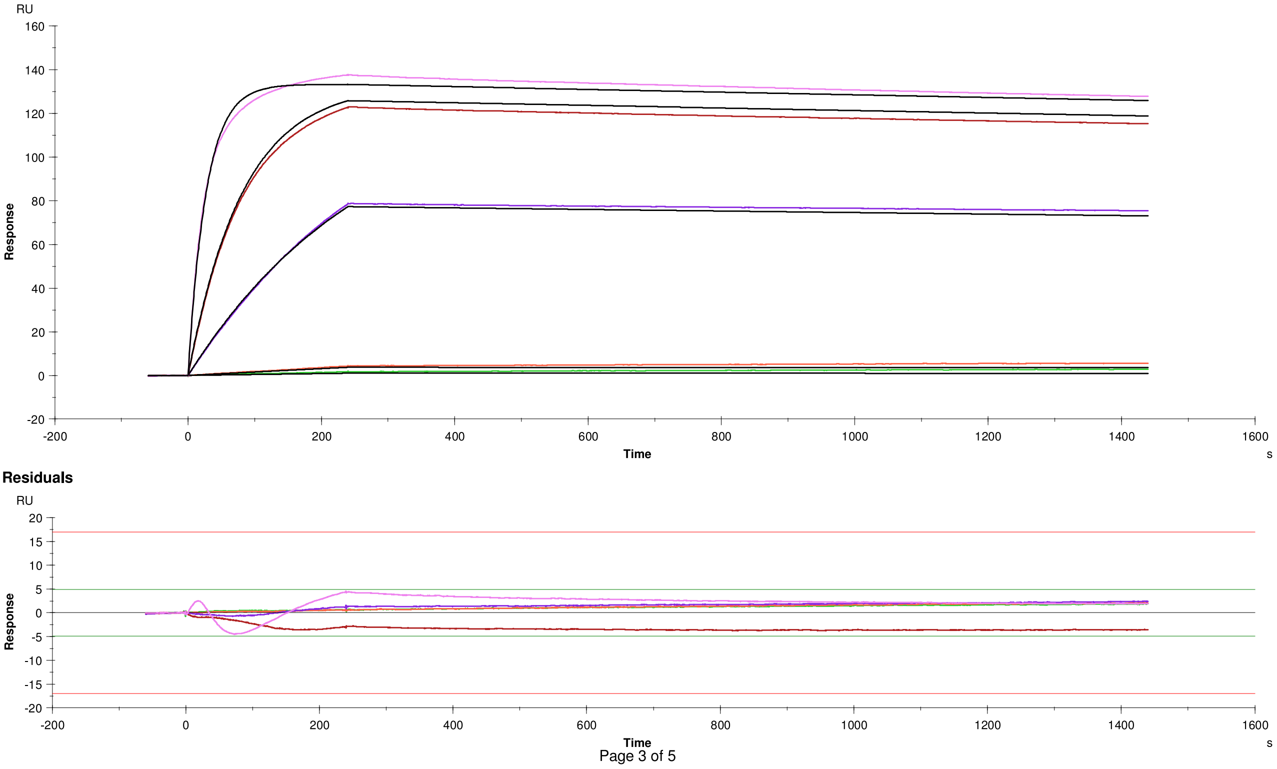

#### Kinetics: ‘GFP 2’, fit: ‘1. 1:1 Binding’ (continued)

**Quality Control**

| | |
|---|---|
| | Kinetic constants are within instrument specifications. |
| | Kinetic constants appear to be uniquely determined. |
| | Bulk contributions (RI) were not evaluated. The RI parameter is set to constant. |
| | Check that sensorgrams have sufficient curvature. |
| | Examine the residual plot. Pay attention to systematic and non-random deviations. |

#### Kinetics: 'GFP 2', fit: '1. 1:1 Binding' (continued)

##### Report table

| Curve | ka (1/Ms) | kd (1/s) | KD (M) | Rmax (RU) | Conc (M) | tc | Flow (ul/min) | kt (RU/Ms) | RI (RU) | Chi² (RU²) | U-value |
|---|---|---|---|---|---|---|---|---|---|---|---|
| | 1.093E+6 | 4.756E-5 | 4.350E-11 | 133.5 | | 1.820E+19 | | | | 4.83 | 7 |
| Cycle: 6 0.03 nM | | | | | 3.000E-11 | | 30.00 | 5.655E+19 | 0.000 | | |
| Cycle: 7 0.11 nM | | | | | 1.100E-10 | | 30.00 | 5.655E+19 | 0.000 | | |
| Cycle: 8 3.33 nM | | | | | 3.330E-9 | | 30.00 | 5.655E+19 | 0.000 | | |
| Cycle: 9 11.11 nM | | | | | 1.111E-8 | | 30.00 | 5.655E+19 | 0.000 | | |
| Cycle: 10 33.33 nM | | | | | 3.333E-8 | | 30.00 | 5.655E+19 | 0.000 | | |

##### Parameters table

| Curve | ka (1/Ms) | SE(ka) | kd (1/s) | SE(kd) | Rmax (RU) | SE(Rmax) | Conc (M) | tc | SE(tc) | f (ul/min) | RI (RU) |
|---|---|---|---|---|---|---|---|---|---|---|---|
| | 1.093E+6 | 4.9E+2 | 4.756E-5 | 2.5E-7 | 133.5 | 0.026 | | 1.820E+19 | 5.7E+21 | | |
| Cycle: 6 0.03 nM | | | | | | | 3E-11 | | | 30 | 0.0 |
| Cycle: 7 0.11 nM | | | | | | | 1.1E-10 | | | 30 | 0.0 |
| Cycle: 8 3.33 nM | | | | | | | 3.33E-09 | | | 30 | 0.0 |
| Cycle: 9 11.11 nM | | | | | | | 1.111E-08 | | | 30 | 0.0 |
| Cycle: 10 33.33 nM | | | | | | | 3.333E-08 | | | 30 | 0.0 |

#### File Properties

**Evaluation File**

| | |
|---|---|
| Name: | 20230221_MCK_GFP-analyte_aGFP-S54C-RSGGH_FC4-3_333-removed.bme |
| Path: | C:\Bia Users\Vaidehi vr358 |
| Size: | 10 861 056 |

**User Information**

| | |
|---|---|
| Performed By: | Administrator |
| Current User: | Administrator |

**Created With Software**

| | |
|---|---|
| Name: | Biacore T200 Evaluation Software |
| Version: | 1.0 |

**Notebook**

**Result**

**Result File**

| | |
|---|---|
| Name: | 20230221_MCK_GFP-analyte_aGFP-S54C-RSGGH_FC4-3.blr |
| Path: | C:\Bia Users\Vaidehi vr358 |
| Size: | 6 625 280 bytes |

**Run Information**

| | |
|---|---|
| Type: | Kinetics/Affinity |
| Cycles: | 21 |
| Start: | 2/21/2023 8:02:06 PM |
| End: | 2/22/2023 7:03:41 AM |

**Instrument**

| | |
|---|---|
| Instrument Type: | BiacoreT200 |
| Instrument Id: | 1646191 |
| IFC: | TYPE105 |

**User Information**

| | |
|---|---|
| Run Performed By: | Administrator |

**Created With Software**

| | |
|---|---|
| Name: | Biacore T200 Control Software |
| Version: | 1.0 |

**Chip Information**

| | |
|---|---|
| Chip Id: | 230220-1251:Vaidehi |
| Chip Lot No: | 10331800 |
| Chip Name: | CM5 |
| First Dock Date: | 2/20/2023 12:54:32 PM |
| Last Modification Date: | 2/20/2023 12:54:32 PM |
| Last Use Date: | 2/21/2023 2:32:53 PM |

**Immobilization in Fc=1**

Immobilization Date:
Immobilization Result:
Ligand:
Final Response [RU]:

**Immobilization in Fc=2**

Immobilization Date:
Immobilization Result:
Ligand:
Final Response [RU]:

**Immobilization in Fc=3**

Immobilization Date:
Immobilization Result:
Ligand:
Final Response [RU]:

#### File Properties (continued)

**Immobilization in Fc=4**
Immobilization Date:
Immobilization Result:
Ligand:
Final Response [RU]:

**Notebook**

**Kinetics: 'GFP 2', fit: '1. 1:1 Binding'**

Curve: Fc=4-3 Ligand: N/A Sample: GFP Temp: 25°C

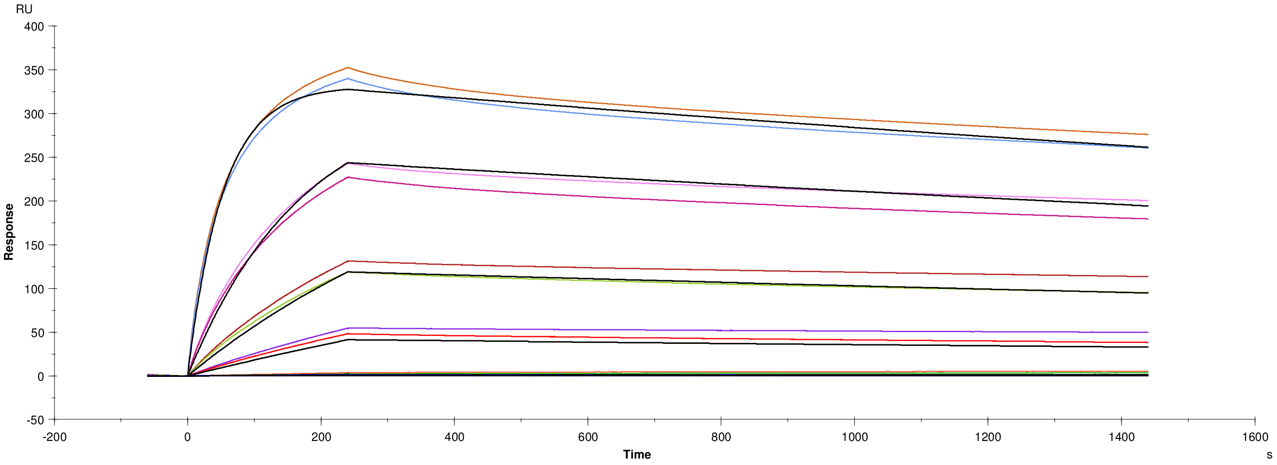

**Residuals**

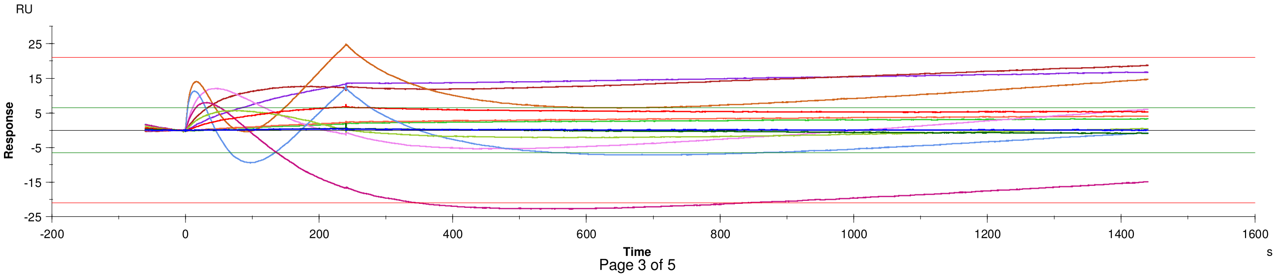

#### Kinetics: 'GFP 2', fit: '1. 1:1 Binding' (continued)

**Quality Control**

| | |
|---|---|
| ✓ | Kinetic constants are within instrument specifications. |
| ✓ | Kinetic constants appear to be uniquely determined. |
| ⇨ | Bulk contributions (RI) were not evaluated. The RI parameter is set to constant. |
| ⇨ | Check that sensorgrams have sufficient curvature. |
| ⇨ | Examine the residual plot. Pay attention to systematic and non-random deviations. |

#### Kinetics: ‘GFP 2’, fit: ‘1. 1:1 Binding’ (continued)

**Report table**

| Curve | ka (1/Ms) | kd (1/s) | KD (M) | Rmax (RU) | Conc (M) | tc | Flow (ul/min) | kt (RU/Ms) | RI (RU) | Chi² (RU²) | U-value |
|---|---|---|---|---|---|---|---|---|---|---|---|
| | 1.690E+5 | 1.883E-4 | 1.115E-9 | 334.7 | | 8.011E+17 | | | | 77.5 | 4 |
| Cycle: 6 0.03 nM | | | | | 3.000E-11 | | 30.00 | 2.489E+18 | 0.000 | | |
| Cycle: 7 0.11 nM | | | | | 1.100E-10 | | 30.00 | 2.489E+18 | 0.000 | | |
| Cycle: 8 3.33 nM | | | | | 3.330E-9 | | 30.00 | 2.489E+18 | 0.000 | | |
| Cycle: 9 11.11 nM | | | | | 1.111E-8 | | 30.00 | 2.489E+18 | 0.000 | | |
| Cycle: 10 33.33 nM | | | | | 3.333E-8 | | 30.00 | 2.489E+18 | 0.000 | | |
| Cycle: 11 111.11 nM | | | | | 1.111E-7 | | 30.00 | 2.489E+18 | 0.000 | | |
| Cycle: 14 111.11 nM | | | | | 1.111E-7 | | 30.00 | 2.489E+18 | 0.000 | | |
| Cycle: 15 33.33 nM | | | | | 3.333E-8 | | 30.00 | 2.489E+18 | 0.000 | | |
| Cycle: 16 11.11 nM | | | | | 1.111E-8 | | 30.00 | 2.489E+18 | 0.000 | | |
| Cycle: 17 3.33 nM | | | | | 3.330E-9 | | 30.00 | 2.489E+18 | 0.000 | | |
| Cycle: 18 0.11 nM | | | | | 1.100E-10 | | 30.00 | 2.489E+18 | 0.000 | | |
| Cycle: 19 0.03 nM | | | | | 3.000E-11 | | 30.00 | 2.489E+18 | 0.000 | | |

**Parameters table**

| Curve | ka (1/Ms) | SE(ka) | kd (1/s) | SE(kd) | Rmax (RU) | SE(Rmax) | Conc (M) | tc | SE(tc) | f (ul/min) | RI (RU) |
|---|---|---|---|---|---|---|---|---|---|---|---|
| | 1.690E+5 | 91 | 1.883E-4 | 3.7E-7 | 334.7 | 0.096 | | 8.011E+17 | 1.9E+20 | | |
| Cycle: 6 0.03 nM | | | | | | | 3E-11 | | | 30 | 0.0 |
| Cycle: 7 0.11 nM | | | | | | | 1.1E-10 | | | 30 | 0.0 |
| Cycle: 8 3.33 nM | | | | | | | 3.33E-09 | | | 30 | 0.0 |
| Cycle: 9 11.11 nM | | | | | | | 1.111E-08 | | | 30 | 0.0 |
| Cycle: 10 33.33 nM | | | | | | | 3.333E-08 | | | 30 | 0.0 |
| Cycle: 11 111.11 nM | | | | | | | 1.111E-07 | | | 30 | 0.0 |
| Cycle: 14 111.11 nM | | | | | | | 1.111E-07 | | | 30 | 0.0 |
| Cycle: 15 33.33 nM | | | | | | | 3.333E-08 | | | 30 | 0.0 |
| Cycle: 16 11.11 nM | | | | | | | 1.111E-08 | | | 30 | 0.0 |
| Cycle: 17 3.33 nM | | | | | | | 3.33E-09 | | | 30 | 0.0 |
| Cycle: 18 0.11 nM | | | | | | | 1.1E-10 | | | 30 | 0.0 |
| Cycle: 19 0.03 nM | | | | | | | 3E-11 | | | 30 | 0.0 |

#### File Properties

**Evaluation File**

| | |
|---|---|
| Name: | 20230425_MCK_GFP-analyte_aGFP-S54C-RSGGH-2b-ligand_FC2-1_Run-2.bme |
| Path: | C:\Bia Users\Vaidehi vr358 |
| Size: | 21 491 200 |

**User Information**

| | |
|---|---|
| Performed By: | Administrator |
| Current User: | Administrator |

**Created With Software**

| | |
|---|---|
| Name: | Biacore T200 Evaluation Software |
| Version: | 1.0 |

**Notebook**

**Result**

**Result File**

| | |
|---|---|
| Name: | 20230425_MCK_GFP-analyte_aGFP-S54C-RSGGH-2b-ligand_FC2-1_Run-2.blr |
| Path: | C:\Bia Users\Vaidehi vr358 |
| Size: | 15 421 952 bytes |

**Run Information**

| | |
|---|---|
| Type: | Kinetics/Affinity |
| Cycles: | 25 |
| Start: | 4/26/2023 4:47:36 AM |
| End: | 4/27/2023 6:53:38 AM |

**Instrument**

| | |
|---|---|
| Instrument Type: | BiacoreT200 |
| Instrument Id: | 1646191 |
| IFC: | TYPE105 |

**User Information**

| | |
|---|---|
| Run Performed By: | Administrator |

**Created With Software**

| | |
|---|---|
| Name: | Biacore T200 Control Software |
| Version: | 1.0 |

**Chip Information**

| | |
|---|---|
| Chip Id: | 230425:Vaidehi PEABS* |
| Chip Lot No: | 10331800 |
| Chip Name: | CM5 |
| First Dock Date: | 4/25/2023 1:51:16 PM |
| Last Modification Date: | 4/25/2023 1:51:16 PM |
| Last Use Date: | 4/25/2023 1:52:41 PM |

**Immobilization in Fc=1**

Immobilization Date:
Immobilization Result:
Ligand:
Final Response [RU]:

**Immobilization in Fc=2**

Immobilization Date:
Immobilization Result:
Ligand:
Final Response [RU]:

**Immobilization in Fc=3**

Immobilization Date:
Immobilization Result:
Ligand:
Final Response [RU]:

#### File Properties (continued)

**Immobilization in Fc=4**
Immobilization Date:
Immobilization Result:
Ligand:
Final Response [RU]:

**Notebook**

C:\Bia Users\Vaidehi vr358\20230425_MCK_GFP-analyte_aGFP-S54C-RSGGH-2b-ligand_FC2-1_Run-2.bme (unsaved)

Instr. id: 1646191 Biacore T200 Evaluation Software, version 1.0 Print date: 10/30/2023 4:36:33 AM

**Kinetics: 'GFP 3', fit: '1. 1:1 Binding'**

Curve: Fc=2-1 Ligand: N/A Sample: GFP Temp: 25°C

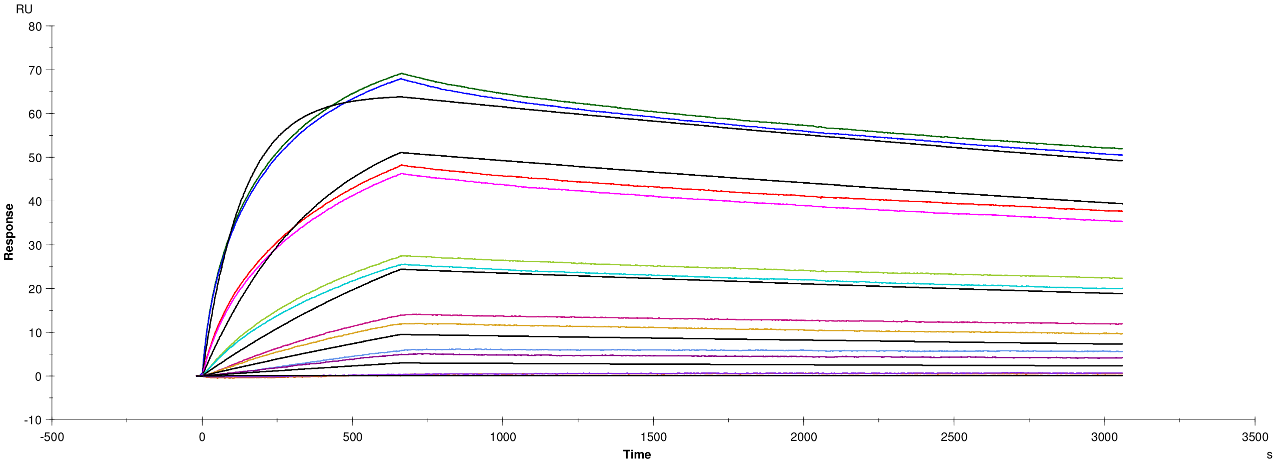

**Residuals**

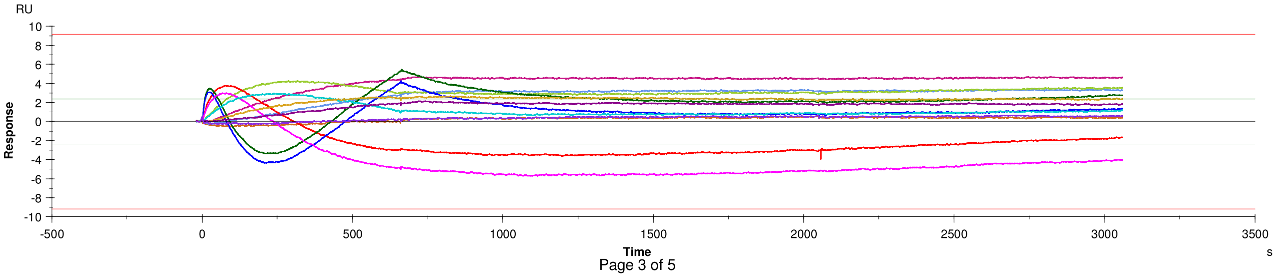

Page 3 of 5

#### Kinetics: ‘GFP 3’, fit: ‘1. 1:1 Binding’ (continued)

**Quality Control**

| | |
|---|---|
| | Kinetic constants are within instrument specifications. |
| | Kinetic constants appear to be uniquely determined. |
| | Bulk contributions (RI) were not evaluated. The RI parameter is set to constant. |
| | Check that sensorgrams have sufficient curvature. |
| | Examine the residual plot. Pay attention to systematic and non-random deviations. |

#### Kinetics: ‘GFP 3’, fit: ‘1. 1:1 Binding’ (continued)

##### Report table

| Curve | ka (1/Ms) | kd (1/s) | KD (M) | Rmax (RU) | Conc (M) | tc | Flow (ul/min) | kt (RU/Ms) | RI (RU) | Chi² (RU²) | U-value |
|---|---|---|---|---|---|---|---|---|---|---|---|
| | 2.222E+5 | 1.087E-4 | 4.894E-10 | 65.25 | | 2.139E+18 | | | | 7.31 | 5 |
| Cycle: 7 0.01 nM | | | | | 1.000E-11 | | 30.00 | 6.647E+18 | 0.000 | | |
| Cycle: 10 0.33 nM | | | | | 3.300E-10 | | 30.00 | 6.647E+18 | 0.000 | | |
| Cycle: 11 1.11 nM | | | | | 1.110E-9 | | 30.00 | 6.647E+18 | 0.000 | | |
| Cycle: 12 3.33 nM | | | | | 3.330E-9 | | 30.00 | 6.647E+18 | 0.000 | | |
| Cycle: 13 11.11 nM | | | | | 1.111E-8 | | 30.00 | 6.647E+18 | 0.000 | | |
| Cycle: 14 33.33 nM | | | | | 3.333E-8 | | 30.00 | 6.647E+18 | 0.000 | | |
| Cycle: 15 33.33 nM | | | | | 3.333E-8 | | 30.00 | 6.647E+18 | 0.000 | | |
| Cycle: 16 11.11 nM | | | | | 1.111E-8 | | 30.00 | 6.647E+18 | 0.000 | | |
| Cycle: 17 3.33 nM | | | | | 3.330E-9 | | 30.00 | 6.647E+18 | 0.000 | | |
| Cycle: 18 1.11 nM | | | | | 1.110E-9 | | 30.00 | 6.647E+18 | 0.000 | | |
| Cycle: 19 0.33 nM | | | | | 3.300E-10 | | 30.00 | 6.647E+18 | 0.000 | | |
| Cycle: 22 0.01 nM | | | | | 1.000E-11 | | 30.00 | 6.647E+18 | 0.000 | | |

##### Parameters table

| Curve | ka (1/Ms) | SE(ka) | kd (1/s) | SE(kd) | Rmax (RU) | SE(Rmax) | Conc (M) | tc | SE(tc) | f (ul/min) | RI (RU) |
|---|---|---|---|---|---|---|---|---|---|---|---|
| | 2.222E+5 | 1.3E+2 | 1.087E-4 | 1.9E-7 | 65.3 | 0.020 | | 2.139E+18 | 1.1E+21 | | |
| Cycle: 7 0.01 nM | | | | | | | 1E-11 | | | 30 | 0.0 |
| Cycle: 10 0.33 nM | | | | | | | 3.3E-10 | | | 30 | 0.0 |
| Cycle: 11 1.11 nM | | | | | | | 1.11E-09 | | | 30 | 0.0 |
| Cycle: 12 3.33 nM | | | | | | | 3.33E-09 | | | 30 | 0.0 |
| Cycle: 13 11.11 nM | | | | | | | 1.111E-08 | | | 30 | 0.0 |
| Cycle: 14 33.33 nM | | | | | | | 3.333E-08 | | | 30 | 0.0 |
| Cycle: 15 33.33 nM | | | | | | | 3.333E-08 | | | 30 | 0.0 |
| Cycle: 16 11.11 nM | | | | | | | 1.111E-08 | | | 30 | 0.0 |
| Cycle: 17 3.33 nM | | | | | | | 3.33E-09 | | | 30 | 0.0 |
| Cycle: 18 1.11 nM | | | | | | | 1.11E-09 | | | 30 | 0.0 |
| Cycle: 19 0.33 nM | | | | | | | 3.3E-10 | | | 30 | 0.0 |
| Cycle: 22 0.01 nM | | | | | | | 1E-11 | | | 30 | 0.0 |

#### File Properties

**Evaluation File**

| | |
|---|---|
| Name: | N/A |

**User Information**

| | |
|---|---|
| Performed By: | Administrator |
| Current User: | Administrator |

**Created With Software**

| | |
|---|---|
| Name: | Biacore T200 Evaluation Software |
| Version: | 1.0 |

**Notebook**

**Result**

**Result File**

| | |
|---|---|
| Name: | 20230831_MCK_FC2-1_3-1_4-1_aGFPS54C-mutants.blr |
| Path: | C:\Bia Users\Vaidehi vr358 |
| Size: | 25 669 120 bytes |

**Run Information**

| | |
|---|---|
| Type: | Method Builder |
| Method: | C:\Bia Users\Methods And Templates\Vaidehi vr358\20230902_MCK_2-1_3-1_4-1_METHOD.Method |
| Cycles: | 19 |
| Start: | 9/2/2023 9:41:03 PM |
| End: | 9/3/2023 3:12:01 PM |

**Instrument**

| | |
|---|---|
| Instrument Type: | BiacoreT200 |
| Instrument Id: | 1646191 |
| IFC: | TYPE105 |

**User Information**

| | |
|---|---|
| Run Performed By: | Administrator |

**Created With Software**

| | |
|---|---|
| Name: | Biacore T200 Control Software |
| Version: | 1.0 |

**Chip Information**

| | |
|---|---|
| Chip Id: | 230806-0953: Vaidehi CM5 |
| Chip Lot No: | |
| Chip Name: | CM5 |
| First Dock Date: | 8/6/2023 9:54:13 PM |
| Last Modification Date: | 8/6/2023 9:54:13 PM |
| Last Use Date: | 8/31/2023 5:44:33 PM |

**Immobilization in Fc=1**

Immobilization Date:
Immobilization Result:
Ligand:
Final Response [RU]:

**Immobilization in Fc=2**

Immobilization Date:
Immobilization Result:
Ligand:
Final Response [RU]:

**Immobilization in Fc=3**

Immobilization Date:
Immobilization Result:
Ligand:
Final Response [RU]:

**Immobilization in Fc=4**

#### File Properties (continued)

Immobilization Date:
Immobilization Result:
Ligand:
Final Response [RU]:

**Notebook**

Unsaved file

Instr. id: 1646191 | Biacore T200 Evaluation Software, version 1.0 | Print date: 10/30/2023 3:00:18 AM

**Kinetics: ‘M1_FC4-1’, fit: ‘1. 1:1 Binding’**

Curve: Fc=4-1 Ligand: N/A Sample: GFP Temp: 25°C

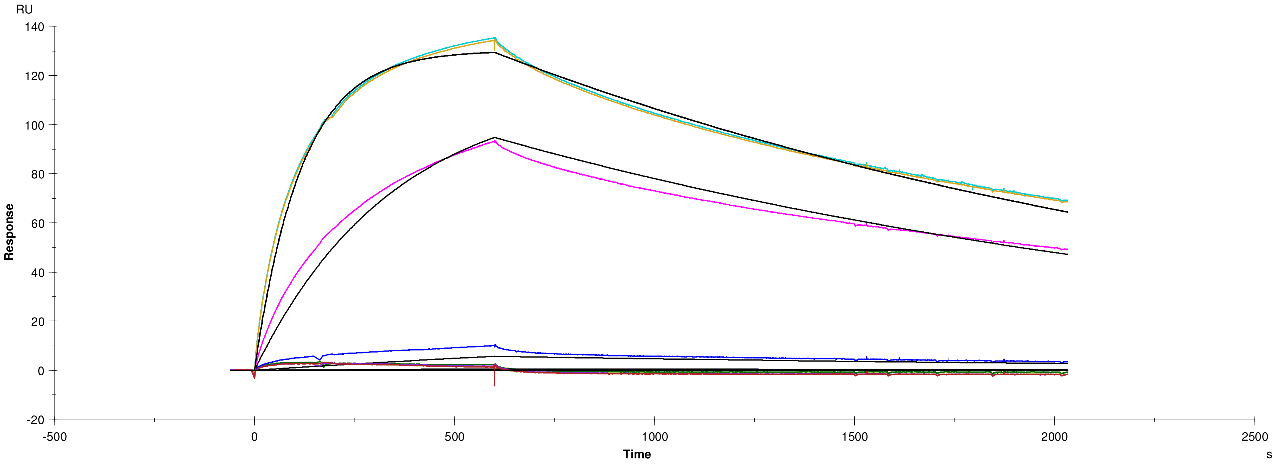

**Residuals**

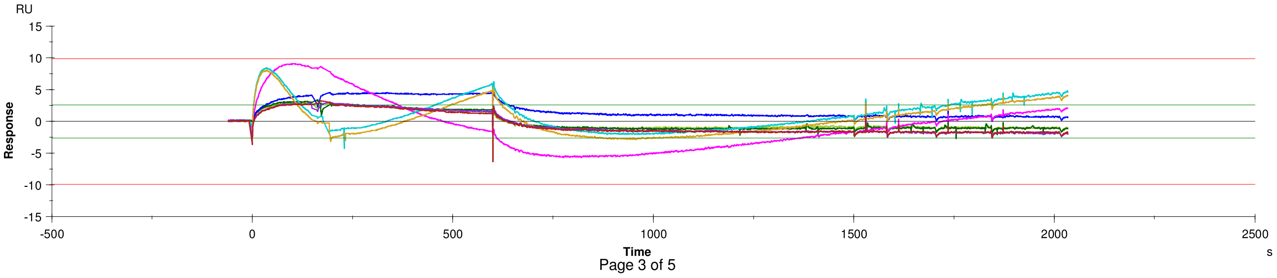

Page 3 of 5

#### Kinetics: ‘M1_FC4-1’, fit: ‘1. 1:1 Binding’ (continued)

**Quality Control**

| | |
|---|---|
| ✓ | Kinetic constants are within instrument specifications. |
| ✓ | Kinetic constants appear to be uniquely determined. |
| ⇨ | Bulk contributions (RI) were not evaluated. The RI parameter is set to constant. |
| ⇨ | Check that sensorgrams have sufficient curvature. |
| ⇨ | Examine the residual plot. Pay attention to systematic and non-random deviations. |

#### Kinetics: 'M1_FC4-1', fit: '1. 1:1 Binding' (continued)

##### Report table

| Curve | ka (1/Ms) | kd (1/s) | KD (M) | Rmax (RU) | Conc (M) | tc | Flow (ul/min) | kt (RU/Ms) | RI (RU) | Chi² (RU²) | U-value |
|---|---|---|---|---|---|---|---|---|---|---|---|
| | 7.248E+4 | 4.868E-4 | 6.716E-9 | 138.1 | | 3.583E+17 | | | | 5.65 | 1 |
| Cycle: 6 0.003 nM | | | | | 3.000E-12 | | 30.00 | 1.113E+18 | 0.000 | | |
| Cycle: 8 0.11 nM | | | | | 1.100E-10 | | 30.00 | 1.113E+18 | 0.000 | | |
| Cycle: 9 1.11 nM | | | | | 1.110E-9 | | 30.00 | 1.113E+18 | 0.000 | | |
| Cycle: 10 33.33 nM | | | | | 3.333E-8 | | 30.00 | 1.113E+18 | 0.000 | | |
| Cycle: 11 111.11 nM | | | | | 1.111E-7 | | 30.00 | 1.113E+18 | 0.000 | | |
| Cycle: 12 111.11 nM | | | | | 1.111E-7 | | 30.00 | 1.113E+18 | 0.000 | | |
| Cycle: 16 0.03 nM | | | | | 3.000E-11 | | 30.00 | 1.113E+18 | 0.000 | | |
| Cycle: 17 0.003 nM | | | | | 3.000E-12 | | 30.00 | 1.113E+18 | 0.000 | | |

##### Parameters table

| Curve | ka (1/Ms) | SE(ka) | kd (1/s) | SE(kd) | Rmax (RU) | SE(Rmax) | Conc (M) | tc | SE(tc) | f (ul/min) | RI (RU) |
|---|---|---|---|---|---|---|---|---|---|---|---|
| | 7.248E+4 | 43 | 4.868E-4 | 2.6E-7 | 138.1 | 0.030 | | 3.583E+17 | 1.6E+20 | | |
| Cycle: 6 0.003 nM | | | | | | | 3E-12 | | | 30 | 0.0 |
| Cycle: 8 0.11 nM | | | | | | | 1.1E-10 | | | 30 | 0.0 |
| Cycle: 9 1.11 nM | | | | | | | 1.11E-09 | | | 30 | 0.0 |
| Cycle: 10 33.33 nM | | | | | | | 3.333E-08 | | | 30 | 0.0 |
| Cycle: 11 111.11 nM | | | | | | | 1.111E-07 | | | 30 | 0.0 |
| Cycle: 12 111.11 nM | | | | | | | 1.111E-07 | | | 30 | 0.0 |
| Cycle: 16 0.03 nM | | | | | | | 3E-11 | | | 30 | 0.0 |
| Cycle: 17 0.003 nM | | | | | | | 3E-12 | | | 30 | 0.0 |

#### File Properties

**Evaluation File**
Name: N/A

**User Information**
Performed By: Administrator
Current User: Administrator

**Created With Software**
Name: Biacore T200 Evaluation Software
Version: 1.0

**Notebook**

**Result**

**Result File**
Name: 20230831_MCK_FC2-1_3-1_4-1_aGFPS54C-mutants.blr
Path: C:\Bia Users\Vaidehi vr358
Size: 25 669 120 bytes

**Run Information**
Type: Method Builder
Method: C:\Bia Users\Methods And Templates\Vaidehi vr358\20230902_MCK_2-1_3-1_4-1_METHOD.Method
Cycles: 19
Start: 9/2/2023 9:41:03 PM
End: 9/3/2023 3:12:01 PM

**Instrument**
Instrument Type: BiacoreT200
Instrument Id: 1646191
IFC: TYPE105

**User Information**
Run Performed By: Administrator

**Created With Software**
Name: Biacore T200 Control Software
Version: 1.0

**Chip Information**
Chip Id: 230806-0953: Vaidehi CM5
Chip Lot No:
Chip Name: CM5
First Dock Date: 8/6/2023 9:54:13 PM
Last Modification Date: 8/6/2023 9:54:13 PM
Last Use Date: 8/31/2023 5:44:33 PM

**Immobilization in Fc=1**
Immobilization Date:
Immobilization Result:
Ligand:
Final Response [RU]:

**Immobilization in Fc=2**
Immobilization Date:
Immobilization Result:
Ligand:
Final Response [RU]:

**Immobilization in Fc=3**
Immobilization Date:
Immobilization Result:
Ligand:
Final Response [RU]:

**Immobilization in Fc=4**

#### File Properties (continued)

Immobilization Date:
Immobilization Result:
Ligand:
Final Response [RU]:

**Notebook**

**Kinetics: ‘M3_FC3-1’, fit: ‘1. 1:1 Binding’**

Curve: Fc=3-1 Ligand: N/A Sample: GFP Temp: 25°C

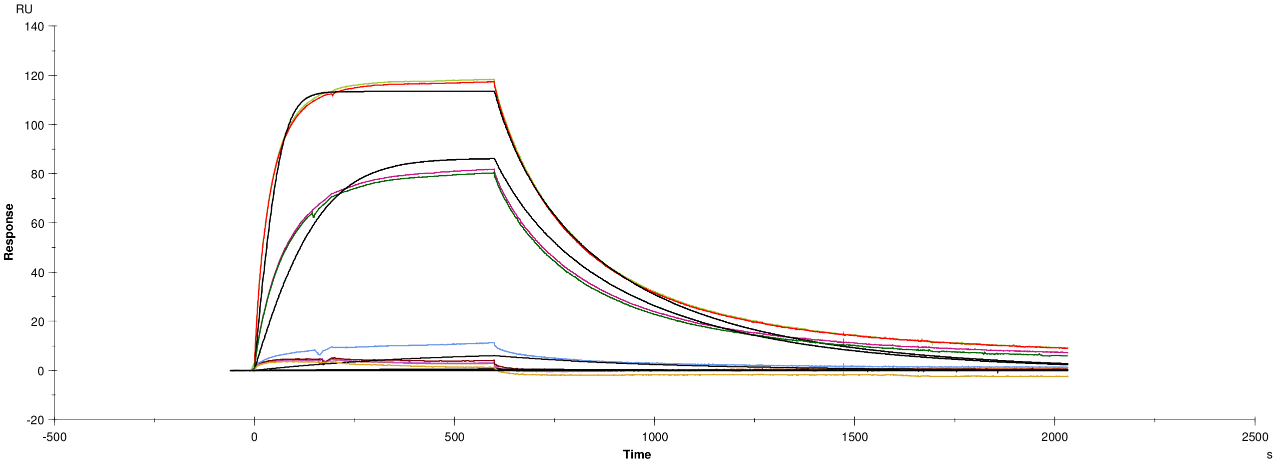

**Residuals**

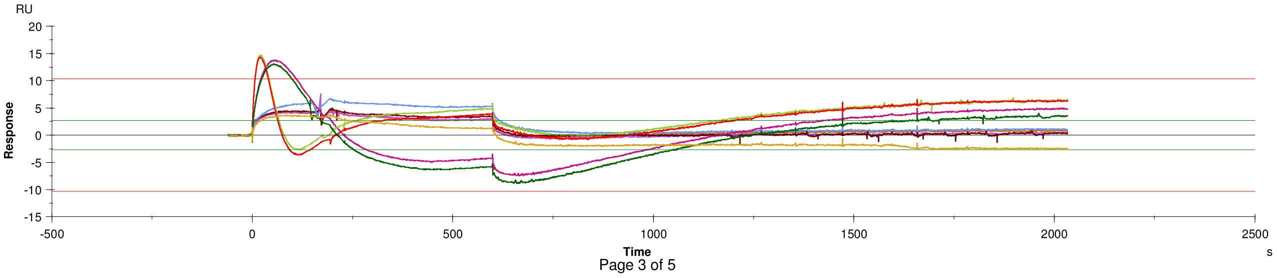

#### Kinetics: 'M3_FC3-1', fit: '1. 1:1 Binding' (continued)

**Quality Control**

| | |
|---|---|
| ✓ | Kinetic constants are within instrument specifications. |
| ✓ | Kinetic constants appear to be uniquely determined. |
| → | Bulk contributions (RI) were not evaluated. The RI parameter is set to constant. |
| → | Check that sensorgrams have sufficient curvature. |
| → | Examine the residual plot. Pay attention to systematic and non-random deviations. |

#### Kinetics: ‘M3_FC3-1’, fit: ‘1. 1:1 Binding’ (continued)

**Report table**

| Curve | ka (1/Ms) | kd (1/s) | KD (M) | Rmax (RU) | Conc (M) | tc | Flow (ul/min) | kt (RU/Ms) | RI (RU) | Chi² (RU²) | U-value |
|---|---|---|---|---|---|---|---|---|---|---|---|
| | 3.827E+5 | 0.006635 | 1.733E-8 | 131.2 | | 8.025E+6 | | | | 12.0 | 5 |
| Cycle: 6 0.003 nM | | | | | 3.000E-12 | | 30.00 | 2.493E+7 | 0.000 | | |
| Cycle: 7 0.03 nM | | | | | 3.000E-11 | | 30.00 | 2.493E+7 | 0.000 | | |
| Cycle: 8 0.11 nM | | | | | 1.100E-10 | | 30.00 | 2.493E+7 | 0.000 | | |
| Cycle: 9 1.11 nM | | | | | 1.110E-9 | | 30.00 | 2.493E+7 | 0.000 | | |
| Cycle: 10 33.33 nM | | | | | 3.333E-8 | | 30.00 | 2.493E+7 | 0.000 | | |
| Cycle: 11 111.11 nM | | | | | 1.111E-7 | | 30.00 | 2.493E+7 | 0.000 | | |
| Cycle: 12 111.11 nM | | | | | 1.111E-7 | | 30.00 | 2.493E+7 | 0.000 | | |
| Cycle: 13 33.33 nM | | | | | 3.333E-8 | | 30.00 | 2.493E+7 | 0.000 | | |
| Cycle: 17 0.003 nM | | | | | 3.000E-12 | | 30.00 | 2.493E+7 | 0.000 | | |

**Parameters table**

| Curve | ka (1/Ms) | SE(ka) | kd (1/s) | SE(kd) | Rmax (RU) | SE(Rmax) | Conc (M) | tc | SE(tc) | f (ul/min) | RI (RU) |
|---|---|---|---|---|---|---|---|---|---|---|---|
| | 3.827E+5 | 1.6E+2 | 0.006635 | 2.5E-6 | 131.2 | 0.030 | | 8.025E+6 | 3.3E+3 | | |
| Cycle: 6 0.003 nM | | | | | | | 3E-12 | | | 30 | 0.0 |
| Cycle: 7 0.03 nM | | | | | | | 3E-11 | | | 30 | 0.0 |
| Cycle: 8 0.11 nM | | | | | | | 1.1E-10 | | | 30 | 0.0 |
| Cycle: 9 1.11 nM | | | | | | | 1.11E-09 | | | 30 | 0.0 |
| Cycle: 10 33.33 nM | | | | | | | 3.333E-08 | | | 30 | 0.0 |
| Cycle: 11 111.11 nM | | | | | | | 1.111E-07 | | | 30 | 0.0 |
| Cycle: 12 111.11 nM | | | | | | | 1.111E-07 | | | 30 | 0.0 |
| Cycle: 13 33.33 nM | | | | | | | 3.333E-08 | | | 30 | 0.0 |
| Cycle: 17 0.003 nM | | | | | | | 3E-12 | | | 30 | 0.0 |

#### File Properties

**Evaluation File**

| | |
|---|---|
| Name: | N/A |

**User Information**

| | |
|---|---|
| Performed By: | Administrator |
| Current User: | Administrator |

**Created With Software**

| | |
|---|---|
| Name: | Biacore T200 Evaluation Software |
| Version: | 1.0 |

**Notebook**

**Result**

**Result File**

| | |
|---|---|
| Name: | 20231027-GFP-mutants-analyte-1-5_aGFP-S54C-RSGGH_FC2-1.blr |
| Path: | C:\Bia Users\Vaidehi vr358 |
| Size: | 24 225 280 bytes |

**Run Information**

| | |
|---|---|
| Type: | Kinetics/Affinity |
| Cycles: | 70 |
| Start: | 10/27/2023 1:54:59 PM |
| End: | 10/29/2023 5:29:48 AM |

**Instrument**

| | |
|---|---|
| Instrument Type: | BiacoreT200 |
| Instrument Id: | 1646191 |
| IFC: | TYPE105 |

**User Information**

| | |
|---|---|
| Run Performed By: | Administrator |

**Created With Software**

| | |
|---|---|
| Name: | Biacore T200 Control Software |
| Version: | 1.0 |

**Chip Information**

| | |
|---|---|
| Chip Id: | 231023 Vaidehi CM5 |
| Chip Lot No: | 10345041 |
| Chip Name: | CM5 |
| First Dock Date: | 10/23/2023 11:57:59 PM |
| Last Modification Date: | 10/23/2023 11:57:59 PM |
| Last Use Date: | 10/23/2023 11:59:23 PM |

**Immobilization in Fc=1**

Immobilization Date:
Immobilization Result:
Ligand:
Final Response [RU]:

**Immobilization in Fc=2**

Immobilization Date:
Immobilization Result:
Ligand:
Final Response [RU]:

**Immobilization in Fc=3**

Immobilization Date:
Immobilization Result:
Ligand:
Final Response [RU]:

**Immobilization in Fc=4**

Immobilization Date:

#### File Properties (continued)

Immobilization Result:
Ligand:
Final Response [RU]:

**Notebook**

**Kinetics: '41', fit: '1. 1:1 Binding'**

Curve: Fc=2-1 Ligand: N/A Sample: 41 Temp: 25°C

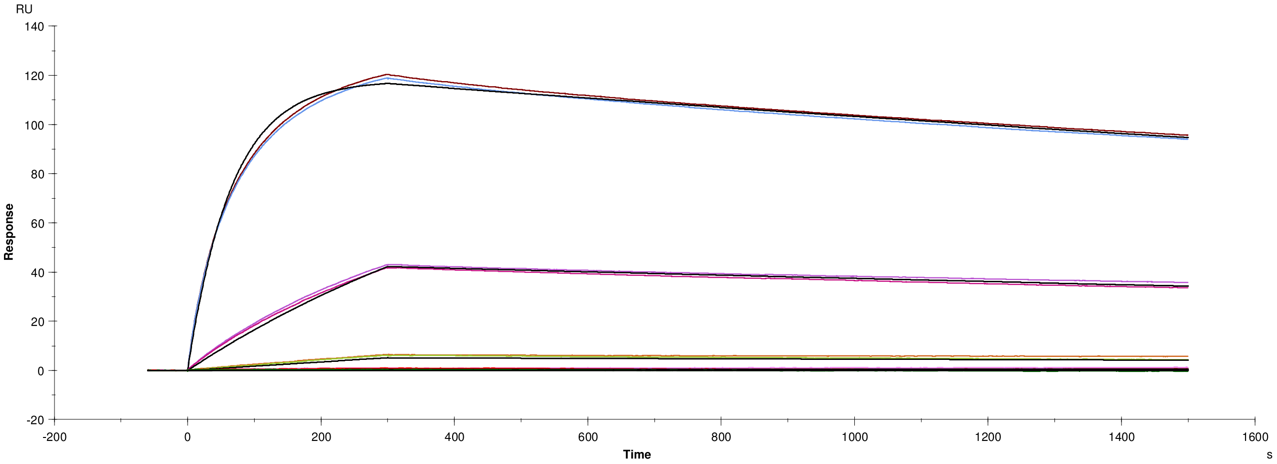

**Residuals**

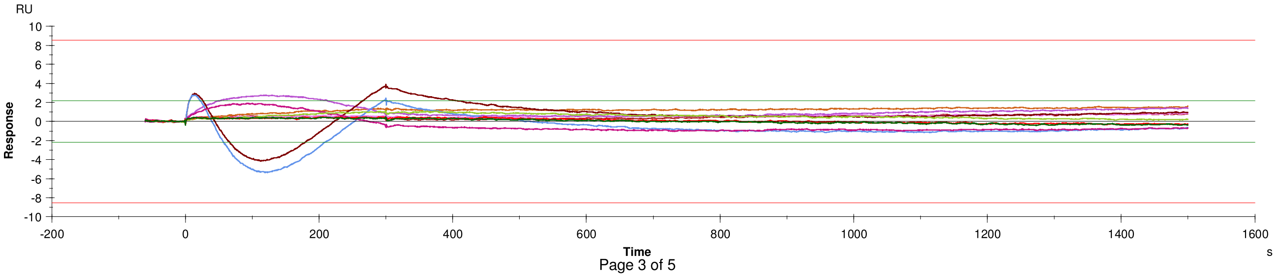

#### Kinetics: ‘41’, fit: ‘1. 1:1 Binding’ (continued)

**Quality Control**

| | |
|---|---|
| ✓ | Kinetic constants are within instrument specifications. |
| ✓ | Kinetic constants appear to be uniquely determined. |
| ➡ | Bulk contributions (RI) were not evaluated. The RI parameter is set to constant. |
| ➡ | Check that sensorgrams have sufficient curvature. |
| ➡ | Examine the residual plot. Pay attention to systematic and non-random deviations. |

#### Kinetics: '41', fit: '1. 1:1 Binding' (continued)

**Report table**

| Curve | ka (1/Ms) | kd (1/s) | KD (M) | Rmax (RU) | Conc (M) | tc | Flow (ul/min) | kt (RU/Ms) | RI (RU) | Chi² (RU²) | U-value |
|---|---|---|---|---|---|---|---|---|---|---|---|
| | 1.503E+5 | 1.740E-4 | 1.158E-9 | 119.3 | | 3.725E+17 | | | | 1.01 | 2 |
| Cycle: 3 0.01 nM | | | | | 1.000E-11 | | 30.00 | 1.158E+18 | 0.000 | | |
| Cycle: 4 0.1 nM | | | | | 1.000E-10 | | 30.00 | 1.158E+18 | 0.000 | | |
| Cycle: 5 1 nM | | | | | 1.000E-9 | | 30.00 | 1.158E+18 | 0.000 | | |
| Cycle: 6 10 nM | | | | | 1.000E-8 | | 30.00 | 1.158E+18 | 0.000 | | |
| Cycle: 7 100 nM | | | | | 1.000E-7 | | 30.00 | 1.158E+18 | 0.000 | | |
| Cycle: 8 100 nM | | | | | 1.000E-7 | | 30.00 | 1.158E+18 | 0.000 | | |
| Cycle: 9 10 nM | | | | | 1.000E-8 | | 30.00 | 1.158E+18 | 0.000 | | |
| Cycle: 10 1 nM | | | | | 1.000E-9 | | 30.00 | 1.158E+18 | 0.000 | | |
| Cycle: 11 0.1 nM | | | | | 1.000E-10 | | 30.00 | 1.158E+18 | 0.000 | | |
| Cycle: 12 0.01 nM | | | | | 1.000E-11 | | 30.00 | 1.158E+18 | 0.000 | | |

**Parameters table**

| Curve | ka (1/Ms) | SE(ka) | kd (1/s) | SE(kd) | Rmax (RU) | SE(Rmax) | Conc (M) | tc | SE(tc) | f (ul/min) | RI (RU) |
|---|---|---|---|---|---|---|---|---|---|---|---|
| | 1.503E+5 | 33 | 1.740E-4 | 1.4E-7 | 119.3 | 0.012 | | 3.725E+17 | 6.2E+19 | | |
| Cycle: 3 0.01 nM | | | | | | | 1E-11 | | | 30 | 0.0 |
| Cycle: 4 0.1 nM | | | | | | | 1E-10 | | | 30 | 0.0 |
| Cycle: 5 1 nM | | | | | | | 1E-09 | | | 30 | 0.0 |
| Cycle: 6 10 nM | | | | | | | 1E-08 | | | 30 | 0.0 |
| Cycle: 7 100 nM | | | | | | | 1E-07 | | | 30 | 0.0 |
| Cycle: 8 100 nM | | | | | | | 1E-07 | | | 30 | 0.0 |
| Cycle: 9 10 nM | | | | | | | 1E-08 | | | 30 | 0.0 |
| Cycle: 10 1 nM | | | | | | | 1E-09 | | | 30 | 0.0 |
| Cycle: 11 0.1 nM | | | | | | | 1E-10 | | | 30 | 0.0 |
| Cycle: 12 0.01 nM | | | | | | | 1E-11 | | | 30 | 0.0 |

#### File Properties

**Evaluation File**
Name: N/A

**User Information**
Performed By: Administrator
Current User: Administrator

**Created With Software**
Name: Biacore T200 Evaluation Software
Version: 1.0

**Notebook**

**Result**

**Result File**
Name: 20231027-GFP-mutants-analyte-1-5_aGFP-S54C-RSGGH_FC2-1.blr
Path: C:\Bia Users\Vaidehi vr358
Size: 24 225 280 bytes

**Run Information**
Type: Kinetics/Affinity
Cycles: 70
Start: 10/27/2023 1:54:59 PM
End: 10/29/2023 5:29:48 AM

**Instrument**
Instrument Type: BiacoreT200
Instrument Id: 1646191
IFC: TYPE105

**User Information**
Run Performed By: Administrator

**Created With Software**
Name: Biacore T200 Control Software
Version: 1.0

**Chip Information**
Chip Id: 231023 Vaidehi CM5
Chip Lot No: 10345041
Chip Name: CM5
First Dock Date: 10/23/2023 11:57:59 PM
Last Modification Date: 10/23/2023 11:57:59 PM
Last Use Date: 10/23/2023 11:59:23 PM

**Immobilization in Fc=1**
Immobilization Date:
Immobilization Result:
Ligand:
Final Response [RU]:

**Immobilization in Fc=2**
Immobilization Date:
Immobilization Result:
Ligand:
Final Response [RU]:

**Immobilization in Fc=3**
Immobilization Date:
Immobilization Result:
Ligand:
Final Response [RU]:

**Immobilization in Fc=4**
Immobilization Date:

#### File Properties (continued)

Immobilization Result:
Ligand:
Final Response [RU]:

**Notebook**

Unsaved file

Instr. id: 1646191 Biacore T200 Evaluation Software, version 1.0 Print date: 10/30/2023 1:56:42 AM

**Kinetics: ‘131’, fit: ‘1. 1:1 Binding’**

Curve: Fc=2-1 Ligand: N/A Sample: 131 Temp: 25°C

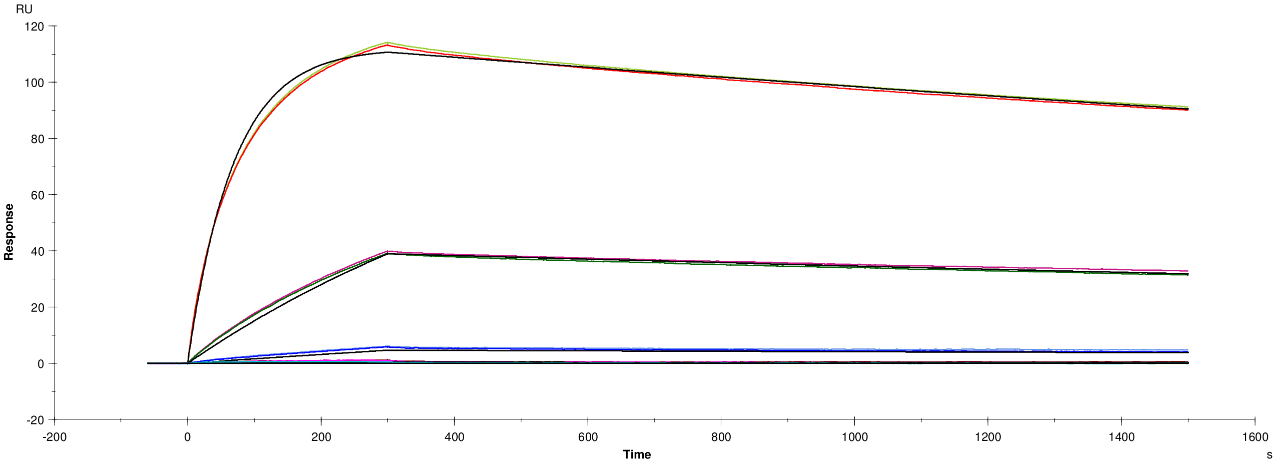

**Residuals**

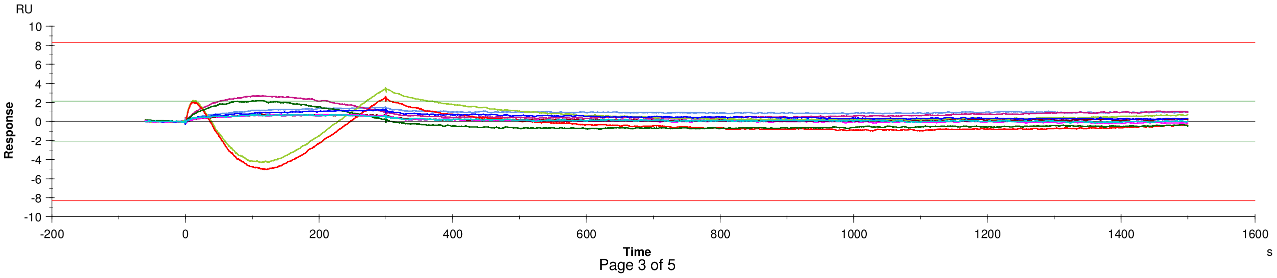

##### Kinetics: '131', fit: '1. 1:1 Binding' (continued)

**Quality Control**

| | |
|---|---|
| ✓ | Kinetic constants are within instrument specifications. |
| ✓ | Kinetic constants appear to be uniquely determined. |
| ⇨ | Bulk contributions (RI) were not evaluated. The RI parameter is set to constant. |
| ⇨ | Check that sensorgrams have sufficient curvature. |
| ⇨ | Examine the residual plot. Pay attention to systematic and non-random deviations. |

#### Kinetics: '131', fit: '1. 1:1 Binding' (continued)

##### Report table

| Curve | ka (1/Ms) | kd (1/s) | KD (M) | Rmax (RU) | Conc (M) | tc | Flow (ul/min) | kt (RU/Ms) | RI (RU) | Chi² (RU²) | U-value |
|---|---|---|---|---|---|---|---|---|---|---|---|
| | 1.445E+5 | 1.680E-4 | 1.162E-9 | 113.5 | | 6.070E+18 | | | | 0.779 | 2 |
| Cycle: 17 0.01 nM | | | | | 1.000E-11 | | 30.00 | 1.886E+19 | 0.000 | | |
| Cycle: 18 0.1 nM | | | | | 1.000E-10 | | 30.00 | 1.886E+19 | 0.000 | | |
| Cycle: 19 1 nM | | | | | 1.000E-9 | | 30.00 | 1.886E+19 | 0.000 | | |
| Cycle: 20 10 nM | | | | | 1.000E-8 | | 30.00 | 1.886E+19 | 0.000 | | |
| Cycle: 21 100 nM | | | | | 1.000E-7 | | 30.00 | 1.886E+19 | 0.000 | | |
| Cycle: 22 100 nM | | | | | 1.000E-7 | | 30.00 | 1.886E+19 | 0.000 | | |
| Cycle: 23 10 nM | | | | | 1.000E-8 | | 30.00 | 1.886E+19 | 0.000 | | |
| Cycle: 24 1 nM | | | | | 1.000E-9 | | 30.00 | 1.886E+19 | 0.000 | | |
| Cycle: 25 0.1 nM | | | | | 1.000E-10 | | 30.00 | 1.886E+19 | 0.000 | | |
| Cycle: 26 0.01 nM | | | | | 1.000E-11 | | 30.00 | 1.886E+19 | 0.000 | | |

##### Parameters table

| Curve | ka (1/Ms) | SE(ka) | kd (1/s) | SE(kd) | Rmax (RU) | SE(Rmax) | Conc (M) | tc | SE(tc) | f (ul/min) | RI (RU) |
|---|---|---|---|---|---|---|---|---|---|---|---|
| | 1.445E+5 | 30 | 1.680E-4 | 1.3E-7 | 113.5 | 0.011 | | 6.070E+18 | 1.5E+21 | | |
| Cycle: 17 0.01 nM | | | | | | | 1E-11 | | | 30 | 0.0 |
| Cycle: 18 0.1 nM | | | | | | | 1E-10 | | | 30 | 0.0 |
| Cycle: 19 1 nM | | | | | | | 1E-09 | | | 30 | 0.0 |
| Cycle: 20 10 nM | | | | | | | 1E-08 | | | 30 | 0.0 |
| Cycle: 21 100 nM | | | | | | | 1E-07 | | | 30 | 0.0 |
| Cycle: 22 100 nM | | | | | | | 1E-07 | | | 30 | 0.0 |
| Cycle: 23 10 nM | | | | | | | 1E-08 | | | 30 | 0.0 |
| Cycle: 24 1 nM | | | | | | | 1E-09 | | | 30 | 0.0 |
| Cycle: 25 0.1 nM | | | | | | | 1E-10 | | | 30 | 0.0 |
| Cycle: 26 0.01 nM | | | | | | | 1E-11 | | | 30 | 0.0 |

#### File Properties

**Evaluation File**

| | |
|---|---|
| Name: | N/A |

**User Information**

| | |
|---|---|
| Performed By: | Administrator |
| Current User: | Administrator |

**Created With Software**

| | |
|---|---|
| Name: | Biacore T200 Evaluation Software |
| Version: | 1.0 |

**Notebook**

**Result**

**Result File**

| | |
|---|---|
| Name: | 20231027-GFP-mutants-analyte-1-5_aGFP-S54C-RSGGH_FC2-1.blr |
| Path: | C:\Bia Users\Vaidehi vr358 |
| Size: | 24 225 280 bytes |

**Run Information**

| | |
|---|---|
| Type: | Kinetics/Affinity |
| Cycles: | 70 |
| Start: | 10/27/2023 1:54:59 PM |
| End: | 10/29/2023 5:29:48 AM |

**Instrument**

| | |
|---|---|
| Instrument Type: | BiacoreT200 |
| Instrument Id: | 1646191 |
| IFC: | TYPE105 |

**User Information**

| | |
|---|---|
| Run Performed By: | Administrator |

**Created With Software**

| | |
|---|---|
| Name: | Biacore T200 Control Software |
| Version: | 1.0 |

**Chip Information**

| | |
|---|---|
| Chip Id: | 231023 Vaidehi CM5 |
| Chip Lot No: | 10345041 |
| Chip Name: | CM5 |
| First Dock Date: | 10/23/2023 11:57:59 PM |
| Last Modification Date: | 10/23/2023 11:57:59 PM |
| Last Use Date: | 10/23/2023 11:59:23 PM |

**Immobilization in Fc=1**

Immobilization Date:
Immobilization Result:
Ligand:
Final Response [RU]:

**Immobilization in Fc=2**

Immobilization Date:
Immobilization Result:
Ligand:
Final Response [RU]:

**Immobilization in Fc=3**

Immobilization Date:
Immobilization Result:
Ligand:
Final Response [RU]:

**Immobilization in Fc=4**

Immobilization Date:

#### File Properties (continued)

Immobilization Result:
Ligand:
Final Response [RU]:

**Notebook**

**Kinetics: ‘140’, fit: ‘1. 1:1 Binding’**

Curve: Fc=2-1 Ligand: N/A Sample: 140 Temp: 25°C

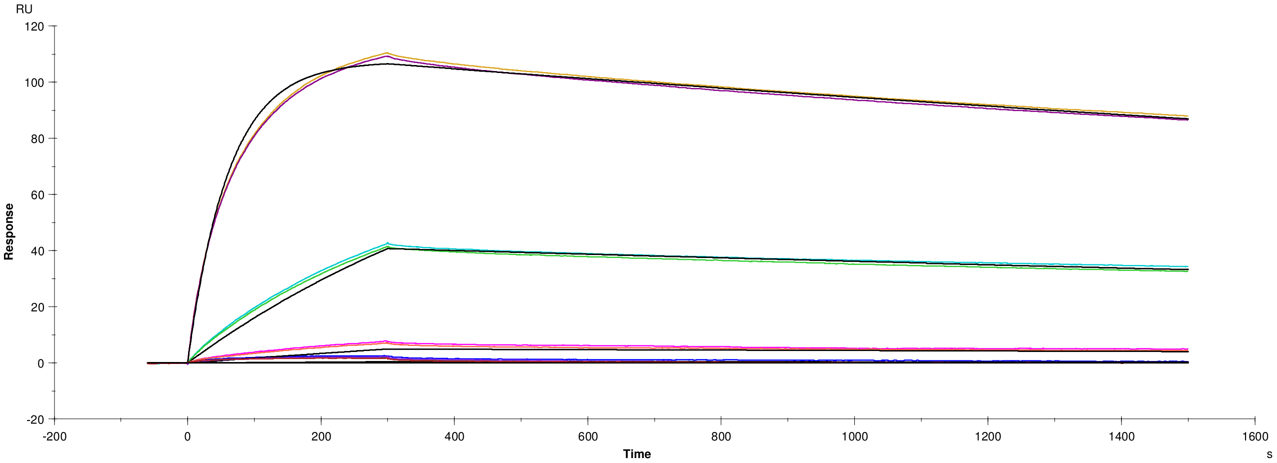

**Residuals**

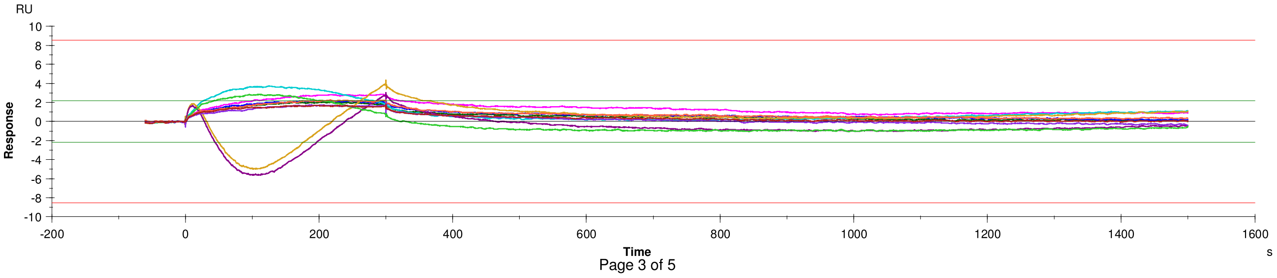

#### Kinetics: ‘140’, fit: ‘1. 1:1 Binding’ (continued)

**Quality Control**

| | |
|---|---|
| ✓ | Kinetic constants are within instrument specifications. |
| ✓ | Kinetic constants appear to be uniquely determined. |
| ⇨ | Bulk contributions (RI) were not evaluated. The RI parameter is set to constant. |
| ⇨ | Check that sensorgrams have sufficient curvature. |
| ⇨ | Examine the residual plot. Pay attention to systematic and non-random deviations. |

#### Kinetics: '140', fit: '1. 1:1 Binding' (continued)

##### Report table

| Curve | ka (1/Ms) | kd (1/s) | KD (M) | Rmax (RU) | Conc (M) | tc | Flow (ul/min) | kt (RU/Ms) | RI (RU) | Chi² (RU²) | U-value |
|---|---|---|---|---|---|---|---|---|---|---|---|
| | 1.619E+5 | 1.693E-4 | 1.046E-9 | 108.5 | | 6.769E+18 | | | | 1.47 | 2 |
| Cycle: 59 0.01 nM | | | | | 1.000E-11 | | 30.00 | 2.103E+19 | 0.000 | | |
| Cycle: 60 0.1 nM | | | | | 1.000E-10 | | 30.00 | 2.103E+19 | 0.000 | | |
| Cycle: 61 1 nM | | | | | 1.000E-9 | | 30.00 | 2.103E+19 | 0.000 | | |
| Cycle: 62 10 nM | | | | | 1.000E-8 | | 30.00 | 2.103E+19 | 0.000 | | |
| Cycle: 63 100 nM | | | | | 1.000E-7 | | 30.00 | 2.103E+19 | 0.000 | | |
| Cycle: 64 100 nM | | | | | 1.000E-7 | | 30.00 | 2.103E+19 | 0.000 | | |
| Cycle: 65 10 nM | | | | | 1.000E-8 | | 30.00 | 2.103E+19 | 0.000 | | |
| Cycle: 66 1 nM | | | | | 1.000E-9 | | 30.00 | 2.103E+19 | 0.000 | | |
| Cycle: 67 0.1 nM | | | | | 1.000E-10 | | 30.00 | 2.103E+19 | 0.000 | | |
| Cycle: 68 0.01 nM | | | | | 1.000E-11 | | 30.00 | 2.103E+19 | 0.000 | | |

##### Parameters table

| Curve | ka (1/Ms) | SE(ka) | kd (1/s) | SE(kd) | Rmax (RU) | SE(Rmax) | Conc (M) | tc | SE(tc) | f (ul/min) | RI (RU) |
|---|---|---|---|---|---|---|---|---|---|---|---|
| | 1.619E+5 | 45 | 1.693E-4 | 1.8E-7 | 108.5 | 0.014 | | 6.769E+18 | 5.6E+21 | | |
| Cycle: 59 0.01 nM | | | | | | | 1E-11 | | | 30 | 0.0 |
| Cycle: 60 0.1 nM | | | | | | | 1E-10 | | | 30 | 0.0 |
| Cycle: 61 1 nM | | | | | | | 1E-09 | | | 30 | 0.0 |
| Cycle: 62 10 nM | | | | | | | 1E-08 | | | 30 | 0.0 |
| Cycle: 63 100 nM | | | | | | | 1E-07 | | | 30 | 0.0 |
| Cycle: 64 100 nM | | | | | | | 1E-07 | | | 30 | 0.0 |
| Cycle: 65 10 nM | | | | | | | 1E-08 | | | 30 | 0.0 |
| Cycle: 66 1 nM | | | | | | | 1E-09 | | | 30 | 0.0 |
| Cycle: 67 0.1 nM | | | | | | | 1E-10 | | | 30 | 0.0 |
| Cycle: 68 0.01 nM | | | | | | | 1E-11 | | | 30 | 0.0 |

#### File Properties

**Evaluation File**
Name: N/A

**User Information**
Performed By: Administrator
Current User: Administrator

**Created With Software**
Name: Biacore T200 Evaluation Software
Version: 1.0

**Notebook**

**Result**

**Result File**
Name: 20231027-GFP-mutants-analyte-1-5_aGFP-S54C-RSGGH_FC2-1.blr
Path: C:\Bia Users\Vaidehi vr358
Size: 24 225 280 bytes

**Run Information**
Type: Kinetics/Affinity
Cycles: 70
Start: 10/27/2023 1:54:59 PM
End: 10/29/2023 5:29:48 AM

**Instrument**
Instrument Type: BiacoreT200
Instrument Id: 1646191
IFC: TYPE105

**User Information**
Run Performed By: Administrator

**Created With Software**
Name: Biacore T200 Control Software
Version: 1.0

**Chip Information**
Chip Id: 231023 Vaidehi CM5
Chip Lot No: 10345041
Chip Name: CM5
First Dock Date: 10/23/2023 11:57:59 PM
Last Modification Date: 10/23/2023 11:57:59 PM
Last Use Date: 10/23/2023 11:59:23 PM

**Immobilization in Fc=1**
Immobilization Date:
Immobilization Result:
Ligand:
Final Response [RU]:

**Immobilization in Fc=2**
Immobilization Date:
Immobilization Result:
Ligand:
Final Response [RU]:

**Immobilization in Fc=3**
Immobilization Date:
Immobilization Result:
Ligand:
Final Response [RU]:

**Immobilization in Fc=4**
Immobilization Date:

#### File Properties (continued)

Immobilization Result:
Ligand:
Final Response [RU]:

**Notebook**

**Kinetics: ‘162’, fit: ‘1. 1:1 Binding’**

Curve: Fc=2-1 Ligand: N/A Sample: 162 Temp: 25°C

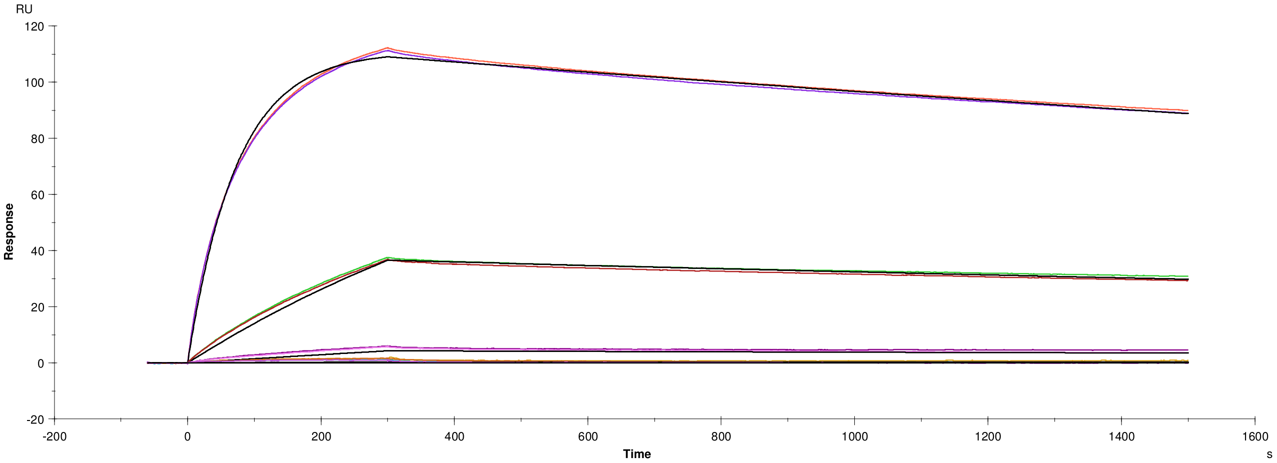

**Residuals**

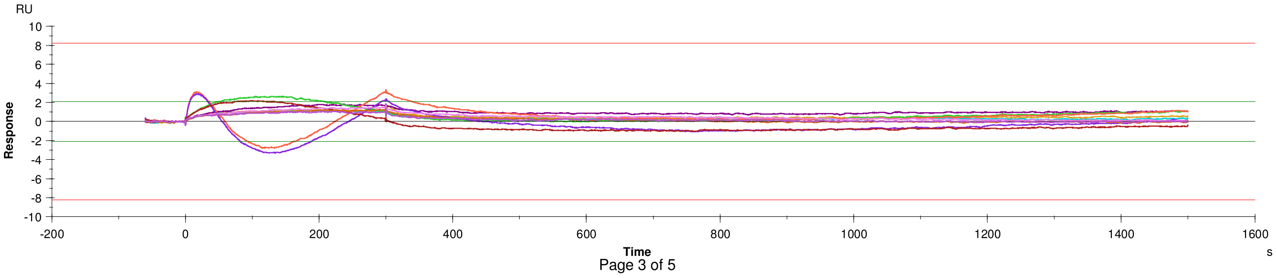

#### Kinetics: ‘162’, fit: ‘1. 1:1 Binding’ (continued)

**Quality Control**

| | |
|---|---|
| ✓ | Kinetic constants are within instrument specifications. |
| ✓ | Kinetic constants appear to be uniquely determined. |
| ➡ | Bulk contributions (RI) were not evaluated. The RI parameter is set to constant. |
| ➡ | Check that sensorgrams have sufficient curvature. |
| ➡ | Examine the residual plot. Pay attention to systematic and non-random deviations. |

#### Kinetics: '162', fit: '1. 1:1 Binding' (continued)

##### Report table

| Curve | ka (1/Ms) | kd (1/s) | KD (M) | Rmax (RU) | Conc (M) | tc | Flow (ul/min) | kt (RU/Ms) | RI (RU) | Chi² (RU²) | U-value |
|---|---|---|---|---|---|---|---|---|---|---|---|
| | 1.354E+5 | 1.709E-4 | 1.263E-9 | 112.3 | | 6.354E+14 | | | | 0.705 | 2 |
| Cycle: 31 0.01 nM | | | | | 1.000E-11 | | 30.00 | 1.974E+15 | 0.000 | | |
| Cycle: 32 0.1 nM | | | | | 1.000E-10 | | 30.00 | 1.974E+15 | 0.000 | | |
| Cycle: 33 1 nM | | | | | 1.000E-9 | | 30.00 | 1.974E+15 | 0.000 | | |
| Cycle: 34 10 nM | | | | | 1.000E-8 | | 30.00 | 1.974E+15 | 0.000 | | |
| Cycle: 35 100 nM | | | | | 1.000E-7 | | 30.00 | 1.974E+15 | 0.000 | | |
| Cycle: 36 100 nM | | | | | 1.000E-7 | | 30.00 | 1.974E+15 | 0.000 | | |
| Cycle: 37 10 nM | | | | | 1.000E-8 | | 30.00 | 1.974E+15 | 0.000 | | |
| Cycle: 38 1 nM | | | | | 1.000E-9 | | 30.00 | 1.974E+15 | 0.000 | | |
| Cycle: 39 0.1 nM | | | | | 1.000E-10 | | 30.00 | 1.974E+15 | 0.000 | | |
| Cycle: 40 0.01 nM | | | | | 1.000E-11 | | 30.00 | 1.974E+15 | 0.000 | | |

##### Parameters table

| Curve | ka (1/Ms) | SE(ka) | kd (1/s) | SE(kd) | Rmax (RU) | SE(Rmax) | Conc (M) | tc | SE(tc) | f (ul/min) | RI (RU) |
|---|---|---|---|---|---|---|---|---|---|---|---|
| | 1.354E+5 | 28 | 1.709E-4 | 1.3E-7 | 112.3 | 0.010 | | 6.354E+14 | 1.4E+13 | | |
| Cycle: 31 0.01 nM | | | | | | | 1E-11 | | | 30 | 0.0 |
| Cycle: 32 0.1 nM | | | | | | | 1E-10 | | | 30 | 0.0 |
| Cycle: 33 1 nM | | | | | | | 1E-09 | | | 30 | 0.0 |
| Cycle: 34 10 nM | | | | | | | 1E-08 | | | 30 | 0.0 |
| Cycle: 35 100 nM | | | | | | | 1E-07 | | | 30 | 0.0 |
| Cycle: 36 100 nM | | | | | | | 1E-07 | | | 30 | 0.0 |
| Cycle: 37 10 nM | | | | | | | 1E-08 | | | 30 | 0.0 |
| Cycle: 38 1 nM | | | | | | | 1E-09 | | | 30 | 0.0 |
| Cycle: 39 0.1 nM | | | | | | | 1E-10 | | | 30 | 0.0 |
| Cycle: 40 0.01 nM | | | | | | | 1E-11 | | | 30 | 0.0 |

#### File Properties

**Evaluation File**

| | |
|---|---|
| Name: | N/A |

**User Information**

| | |
|---|---|
| Performed By: | Administrator |
| Current User: | Administrator |

**Created With Software**

| | |
|---|---|
| Name: | Biacore T200 Evaluation Software |
| Version: | 1.0 |

**Notebook**

**Result**

**Result File**

| | |
|---|---|
| Name: | 20231027-GFP-mutants-analyte-1-5_aGFP-S54C-RSGGH_FC2-1.blr |
| Path: | C:\Bia Users\Vaidehi vr358 |
| Size: | 24 225 280 bytes |

**Run Information**

| | |
|---|---|
| Type: | Kinetics/Affinity |
| Cycles: | 70 |
| Start: | 10/27/2023 1:54:59 PM |
| End: | 10/29/2023 5:29:48 AM |

**Instrument**

| | |
|---|---|
| Instrument Type: | BiacoreT200 |
| Instrument Id: | 1646191 |
| IFC: | TYPE105 |

**User Information**

| | |
|---|---|
| Run Performed By: | Administrator |

**Created With Software**

| | |
|---|---|
| Name: | Biacore T200 Control Software |
| Version: | 1.0 |

**Chip Information**

| | |
|---|---|
| Chip Id: | 231023 Vaidehi CM5 |
| Chip Lot No: | 10345041 |
| Chip Name: | CM5 |
| First Dock Date: | 10/23/2023 11:57:59 PM |
| Last Modification Date: | 10/23/2023 11:57:59 PM |
| Last Use Date: | 10/23/2023 11:59:23 PM |

**Immobilization in Fc=1**
Immobilization Date:
Immobilization Result:
Ligand:
Final Response [RU]:

**Immobilization in Fc=2**
Immobilization Date:
Immobilization Result:
Ligand:
Final Response [RU]:

**Immobilization in Fc=3**
Immobilization Date:
Immobilization Result:
Ligand:
Final Response [RU]:

**Immobilization in Fc=4**
Immobilization Date:

#### File Properties (continued)

Immobilization Result:
Ligand:
Final Response [RU]:

**Notebook**

Unsaved file

Instr. id: 1646191 Biacore T200 Evaluation Software, version 1.0 Print date: 10/30/2023 1:52:32 AM

**Kinetics: ‘166’, fit: ‘1. 1:1 Binding’**

Curve: Fc=2-1 Ligand: N/A Sample: 166 Temp: 25°C

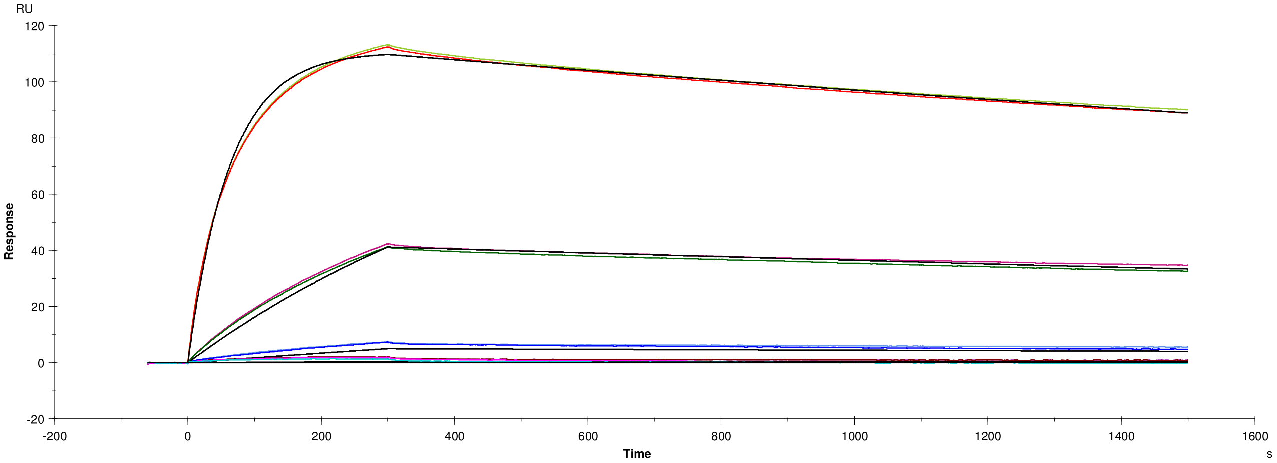

**Residuals**

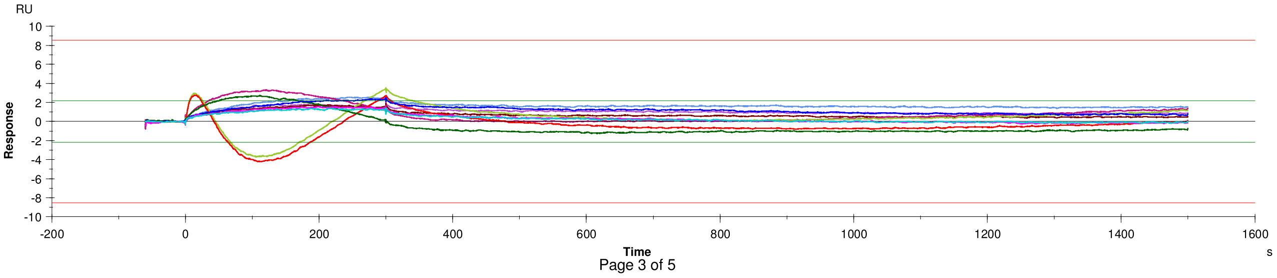

#### Kinetics: ‘166’, fit: ‘1. 1:1 Binding’ (continued)

**Quality Control**

| | |
|---|---|
| ✓ | Kinetic constants are within instrument specifications. |
| ✓ | Kinetic constants appear to be uniquely determined. |
| ➡ | Bulk contributions (RI) were not evaluated. The RI parameter is set to constant. |
| ➡ | Check that sensorgrams have sufficient curvature. |
| ➡ | Examine the residual plot. Pay attention to systematic and non-random deviations. |

#### Kinetics: ‘166’, fit: ‘1. 1:1 Binding’ (continued)

**Report table**

| Curve | ka (1/Ms) | kd (1/s) | KD (M) | Rmax (RU) | Conc (M) | tc | Flow (ul/min) | kt (RU/Ms) | RI (RU) | Chi² (RU²) | U-value |
|---|---|---|---|---|---|---|---|---|---|---|---|
| | 1.579E+5 | 1.753E-4 | 1.110E-9 | 112.0 | | 3.735E+18 | | | | 1.32 | 2 |
| Cycle: 45 0.01 nM | | | | | 1.000E-11 | | 30.00 | 1.161E+19 | 0.000 | | |
| Cycle: 46 0.1 nM | | | | | 1.000E-10 | | 30.00 | 1.161E+19 | 0.000 | | |
| Cycle: 47 1 nM | | | | | 1.000E-9 | | 30.00 | 1.161E+19 | 0.000 | | |
| Cycle: 48 10 nM | | | | | 1.000E-8 | | 30.00 | 1.161E+19 | 0.000 | | |
| Cycle: 49 100 nM | | | | | 1.000E-7 | | 30.00 | 1.161E+19 | 0.000 | | |
| Cycle: 50 100 nM | | | | | 1.000E-7 | | 30.00 | 1.161E+19 | 0.000 | | |
| Cycle: 51 10 nM | | | | | 1.000E-8 | | 30.00 | 1.161E+19 | 0.000 | | |
| Cycle: 52 1 nM | | | | | 1.000E-9 | | 30.00 | 1.161E+19 | 0.000 | | |
| Cycle: 53 0.1 nM | | | | | 1.000E-10 | | 30.00 | 1.161E+19 | 0.000 | | |
| Cycle: 54 0.01 nM | | | | | 1.000E-11 | | 30.00 | 1.161E+19 | 0.000 | | |

**Parameters table**

| Curve | ka (1/Ms) | SE(ka) | kd (1/s) | SE(kd) | Rmax (RU) | SE(Rmax) | Conc (M) | tc | SE(tc) | f (ul/min) | RI (RU) |
|---|---|---|---|---|---|---|---|---|---|---|---|
| | 1.579E+5 | 41 | 1.753E-4 | 1.7E-7 | 112.0 | 0.013 | | 3.735E+18 | 3.4E+21 | | |
| Cycle: 45 0.01 nM | | | | | | | 1E-11 | | | 30 | 0.0 |
| Cycle: 46 0.1 nM | | | | | | | 1E-10 | | | 30 | 0.0 |
| Cycle: 47 1 nM | | | | | | | 1E-09 | | | 30 | 0.0 |
| Cycle: 48 10 nM | | | | | | | 1E-08 | | | 30 | 0.0 |
| Cycle: 49 100 nM | | | | | | | 1E-07 | | | 30 | 0.0 |
| Cycle: 50 100 nM | | | | | | | 1E-07 | | | 30 | 0.0 |
| Cycle: 51 10 nM | | | | | | | 1E-08 | | | 30 | 0.0 |
| Cycle: 52 1 nM | | | | | | | 1E-09 | | | 30 | 0.0 |
| Cycle: 53 0.1 nM | | | | | | | 1E-10 | | | 30 | 0.0 |
| Cycle: 54 0.01 nM | | | | | | | 1E-11 | | | 30 | 0.0 |

### Image Report: admin 2023-03-11 22h39m23s analysed

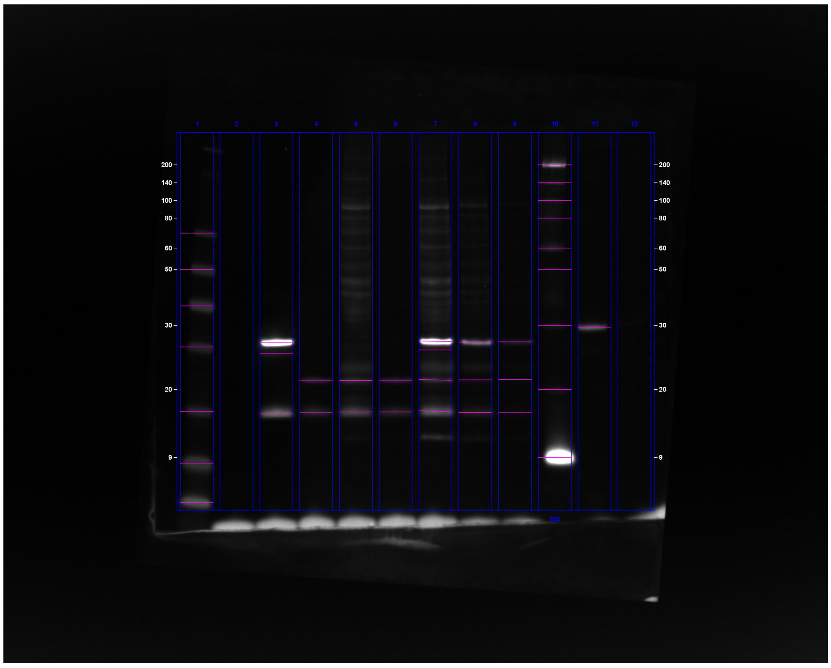

**C:\Users\oded\OneDrive - University Of Cambridge\Vendruscolo\Results\Western\ChemiDoc Images 2023-03-11_22.57.16\admin 2023-03-11 22h39m23s analysed.scn**

---

#### Acquisition Information

| | |
|---|---|
| Imager | ChemiDoc™ MP |
| Exposure Time (sec) | 0.050 (Manual) |
| Flat Field | Applied (Red Epi) |
| Serial Number | 734BR-3787 |
| Software Version | 2.4.0.03 |
| Application | Alexa 647 |
| Excitation Source | Red Epi Illumination |
| Emission Filter | 700/50 Filter |

#### Image Information

| | |
|---|---|
| Acquisition Date | 11/03/2023 10:39:23 PM |
| User Name | admin |
| Image Area (mm) | X: 129.0 Y: 103.2 |
| Pixel Size (µm) | X: 46.9 Y: 46.9 |
| Data Range (Int) | 370 - 20506 |

#### Analysis Settings

| | |
|---|---|
| Detection | Lane detection:<br>Manually created lanes |

| | |
|---|---|
| | Band detection:<br>Automatically detected bands with custom sensitivity: 70<br>Manually adjusted bands<br><br>Lane Background Subtraction:<br>Lane background subtracted with disk size: 17<br><br>Lane width: 5.16 mm |
| Mol. Weight Analysis | Standard: BiotinMarker 7227<br>Standard lanes: 10<br>Regression method: Point to Point (semi-log) |

#### Lane Statistics

| Lane No. | Adj. Total Band Vol. (Int) | Total Band Vol. (Int) | Adj. Total Lane Vol. (Int) | Total Lane Vol. (Int) | Bkgd. Vol. (Int) | Norm. Factor |
|---|---|---|---|---|---|---|
| 1 | 17,781,390 | 38,119,070 | 20,854,790 | 97,962,920 | 77,108,130 | N/A |
| 2 | N/A | N/A | 898,810 | 70,482,280 | 69,583,470 | N/A |
| 3 | 31,331,300 | 38,048,010 | 35,240,260 | 107,671,410 | 72,431,150 | N/A |
| 4 | 2,727,890 | 6,681,070 | 5,022,050 | 83,554,460 | 78,532,410 | N/A |
| 5 | 4,674,230 | 9,063,890 | 16,165,710 | 113,572,250 | 97,406,540 | N/A |
| 6 | 2,429,900 | 5,564,020 | 4,737,040 | 86,070,710 | 81,333,670 | N/A |
| 7 | 24,322,100 | 35,600,840 | 38,091,790 | 138,959,700 | 100,867,910 | N/A |
| 8 | 5,952,210 | 13,581,590 | 10,631,610 | 96,866,990 | 86,235,380 | N/A |
| 9 | 1,455,080 | 8,770,850 | 3,196,380 | 83,310,810 | 80,114,430 | N/A |
| 10 | 53,907,040 | 73,667,660 | 58,727,790 | 144,760,550 | 86,032,760 | N/A |
| 11 | 3,065,480 | 5,109,610 | 4,723,730 | 82,499,230 | 77,775,500 | N/A |
| 12 | N/A | N/A | 1,604,020 | 76,785,500 | 75,181,480 | N/A |

#### Lane And Band Analysis

##### Lane 1

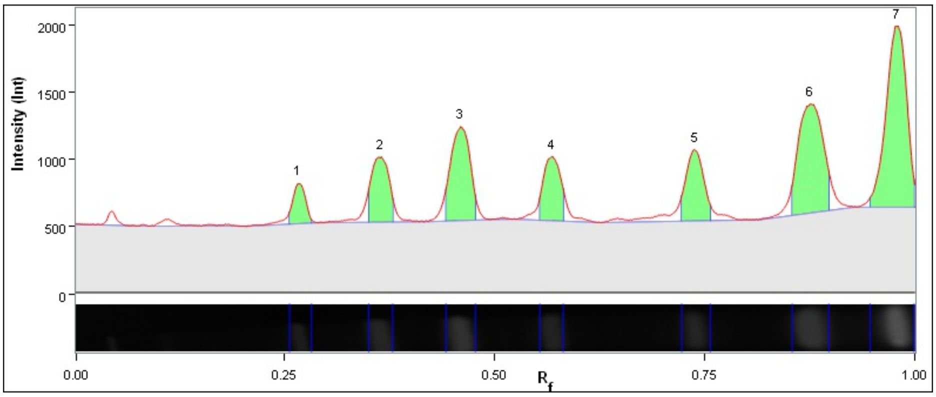

| Band No. | Band Label | Mol. Wt. (KDa) | Relative Front | Adj. Volume (Int) | Volume (Int) | Abs. Quant. | Rel. Quant. | Band % | Lane % |
|---|---|---|---|---|---|---|---|---|---|
| 1 | | 69.3 | 0.266 | 734,690 | 2,727,120 | N/A | N/A | 4.1 | 3.5 |
| 2 | | 49.9 | 0.363 | 1,654,620 | 3,865,290 | N/A | N/A | 9.3 | 7.9 |
| 3 | | 35.8 | 0.459 | 2,596,330 | 5,284,730 | N/A | N/A | 14.6 | 12.4 |
| 4 | | 26.2 | 0.568 | 1,555,840 | 3,751,660 | N/A | N/A | 8.7 | 7.5 |
| 5 | | 15.5 | 0.738 | 1,816,760 | 4,493,500 | N/A | N/A | 10.2 | 8.7 |
| 6 | | 9.0 | 0.875 | 3,894,440 | 7,611,230 | N/A | N/A | 21.9 | 18.7 |
| 7 | | 9.0 | 0.979 | 5,528,710 | 10,385,540 | N/A | N/A | 31.1 | 26.5 |

| | |
|---|---|
| Band Detection | Automatically detected bands with custom sensitivity: 70 |
| Lane Background | Lane background subtracted with disk size: 17 |
| Lane Width | 5.16 mm |
| Regression Equation | A single equation is not available for this method |

##### Lane 2

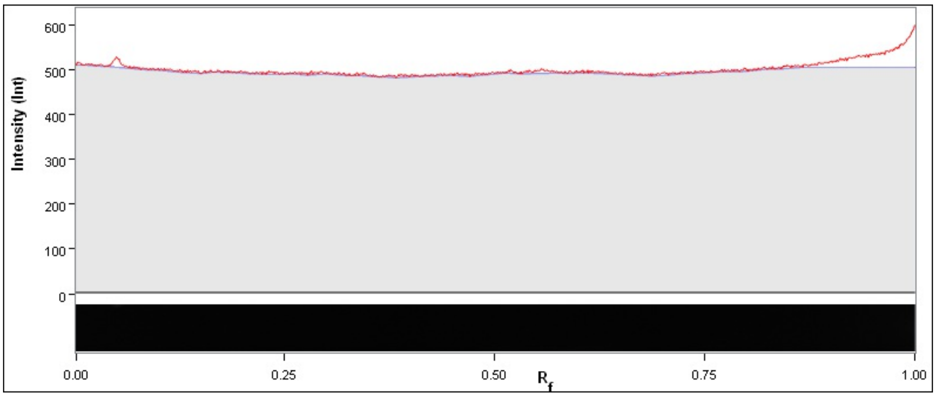

| Band No. | Band Label | Mol. Wt. (KDa) | Relative Front | Adj. Volume (Int) | Volume (Int) | Abs. Quant. | Rel. Quant. | Band % | Lane % |
|---|---|---|---|---|---|---|---|---|---|

| | |
|---|---|
| Band Detection | Automatically detected bands with custom sensitivity: 70 |
| Lane Background | Lane background subtracted with disk size: 17 |
| Lane Width | 5.16 mm |
| Regression Equation | A single equation is not available for this method |

##### Lane 3

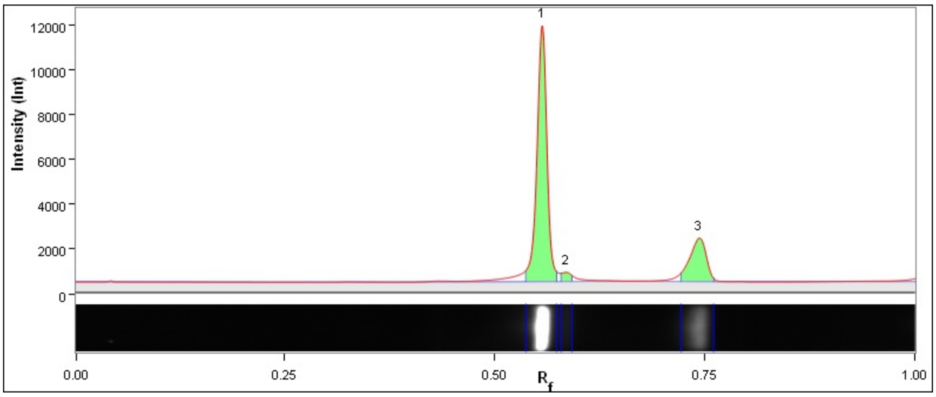

| Band No. | Band Label | Mol. Wt. (KDa) | Relative Front | Adj. Volume (Int) | Volume (Int) | Abs. Quant. | Rel. Quant. | Band % | Lane % |
|---|---|---|---|---|---|---|---|---|---|
| 1 | | 26.9 | 0.557 | 23,927,750 | 26,688,750 | N/A | N/A | 76.4 | 67.9 |
| 2 | | 25.2 | 0.584 | 743,050 | 1,752,520 | N/A | N/A | 2.4 | 2.1 |
| 3 | | 15.1 | 0.743 | 6,660,500 | 9,606,740 | N/A | N/A | 21.3 | 18.9 |

| | |
|---|---|
| Band Detection | Automatically detected bands with custom sensitivity: 70 |

| | |
|---|---|
| Lane Background | Lane background subtracted with disk size: 17 |
| Lane Width | 5.16 mm |
| Regression Equation | A single equation is not available for this method |

##### Lane 4

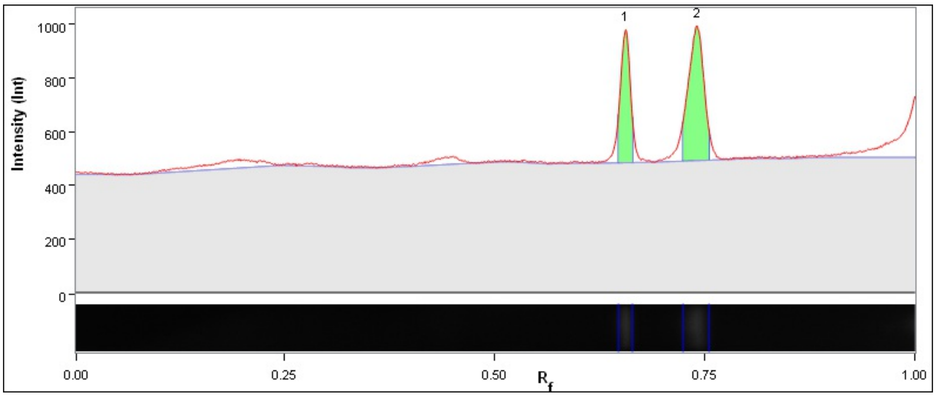

| Band No. | Band Label | Mol. Wt. (KDa) | Relative Front | Adj. Volume (Int) | Volume (Int) | Abs. Quant. | Rel. Quant. | Band % | Lane % |
|---|---|---|---|---|---|---|---|---|---|
| 1 | | 21.2 | 0.656 | 1,015,190 | 2,403,500 | N/A | N/A | 37.2 | 20.2 |
| 2 | | 15.3 | 0.741 | 1,712,700 | 4,277,570 | N/A | N/A | 62.8 | 34.1 |

| | |
|---|---|
| Band Detection | Automatically detected bands with custom sensitivity: 70 |
| Lane Background | Lane background subtracted with disk size: 17 |
| Lane Width | 5.16 mm |
| Regression Equation | A single equation is not available for this method |

##### Lane 5

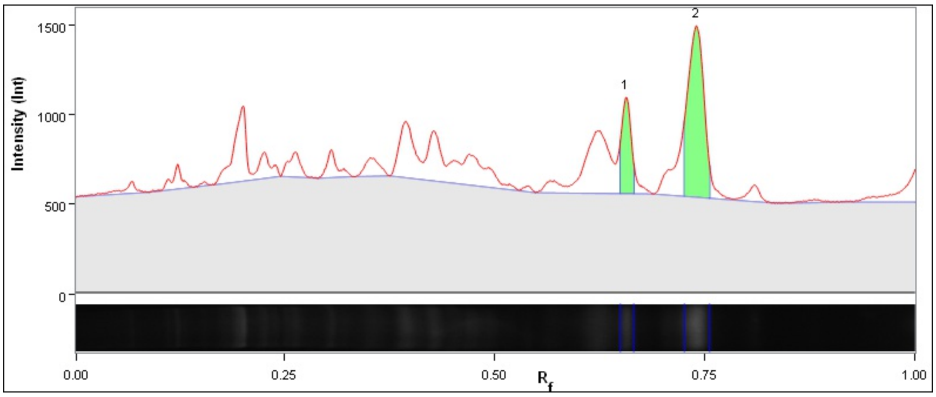

| Band No. | Band Label | Mol. Wt. (KDa) | Relative Front | Adj. Volume (Int) | Volume (Int) | Abs. Quant. | Rel. Quant. | Band % | Lane % |
|---|---|---|---|---|---|---|---|---|---|
| 1 | | 21.2 | 0.657 | 1,147,960 | 2,721,180 | N/A | N/A | 24.6 | 7.1 |
| 2 | | 15.4 | 0.740 | 3,526,270 | 6,342,710 | N/A | N/A | 75.4 | 21.8 |

| | |
|---|---|
| Band Detection | Automatically detected bands with custom sensitivity: 70 |
| Lane Background | Lane background subtracted with disk size: 17 |

| | |
|---|---|
| Lane Width | 5.16 mm |
| Regression Equation | A single equation is not available for this method |

##### Lane 6

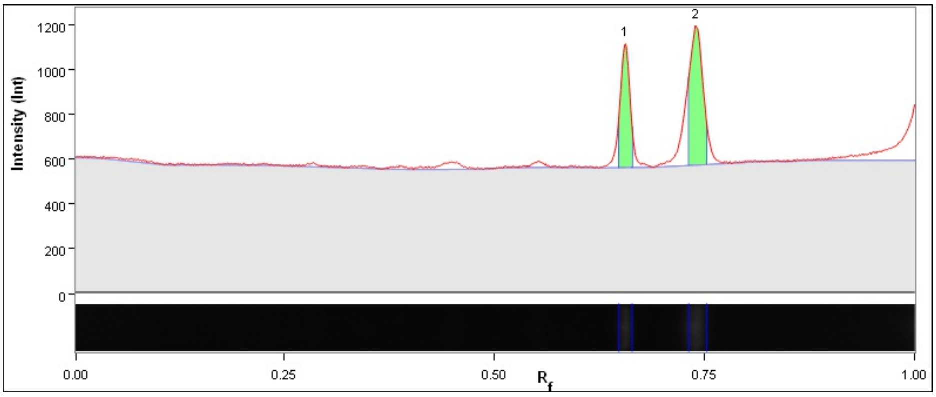

| Band No. | Band Label | Mol. Wt. (KDa) | Relative Front | Adj. Volume (Int) | Volume (Int) | Abs. Quant. | Rel. Quant. | Band % | Lane % |
|---|---|---|---|---|---|---|---|---|---|
| 1 | | 21.2 | 0.656 | 950,290 | 2,277,330 | N/A | N/A | 39.1 | 20.1 |
| 2 | | 15.4 | 0.740 | 1,479,610 | 3,286,690 | N/A | N/A | 60.9 | 31.2 |

| | |
|---|---|
| Band Detection | Automatically detected bands with custom sensitivity: 70 |
| Lane Background | Lane background subtracted with disk size: 17 |
| Lane Width | 5.16 mm |
| Regression Equation | A single equation is not available for this method |

##### Lane 7

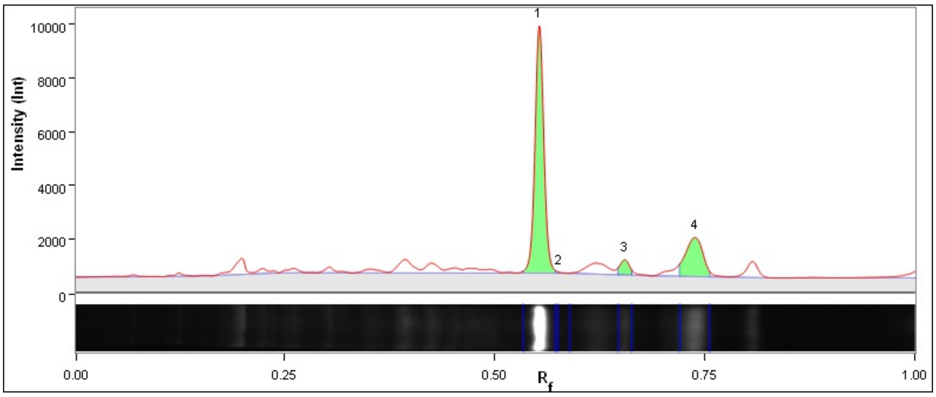

| Band No. | Band Label | Mol. Wt. (KDa) | Relative Front | Adj. Volume (Int) | Volume (Int) | Abs. Quant. | Rel. Quant. | Band % | Lane % |
|---|---|---|---|---|---|---|---|---|---|
| 1 | | 27.1 | 0.553 | 17,788,870 | 22,277,970 | N/A | N/A | 73.1 | 46.7 |
| 2 | | 25.7 | 0.576 | 32,890 | 1,729,970 | N/A | N/A | 0.1 | 0.1 |
| 3 | | 21.3 | 0.655 | 1,105,170 | 2,803,350 | N/A | N/A | 4.5 | 2.9 |
| 4 | | 15.5 | 0.738 | 5,395,170 | 8,789,550 | N/A | N/A | 22.2 | 14.2 |

| | |
|---|---|
| Band Detection | Automatically detected bands with custom sensitivity: 70 |

| | |
|---|---|
| Lane Background | Lane background subtracted with disk size: 17 |
| Lane Width | 5.16 mm |
| Regression Equation | A single equation is not available for this method |

##### Lane 8

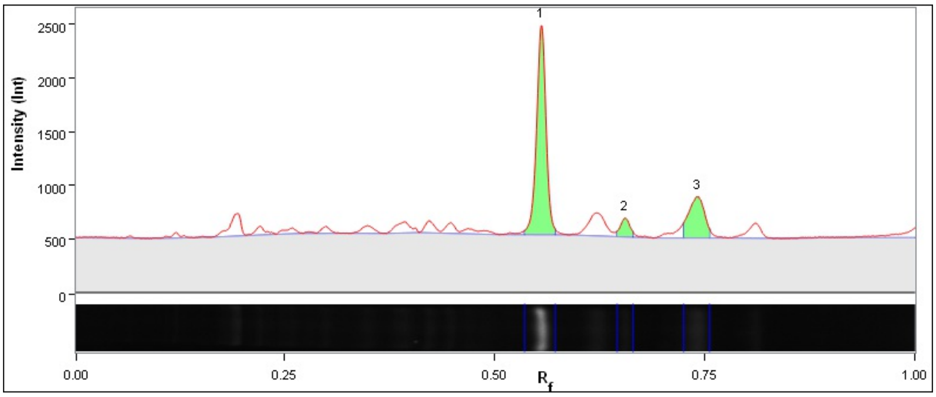

| Band No. | Band Label | Mol. Wt. (KDa) | Relative Front | Adj. Volume (Int) | Volume (Int) | Abs. Quant. | Rel. Quant. | Band % | Lane % |
|---|---|---|---|---|---|---|---|---|---|
| 1 | | 27.0 | 0.555 | 4,221,360 | 7,516,190 | N/A | N/A | 70.9 | 39.7 |
| 2 | | 21.3 | 0.655 | 372,130 | 2,057,880 | N/A | N/A | 6.3 | 3.5 |
| 3 | | 15.3 | 0.741 | 1,358,720 | 4,007,520 | N/A | N/A | 22.8 | 12.8 |

| | |
|---|---|
| Band Detection | Automatically detected bands with custom sensitivity: 70 |
| Lane Background | Lane background subtracted with disk size: 17 |
| Lane Width | 5.16 mm |
| Regression Equation | A single equation is not available for this method |

##### Lane 9

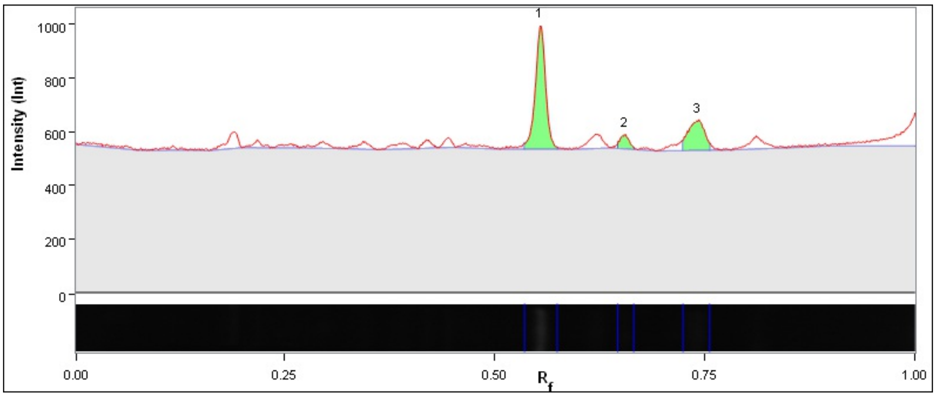

| Band No. | Band Label | Mol. Wt. (KDa) | Relative Front | Adj. Volume (Int) | Volume (Int) | Abs. Quant. | Rel. Quant. | Band % | Lane % |
|---|---|---|---|---|---|---|---|---|---|
| 1 | | 27.0 | 0.554 | 978,780 | 4,141,280 | N/A | N/A | 67.3 | 30.6 |
| 2 | | 21.3 | 0.654 | 97,790 | 1,681,240 | N/A | N/A | 6.7 | 3.1 |
| 3 | | 15.3 | 0.741 | 378,510 | 2,948,330 | N/A | N/A | 26.0 | 11.8 |

| | |
|---|---|
| Band Detection | Automatically detected bands with custom sensitivity: 70 |
| Lane Background | Lane background subtracted with disk size: 17 |
| Lane Width | 5.16 mm |
| Regression Equation | A single equation is not available for this method |

##### Lane 10 - BiotinMarker 7227

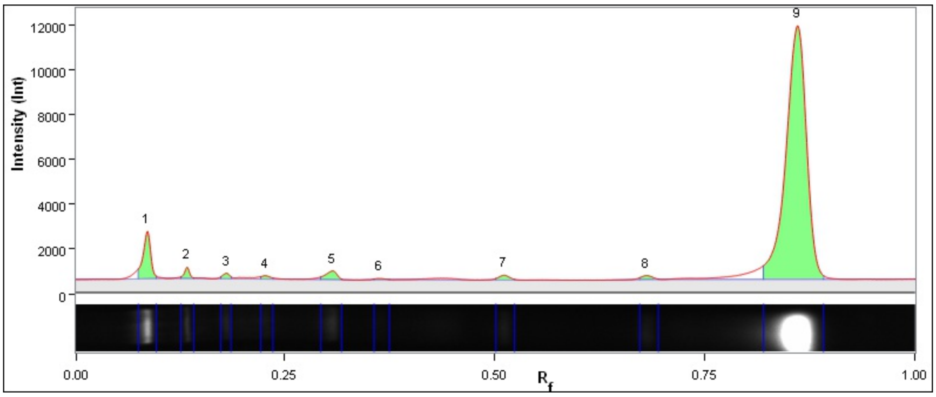

| Band No. | Band Label | Mol. Wt. (KDa) | Relative Front | Adj. Volume (Int) | Volume (Int) | Abs. Quant. | Rel. Quant. | Band % | Lane % |
|---|---|---|---|---|---|---|---|---|---|
| 1 | | 200.0 | 0.086 | 3,260,510 | 5,206,850 | N/A | N/A | 6.0 | 5.6 |
| 2 | | 140.0 | 0.133 | 537,790 | 1,998,150 | N/A | N/A | 1.0 | 0.9 |
| 3 | | 100.0 | 0.180 | 336,600 | 1,483,900 | N/A | N/A | 0.6 | 0.6 |
| 4 | | 80.0 | 0.227 | 253,440 | 1,589,720 | N/A | N/A | 0.5 | 0.4 |
| 5 | | 60.0 | 0.306 | 893,750 | 3,040,180 | N/A | N/A | 1.7 | 1.5 |
| 6 | | 50.0 | 0.362 | 120,670 | 1,717,540 | N/A | N/A | 0.2 | 0.2 |
| 7 | | 30.0 | 0.511 | 464,420 | 2,353,670 | N/A | N/A | 0.9 | 0.8 |
| 8 | | 20.0 | 0.680 | 399,080 | 2,360,930 | N/A | N/A | 0.7 | 0.7 |
| 9 | | 9.0 | 0.860 | 47,640,780 | 53,916,720 | N/A | N/A | 88.4 | 81.1 |

| | |
|---|---|
| Band Detection | Automatically detected bands with custom sensitivity: 70 |
| Lane Background | Lane background subtracted with disk size: 17 |
| Lane Width | 5.16 mm |
| Regression Equation | A single equation is not available for this method |

##### Lane 11

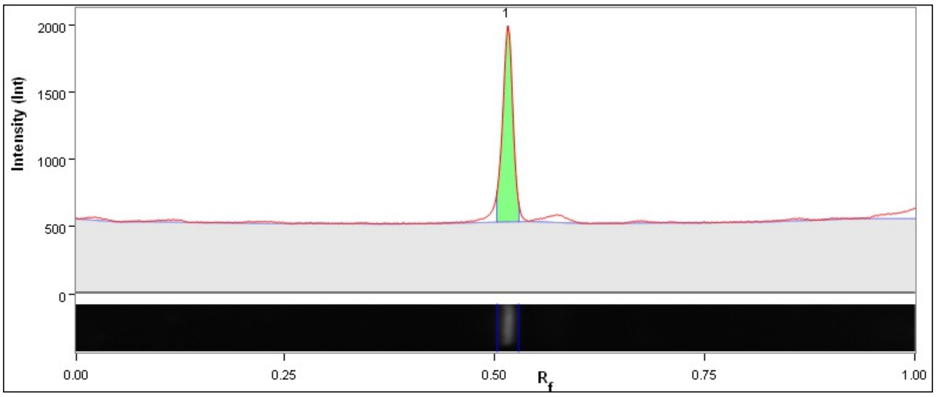

| Band No. | Band Label | Mol. Wt. (KDa) | Relative Front | Adj. Volume (Int) | Volume (Int) | Abs. Quant. | Rel. Quant. | Band % | Lane % |
|---|---|---|---|---|---|---|---|---|---|
| 1 | | 29.7 | 0.515 | 3,065,480 | 5,109,610 | N/A | N/A | 100.0 | 64.9 |

| | |
|---|---|
| Band Detection | Automatically detected bands with custom sensitivity: 70 |
| Lane Background | Lane background subtracted with disk size: 17 |
| Lane Width | 5.16 mm |
| Regression Equation | A single equation is not available for this method |

**Lane 12**

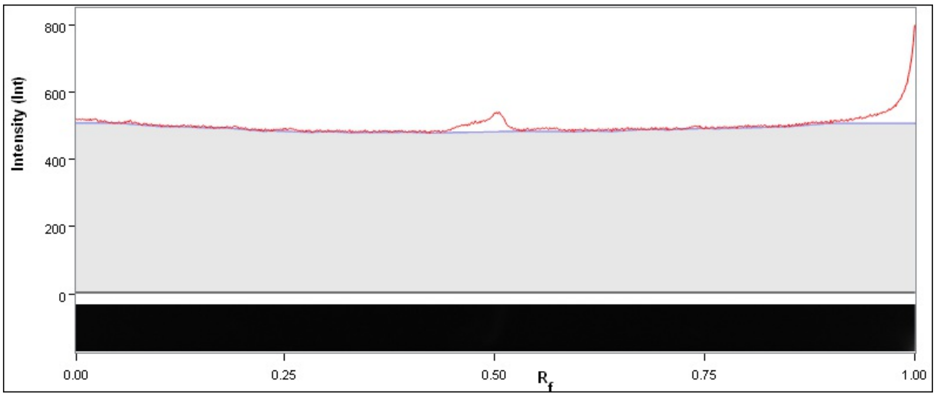

| Band No. | Band Label | Mol. Wt. (KDa) | Relative Front | Adj. Volume (Int) | Volume (Int) | Abs. Quant. | Rel. Quant. | Band % | Lane % |
|---|---|---|---|---|---|---|---|---|---|

| | |
|---|---|
| Band Detection | Automatically detected bands with custom sensitivity: 70 |
| Lane Background | Lane background subtracted with disk size: 17 |
| Lane Width | 5.16 mm |
| Regression Equation | A single equation is not available for this method |

### Image Report: admin 2023-03-25 20h41m30s

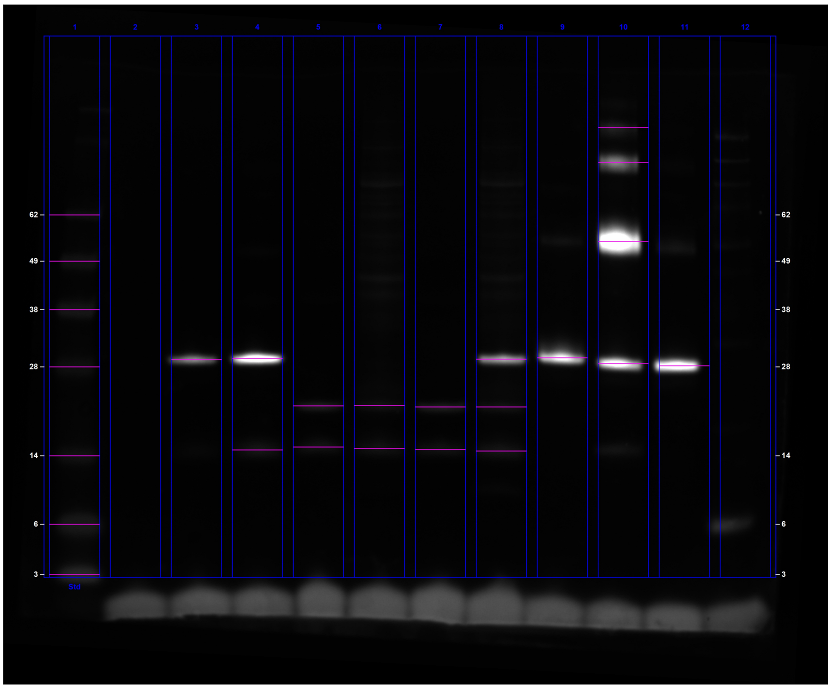

**C:\Users\oded\OneDrive - University Of Cambridge\Vendruscolo\tmp\ChemiDoc Images 2023-03-25_20.48.53\admin 2023-03-25 20h41m30s.scn**

#### Acquisition Information

| | |
|---|---|
| Imager | ChemiDoc™ MP |
| Exposure Time (sec) | 0.050 (Manual) |
| Flat Field | Applied (Red Epi) |
| Serial Number | 734BR-3787 |
| Software Version | 2.4.0.03 |
| Application | Alexa 647 |
| Excitation Source | Red Epi Illumination |
| Emission Filter | 700/50 Filter |

#### Image Information

| | |
|---|---|
| Acquisition Date | 25/03/2023 8:41:30 PM |
| User Name | admin |
| Image Area (mm) | X: 89.3 Y: 73.7 |
| Pixel Size (μm) | X: 39.5 Y: 39.5 |
| Data Range (Int) | 0 - 19859 |

#### Analysis Settings

| | |
|---|---|
| Detection | Lane detection:<br>Manually created lanes |

| | |
|---|---|
| | Band detection:<br>Automatically detected bands with sensitivity: High<br>Manually adjusted bands<br><br>Lane Background Subtraction:<br>Lane background subtracted with disk size: 17<br><br>Lane width: 5.45 mm |
| Mol. Weight Analysis | Standard: SeeBlue2<br>Standard lanes: first<br>Regression method: Point to Point (semi-log) |

#### Lane Statistics

| Lane No. | Adj. Total Band Vol. (Int) | Total Band Vol. (Int) | Adj. Total Lane Vol. (Int) | Total Lane Vol. (Int) | Bkgd. Vol. (Int) | Norm. Factor |
|---|---|---|---|---|---|---|
| 1 | 19,744,764 | 43,434,258 | 27,616,008 | 152,021,214 | 124,405,206 | N/A |
| 2 | N/A | N/A | 1,952,562 | 112,140,042 | 110,187,480 | N/A |
| 3 | 14,245,326 | 17,762,532 | 18,093,318 | 128,832,660 | 110,739,342 | N/A |
| 4 | 45,786,054 | 53,543,310 | 50,559,198 | 162,096,870 | 111,537,672 | N/A |
| 5 | 5,376,342 | 11,211,948 | 9,178,380 | 121,011,096 | 111,832,716 | N/A |
| 6 | 4,415,034 | 10,828,032 | 14,767,242 | 142,598,436 | 127,831,194 | N/A |
| 7 | 5,761,776 | 11,477,874 | 9,198,942 | 122,714,706 | 113,515,764 | N/A |
| 8 | 24,762,030 | 36,460,290 | 34,523,322 | 167,168,094 | 132,644,772 | N/A |
| 9 | 40,702,686 | 46,981,686 | 47,575,500 | 165,631,878 | 118,056,378 | N/A |
| 10 | 159,679,800 | 190,170,762 | 168,640,968 | 294,771,174 | 126,130,206 | N/A |
| 11 | 42,911,238 | 48,692,058 | 58,633,854 | 167,621,424 | 108,987,570 | N/A |
| 12 | N/A | N/A | 16,952,196 | 128,108,850 | 111,156,654 | N/A |

#### Lane And Band Analysis

##### Lane 1 - SeeBlue2

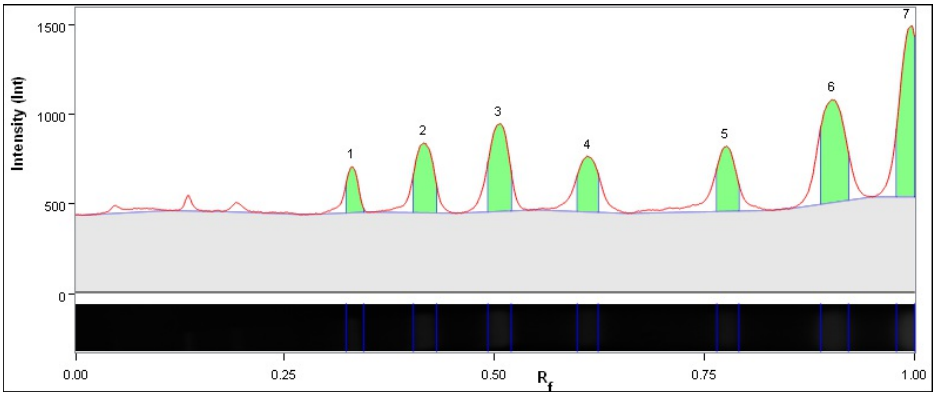

| Band No. | Band Label | Mol. Wt. (KDa) | Relative Front | Adj. Volume (Int) | Volume (Int) | Abs. Quant. | Rel. Quant. | Band % | Lane % |
|---|---|---|---|---|---|---|---|---|---|
| 1 | | 62.0 | 0.330 | 1,002,570 | 3,613,254 | N/A | N/A | 5.1 | 3.6 |
| 2 | | 49.0 | 0.416 | 2,444,118 | 5,862,792 | N/A | N/A | 12.4 | 8.9 |
| 3 | | 38.0 | 0.505 | 3,109,002 | 6,587,844 | N/A | N/A | 15.7 | 11.3 |
| 4 | | 28.0 | 0.611 | 1,912,818 | 5,038,656 | N/A | N/A | 9.7 | 6.9 |
| 5 | | 14.0 | 0.775 | 2,165,082 | 5,482,326 | N/A | N/A | 11.0 | 7.8 |
| 6 | | 6.0 | 0.902 | 4,476,582 | 9,088,266 | N/A | N/A | 22.7 | 16.2 |
| 7 | | 3.0 | 0.995 | 4,634,592 | 7,761,120 | N/A | N/A | 23.5 | 16.8 |

| | |
|---|---|
| Band Detection | Automatically detected bands with sensitivity: High |
| Lane Background | Lane background subtracted with disk size: 17 |
| Lane Width | 5.45 mm |
| Regression Equation | A single equation is not available for this method |

#### Lane 2

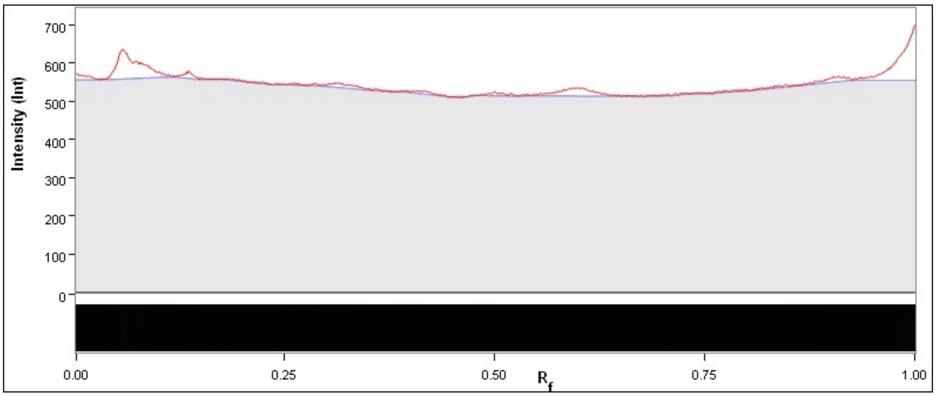

| Band No. | Band Label | Mol. Wt. (KDa) | Relative Front | Adj. Volume (Int) | Volume (Int) | Abs. Quant. | Rel. Quant. | Band % | Lane % |
|---|---|---|---|---|---|---|---|---|---|

| | |
|---|---|
| Band Detection | Automatically detected bands with sensitivity: High |
| Lane Background | Lane background subtracted with disk size: 17 |
| Lane Width | 5.45 mm |
| Regression Equation | A single equation is not available for this method |

#### Lane 3

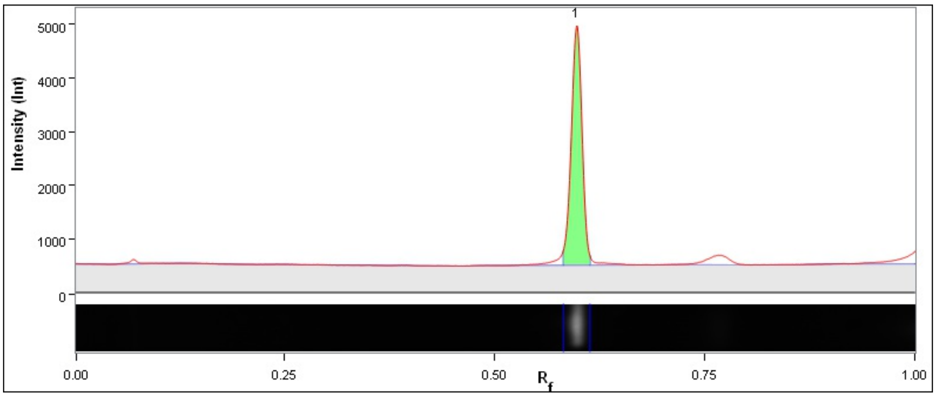

| Band No. | Band Label | Mol. Wt. (KDa) | Relative Front | Adj. Volume (Int) | Volume (Int) | Abs. Quant. | Rel. Quant. | Band % | Lane % |
|---|---|---|---|---|---|---|---|---|---|
| 1 | | 29.1 | 0.598 | 14,245,326 | 17,762,532 | N/A | N/A | 100.0 | 78.7 |

| | |
|---|---|
| Band Detection | Automatically detected bands with sensitivity: High |
| Lane Background | Lane background subtracted with disk size: 17 |
| Lane Width | 5.45 mm |

| | |
|---|---|
| Regression Equation | A single equation is not available for this method |

##### Lane 4

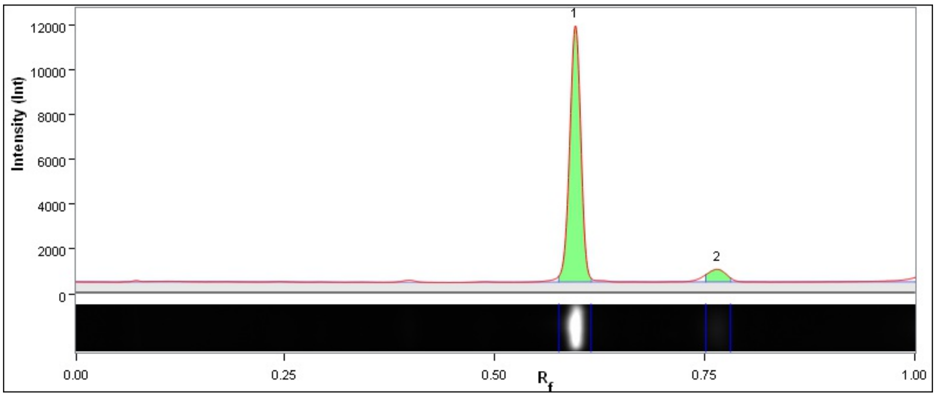

| Band No. | Band Label | Mol. Wt. (KDa) | Relative Front | Adj. Volume (Int) | Volume (Int) | Abs. Quant. | Rel. Quant. | Band % | Lane % |
|---|---|---|---|---|---|---|---|---|---|
| 1 | | 29.3 | 0.596 | 42,628,062 | 46,973,268 | N/A | N/A | 93.1 | 84.3 |
| 2 | | 14.7 | 0.764 | 3,157,992 | 6,570,042 | N/A | N/A | 6.9 | 6.2 |

| | |
|---|---|
| Band Detection | Automatically detected bands with sensitivity: High |
| Lane Background | Lane background subtracted with disk size: 17 |
| Lane Width | 5.45 mm |
| Regression Equation | A single equation is not available for this method |

##### Lane 5

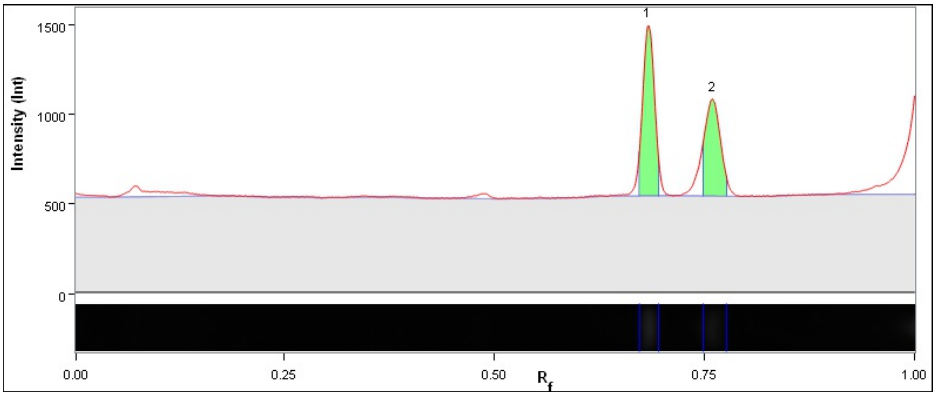

| Band No. | Band Label | Mol. Wt. (KDa) | Relative Front | Adj. Volume (Int) | Volume (Int) | Abs. Quant. | Rel. Quant. | Band % | Lane % |
|---|---|---|---|---|---|---|---|---|---|
| 1 | | 20.7 | 0.683 | 3,127,218 | 5,783,442 | N/A | N/A | 58.2 | 34.1 |
| 2 | | 15.0 | 0.759 | 2,249,124 | 5,428,506 | N/A | N/A | 41.8 | 24.5 |

| | |
|---|---|
| Band Detection | Automatically detected bands with sensitivity: High |
| Lane Background | Lane background subtracted with disk size: 17 |
| Lane Width | 5.45 mm |
| Regression Equation | A single equation is not available for this method |

#### Lane 6

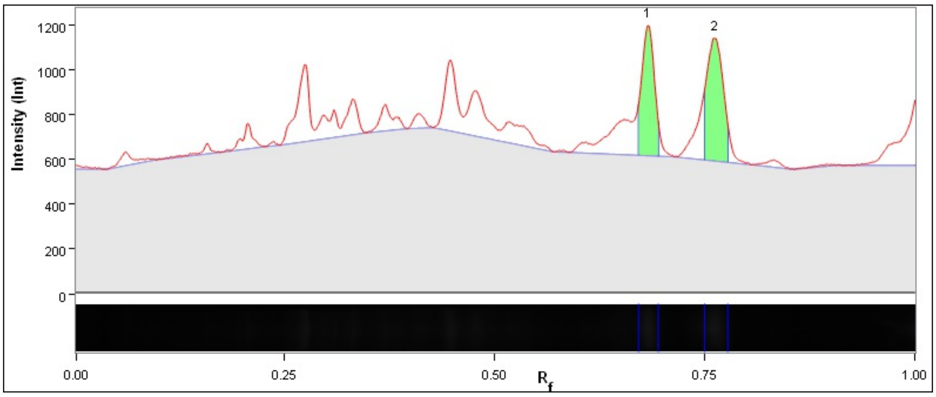

| Band No. | Band Label | Mol. Wt. (KDa) | Relative Front | Adj. Volume (Int) | Volume (Int) | Abs. Quant. | Rel. Quant. | Band % | Lane % |
|---|---|---|---|---|---|---|---|---|---|
| 1 | | 20.7 | 0.682 | 2,008,038 | 5,032,998 | N/A | N/A | 45.5 | 13.6 |
| 2 | | 14.8 | 0.762 | 2,406,996 | 5,795,034 | N/A | N/A | 54.5 | 16.3 |

| | |
|---|---|
| Band Detection | Automatically detected bands with sensitivity: High |
| Lane Background | Lane background subtracted with disk size: 17 |
| Lane Width | 5.45 mm |
| Regression Equation | A single equation is not available for this method |

#### Lane 7

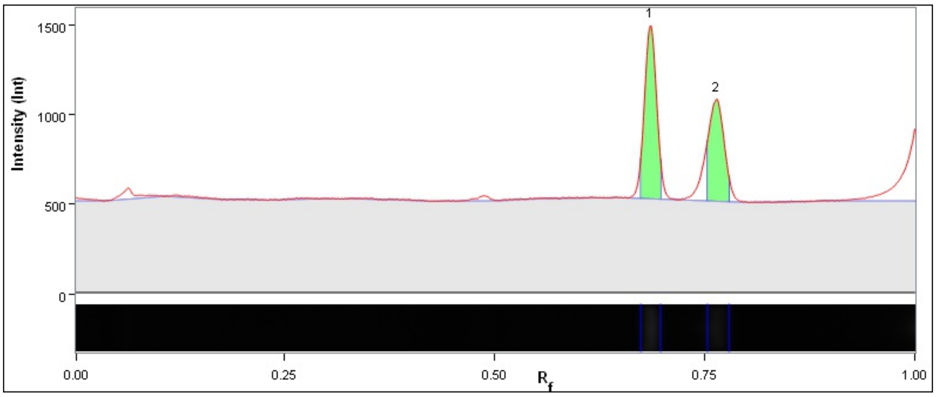

| Band No. | Band Label | Mol. Wt. (KDa) | Relative Front | Adj. Volume (Int) | Volume (Int) | Abs. Quant. | Rel. Quant. | Band % | Lane % |
|---|---|---|---|---|---|---|---|---|---|
| 1 | | 20.5 | 0.685 | 3,370,650 | 6,153,834 | N/A | N/A | 58.5 | 36.6 |
| 2 | | 14.7 | 0.764 | 2,391,126 | 5,324,040 | N/A | N/A | 41.5 | 26.0 |

| | |
|---|---|
| Band Detection | Automatically detected bands with sensitivity: High |
| Lane Background | Lane background subtracted with disk size: 17 |
| Lane Width | 5.45 mm |
| Regression Equation | A single equation is not available for this method |

#### Lane 8

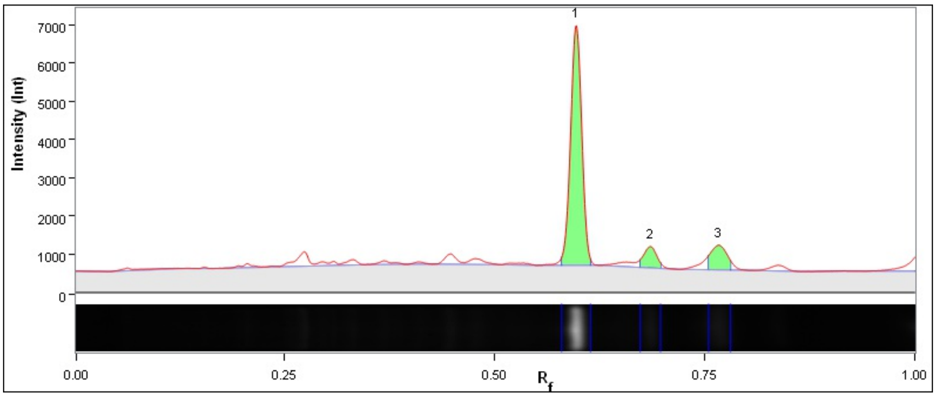

| Band No. | Band Label | Mol. Wt. (KDa) | Relative Front | Adj. Volume (Int) | Volume (Int) | Abs. Quant. | Rel. Quant. | Band % | Lane % |
|---|---|---|---|---|---|---|---|---|---|
| 1 | | 29.2 | 0.597 | 20,008,206 | 25,131,042 | N/A | N/A | 80.8 | 58.0 |
| 2 | | 20.5 | 0.685 | 1,969,950 | 5,267,046 | N/A | N/A | 8.0 | 5.7 |
| 3 | | 14.5 | 0.766 | 2,783,874 | 6,062,202 | N/A | N/A | 11.2 | 8.1 |

| | |
|---|---|
| Band Detection | Automatically detected bands with sensitivity: High |
| Lane Background | Lane background subtracted with disk size: 17 |
| Lane Width | 5.45 mm |
| Regression Equation | A single equation is not available for this method |

##### Lane 9

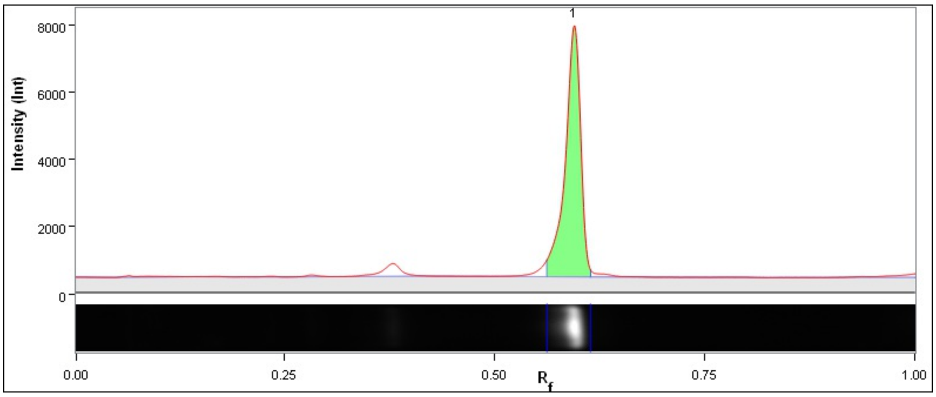

| Band No. | Band Label | Mol. Wt. (KDa) | Relative Front | Adj. Volume (Int) | Volume (Int) | Abs. Quant. | Rel. Quant. | Band % | Lane % |
|---|---|---|---|---|---|---|---|---|---|
| 1 | | 29.4 | 0.594 | 40,702,686 | 46,981,686 | N/A | N/A | 100.0 | 85.6 |

| | |
|---|---|
| Band Detection | Automatically detected bands with sensitivity: High |
| Lane Background | Lane background subtracted with disk size: 17 |
| Lane Width | 5.45 mm |
| Regression Equation | A single equation is not available for this method |

##### Lane 10

6

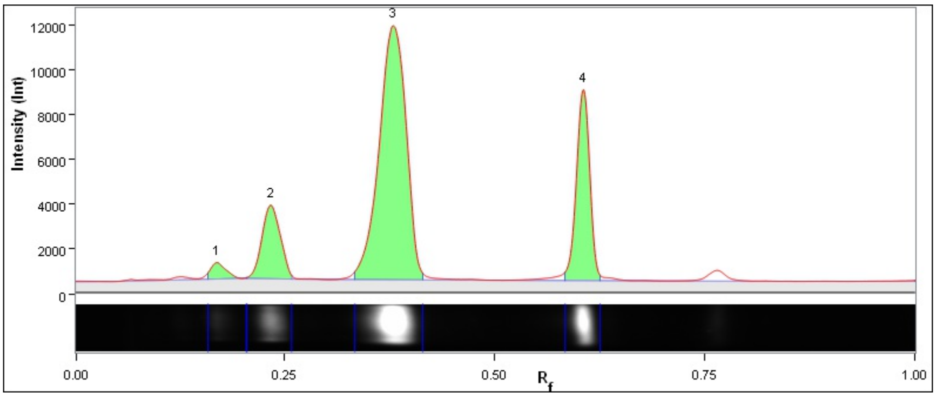

| Band No. | Band Label | Mol. Wt. (KDa) | Relative Front | Adj. Volume (Int) | Volume (Int) | Abs. Quant. | Rel. Quant. | Band % | Lane % |
|---|---|---|---|---|---|---|---|---|---|
| 1 | | 62.0 | 0.169 | 3,572,544 | 10,247,190 | N/A | N/A | 2.2 | 2.1 |
| 2 | | 62.0 | 0.234 | 19,669,278 | 27,406,938 | N/A | N/A | 12.3 | 11.7 |
| 3 | | 54.2 | 0.380 | 99,372,558 | 110,198,106 | N/A | N/A | 62.2 | 58.9 |
| 4 | | 28.5 | 0.605 | 37,065,420 | 42,318,528 | N/A | N/A | 23.2 | 22.0 |

| | |
|---|---|
| Band Detection | Automatically detected bands with sensitivity: High |
| Lane Background | Lane background subtracted with disk size: 17 |
| Lane Width | 5.45 mm |
| Regression Equation | A single equation is not available for this method |

#### Lane 11

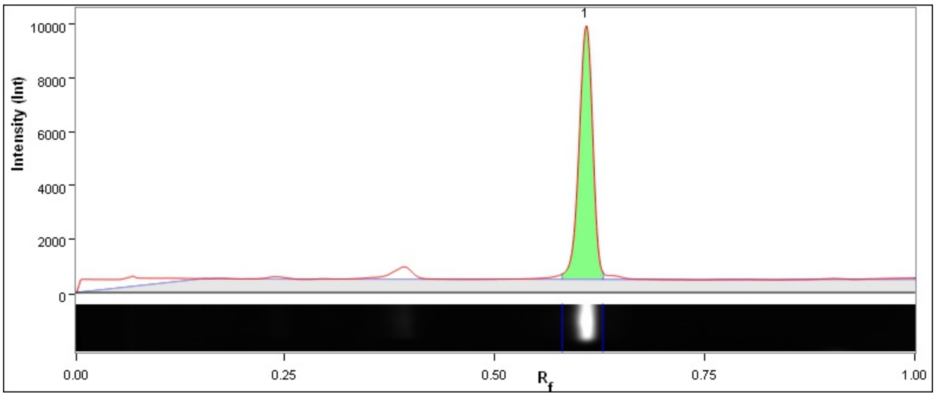

| Band No. | Band Label | Mol. Wt. (KDa) | Relative Front | Adj. Volume (Int) | Volume (Int) | Abs. Quant. | Rel. Quant. | Band % | Lane % |
|---|---|---|---|---|---|---|---|---|---|
| 1 | | 28.2 | 0.609 | 42,911,238 | 48,692,058 | N/A | N/A | 100.0 | 73.2 |

| | |
|---|---|
| Band Detection | Automatically detected bands with sensitivity: High |
| Lane Background | Lane background subtracted with disk size: 17 |
| Lane Width | 5.45 mm |
| Regression Equation | A single equation is not available for this method |

#### Lane 12

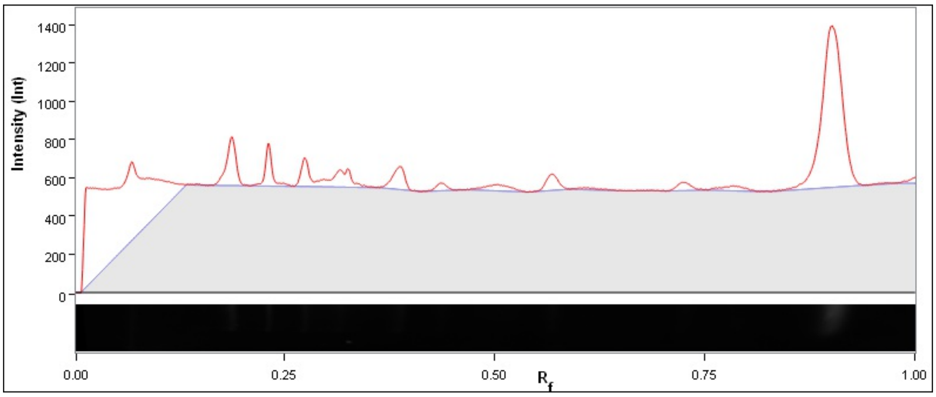

| Band No. | Band Label | Mol. Wt. (KDa) | Relative Front | Adj. Volume (Int) | Volume (Int) | Abs. Quant. | Rel. Quant. | Band % | Lane % |
|---|---|---|---|---|---|---|---|---|---|

| | |
|---|---|
| Band Detection | Automatically detected bands with sensitivity: High |
| Lane Background | Lane background subtracted with disk size: 17 |
| Lane Width | 5.45 mm |
| Regression Equation | A single equation is not available for this method |

8

### Image Report: admin 2023-03-28 21h12m43s

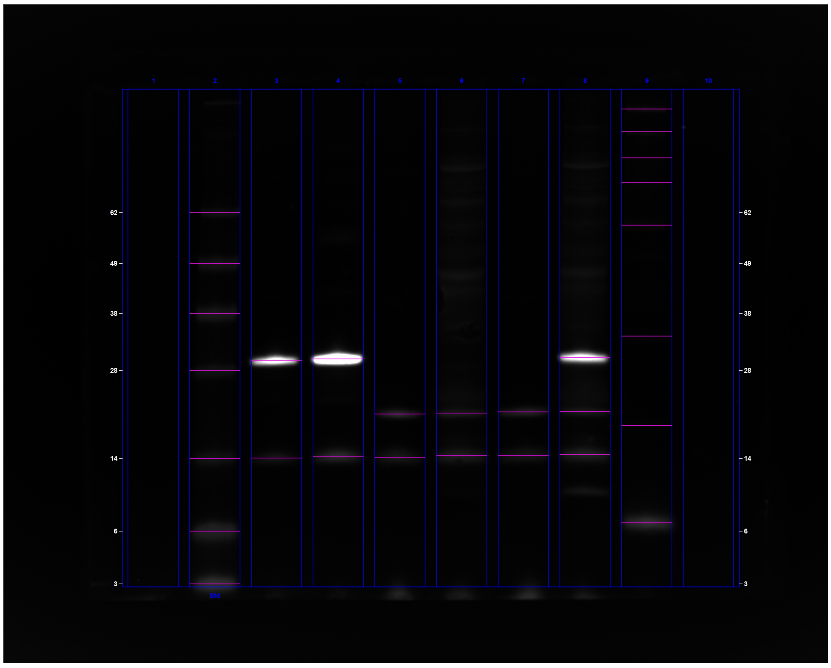

**C:\Users\oded\OneDrive - University Of Cambridge\Vendruscolo\tmp\ChemiDoc Images 2023-03-28_21.19.37\admin 2023-03-28 21h12m43s.scn**

---

#### Acquisition Information

| | |
|---|---|
| Imager | ChemiDoc™ MP |
| Exposure Time (sec) | 0.050 (Manual) |
| Flat Field | Applied (Red Epi) |
| Serial Number | 734BR-3787 |
| Software Version | 2.4.0.03 |
| Application | Alexa 647 |
| Excitation Source | Red Epi Illumination |
| Emission Filter | 700/50 Filter |

#### Image Information

| | |
|---|---|
| Acquisition Date | 28/03/2023 10:12:42 PM |
| User Name | admin |
| Image Area (mm) | X: 96.8 Y: 77.4 |
| Pixel Size (µm) | X: 35.2 Y: 35.2 |
| Data Range (Int) | 401 - 31797 |

#### Analysis Settings

| | |
|---|---|
| Detection | Lane detection:<br>Manually created lanes |

| | |
|---|---|
| | Band detection:<br>Automatically detected bands with custom sensitivity: 77<br>Manually adjusted bands<br><br>Lane Background Subtraction:<br>Lane background subtracted with disk size: 17<br><br>Lane width: 5.91 mm |
| Mol. Weight Analysis | Standard: SeeBlue2<br>Standard lanes: 2<br>Regression method: Point to Point (semi-log) |

#### Lane Statistics

| Lane No. | Adj. Total Band Vol. (Int) | Total Band Vol. (Int) | Adj. Total Lane Vol. (Int) | Total Lane Vol. (Int) | Bkgd. Vol. (Int) | Norm. Factor |
|---|---|---|---|---|---|---|
| 1 | N/A | N/A | 1,507,800 | 138,787,152 | 137,279,352 | N/A |
| 2 | 27,142,080 | 58,550,688 | 35,168,784 | 187,128,312 | 151,959,528 | N/A |
| 3 | 36,335,712 | 44,006,256 | 40,204,584 | 180,965,400 | 140,760,816 | N/A |
| 4 | 104,165,040 | 112,990,752 | 111,963,936 | 253,812,720 | 141,848,784 | N/A |
| 5 | 6,639,696 | 12,717,264 | 10,703,616 | 151,595,640 | 140,892,024 | N/A |
| 6 | 6,804,336 | 13,813,464 | 22,984,416 | 185,072,328 | 162,087,912 | N/A |
| 7 | 6,875,400 | 14,118,552 | 10,852,296 | 151,504,920 | 140,652,624 | N/A |
| 8 | 45,965,304 | 60,649,680 | 60,612,720 | 227,901,576 | 167,288,856 | N/A |
| 9 | 8,779,680 | 16,384,032 | 16,441,152 | 159,750,192 | 143,309,040 | N/A |
| 10 | N/A | N/A | 834,960 | 139,817,328 | 138,982,368 | N/A |

#### Lane And Band Analysis

##### Lane 1

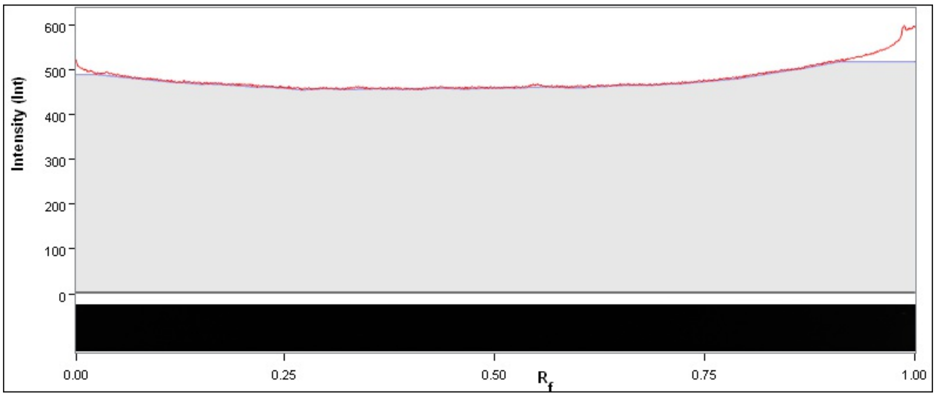

| Band No. | Band Label | Mol. Wt. (KDa) | Relative Front | Adj. Volume (Int) | Volume (Int) | Abs. Quant. | Rel. Quant. | Band % | Lane % |
|---|---|---|---|---|---|---|---|---|---|

| | |
|---|---|
| Band Detection | Automatically detected bands with custom sensitivity: 77 |
| Lane Background | Lane background subtracted with disk size: 17 |
| Lane Width | 5.91 mm |
| Regression Equation | A single equation is not available for this method |

##### Lane 2 - SeeBlue2

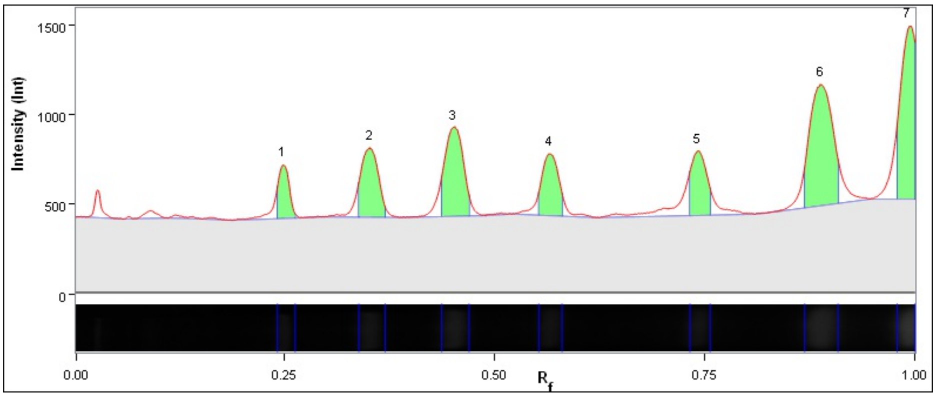

| Band No. | Band Label | Mol. Wt. (KDa) | Relative Front | Adj. Volume (Int) | Volume (Int) | Abs. Quant. | Rel. Quant. | Band % | Lane % |
|---|---|---|---|---|---|---|---|---|---|
| 1 | | 62.0 | 0.248 | 1,475,040 | 4,613,784 | N/A | N/A | 5.4 | 4.2 |
| 2 | | 49.0 | 0.350 | 3,070,032 | 7,757,568 | N/A | N/A | 11.3 | 8.7 |
| 3 | | 38.0 | 0.451 | 4,248,216 | 9,176,328 | N/A | N/A | 15.7 | 12.1 |
| 4 | | 28.0 | 0.565 | 2,612,568 | 6,856,584 | N/A | N/A | 9.6 | 7.4 |
| 5 | | 14.0 | 0.742 | 2,499,336 | 6,216,168 | N/A | N/A | 9.2 | 7.1 |
| 6 | | 6.0 | 0.888 | 7,245,168 | 14,093,016 | N/A | N/A | 26.7 | 20.6 |
| 7 | | 3.0 | 0.994 | 5,991,720 | 9,837,240 | N/A | N/A | 22.1 | 17.0 |

| | |
|---|---|
| Band Detection | Automatically detected bands with custom sensitivity: 77 |
| Lane Background | Lane background subtracted with disk size: 17 |
| Lane Width | 5.91 mm |
| Regression Equation | A single equation is not available for this method |

#### Lane 3

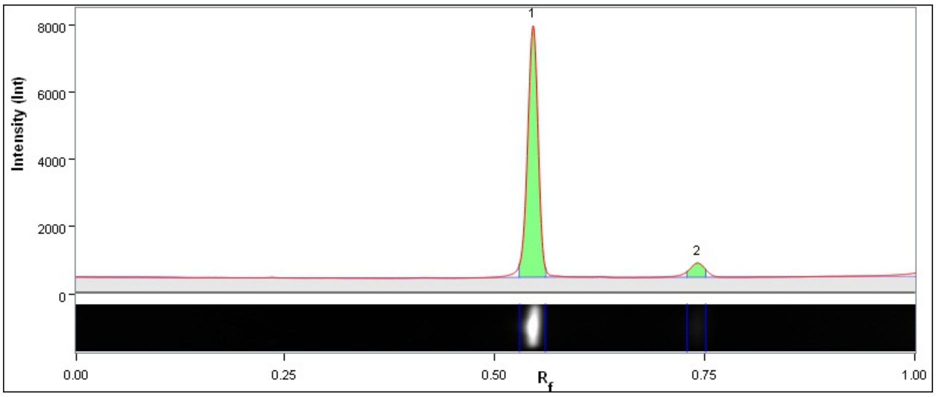

| Band No. | Band Label | Mol. Wt. (KDa) | Relative Front | Adj. Volume (Int) | Volume (Int) | Abs. Quant. | Rel. Quant. | Band % | Lane % |
|---|---|---|---|---|---|---|---|---|---|
| 1 | | 29.5 | 0.545 | 33,958,008 | 38,393,040 | N/A | N/A | 93.5 | 84.5 |
| 2 | | 14.0 | 0.741 | 2,377,704 | 5,613,216 | N/A | N/A | 6.5 | 5.9 |

| | |
|---|---|
| Band Detection | Automatically detected bands with custom sensitivity: 77 |
| Lane Background | Lane background subtracted with disk size: 17 |
| Lane Width | 5.91 mm |
| Regression Equation | A single equation is not available for this method |

#### Lane 4

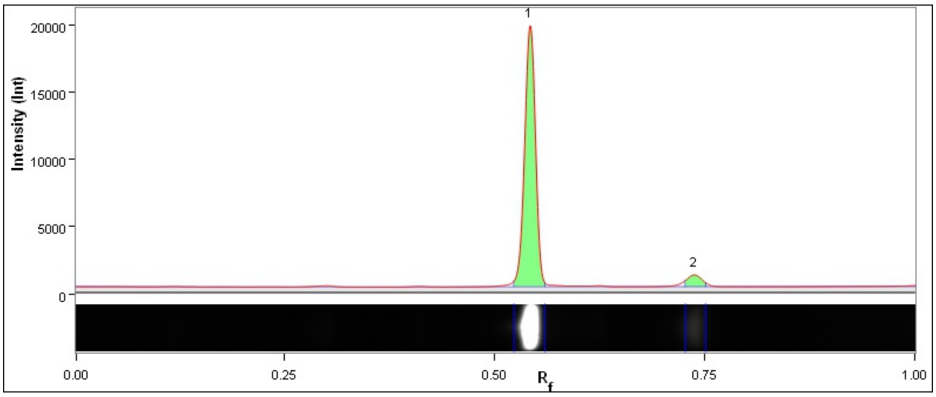

| Band No. | Band Label | Mol. Wt. (KDa) | Relative Front | Adj. Volume (Int) | Volume (Int) | Abs. Quant. | Rel. Quant. | Band % | Lane % |
|---|---|---|---|---|---|---|---|---|---|
| 1 | | 29.8 | 0.542 | 98,621,712 | 103,882,968 | N/A | N/A | 94.7 | 88.1 |
| 2 | | 14.2 | 0.738 | 5,543,328 | 9,107,784 | N/A | N/A | 5.3 | 5.0 |

| | |
|---|---|
| Band Detection | Automatically detected bands with custom sensitivity: 77 |
| Lane Background | Lane background subtracted with disk size: 17 |
| Lane Width | 5.91 mm |
| Regression Equation | A single equation is not available for this method |

#### Lane 5

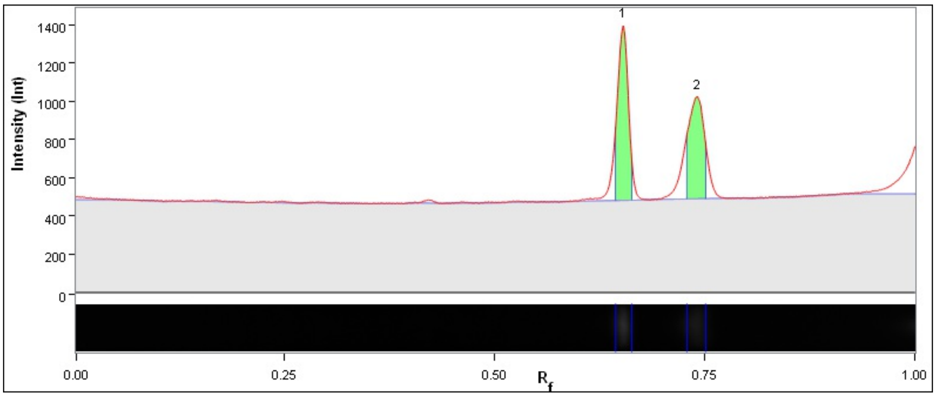

| Band No. | Band Label | Mol. Wt. (KDa) | Relative Front | Adj. Volume (Int) | Volume (Int) | Abs. Quant. | Rel. Quant. | Band % | Lane % |
|---|---|---|---|---|---|---|---|---|---|
| 1 | | 19.9 | 0.653 | 3,664,752 | 6,464,472 | N/A | N/A | 55.2 | 34.2 |
| 2 | | 14.1 | 0.741 | 2,974,944 | 6,252,792 | N/A | N/A | 44.8 | 27.8 |

| | |
|---|---|
| Band Detection | Automatically detected bands with custom sensitivity: 77 |
| Lane Background | Lane background subtracted with disk size: 17 |
| Lane Width | 5.91 mm |
| Regression Equation | A single equation is not available for this method |

4

#### Lane 6

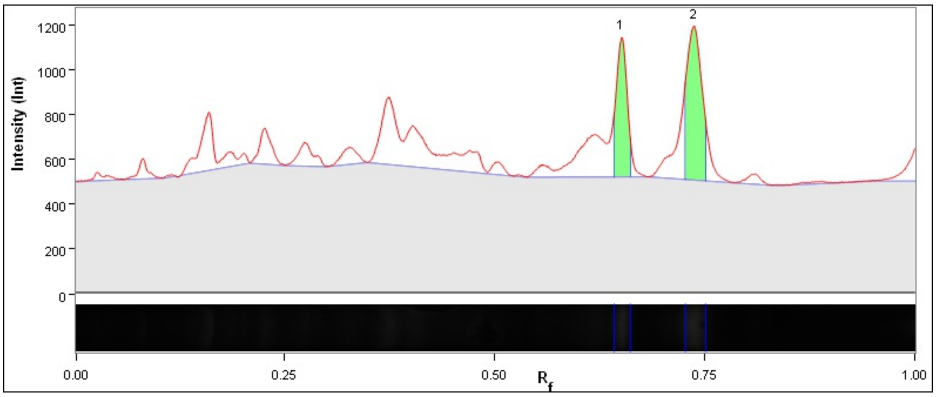

| Band No. | Band Label | Mol. Wt. (KDa) | Relative Front | Adj. Volume (Int) | Volume (Int) | Abs. Quant. | Rel. Quant. | Band % | Lane % |
|---|---|---|---|---|---|---|---|---|---|
| 1 | | 20.0 | 0.651 | 2,718,744 | 5,894,784 | N/A | N/A | 40.0 | 11.8 |
| 2 | | 14.3 | 0.736 | 4,085,592 | 7,918,680 | N/A | N/A | 60.0 | 17.8 |

| | |
|---|---|
| Band Detection | Automatically detected bands with custom sensitivity: 77 |
| Lane Background | Lane background subtracted with disk size: 17 |
| Lane Width | 5.91 mm |
| Regression Equation | A single equation is not available for this method |

#### Lane 7

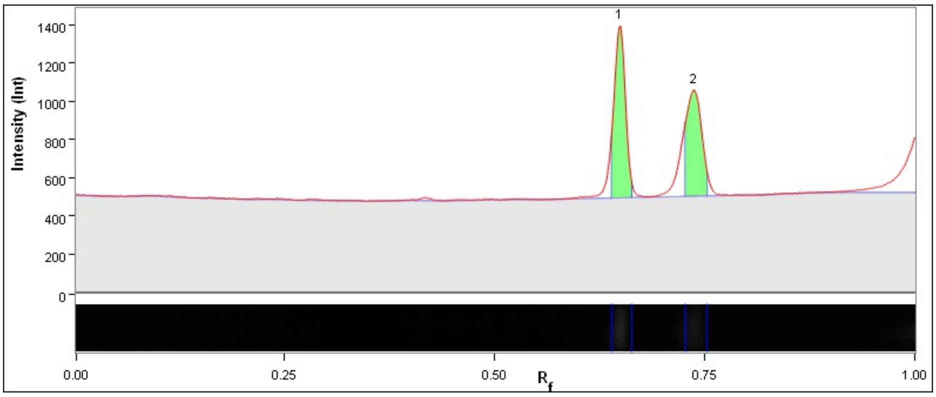

| Band No. | Band Label | Mol. Wt. (KDa) | Relative Front | Adj. Volume (Int) | Volume (Int) | Abs. Quant. | Rel. Quant. | Band % | Lane % |
|---|---|---|---|---|---|---|---|---|---|
| 1 | | 20.2 | 0.648 | 3,799,320 | 7,262,472 | N/A | N/A | 55.3 | 35.0 |
| 2 | | 14.3 | 0.736 | 3,076,080 | 6,856,080 | N/A | N/A | 44.7 | 28.3 |

| | |
|---|---|
| Band Detection | Automatically detected bands with custom sensitivity: 77 |
| Lane Background | Lane background subtracted with disk size: 17 |
| Lane Width | 5.91 mm |
| Regression Equation | A single equation is not available for this method |

#### Lane 8

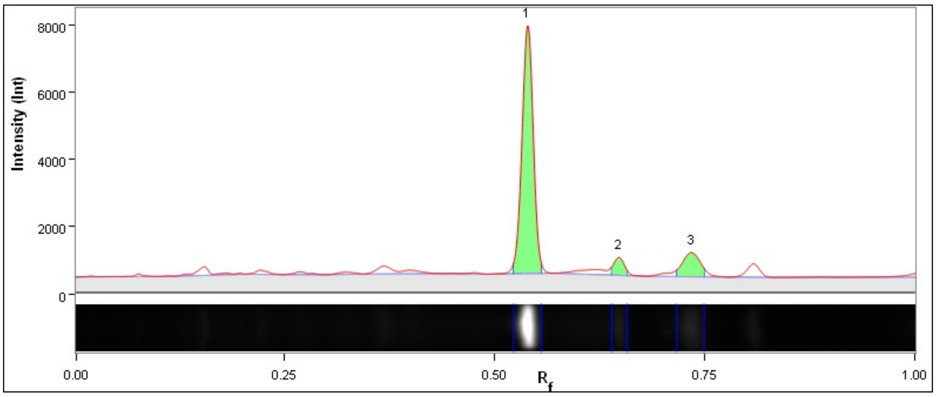

| Band No. | Band Label | Mol. Wt. (KDa) | Relative Front | Adj. Volume (Int) | Volume (Int) | Abs. Quant. | Rel. Quant. | Band % | Lane % |
|---|---|---|---|---|---|---|---|---|---|
| 1 | | 30.1 | 0.539 | 37,993,704 | 44,271,696 | N/A | N/A | 82.7 | 62.7 |
| 2 | | 20.2 | 0.648 | 2,445,912 | 5,696,544 | N/A | N/A | 5.3 | 4.0 |
| 3 | | 14.4 | 0.734 | 5,525,688 | 10,681,440 | N/A | N/A | 12.0 | 9.1 |

| | |
|---|---|
| Band Detection | Automatically detected bands with custom sensitivity: 77 |
| Lane Background | Lane background subtracted with disk size: 17 |
| Lane Width | 5.91 mm |
| Regression Equation | A single equation is not available for this method |

#### Lane 9

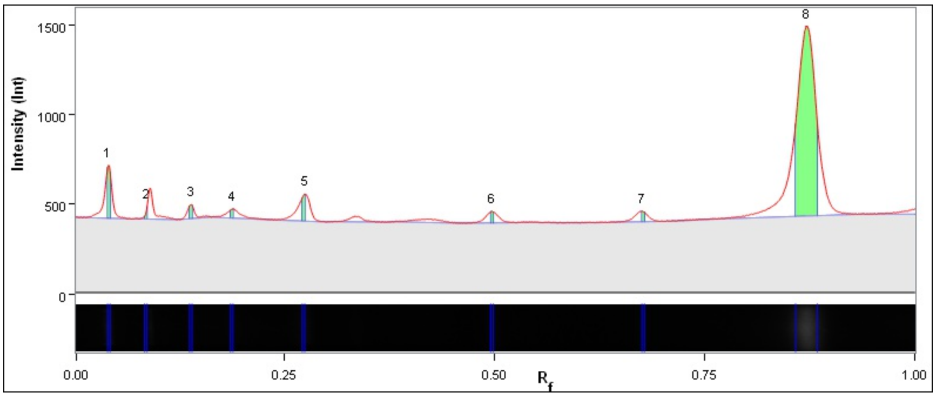

| Band No. | Band Label | Mol. Wt. (KDa) | Relative Front | Adj. Volume (Int) | Volume (Int) | Abs. Quant. | Rel. Quant. | Band % | Lane % |
|---|---|---|---|---|---|---|---|---|---|
| 1 | | 62.0 | 0.040 | 350,280 | 878,304 | N/A | N/A | 4.0 | 2.1 |
| 2 | | 62.0 | 0.085 | 60,144 | 583,296 | N/A | N/A | 0.7 | 0.4 |
| 3 | | 62.0 | 0.138 | 95,928 | 621,600 | N/A | N/A | 1.1 | 0.6 |
| 4 | | 62.0 | 0.188 | 63,336 | 592,872 | N/A | N/A | 0.7 | 0.4 |
| 5 | | 58.5 | 0.273 | 184,632 | 692,664 | N/A | N/A | 2.1 | 1.1 |
| 6 | | 33.7 | 0.496 | 79,800 | 574,728 | N/A | N/A | 0.9 | 0.5 |
| 7 | | 18.2 | 0.675 | 69,720 | 573,720 | N/A | N/A | 0.8 | 0.4 |
| 8 | | 6.6 | 0.871 | 7,875,840 | 11,866,848 | N/A | N/A | 89.7 | 47.9 |

| | |
|---|---|
| Band Detection | Automatically detected bands with custom sensitivity: 77 |
| Lane Background | Lane background subtracted with disk size: 17 |

| | |
|---|---|
| Lane Width | 5.91 mm |
| Regression Equation | A single equation is not available for this method |

##### Lane 10

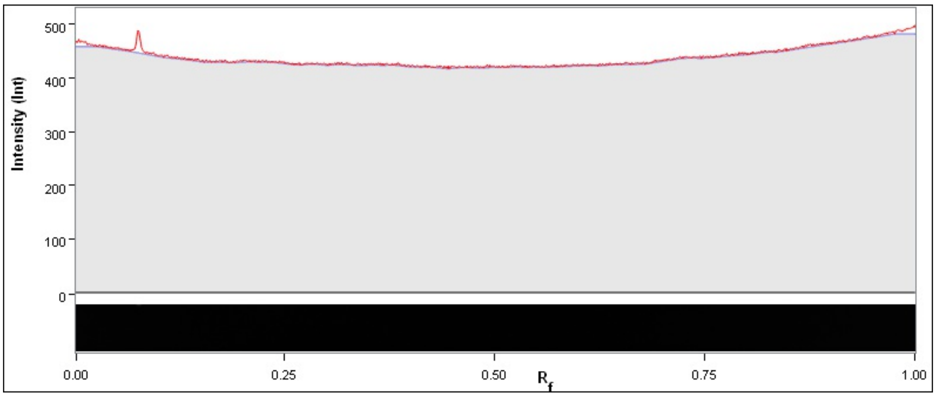

| Band No. | Band Label | Mol. Wt. (KDa) | Relative Front | Adj. Volume (Int) | Volume (Int) | Abs. Quant. | Rel. Quant. | Band % | Lane % |
|---|---|---|---|---|---|---|---|---|---|

| | |
|---|---|
| Band Detection | Automatically detected bands with custom sensitivity: 77 |
| Lane Background | Lane background subtracted with disk size: 17 |
| Lane Width | 5.91 mm |
| Regression Equation | A single equation is not available for this method |

### Image Report: admin 2023-10-11 13h49m54s

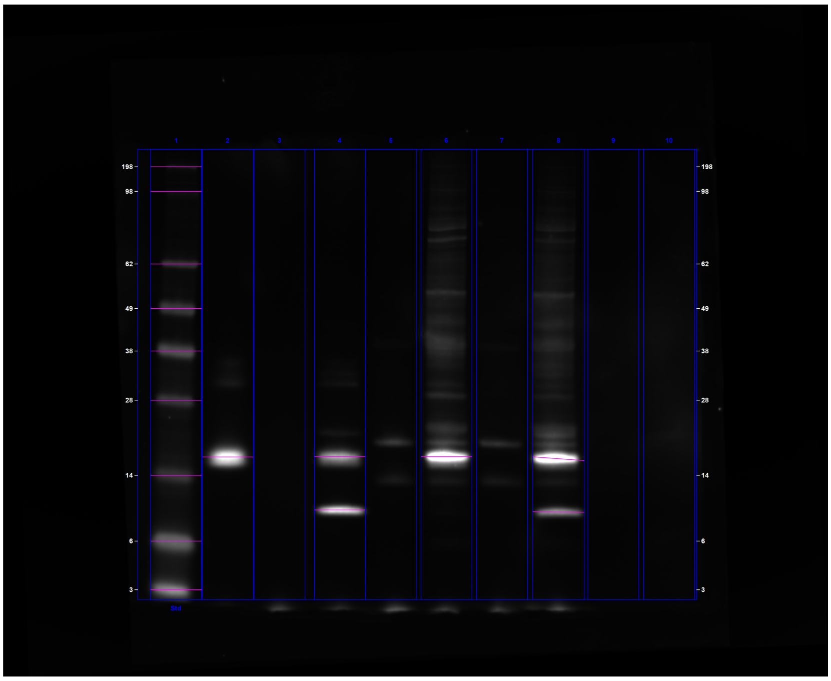

**C:\Users\oded\OneDrive - University Of Cambridge\Vendruscolo\Results\Western\ChemiDoc Images 2023-10-11_19.07.33\admin 2023-10-11 13h49m54s.scn**

---

#### Acquisition Information

| | |
|---|---|
| Imager | ChemiDoc™ MP |
| Exposure Time (sec) | 0.500 (Manual) |
| Flat Field | Applied (Red Epi) |
| Serial Number | 734BR-3787 |
| Software Version | 2.4.0.03 |
| Application | Alexa 647 |
| Excitation Source | Red Epi Illumination |
| Emission Filter | 700/50 Filter |

#### Image Information

| | |
|---|---|
| Acquisition Date | 11/10/2023 2:49:54 PM |
| User Name | admin |
| Image Area (mm) | X: 111.8 Y: 91.0 |
| Pixel Size (µm) | X: 39.4 Y: 39.4 |
| Data Range (Int) | 0 - 46274 |

#### Analysis Settings

| | |
|---|---|
| Detection | Lane detection:<br>Manually created lanes |

| | |
|---|---|
| | Band detection:<br><br>Manually adjusted bands<br><br>Lane Background Subtraction:<br>Lane background subtracted with disk size: 17<br><br>Lane width: Variable |
| Mol. Weight Analysis | Standard: SeeBlue2-full<br>Standard lanes: first<br>Regression method: Point to Point (semi-log) |

#### Lane Statistics

| Lane No. | Adj. Total Band Vol. (Int) | Total Band Vol. (Int) | Adj. Total Lane Vol. (Int) | Total Lane Vol. (Int) | Bkgd. Vol. (Int) | Norm. Factor |
|---|---|---|---|---|---|---|
| 1 | 259,340,900 | 394,815,750 | 312,136,300 | 723,759,225 | 411,622,925 | N/A |
| 2 | 157,019,800 | 176,464,750 | 184,947,525 | 454,559,000 | 269,611,475 | N/A |
| 3 | N/A | N/A | 22,922,900 | 275,679,950 | 252,757,050 | N/A |
| 4 | 183,521,975 | 210,867,125 | 214,878,475 | 480,766,300 | 265,887,825 | N/A |
| 5 | N/A | N/A | 36,054,200 | 305,802,350 | 269,748,150 | N/A |
| 6 | 198,652,475 | 220,619,875 | 334,509,875 | 761,133,275 | 426,623,400 | N/A |
| 7 | N/A | N/A | 33,145,875 | 329,332,325 | 296,186,450 | N/A |
| 8 | 248,770,137 | 287,914,926 | 350,548,146 | 814,955,790 | 464,407,644 | N/A |
| 9 | N/A | N/A | 21,283,325 | 304,686,550 | 283,403,225 | N/A |
| 10 | N/A | N/A | 40,710,075 | 373,178,050 | 332,467,975 | N/A |

#### Lane And Band Analysis

##### Lane 1 - SeeBlue2-full

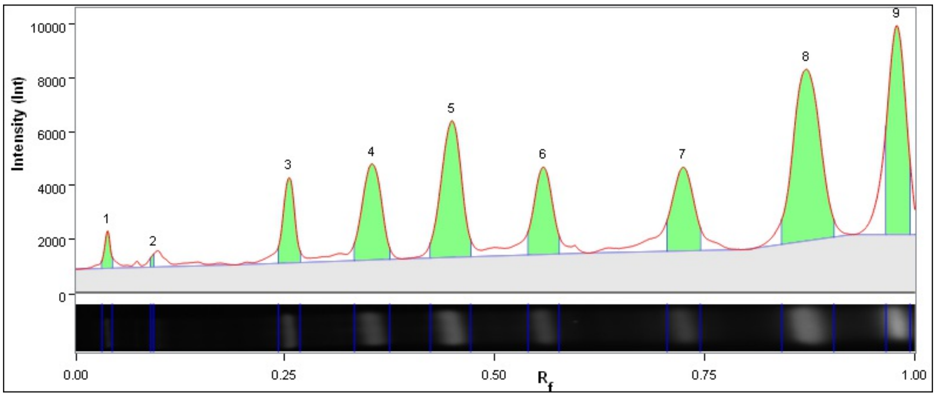

| Band No. | Band Label | Mol. Wt. (KDa) | Relative Front | Adj. Volume (Int) | Volume (Int) | Abs. Quant. | Rel. Quant. | Band % | Lane % |
|---|---|---|---|---|---|---|---|---|---|
| 1 | | 198.0 | 0.039 | 3,266,550 | 6,695,500 | N/A | N/A | 1.3 | 1.0 |
| 2 | | 98.0 | 0.094 | 506,800 | 1,588,475 | N/A | N/A | 0.2 | 0.2 |
| 3 | | 62.0 | 0.255 | 14,820,750 | 23,421,650 | N/A | N/A | 5.7 | 4.7 |
| 4 | | 49.0 | 0.354 | 27,891,150 | 43,129,275 | N/A | N/A | 10.8 | 8.9 |
| 5 | | 38.0 | 0.448 | 41,808,200 | 60,506,425 | N/A | N/A | 16.1 | 13.4 |
| 6 | | 28.0 | 0.558 | 23,892,750 | 39,825,625 | N/A | N/A | 9.2 | 7.7 |
| 7 | | 14.0 | 0.724 | 25,997,475 | 44,299,850 | N/A | N/A | 10.0 | 8.3 |
| 8 | | 6.0 | 0.871 | 69,950,125 | 105,619,850 | N/A | N/A | 27.0 | 22.4 |
| 9 | | 3.0 | 0.979 | 51,207,100 | 69,729,100 | N/A | N/A | 19.7 | 16.4 |

| | |
|---|---|
| Lane Background | Lane background subtracted with disk size: 17 |
| Lane Width | 6.90 mm |
| Regression Equation | A single equation is not available for this method |

##### Lane 2

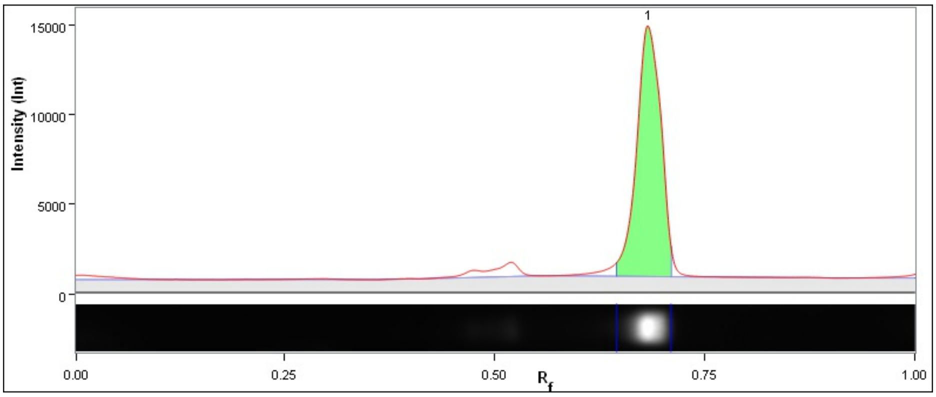

| Band No. | Band Label | Mol. Wt. (KDa) | Relative Front | Adj. Volume (Int) | Volume (Int) | Abs. Quant. | Rel. Quant. | Band % | Lane % |
|---|---|---|---|---|---|---|---|---|---|
| 1 | | 16.6 | 0.684 | 157,019,800 | 176,464,750 | N/A | N/A | 100.0 | 84.9 |

| | |
|---|---|
| Lane Background | Lane background subtracted with disk size: 17 |
| Lane Width | 6.90 mm |
| Regression Equation | A single equation is not available for this method |

##### Lane 3

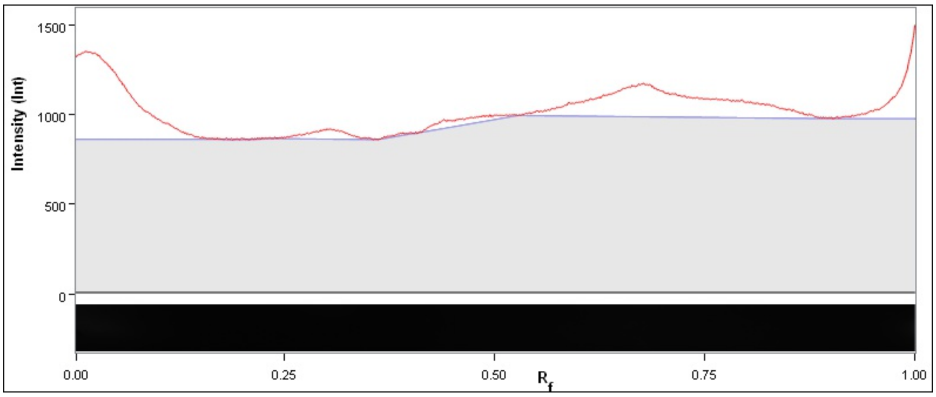

| Band No. | Band Label | Mol. Wt. (KDa) | Relative Front | Adj. Volume (Int) | Volume (Int) | Abs. Quant. | Rel. Quant. | Band % | Lane % |
|---|---|---|---|---|---|---|---|---|---|

| | |
|---|---|
| Lane Background | Lane background subtracted with disk size: 17 |
| Lane Width | 6.90 mm |
| Regression Equation | A single equation is not available for this method |

#### Lane 4

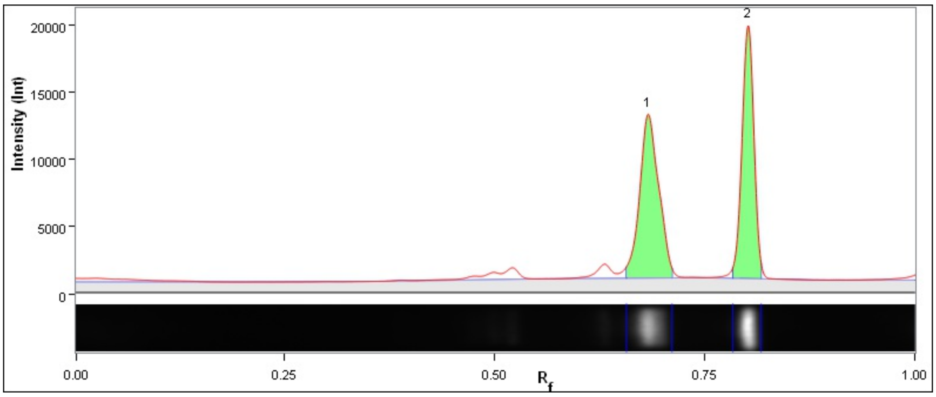

| Band No. | Band Label | Mol. Wt. (KDa) | Relative Front | Adj. Volume (Int) | Volume (Int) | Abs. Quant. | Rel. Quant. | Band % | Lane % |
|---|---|---|---|---|---|---|---|---|---|
| 1 | | 16.6 | 0.683 | 93,530,150 | 110,710,600 | N/A | N/A | 51.0 | 43.5 |
| 2 | | 9.0 | 0.801 | 89,991,825 | 100,156,525 | N/A | N/A | 49.0 | 41.9 |

| | |
|---|---|
| Lane Background | Lane background subtracted with disk size: 17 |
| Lane Width | 6.90 mm |
| Regression Equation | A single equation is not available for this method |

#### Lane 5

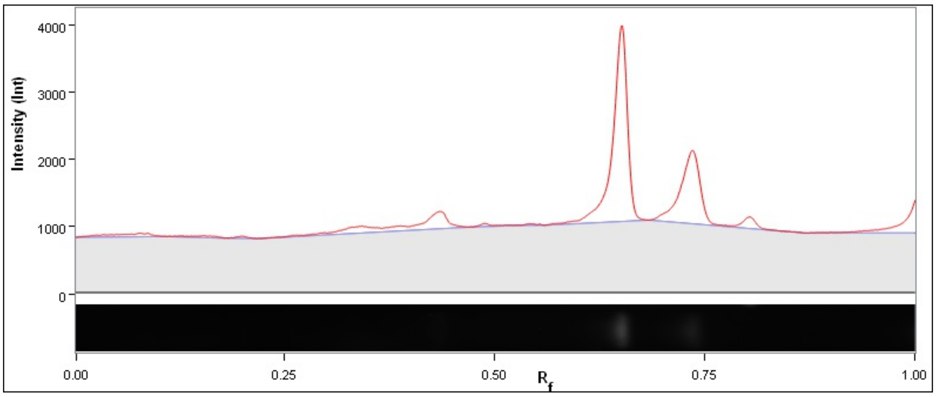

| Band No. | Band Label | Mol. Wt. (KDa) | Relative Front | Adj. Volume (Int) | Volume (Int) | Abs. Quant. | Rel. Quant. | Band % | Lane % |
|---|---|---|---|---|---|---|---|---|---|

| | |
|---|---|
| Lane Background | Lane background subtracted with disk size: 17 |
| Lane Width | 6.90 mm |
| Regression Equation | A single equation is not available for this method |

#### Lane 6

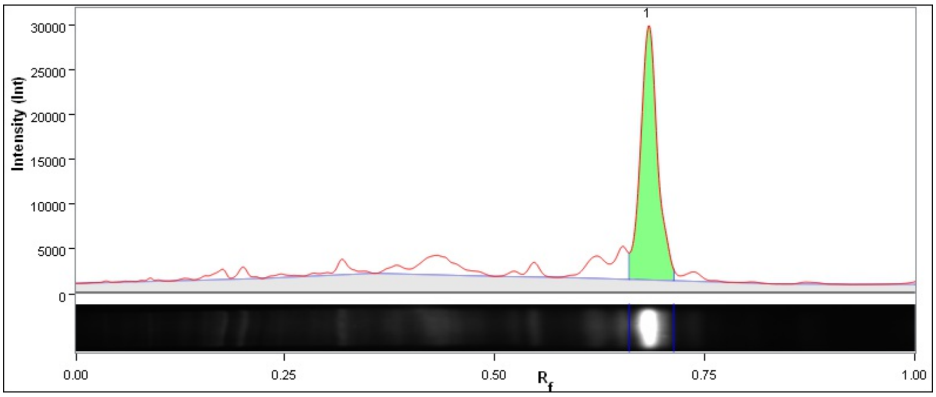

| Band No. | Band Label | Mol. Wt. (KDa) | Relative Front | Adj. Volume (Int) | Volume (Int) | Abs. Quant. | Rel. Quant. | Band % | Lane % |
|---|---|---|---|---|---|---|---|---|---|
| 1 | | 16.6 | 0.683 | 198,652,475 | 220,619,875 | N/A | N/A | 100.0 | 59.4 |

| | |
|---|---|
| Lane Background | Lane background subtracted with disk size: 17 |
| Lane Width | 6.90 mm |
| Regression Equation | A single equation is not available for this method |

**Lane 7**

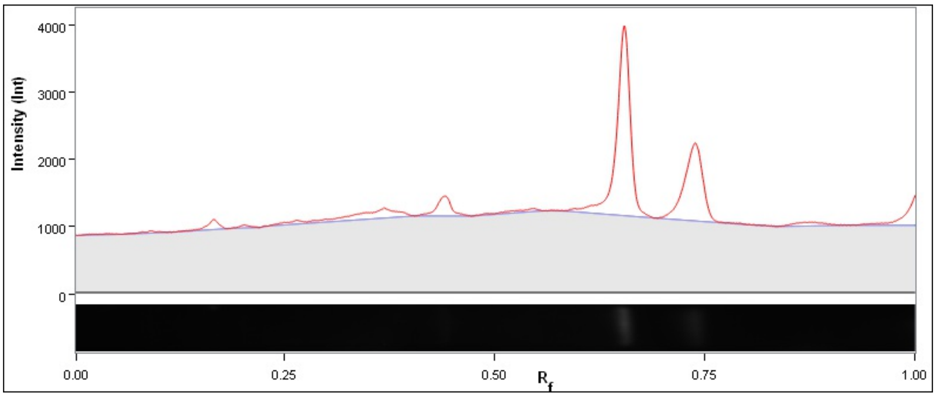

| Band No. | Band Label | Mol. Wt. (KDa) | Relative Front | Adj. Volume (Int) | Volume (Int) | Abs. Quant. | Rel. Quant. | Band % | Lane % |
|---|---|---|---|---|---|---|---|---|---|

| | |
|---|---|
| Lane Background | Lane background subtracted with disk size: 17 |
| Lane Width | 6.90 mm |
| Regression Equation | A single equation is not available for this method |

**Lane 8**

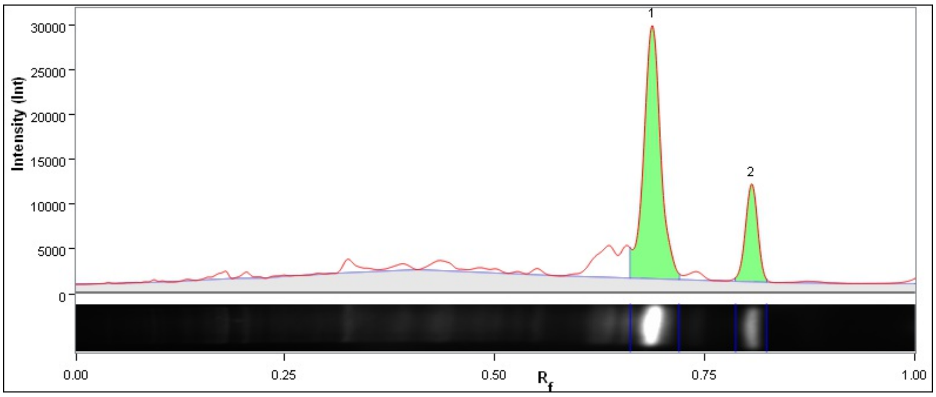

| Band No. | Band Label | Mol. Wt. (KDa) | Relative Front | Adj. Volume (Int) | Volume (Int) | Abs. Quant. | Rel. Quant. | Band % | Lane % |
|---|---|---|---|---|---|---|---|---|---|
| 1 | | 16.3 | 0.688 | 194,559,285 | 220,717,584 | N/A | N/A | 78.2 | 55.5 |
| 2 | | 8.7 | 0.806 | 54,210,852 | 67,197,342 | N/A | N/A | 21.8 | 15.5 |

| | |
|---|---|
| Lane Background | Lane background subtracted with disk size: 17 |
| Lane Width | 6.98 mm |
| Regression Equation | A single equation is not available for this method |

#### Lane 9

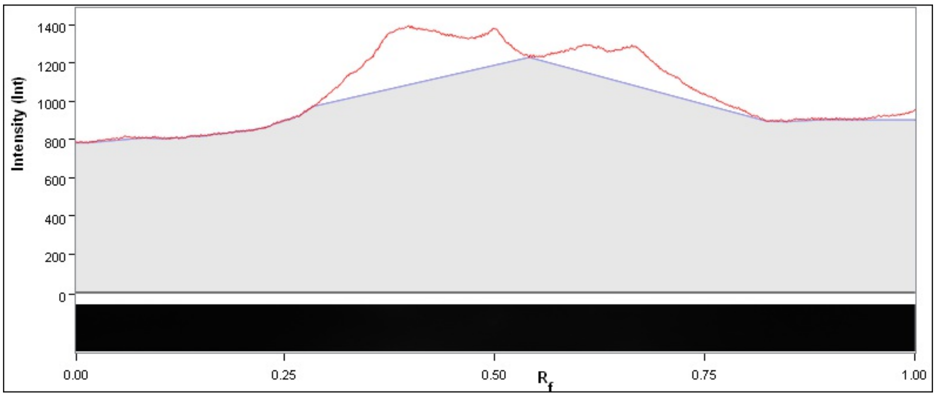

| Band No. | Band Label | Mol. Wt. (KDa) | Relative Front | Adj. Volume (Int) | Volume (Int) | Abs. Quant. | Rel. Quant. | Band % | Lane % |
|---|---|---|---|---|---|---|---|---|---|

| | |
|---|---|
| Lane Background | Lane background subtracted with disk size: 17 |
| Lane Width | 6.90 mm |
| Regression Equation | A single equation is not available for this method |

#### Lane 10

6

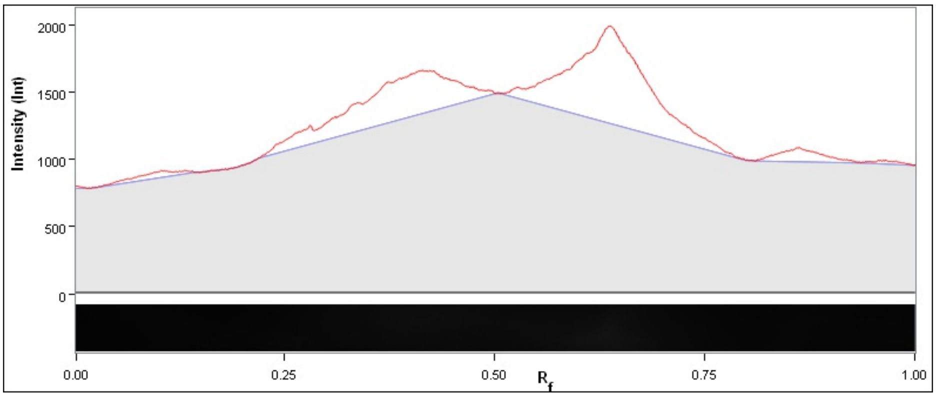

| Band No. | Band Label | Mol. Wt. (KDa) | Relative Front | Adj. Volume (Int) | Volume (Int) | Abs. Quant. | Rel. Quant. | Band % | Lane % |
|---|---|---|---|---|---|---|---|---|---|

| | |
|---|---|
| Lane Background | Lane background subtracted with disk size: 17 |
| Lane Width | 6.90 mm |
| Regression Equation | A single equation is not available for this method |

### Image Report: admin 2023-10-12 14h11m42s

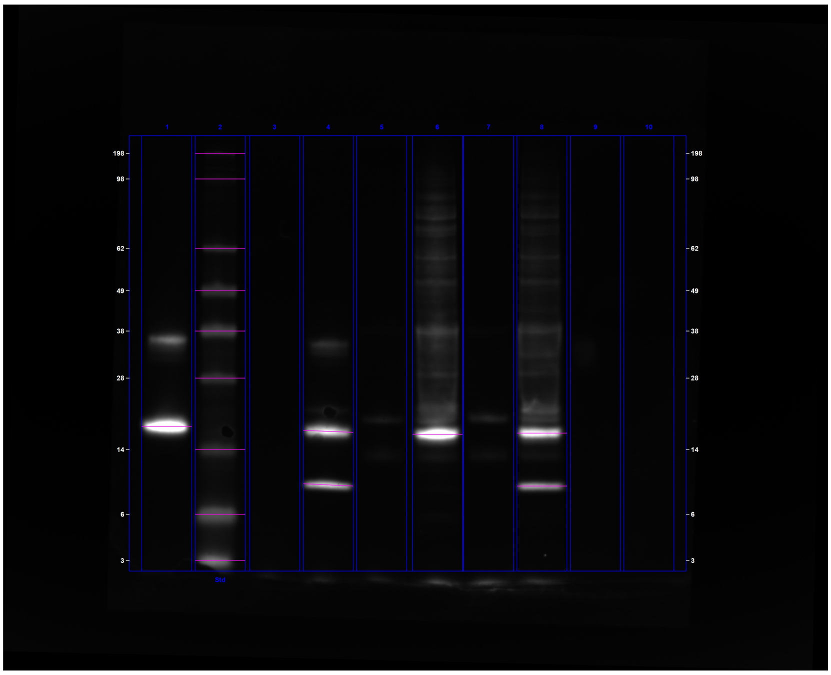

**C:\Users\oded\OneDrive - University Of Cambridge\Vendruscolo\Results\Western\ChemiDoc Images 2023-10-12_19.02.38\admin 2023-10-12 14h11m42s.scn**

---

#### Acquisition Information

| | |
|---|---|
| Imager | ChemiDoc™ MP |
| Exposure Time (sec) | 0.200 (Manual) |
| Flat Field | Applied (Red Epi) |
| Serial Number | 734BR-3787 |
| Software Version | 2.4.0.03 |
| Application | Alexa 647 |
| Excitation Source | Red Epi Illumination |
| Emission Filter | 700/50 Filter |

#### Image Information

| | |
|---|---|
| Acquisition Date | 12/10/2023 3:11:42 PM |
| User Name | admin |
| Image Area (mm) | X: 115.0 Y: 92.9 |
| Pixel Size (µm) | X: 41.0 Y: 41.0 |
| Data Range (Int) | 0 - 24086 |

#### Analysis Settings

| | |
|---|---|
| Detection | Lane detection:<br>Manually created lanes |

| | |
|---|---|
| | Band detection:<br><br>Manually adjusted bands<br><br>Lane Background Subtraction:<br>Lane background subtracted with disk size: 17<br><br>Lane width: 6.98 mm |
| Mol. Weight Analysis | Standard: SeeBlue2-full<br>Standard lanes: 2<br>Regression method: Point to Point (semi-log) |

#### Lane Statistics

| Lane No. | Adj. Total Band Vol. (Int) | Total Band Vol. (Int) | Adj. Total Lane Vol. (Int) | Total Lane Vol. (Int) | Bkgd. Vol. (Int) | Norm. Factor |
|---|---|---|---|---|---|---|
| 1 | 109,744,520 | 121,707,590 | 146,390,910 | 325,867,730 | 179,476,820 | N/A |
| 2 | 94,940,240 | 173,478,880 | 107,879,790 | 338,760,360 | 230,880,570 | N/A |
| 3 | N/A | N/A | 2,556,290 | 171,606,330 | 169,050,040 | N/A |
| 4 | 96,699,060 | 116,670,150 | 116,051,520 | 293,162,790 | 177,111,270 | N/A |
| 5 | N/A | N/A | 8,463,280 | 181,067,170 | 172,603,890 | N/A |
| 6 | 87,943,720 | 104,739,890 | 168,372,420 | 436,918,700 | 268,546,280 | N/A |
| 7 | N/A | N/A | 11,267,600 | 185,231,660 | 173,964,060 | N/A |
| 8 | 93,459,710 | 117,892,620 | 157,031,550 | 407,621,920 | 250,590,370 | N/A |
| 9 | N/A | N/A | 4,614,310 | 171,628,770 | 167,014,460 | N/A |
| 10 | N/A | N/A | 1,434,970 | 168,972,860 | 167,537,890 | N/A |

#### Lane And Band Analysis

##### Lane 1

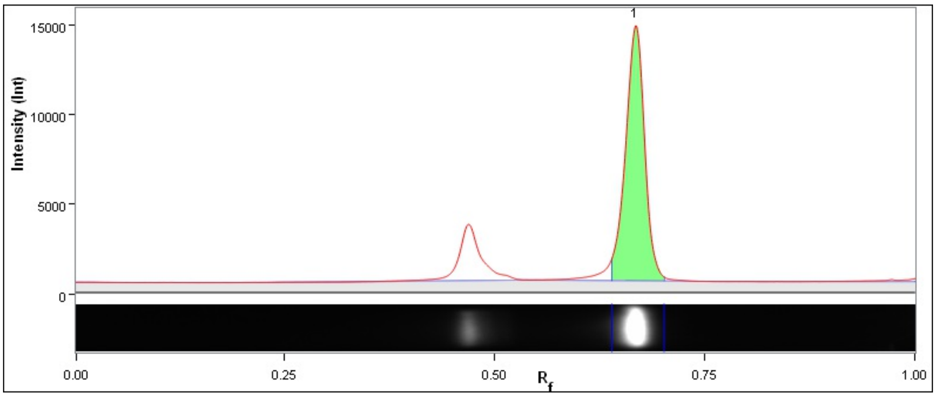

| Band No. | Band Label | Mol. Wt. (KDa) | Relative Front | Adj. Volume (Int) | Volume (Int) | Abs. Quant. | Rel. Quant. | Band % | Lane % |
|---|---|---|---|---|---|---|---|---|---|
| 1 | | 17.5 | 0.668 | 109,744,520 | 121,707,590 | N/A | N/A | 100.0 | 75.0 |

| | |
|---|---|
| Lane Background | Lane background subtracted with disk size: 17 |
| Lane Width | 6.98 mm |
| Regression Equation | A single equation is not available for this method |

##### Lane 2 - SeeBlue2-full

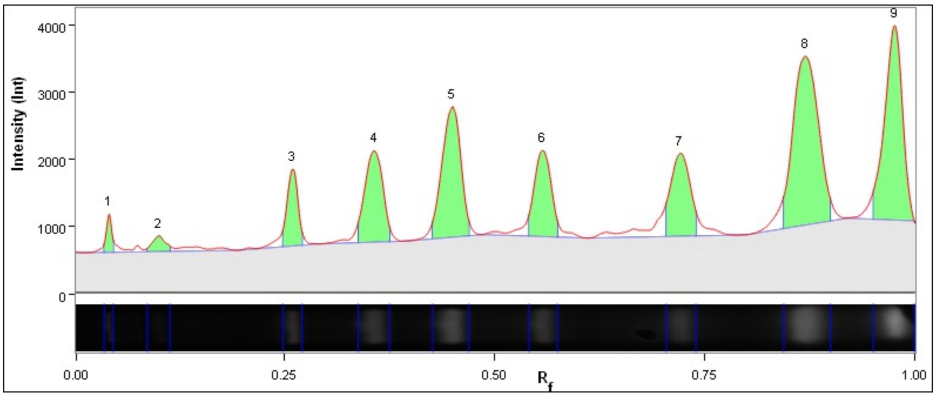

| Band No. | Band Label | Mol. Wt. (KDa) | Relative Front | Adj. Volume (Int) | Volume (Int) | Abs. Quant. | Rel. Quant. | Band % | Lane % |
|---|---|---|---|---|---|---|---|---|---|
| 1 | | 198.0 | 0.041 | 1,177,420 | 3,151,970 | N/A | N/A | 1.2 | 1.1 |
| 2 | | 98.0 | 0.099 | 1,203,090 | 6,060,500 | N/A | N/A | 1.3 | 1.1 |
| 3 | | 62.0 | 0.259 | 4,884,950 | 9,598,540 | N/A | N/A | 5.1 | 4.5 |
| 4 | | 49.0 | 0.356 | 9,675,720 | 17,831,130 | N/A | N/A | 10.2 | 9.0 |
| 5 | | 38.0 | 0.449 | 14,461,900 | 24,838,530 | N/A | N/A | 15.2 | 13.4 |
| 6 | | 28.0 | 0.557 | 8,450,530 | 16,701,140 | N/A | N/A | 8.9 | 7.8 |
| 7 | | 14.0 | 0.721 | 9,223,520 | 17,859,010 | N/A | N/A | 9.7 | 8.5 |
| 8 | | 6.0 | 0.870 | 25,322,520 | 41,586,930 | N/A | N/A | 26.7 | 23.5 |
| 9 | | 3.0 | 0.976 | 20,540,590 | 35,851,130 | N/A | N/A | 21.6 | 19.0 |

| | |
|---|---|
| Lane Background | Lane background subtracted with disk size: 17 |
| Lane Width | 6.98 mm |
| Regression Equation | A single equation is not available for this method |

#### Lane 3

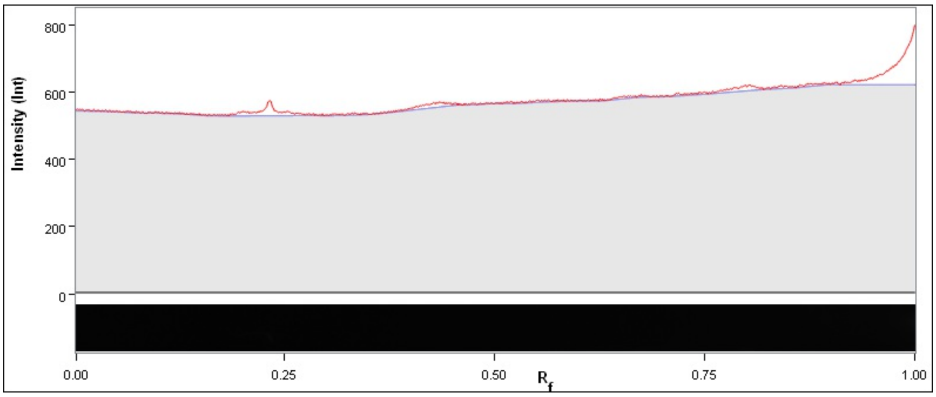

| Band No. | Band Label | Mol. Wt. (KDa) | Relative Front | Adj. Volume (Int) | Volume (Int) | Abs. Quant. | Rel. Quant. | Band % | Lane % |
|---|---|---|---|---|---|---|---|---|---|

| | |
|---|---|
| Lane Background | Lane background subtracted with disk size: 17 |
| Lane Width | 6.98 mm |
| Regression Equation | A single equation is not available for this method |

**Lane 4**

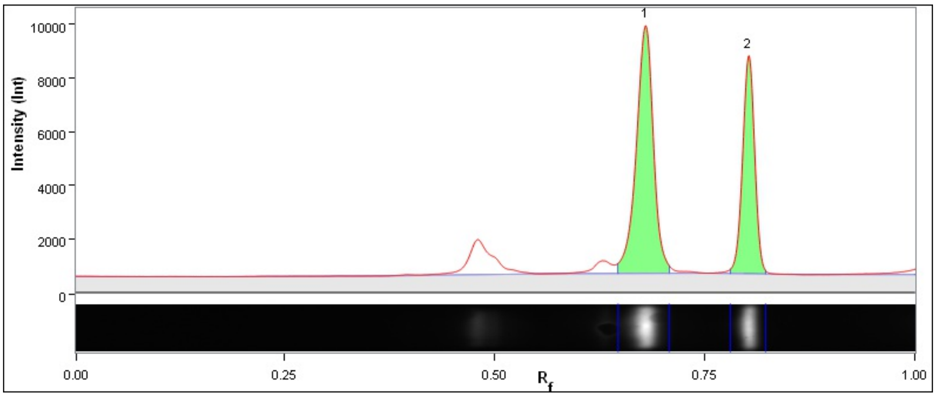

| Band No. | Band Label | Mol. Wt. (KDa) | Relative Front | Adj. Volume (Int) | Volume (Int) | Abs. Quant. | Rel. Quant. | Band % | Lane % |
|---|---|---|---|---|---|---|---|---|---|
| 1 | | 16.7 | 0.679 | 59,712,500 | 71,603,660 | N/A | N/A | 61.8 | 51.5 |
| 2 | | 8.8 | 0.802 | 36,986,560 | 45,066,490 | N/A | N/A | 38.2 | 31.9 |

| | |
|---|---|
| Lane Background | Lane background subtracted with disk size: 17 |
| Lane Width | 6.98 mm |
| Regression Equation | A single equation is not available for this method |

**Lane 5**

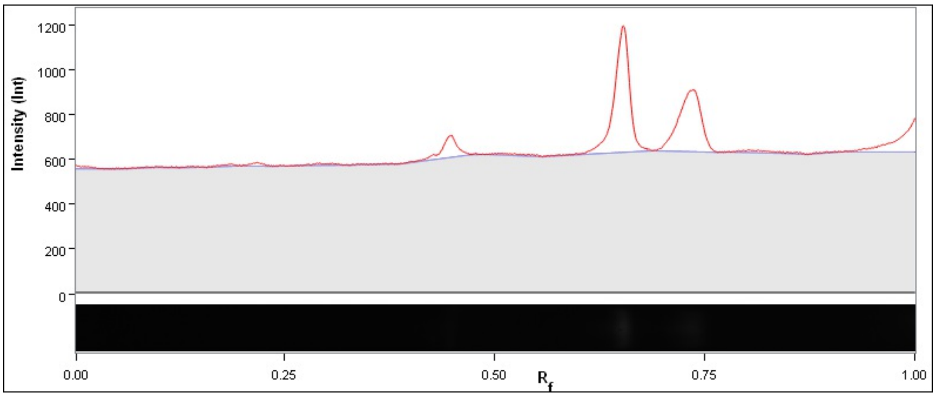

| Band No. | Band Label | Mol. Wt. (KDa) | Relative Front | Adj. Volume (Int) | Volume (Int) | Abs. Quant. | Rel. Quant. | Band % | Lane % |
|---|---|---|---|---|---|---|---|---|---|

| | |
|---|---|
| Lane Background | Lane background subtracted with disk size: 17 |
| Lane Width | 6.98 mm |
| Regression Equation | A single equation is not available for this method |

**Lane 6**

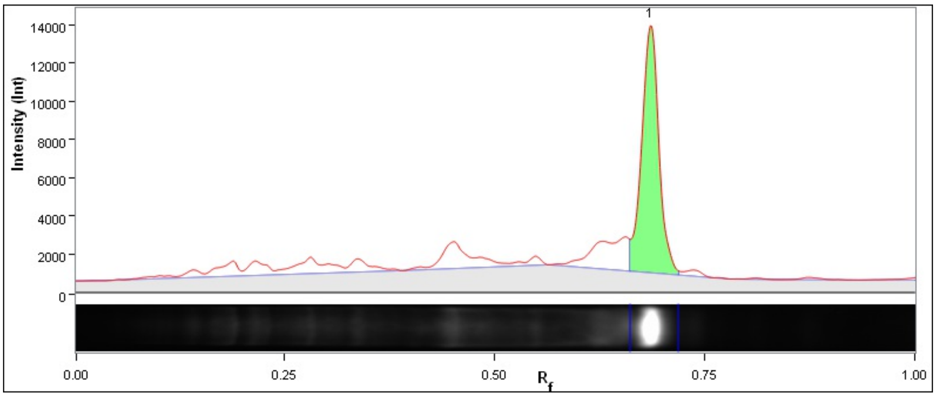

| Band No. | Band Label | Mol. Wt. (KDa) | Relative Front | Adj. Volume (Int) | Volume (Int) | Abs. Quant. | Rel. Quant. | Band % | Lane % |
|---|---|---|---|---|---|---|---|---|---|
| 1 | | 16.2 | 0.686 | 87,943,720 | 104,739,890 | N/A | N/A | 100.0 | 52.2 |

| | |
|---|---|
| Lane Background | Lane background subtracted with disk size: 17 |
| Lane Width | 6.98 mm |
| Regression Equation | A single equation is not available for this method |

##### Lane 7

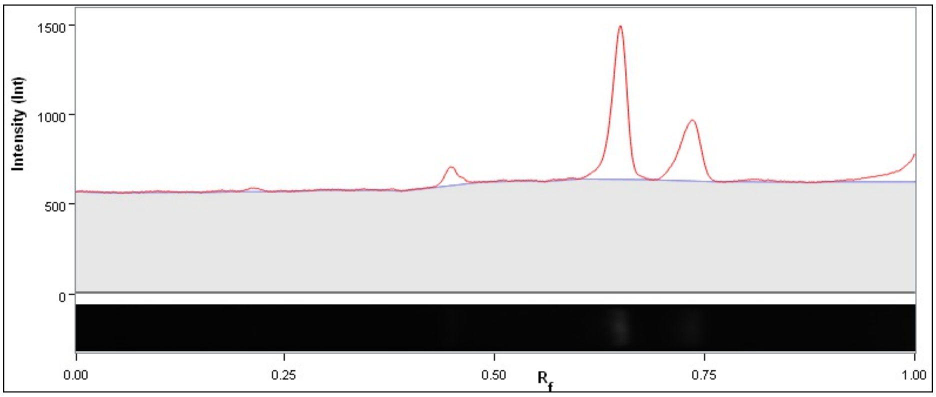

| Band No. | Band Label | Mol. Wt. (KDa) | Relative Front | Adj. Volume (Int) | Volume (Int) | Abs. Quant. | Rel. Quant. | Band % | Lane % |
|---|---|---|---|---|---|---|---|---|---|

| | |
|---|---|
| Lane Background | Lane background subtracted with disk size: 17 |
| Lane Width | 6.98 mm |
| Regression Equation | A single equation is not available for this method |

##### Lane 8

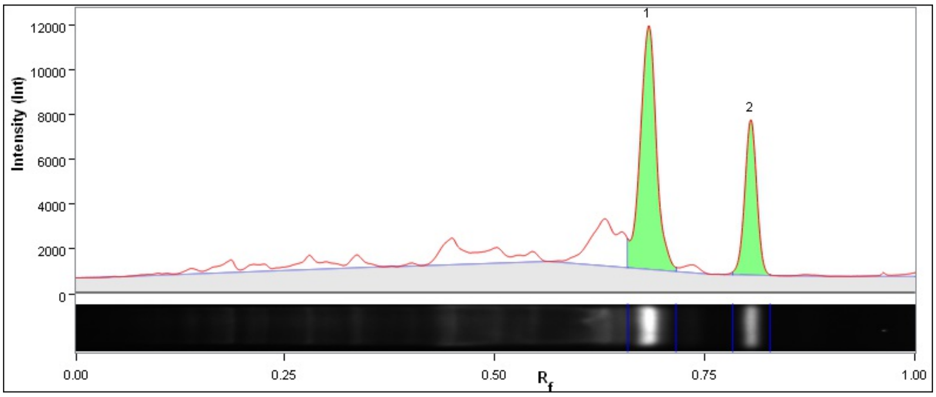

| Band No. | Band Label | Mol. Wt. (KDa) | Relative Front | Adj. Volume (Int) | Volume (Int) | Abs. Quant. | Rel. Quant. | Band % | Lane % |
|---|---|---|---|---|---|---|---|---|---|
| 1 | | 16.4 | 0.683 | 63,452,500 | 78,796,190 | N/A | N/A | 67.9 | 40.4 |
| 2 | | 8.7 | 0.805 | 30,007,210 | 39,096,430 | N/A | N/A | 32.1 | 19.1 |

| | |
|---|---|
| Lane Background | Lane background subtracted with disk size: 17 |
| Lane Width | 6.98 mm |
| Regression Equation | A single equation is not available for this method |

#### Lane 9

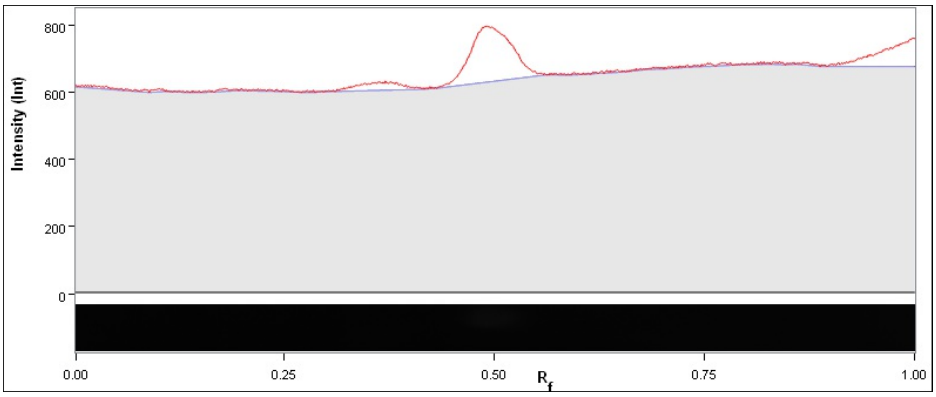

| Band No. | Band Label | Mol. Wt. (KDa) | Relative Front | Adj. Volume (Int) | Volume (Int) | Abs. Quant. | Rel. Quant. | Band % | Lane % |
|---|---|---|---|---|---|---|---|---|---|

| | |
|---|---|
| Lane Background | Lane background subtracted with disk size: 17 |
| Lane Width | 6.98 mm |
| Regression Equation | A single equation is not available for this method |

#### Lane 10

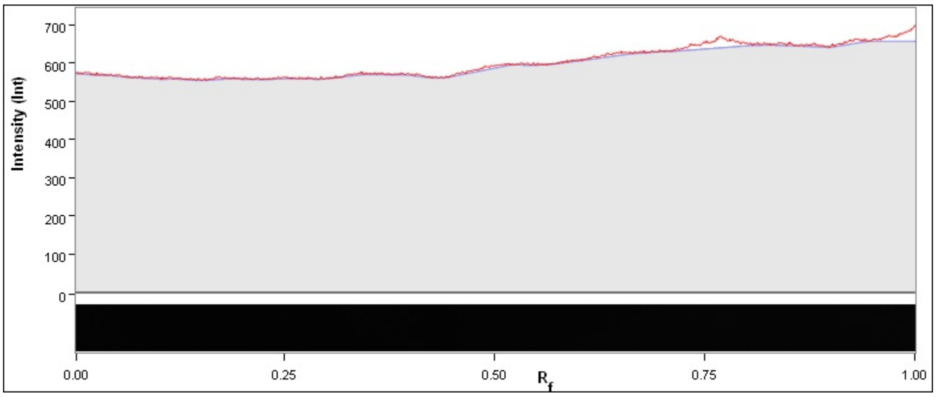

| Band No. | Band Label | Mol. Wt. (KDa) | Relative Front | Adj. Volume (Int) | Volume (Int) | Abs. Quant. | Rel. Quant. | Band % | Lane % |
|---|---|---|---|---|---|---|---|---|---|

| | |
|---|---|
| Lane Background | Lane background subtracted with disk size: 17 |
| Lane Width | 6.98 mm |
| Regression Equation | A single equation is not available for this method |

7

### Image Report: admin 2023-10-13 14h06m50s

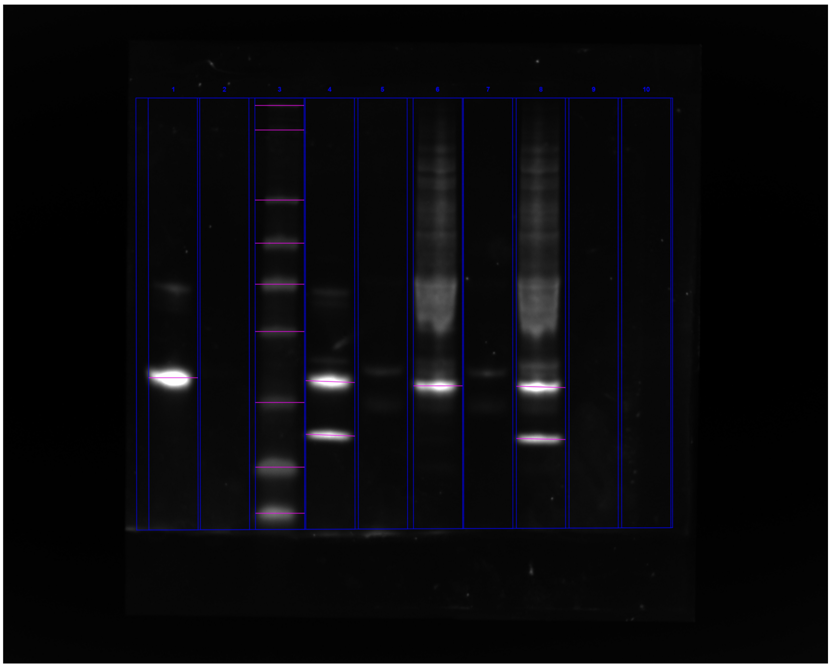

**C:\Users\oded\OneDrive - University Of Cambridge\Vendruscolo\Results\Western\ChemiDoc Images 2023-10-13_14.07.42\admin 2023-10-13 14h06m50s.scn**

---

#### Acquisition Information

| | |
|---|---|
| Imager | ChemiDoc™ MP |
| Exposure Time (sec) | 0.200 (Manual) |
| Flat Field | Applied (Red Epi) |
| Serial Number | 734BR-3787 |
| Software Version | 2.4.0.03 |
| Application | Alexa 647 |
| Excitation Source | Red Epi Illumination |
| Emission Filter | 700/50 Filter |

#### Image Information

| | |
|---|---|
| Acquisition Date | 13/10/2023 3:06:50 PM |
| User Name | admin |
| Image Area (mm) | X: 117.5 Y: 94.0 |
| Pixel Size (μm) | X: 42.7 Y: 42.7 |
| Data Range (Int) | 397 - 33545 |

#### Analysis Settings

| | |
|---|---|
| Detection | Lane detection:<br>Manually created lanes |

| | |
|---|---|
| | Band detection:<br>Automatically detected bands with sensitivity: Low<br>Manually adjusted bands<br><br>Lane Background Subtraction:<br>Lane background subtracted with disk size: 7<br><br>Lane width: 7.00 mm |

#### Lane Statistics

| Lane No. | Adj. Total Band Vol. (Int) | Total Band Vol. (Int) | Adj. Total Lane Vol. (Int) | Total Lane Vol. (Int) | Bkgd. Vol. (Int) | Norm. Factor |
|---|---|---|---|---|---|---|
| 1 | 134,328,956 | 152,699,416 | 154,314,324 | 384,231,664 | 229,917,340 | N/A |
| 2 | N/A | N/A | 4,811,924 | 227,122,616 | 222,310,692 | N/A |
| 3 | 110,721,156 | 250,304,508 | 115,003,524 | 395,984,560 | 280,981,036 | N/A |
| 4 | 126,761,176 | 155,714,720 | 141,992,020 | 374,855,456 | 232,863,436 | N/A |
| 5 | N/A | N/A | 13,780,592 | 233,733,292 | 219,952,700 | N/A |
| 6 | 70,694,168 | 87,575,180 | 185,675,552 | 519,711,736 | 334,036,184 | N/A |
| 7 | N/A | N/A | 13,248,248 | 225,282,864 | 212,034,616 | N/A |
| 8 | 119,098,276 | 153,574,356 | 184,316,812 | 599,866,244 | 415,549,432 | N/A |
| 9 | N/A | N/A | 4,797,164 | 232,455,568 | 227,658,404 | N/A |
| 10 | N/A | N/A | 4,521,480 | 224,646,052 | 220,124,572 | N/A |

#### Lane And Band Analysis

##### Lane 1

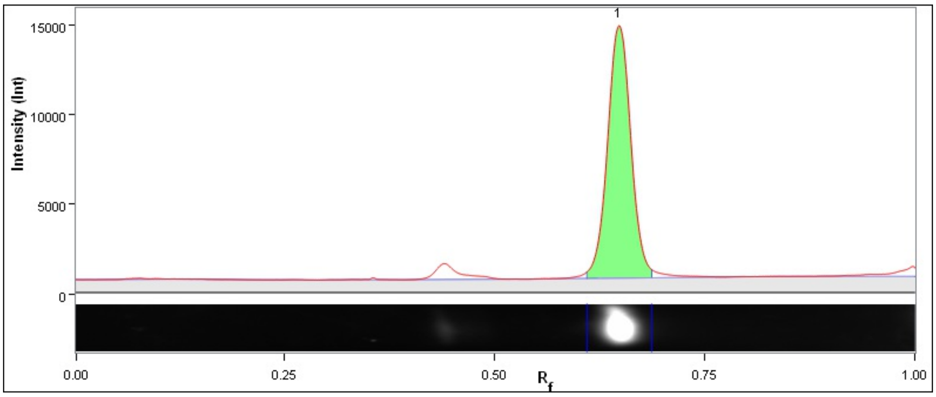

| Band No. | Band Label | Mol. Wt. (KDa) | Relative Front | Adj. Volume (Int) | Volume (Int) | Abs. Quant. | Rel. Quant. | Band % | Lane % |
|---|---|---|---|---|---|---|---|---|---|
| 1 | | N/A | 0.648 | 134,328,956 | 152,699,416 | N/A | N/A | 100.0 | 87.0 |

| | |
|---|---|
| Band Detection | Automatically detected bands with sensitivity: Low |
| Lane Background | Lane background subtracted with disk size: 7 |
| Lane Width | 7.00 mm |

##### Lane 2

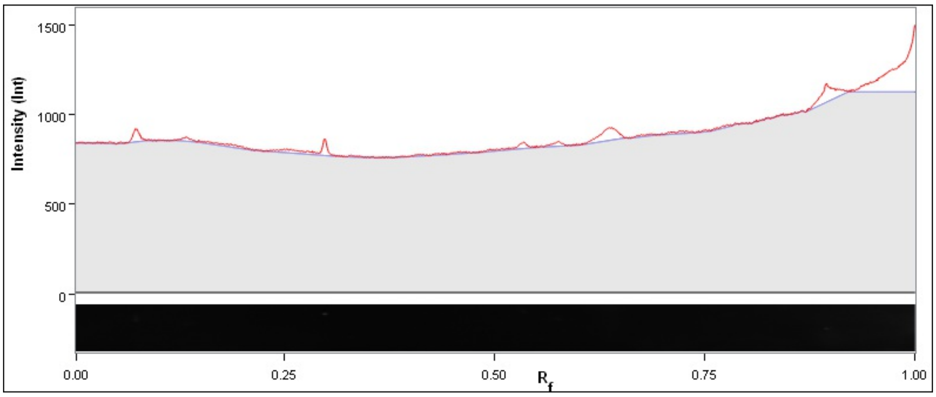

| Band No. | Band Label | Mol. Wt. (KDa) | Relative Front | Adj. Volume (Int) | Volume (Int) | Abs. Quant. | Rel. Quant. | Band % | Lane % |
|---|---|---|---|---|---|---|---|---|---|

| | |
|---|---|
| Band Detection | Automatically detected bands with sensitivity: Low |
| Lane Background | Lane background subtracted with disk size: 7 |
| Lane Width | 7.00 mm |

#### Lane 3

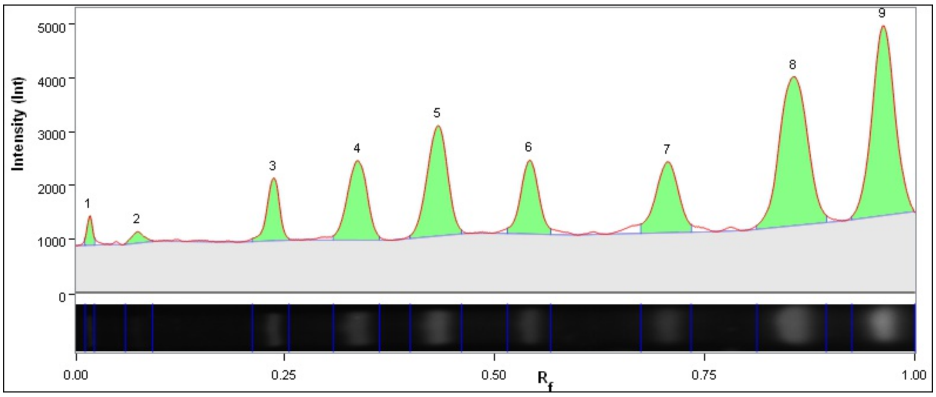

| Band No. | Band Label | Mol. Wt. (KDa) | Relative Front | Adj. Volume (Int) | Volume (Int) | Abs. Quant. | Rel. Quant. | Band % | Lane % |
|---|---|---|---|---|---|---|---|---|---|
| 1 | | N/A | 0.017 | 1,071,084 | 3,767,736 | N/A | N/A | 1.0 | 0.9 |
| 2 | | N/A | 0.074 | 968,584 | 8,950,792 | N/A | N/A | 0.9 | 0.8 |
| 3 | | N/A | 0.237 | 5,366,244 | 16,457,728 | N/A | N/A | 4.8 | 4.7 |
| 4 | | N/A | 0.337 | 10,614,408 | 24,685,280 | N/A | N/A | 9.6 | 9.2 |
| 5 | | N/A | 0.432 | 15,263,480 | 32,099,064 | N/A | N/A | 13.8 | 13.3 |
| 6 | | N/A | 0.541 | 8,885,848 | 23,614,688 | N/A | N/A | 8.0 | 7.7 |
| 7 | | N/A | 0.706 | 10,933,716 | 28,378,232 | N/A | N/A | 9.9 | 9.5 |
| 8 | | N/A | 0.856 | 28,416,608 | 55,255,044 | N/A | N/A | 25.7 | 24.7 |
| 9 | | N/A | 0.963 | 29,201,184 | 57,095,944 | N/A | N/A | 26.4 | 25.4 |

| | |
|---|---|
| Band Detection | Automatically detected bands with sensitivity: Low |
| Lane Background | Lane background subtracted with disk size: 7 |
| Lane Width | 7.00 mm |

**Lane 4**

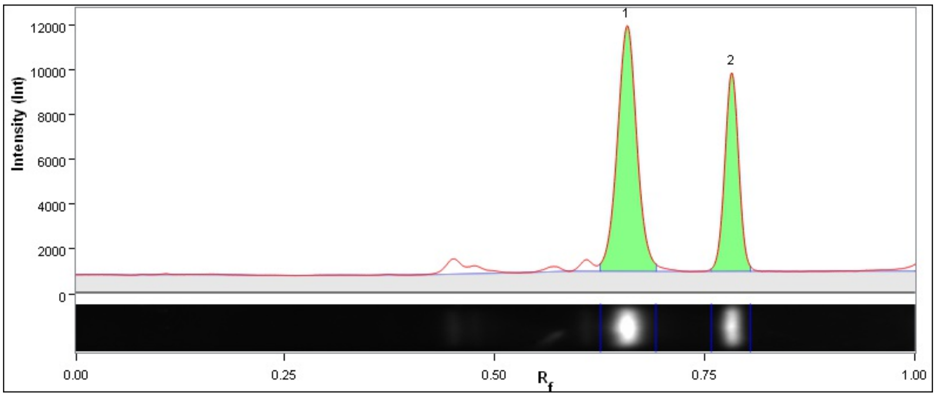

| Band No. | Band Label | Mol. Wt. (KDa) | Relative Front | Adj. Volume (Int) | Volume (Int) | Abs. Quant. | Rel. Quant. | Band % | Lane % |
|---|---|---|---|---|---|---|---|---|---|
| 1 | | N/A | 0.658 | 80,439,540 | 97,398,944 | N/A | N/A | 63.5 | 56.7 |
| 2 | | N/A | 0.782 | 46,321,636 | 58,315,776 | N/A | N/A | 36.5 | 32.6 |

| | |
|---|---|
| Band Detection | Automatically detected bands with sensitivity: Low |
| Lane Background | Lane background subtracted with disk size: 7 |
| Lane Width | 7.00 mm |

**Lane 5**

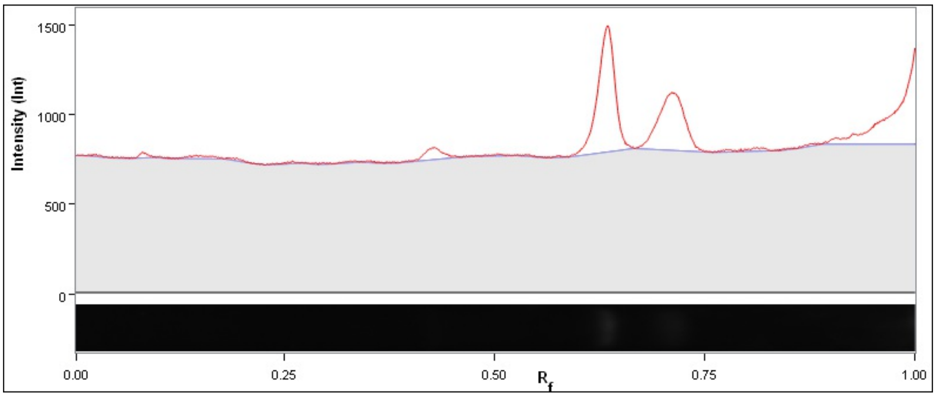

| Band No. | Band Label | Mol. Wt. (KDa) | Relative Front | Adj. Volume (Int) | Volume (Int) | Abs. Quant. | Rel. Quant. | Band % | Lane % |
|---|---|---|---|---|---|---|---|---|---|

| | |
|---|---|
| Band Detection | Automatically detected bands with sensitivity: Low |
| Lane Background | Lane background subtracted with disk size: 7 |
| Lane Width | 7.00 mm |

**Lane 6**

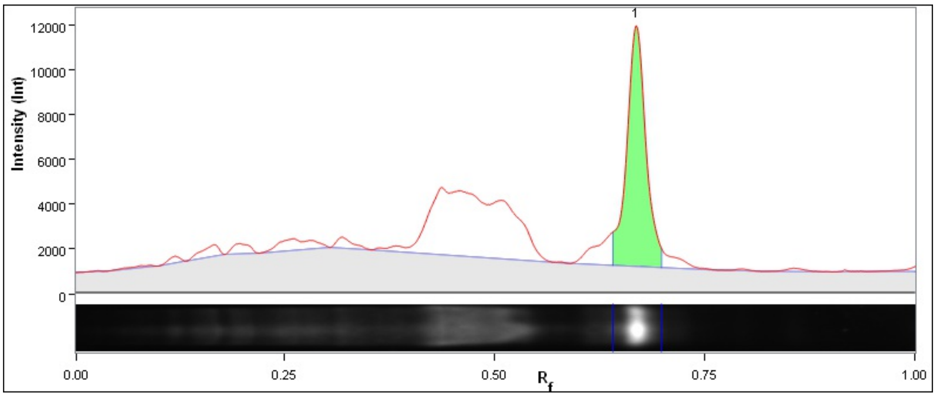

| Band No. | Band Label | Mol. Wt. (KDa) | Relative Front | Adj. Volume (Int) | Volume (Int) | Abs. Quant. | Rel. Quant. | Band % | Lane % |
|---|---|---|---|---|---|---|---|---|---|
| 1 | | N/A | 0.668 | 70,694,168 | 87,575,180 | N/A | N/A | 100.0 | 38.1 |

| | |
|---|---|
| Band Detection | Automatically detected bands with sensitivity: Low |
| Lane Background | Lane background subtracted with disk size: 7 |
| Lane Width | 7.00 mm |

**Lane 7**

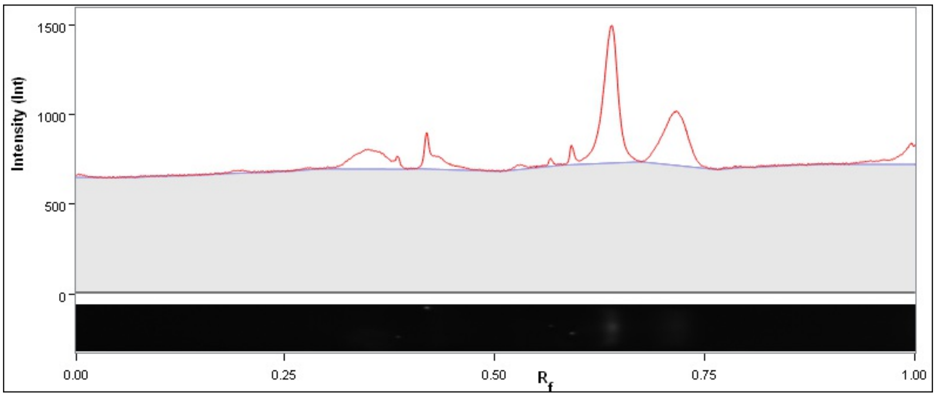

| Band No. | Band Label | Mol. Wt. (KDa) | Relative Front | Adj. Volume (Int) | Volume (Int) | Abs. Quant. | Rel. Quant. | Band % | Lane % |
|---|---|---|---|---|---|---|---|---|---|

| | |
|---|---|
| Band Detection | Automatically detected bands with sensitivity: Low |
| Lane Background | Lane background subtracted with disk size: 7 |
| Lane Width | 7.00 mm |

**Lane 8**

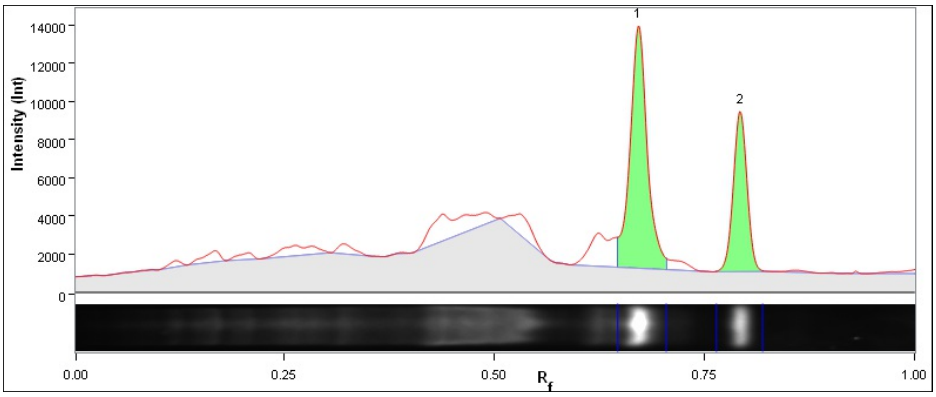

| Band No. | Band Label | Mol. Wt. (KDa) | Relative Front | Adj. Volume (Int) | Volume (Int) | Abs. Quant. | Rel. Quant. | Band % | Lane % |
|---|---|---|---|---|---|---|---|---|---|
| 1 | | N/A | 0.672 | 77,819,968 | 96,745,240 | N/A | N/A | 65.3 | 42.2 |
| 2 | | N/A | 0.793 | 41,278,308 | 56,829,116 | N/A | N/A | 34.7 | 22.4 |

| | |
|---|---|
| Band Detection | Automatically detected bands with sensitivity: Low |
| Lane Background | Lane background subtracted with disk size: 7 |
| Lane Width | 7.00 mm |

**Lane 9**

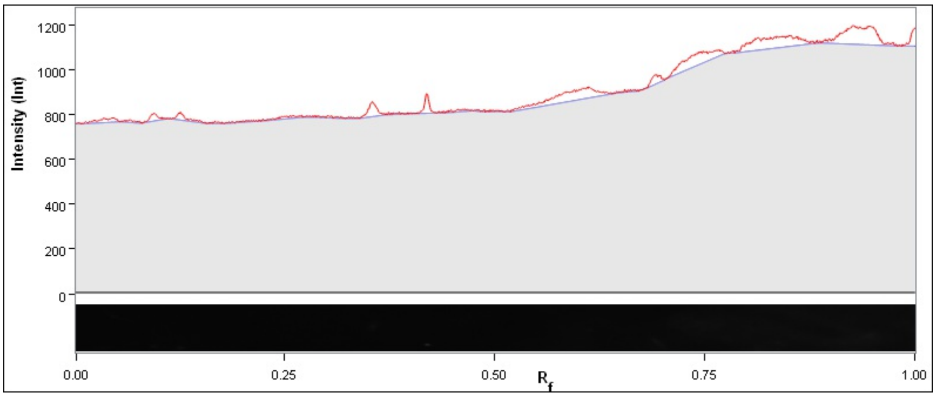

| Band No. | Band Label | Mol. Wt. (KDa) | Relative Front | Adj. Volume (Int) | Volume (Int) | Abs. Quant. | Rel. Quant. | Band % | Lane % |
|---|---|---|---|---|---|---|---|---|---|

| | |
|---|---|
| Band Detection | Automatically detected bands with sensitivity: Low |
| Lane Background | Lane background subtracted with disk size: 7 |
| Lane Width | 7.00 mm |

**Lane 10**

6

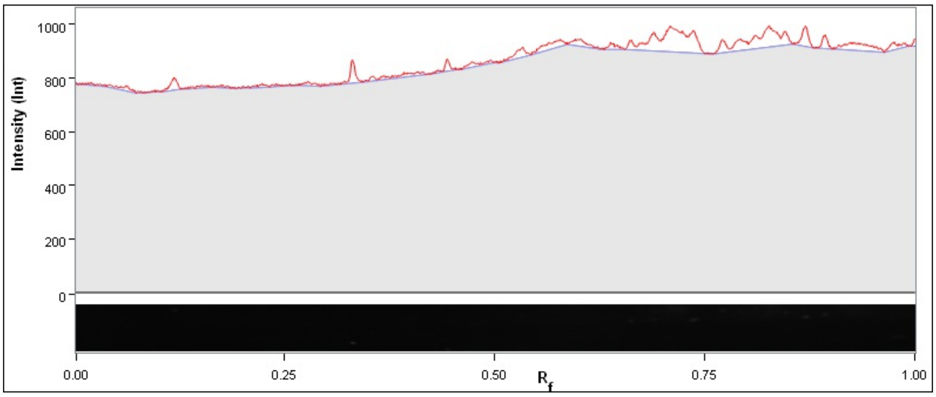

| Band No. | Band Label | Mol. Wt. (KDa) | Relative Front | Adj. Volume (Int) | Volume (Int) | Abs. Quant. | Rel. Quant. | Band % | Lane % |
|---|---|---|---|---|---|---|---|---|---|

| | |
|---|---|
| Band Detection | Automatically detected bands with sensitivity: Low |
| Lane Background | Lane background subtracted with disk size: 7 |
| Lane Width | 7.00 mm |

7

[9] A. Rosace, A. Bennett, M. Oeller, M. M. Mortensen, L. Sakhnini, N. Lorenzen, C. Poulsen, and P. Sormanni, “Automated optimisation of solubility and conforma-

tional stability of antibodies and proteins," *Nat. Commun.*, vol. 14, no. 1, p. 1937, 2023.

[15] M. Badertscher, P. Bühlmann, and E. Pretsch, *Structure Determination of Organic Compounds.* Springer-Verlag, 2009.

[16] M. T. Marty, A. J. Baldwin, E. G. Marklund, G. K. A. Hochberg, J. L. P. Benesch, and C. V. Robinson, "Bayesian deconvolution of mass and ion mobility spectra: From binary interactions to polydisperse ensembles," *Anal. Chem.*, vol. 87, no. 8, pp. 4370–4376, 2015.
